# Common Garlic (*Allium sativum* L.) has Potent Anti-*Bacillus anthracis* Activity

**DOI:** 10.1101/162214

**Authors:** Rajinder Kaur, Atul Tiwari, Manish Manish, Indresh K Maurya, Rakesh Bhatnagar, Samer Singh

**Affiliations:** Department of Microbial Biotechnology, Panjab University, Chandigarh-160014, India; School of Biotechnology, Jawaharlal Nehru University, New Delhi-110067, India; Centre of Experimental Medicine and Surgery, Institute of Medical Sciences, Banaras Hindu University, Varanasi-221005, India

**Keywords:** Anthrax, Garlic, Traditional medicine, Edible plants, *Bacillus anthracis*, *Allium sativum*

## Abstract

**Ethnopharmacological Relevance:** Gastrointestinal anthrax, a disease caused by *Bacillus anthracis*, remains an important but relatively neglected endemic disease of animals and humans in remote areas of the Indian subcontinent and some parts of Africa. Its initial symptoms include diarrhea and stomachache. In the current study, several common plants indicated for diarrhea, dysentery, stomachache or as stomachic as per traditional knowledge in the Indian subcontinent, *i.e.*, *Aegle marmelos* (L.) Correa (Bael), *Allium cepa* L. (Onion), *Allium sativum* L. (Garlic)*, Azadirachta indica* A. Juss. (Neem), *Berberis asiatica* Roxb. ex DC. (Daruharidra), *Coriandrum sativum* L. (Coriander), *Curcuma longa* L. (Turmeric), *Cynodon dactylon* (L.) Pers. (Bermuda grass), *Mangifera indica L.* (Mango), *Morus indica* L. (Black mulberry), *Ocimum tenuiflorum* L. *(Ocimum sanctum L.*, Holy Basil), *Ocimum gratissimum* L. (Ram Tulsi), *Psidium guajava* L. (Guava), *Zingiber officinale* Roscoe (Ginger), were evaluated for their anti-*Bacillus anthracis* property. The usage of *Azadirachta indica* A. Juss. and *Curcuma longa* L. by Santals (India), and *Allium sp.* by biblical people to alleviate anthrax-like symptoms is well documented, but the usage of other plants is traditionally only indicated for different gastrointestinal disturbances/conditions.

**Aim of the Study:** Evaluate the above listed commonly available edible plants from the Indian subcontinent that are used in the traditional medicine to treat gastrointestinal diseases including those also indicated for anthrax-like symptoms for the presence of potent anti-*B. anthracis* activity in a form amenable to use by the general population in the endemic areas.

**Materials and Methods:** Aqueous extracts made from fourteen plants indicated above were screened for their anti-*B. anthracis* activity using agar-well diffusion assay (AWDA) and broth microdilution methods. The Aqueous Garlic Extract (AGE) that displayed most potent anti-*B. anthracis* activity was assessed for its thermostability, stability under pH extremes encountered in the gastrointestinal tract, and potential antagonistic interaction with bile salts as well as the FDA-approved antibiotics used for anthrax control. The bioactive fractions from the AGE were isolated by TLC coupled bioautography followed by their characterization using GC-MS.

**Results:** Garlic (*Allium sativum* L.) extract was identified as the most promising candidate with bactericidal activity against *B. anthracis*. It consistently inhibited the growth of *B. anthracis* in AWDA and decreased the viable colony-forming unit counts in liquid-broth cultures by 6-logs within 6-12 h. The AGE displayed acceptable thermostability (>80% anti-*B. anthracis* activity retained on incubation at 50°C for 12 h) and stability in gastric pH range (2-8). It did not antagonize the activity of FDA-approved antibiotics used for anthrax control. GC-MS analysis of the TLC separated bioactive fractions of AGE indicated the presence of previously unreported constituents such as phthalic acid derivatives, acid esters, phenyl group-containing compounds, steroids *etc*.

**Conclusion:** The Aqueous Garlic Extract (AGE) displayed potent anti-*B. anthracis* activity. It was better than that displayed by *Azadirachta indica* A. Juss. (Neem) and *Mangifera indica* L. while *Curcuma longa* L. (Turmeric) did not show any activity under the assay conditions used. Further work should be undertaken to explore the possible application of AGE in preventing anthrax incidences in endemic areas.

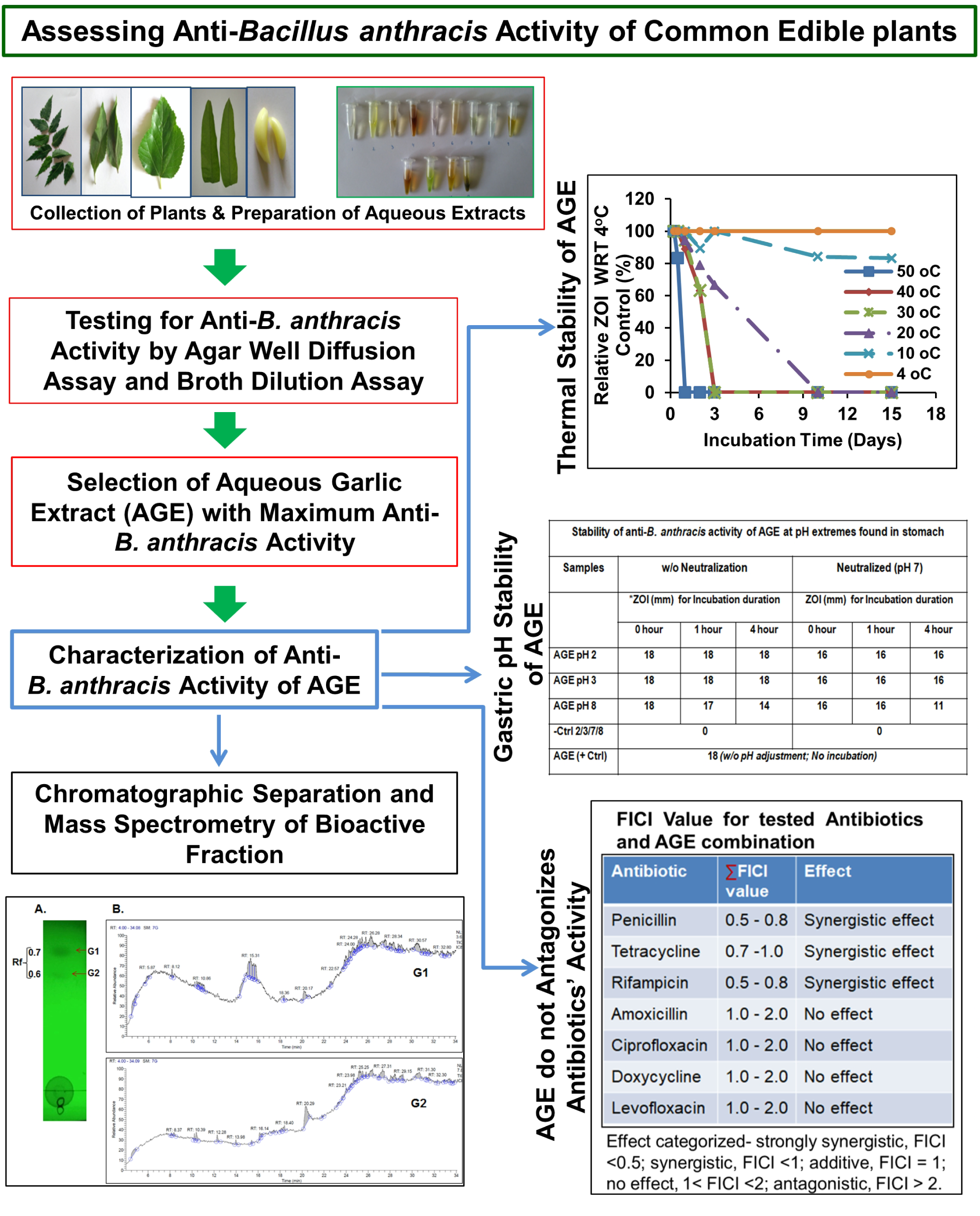

## 1. Introduction

Anthrax remains one of the major enzootic diseases in many poorer regions of sub-Saharan Africa, Asia, Central and South America (Shadomy *et al*., 2016; Turnbull, 2008). The causative agent *Bacillus anthracis* is endemic to these regions, primarily, due to favorable environmental conditions, availability of a wide range of animal hosts (*e.g.,* cattle, horses, sheep, *etc.*), inadequate surveillance and the poor healthcare facilities. The cutaneous anthrax is the most commonly observed form followed by the gastrointestinal anthrax under natural settings (Kaur *et al*., 2013; Turnbull, 2008). In case of humans, the activities like handling of the infected animals, their skins, carcasses and the consumption of the contaminated meat remain the primary means of acquiring the disease. The animals generally acquire anthrax while grazing on the contaminated pastures or by eating the infected carcasses. Inhalational or pulmonary anthrax is the rarest form that is mostly associated with the exposure to a large dosage of *B. anthracis* spores present in the air mainly due to bioterror activity (CDC, 2016).

The recent estimate of about 2000-20,000 human anthrax cases worldwide per year (Shadomy *et al*., 2016) could be a gross underestimate of the underlying problem due to the similarity of the disease symptoms with other common diseases. Initial symptoms of the gastrointestinal anthrax are very similar to diarrhea, while that of the inhalational anthrax are similar to flu. The lack of education, the absence of comprehensive surveillance, the unavailability of the diagnostic tools and the lack of medical infrastructure needed to treat anthrax in fulminant stage remain the major hindrances to proper anthrax control/management (Kaur *et al*., 2013; Shadomy *et al*., 2016; Turnbull, 2008). Prevention, diagnosis and treatment of anthrax had remained a priority (CDC, 2016; Kaur *et al*., 2013; Turnbull *et al*., 1999). Vaccines (Kaur *et al*., 2013; Shadomy *et al*., 2016; Turnbull, 2008), diagnostic kits and therapeutic antibiotics to manage anthrax infection (CDC, 2016; FDA, 2016) are available. ‘Centers for Disease Control’ and ‘Anthrax prevention expert panel’, recommends an extended combination of three or more antibiotics for the treatment of individuals exposed to anthrax spores (CDC, 2016; Hendricks *et al*., 2014). However, antibiotics are of no use if sufficient anthrax toxins (*e.g.*, Lethal toxin and Edema Toxin) have already accumulated in the body due to delay in the starting of antibiotics treatment. In the absence of sufficient medical support in anthrax endemic regions of Asia and Africa, people still use and rely on the traditional medicine that is based on empirical evidence and the knowledge of symptomatic treatment gathered over thousands of years to treat such diseases. The symptoms of anthrax are so similar to other less serious diseases that there is always a chance of wrong diagnosis or treatment.

The identification of edible plants or the plant parts with anti-*B. anthracis* activity could help fight endemic anthrax in poorer parts of the world. The conscious evidence-based inclusion of such plant(s) or the plant part(s) in the regular diet could effectively decrease the anthrax incidences. The current study had been undertaken to evaluate and characterize the anti-*B. anthracis* activity of fourteen commonly available edible plants which are purported to have antimicrobial or stomachic properties and also indicated in the traditional medicine for diarrhea, dysentery and stomachache (See Supplementary Table 1) (Brijesh *et al*., 2009; Chatterjee 1983; Chopra *et al*., 1956; Duke *et al*., 2002, 2008; Eigner *et al*., 1999; Ghosh *et al*., 1993; Hughes *et al*., 1991; Jaiarj *et al* 1999; Khare, 2007; Mazumder *et al*., 2006; Ndounga *et al*. 1997; Offiah *et al*., 1999; Parekh *et al*., 2007; Singh *et al*., 1983; U.S. Department of Agriculture, 1992-2016; Venkatesan *et al*., 2009). Although *Azadirachta indica* A. Juss. (Neem), *Allium cepa* L. (Onion), and *Curcuma longa* L. (Turmeric) have been indicated in traditional/folk medicine for treating anthrax-like symptoms their characterization and validation had been lacking (Duke *et al*., 2002, 2008; U.S. Department of Agriculture, 1992-2016). During our study, an evaluation of the acetone extracts of nine medicinal plants from South Africa against *B. anthracis* Sterne strain had been published (Elisha *et al*., 2016). The acetone extracts from *Maesa lanceolata, Bolusanthus speciosus, Hypericum roeperianum, Morus mesozygia and Pittosporum viridiflorum* displayed good anti-*B. anthracis* activity but variable cytotoxicity to vertebrate cells.

The identification of commonly used edible plant(s) with a sufficiently high concentration of water-extractable anti-*B. anthracis* constituents was performed using *B. anthracis* Sterne 34F2 (pXO1^+^, pXO2^-^) - an attenuated veterinary anthrax vaccine strain, as a surrogate for the wild type pathogenic *B. anthracis*. Several plants such as *Allium sativum* L. (Garlic), *Azadiracta indica* A. Juss. (Neem), *Mangifera indica* L. *(Mango), Berberis asiatica* Roxb. ex DC. (Daruharidra), *Psidium guajava* L. (Guava) and *Allium cepa* L. (Onion) displayed varying level of anti-*B. anthracis* activity. Interestingly*, Allium sativum* L. (Garlic), a common spice and herb used in food appeared to contain the highest concentration of water-extractable anti-*B. anthracis* activity among the tested plants, even more than *Azadirachta indica* A. Juss. (Neem) and *Allium cepa* L. (Onion) which are traditionally indicated for anthrax (Duke *et al*., 2002, 2008; U.S. Department of Agriculture, 1992-2016). The Aqueous Garlic Extract (AGE) exposure changed the morphology of *B. anthracis* cells within 3 hours (h) and decreased the number of viable cells in the liquid growth medium by 6 logs within 6-12 h. The preliminary characterization of the bioactive principles present in AGE that may be responsible for its anti-*B. anthracis* activity indicated it to be different from the ones already ascribed in the literature for the general antibacterial activity of Garlic (Duke *et al*., 2002; Goncagul and Ayaz, 2010; Sharifi-Rad *et al*., 2016; U.S. Department of Agriculture, 1992-2016). Furthermore, the AGE did not seem to antagonize the activity of antibiotics approved by Food and Drug Administration, USA (FDA) for the anthrax control, suggesting its safer interaction with the antibiotics.

## 2. Material and Methods

### 2.1 Bacterial strains and Chemicals

#### Escherichia coli DH5α and B. anthracis

Sterne 34F2 *-* an avirulent (pXO1^+^, pXO2^-^) vaccine strain were used in the current study. All bacterial growth media such as Muller-Hinton Broth (MHB), Muller-Hinton Agar (MHA), Luria Broth (LB), Luria Agar (LA), Nutrient Agar (NA), Nutrient Broth (NB) and 3-(4,5-dimethylthiazol-2-yl)-2,5-diphenyltetrazolium bromide (MTT) were from HiMedia Laboratories Ltd (India). The antibiotics and other routine chemicals were from Sigma Aldrich Inc (St. Louis, MO, USA). The organic solvents were purchased from Merck Millipore (Merck Life Science Private Limited, India).

#### Biosafety and bioethics statement

All experiments performed were approved by Institutional Biosafety committee, Panjab University, Chandigarh (IBSC approval number: IBSC-PANUNI- 042).

### 2.2 Collection of plant samples

The dried cloves/bulbs of Garlic (*Allium sativum* L.) and Onion (*Allium cepa* L.), and the dried rhizomes of turmeric (*Curcuma longa* L.) were purchased from the local market of Chandigarh, India. Fresh leaves of other identified plants from the premises of Panjab University, Chandigarh, India were collected in sterile polythene bags on the day of experimentation. The plants/ plant-parts used in the current study were identified by Professor Richa Puri, a noted Angiosperms Taxonomist at Department of Botany, Panjab University, Chandigarh. The voucher specimens of different plants were deposited in the herbarium of the Panjab University, Chandigarh (PAN) and the Acc. No. are given in Sup. Table 2. The Garlic (*Allium sativum* L.) batch with Acc. No. 21146 was used for all the experiments described in the current study including where it was compared (referred as sample No. 1) with other batches of Garlic (Acc. No. 21147 and 21148 referred to as sample No. 2 and 3, respectively) for assesing the batch to batch anti-*B. anthracis* activity variability.

### 2.3 Preparation of the aqueous plant extract

The freshly collected plant parts (see Sup. Table 2) were surface-sterilized by dipping in 70% ethanol (except turmeric, which was used as such due to the apparent absorption and retention of ethanol in the dried rhizome) followed by drying for 15 minutes at room temperature under aseptic conditions. The weighted plant samples (*e.g.,* 2g) were crushed in sterile ultrapure water (*e.g.,* 5 ml) using a sterile mortar pestle. The aqueous extract was obtained by separating the insoluble portion by decanting and passing it through a sterile Whatman No. 1 filter paper. The soluble filtered aqueous extracts of the plants were kept at 4° C until use (usually within 2-4 h) unless noted otherwise. The strength of aqueous extracts was calculated as initial plant material weight to volume of the extractant used, *i.e.,* the plant material (in mg) used to prepare per mL of aqueous extract (*e.g.,* 200 mg plant material used to prepare one mL of aqueous extract is reported as 200 mg/ml or 20% w/v; Note: evaporation yielded about 40-48 mg of residue per ml of 20% w/v AGE).

### 2.4 Anti-*B. anthracis* activity assessment of aqueous plant extracts

#### 2.4.1 Agar-well diffusion assay (AWDA)

The anti-*B. anthracis* activity of aqueous plant extracts was assessed by AWDA (Valgas *et al*. 2007; CLSI document M02-A11, 2012) using Sterne 34F2 strain as a surrogate for the wild type pathogenic *B. anthracis.* Briefly, the overnight grown *B. anthracis* Sterne strain culture was used to re-inoculate the fresh Muller-Hinton Broth (MHB) medium and the culture was allowed to reach the optical density at 600nm (OD_600_) of 0.3 -0.4. This culture was harvested, diluted in fresh medium to adjust its OD_600_ to 0.1 then further diluted to get evenly distributed lawn of the cells when 50-100 μl of diluted culture is spread on the Muller-Hinton Agar (MHA) plate or a mat when mixed with molten 1% MHA and incubated at 37^0^C for 12-16 h. The test wells in the agar plates were made using the wide mouth of a 200 μl pipette tips.

A fixed volume of the plant extracts (40% w/v) was loaded in the different wells of MHA plate for estimating their relative growth inhibitory potential. Antibiotic Rifampicin (2-8 μg) and the extractant ultrapure water were used as positive (+ Ctrl) and negative control (- Ctrl) for the growth inhibition, respectively. The plates were incubated for 12-72 h at 37°C. The zones of inhibition (ZOI) were measured at different time intervals by observing the plates against a black, non-reflective flat surface, under a reflected light source. The reported ZOI values are rounded off to the closest millimeter (mm).

The thermal stability of the anti*-B. anthracis* activity was evaluated by incubating the AGE samples (200μl aliquots of AGE (40% w/v)) at 4°C to 100°C for 0 h to 15 days, and determining the residual activity in AGE aliquots (50 µl) by AWDA as described above. The relative ZOI (%) was calculated as equal to ‘(Observed ZOI for sample incubated at test temperature (^0^C) / Observed ZOI for given sample incubated at 4°C temperature) *100.

#### 2.4.2 Growth assay

The overnight grown culture of *B. anthracis* Sterne strain was re-inoculated in MHB medium, incubated for 1.5 - 2 h at 37^0^C with agitation (150 rpm) to allow it to reach OD_600_ of about 0.4, then diluted with MHB medium to adjust its OD_600_ to 0.1. Different volumes (0 - 500 µl) of AGE (40% w/v) were added to the 5 ml of the diluted broth cultures (*∼* 0 - 36 mg/mL), followed by incubation at 37^0^ C with monitoring of the optical density, *i.e.,* OD_600_, at regular time intervals (0 - 24 h).

### 2.5 Verification/identification (DNA barcoding) of plant sample

DNA from the Garlic (*Allium sativum* L.) sample (PAN Acc. No. 21146) was extracted using Plant DNA isolation kit (Hi-Media). The conserved region of the *rbcL* gene was PCR amplified using the forward primer 5’ TGTAAAACGACGGCCAGTATGTCACCACAAACAGAGACTA AAGC 3’ and the reverse primer 5’ CAGGAAACAGCTATGACGTAAAATCAAGTCCACCACG 3’.

The program used was 1 min of initial denaturation at 94°C; 30 cycles of 45 seconds of denaturation at 94°C, 30 sec of annealing at 60°C, 1 min 30 sec of extension at 72°C; followed by a final extension of 10 min at 72°C. The amplified product was purified and sequenced on commercially available Sanger sequencing platform. The sequence of the *rbcL* gene fragment was aligned with reference sequences using nucleotide blast available at https://blast.ncbi.nlm.nih.gov/Blast.cgi?PROGRAM=blastn&PAGE_TYPE=BlastSearch&BLAST_SPEC=&LINK_LOC=blasttab&LAST_PAGE=blastn.

### 2.6 Morphological characterization of AGE exposed *B. anthracis* cells by Scanning Electron Microscopy

The exponentially growing *B. anthracis* cells were exposed to AGE for 0, 3 and 8 h followed by processing for observation by scanning electron microscope, essentially as described (Murtey, et al., 2016). Briefly, the *B. anthracis* cells were fixed with 2% glutaraldehyde for 1hr at room temperature, washed with phosphate buffer (100mM, pH 7.2), incubated with Osmium tetroxide for 1 h, dehydrated by serial passage in 10, 30, 50, 75, 90, and 100% ethanol and finally sputter coated with gold. The images were taken at different magnification using JSM 6100 (JEOL) Scanning Electron Microscope (SAIF, CIL, DST-supported facility at Panjab University, Chandigarh).

### 2.7 Evaluations of the ability of AGE to kill *B. anthracis* cells

#### 2.7.1 Time-Kill Assay

Time-kill assay was performed to assess the bactericidal activity of the extract (CLSI document M26-A, 1998). Briefly, the exponentially growing *B. anthracis* culture (OD_600_ ∼0.4) was centrifuged at 10,000 g for 5 minutes to pellet down the cells. The pelleted cells were washed with saline or MHB medium, followed by resuspension in the normal saline or MHB medium so as to have approximately 10^6^ colony forming units (CFU) /ml. It was supplemented with AGE at a final concentration of 0 to 3.6% w/v and incubated at 37°C for 0 - 24 h with shaking (150rpm), followed by plating of the AGE exposed *B. anthracis* culture on MHA (2% agar) medium plates and incubation at 37°C for 15-18 h. The colonies obtained were counted and the remaining CFU/ml in the AGE supplemented MHB medium or saline suspension was calculated for different AGE exposure durations (0-12 hrs).

#### 2.7.2. Effect of the exposure of AGE to pH extremes encountered in gastrointestinal tract, on its anti-B. anthracis activity

To assess possible utility of garlic as anti-*B. anthracis* agent for gastrointestinal infection the stability of AGE in conditions mimicking stomach and intestinal pH (*i.e.,* pH 2 - 8) was examined. The pH of AGE aliquots was adjusted in the range of 2 - 8 using 1N HCl or 1N NaOH and incubated at 37°C for different time durations (0-4 h). A 30µl aliquot of the incubated AGE samples (40% w/v) were assayed for residual anti-*B. anthracis* activity using AWDA as outlined above in section 2.4.1, after with or without pH neutralization to pH 7. The controls included AGE without any pH adjustment (+ Ctrl) and normal saline solutions of matching pH (2, 3 7, and 8) as pH adjusted solvent controls (- Ctrl). The ZOI produced by different samples were measured after overnight incubation of plates at 37°C.

#### 2.7.3 Relative efficacy of AGE in inhibiting growth of E. coli and B. anthracis

The relative potency of AGE in inhibiting the growth of laboratory strain *E.coli* DH5α and *B. anthracis* Sterne was assessed head to head by AWDA. The MHA plates (with and without (0.0025%) sodium deoxycholate supplementation) were divided into two halves/sectors - on one-half *E. coli* culture was spread, and on the other-half *B. anthracis* culture was spread. On this plate 50µl aliquots of AGE (40% w/v), with and without pH adjustment to pH 8, were placed in the wells already punched on the midline of two halves, followed by overnight incubation of plates at 37°C. The ZOI were measured for their relative antibacterial activity assessment as indicated in section 2.4.1. Rifampicin (8µg) was used as a positive control (+Ctrl) while autoclaved ultrapure water was used as a negative or solvent control (-Ctrl).

#### 2.7.4 Interaction of AGE with antibiotics used for Anthrax treatment

The relative potency and possible antagonistic interaction of AGE with the commonly employed antibiotics used to treat anthrax, *i.e.,* Amoxicillin (Am), Cefixime (Ce), Ciprofloxacin (C), Doxycycline (Dox), Levofloxacin (L), Penicillin (P), Rifampicin(R), Sulfamethoxazole (S), Tetracycline (T) was evaluated using AWDA as described above in section 2.4.1 (EUCAST 1998). The antibiotics Ciprofloxacin, Doxycycline, Levofloxacin and Penicillin are approved by FDA for anthrax treatment (FDA, 2016) while Sulfamethoxazole, a Folic acid biosynthesis inhibitor that is known to be ineffective against *B. anthracis* (Weiss *et al*., 2015), was used as a negative control. The ZOI remaining after different durations of incubation at 37°C were noted.

#### 2.7.5 Minimum Inhibitory concentration (MIC) determination & fractional inhibitory concentration index (FICI) assessment

The checkerboard broth dilution method (White *et al*., 1996) was used to determine the Minimum Inhibitory Concentration (MIC) of AGE alone and in combination with different antibiotics (*i.e.,* synergistic, no effect, antagonistic as *fractional inhibitory concentration index*). Briefly, the *B. anthracis* Sterne strain cells grown overnight in Mueller Hinton Broth (MHB) were inoculated in fresh MHB (100µl of culture in 5ml MHB) followed by incubation at 37°C with agitation (150rpm) until the OD_600_ reached 0.4 (1.5 - 2 h). It was diluted in fresh MHB to adjust OD_600_ to 0.2 or 0.1. A 100µl aliquot was inoculated in wells of 96-well plate already supplemented with different concentrations of antibiotic and AGE in 100µl MHB followed by overnight incubation at 37°C with agitation at 150rpm. Different controls such as media control, *i.e.,* only MHB; no treatment or bacterial cell control, *i.e.,* cells in media; AGE control, *i.e.,* AGE alone in media; antibiotic control, *i.e.,* antibiotic in media were also included/set for experiments as applicable. The bacterial growth in different wells were assessed by directly measuring OD_620_ using ELISA plate reader. The cell viability was assesed by MTT assay (Singh *et al*., 2002). The culturability of surviving cells in the wells with no observable increase in OD were assessed by platting them on MHA plate. The synergistic, antagonist or indifferent interaction of AGE with antibiotics approved by FDA for anthrax treatment, was determined using the checkerboard method essentially as described previously (Mor *et al*., 2015). Briefly, different concentrations of antibiotics and AGE both alone and in combinations were supplemented to test-organism containing broth (1-2×10^6^ CFU/ml) in 96 well plates. The fractional inhibitory concentration index (FICI) was calculated using the formula: ∑FICI = FIC(A) + FIC(B), where, FIC(A) = MIC(A) in combination / MIC(A) alone; and FIC(B) = MIC(B) in combination/ MIC(B) alone. The interaction was categorized as follows: strongly synergistic effect, FIC < 0.5; synergistic effect, FIC < 1; additive effect, FIC = 1; no effect, 1 < FIC < 2; antagonistic effect, FIC > 2 (Mor *et al*., 2015).

### 2.8 Plasmid loss assay

To assess the AGE exposure induced curing or loss of pXO1 plasmid, the *B. anthracis* cells were grown in the presence AGE concentrations (0-3.6% w/v) that retarded/delayed or inhibited its growth in the broth culture. The loss of virulence plasmid pXO1 from cells was assessed at different time intervals (0 - 24 h). The AGE exposed cultures were plated on MHA plates and the colonies that developed on incubation at 37°C for 12-16 h were examined for the loss of pXO1 by assessing the presence of pXO1 encoded toxin gene *pagA* (encodes for Protective Antigen or PA) by colony PCR. Primers, PA-1588F (5’GCA TTT GGA TTT AAC GAA CCG A 3’) and PA-2016R (5’TCC ATC TTG CCG TAA ACT AGA A 3’) were used for *pagA* amplification. The presence of chromosomal gene *phoP,* a component of PhoPR two-component system (Aggarwal *et al*., 2017), was assessed using primers PhoP-F (5’GCG CCC ATG GGC ATG AAC AAT CG 3’) and PhoP-R (5’GCG CCT CGA GTT CAT CCC CTT TTG GC 3’). The condition used for the colony PCR was 30 cycles of ‘45 sec of denaturation at 94°C, 30 sec of annealing at 58°C for *pagA/* 60°C for *phoP*, 1 min 30 sec of extension at 72°C’, followed by 10 min of final extension at 72°C.

### 2.9. Thin Layer Chromatography (TLC), TLC - Bioautography and Mass Spectrometry

A 5-10 µl aliquot of the filtered Aqueous Garlic Extract (AGE) was applied 1 cm away from the base of silica gel 60 TLC plate (Merck Millipore). The applied sample was air dried for 10 min at 37°C, followed by developing the chromatogram with Toluene-Acetone (7:3) solvent system. The TLC plates were run in duplicate. The developed chromatograms/ plates were air-dried by keeping them at 37°C for 20 min. The chromatograms were observed under short- and long- UV radiation. One plate was used for the bioautography (Dewanjee *et al*., 2015) to test the ability of fractionated components to inhibit the growth of *B. anthracis* while the other one was used as a reference plate - that was later used for mass spectrometry to identify the anti-*B. anthracis* activity displaying bioactive principles. The positions of UV-fluorescent bands on the plate were marked as G1 and G2. The retardation factor (Rf) of G1 and G2 bands was also determined as described (Fried *et al*.1996). For bioautography, the agar overlay bioassay was performed. The developed plates were covered with molten 1% MHA medium mixed with exponentially growing *B. anthracis* cells, followed by incubation at 37°C for 15-18 h. The presence/location of the zone of growth inhibition on the plates was noted. As negative controls, TLC plates developed with Toluene-Acetone (7:3) mixture without the loading of AGE and the chromatography media (*i.e.,* silica) alone were also assessed by bioautography. For mass spectrometry the areas corresponding to the zone of inhibition and encompassing the UV-fluorescent bands were scratched from the duplicate TLC plate with the help of sterile blades, placed in separate 1.5 ml tubes containing 100μl of absolute ethanol and then processed for Gas chromatography – Mass spectrometry (GC-MS) analysis. The scraped amount from a single plate was sufficient for the GC-MS analysis. As a control for the process, the butanolic Garlic extract was also prepared and processed for GC-MS analysis. The samples were analyzed using Thermo Scientific Gas Chromatograph - Mass Spectrometer or GC-MS (Thermo Trace 1300 GC and MS Thermo TSQ 8000) fitted with column TG 5MS (30m X 0.25mm, 0.25µm) using He as a carrier gas. The NIST 2.0 Library from National Institute of Standards and Technology was employed to identify potential compound hits based on the similarity of their spectrum with the already stored spectrum of the known components.

## 3. Results

### 3.1 Several common plants have anti-*Bacillus anthracis* activity

The aqueous extracts from different plants prevented the growth of *B. anthracis* cells in AWDA (Figure 1A-B). The growth inhibitory potential of the aqueous extracts of tested edible plants from one representative evaluation experiment is summarized in Sup. Table 2. Among the tested plants Garlic (*Allium sativum* L.) displayed the highest anti-*B. anthracis* content on wet weight basis comparison. (Figure 1A-B, Sup. Table 2). The aqueous extract of Garlic (AGE) produced a zone of inhibition (ZOI) of 18 mm diameter which was sustained up to 72 h of incubation similar to that produced by the +Ctrl Rifampicin (8µg). The ZOI produced by others such as Neem (*Azadirachta indica* A. Juss.) and Mango (*Mangifera indica* L.) were comparatively smaller measuring just about 10-11 mm by 12-24 h of incubation that usually disappeared by 72 h of incubation. The extracts from Daruharidra (*Berberis asiatica* Roxb. ex DC), Onion (*Allium cepa* L.) and Guava (*Psidium guajava* L.) did not produce a ZOI in the shown AWDA when 50 μl of the 40 % w/v aqueous extract was evaluated (Figure 1A-B, Sup. Table 2). Based upon multiple activity evaluation experiments (data not shown), it may be deduced that the activity in the extracts of Neem, Mango, Daruharidra, Onion, and Guava are variable (batch to batch, plant to plant, and time to time). However, on an average the anti-*B. anthracis* activity content in the aqueous extracts made from different plants were in the order: Garlic (*Allium sativum* L.)>> Neem (*Azadirachta indica* A. Juss.), Mango (*Mangifera indica* L.)> Daruharidra (*Berberis asiatica* Roxb. ex DC), Onion (*Allium cepa* L.), Guava (*Psidium guajava* L.). As the Garlic extract showed maximum potency on weight to weight basis and the minimum experiment to experiment and batch to batch variability, it was selected for further characterization.

**Figure 1.**
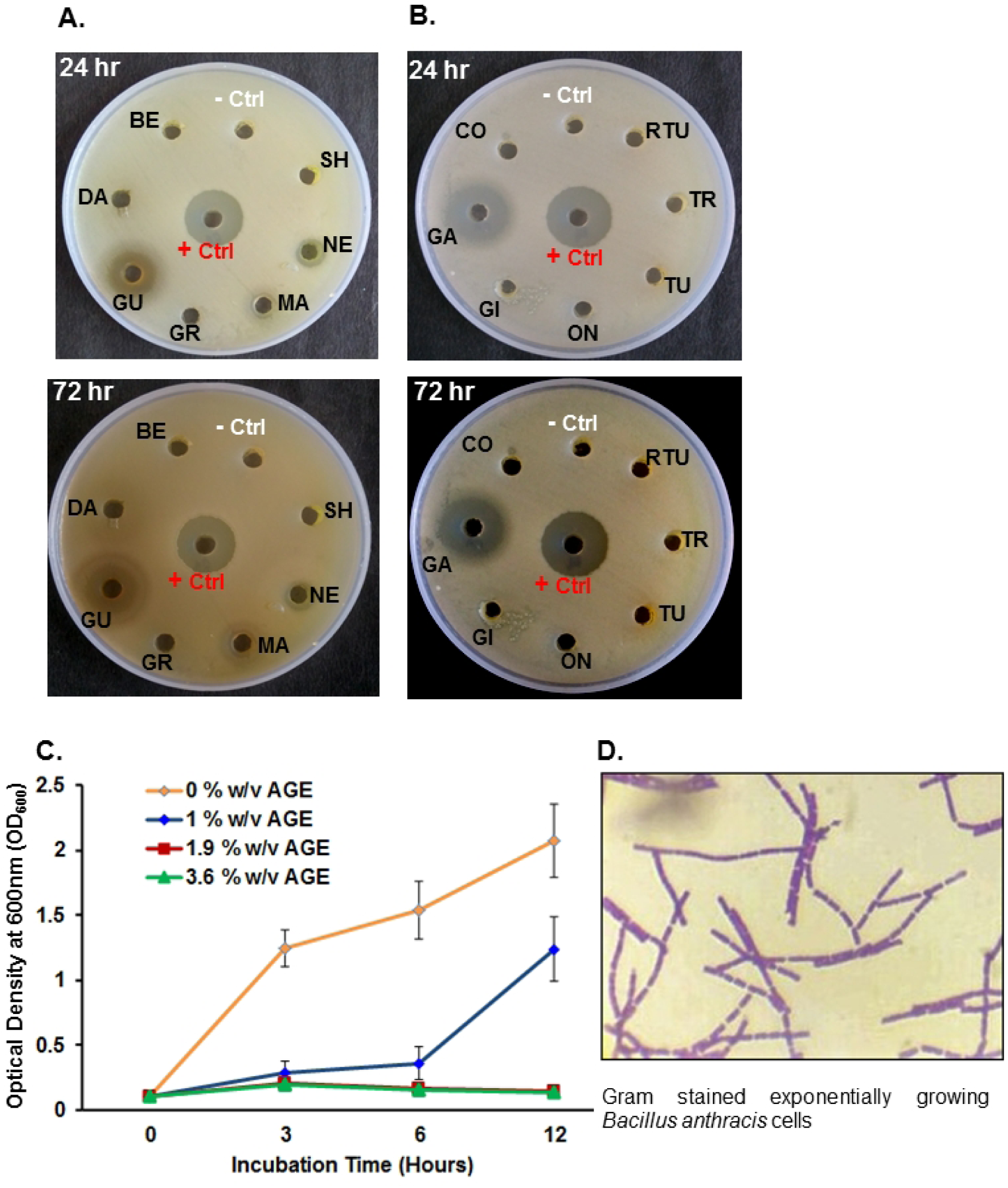
Common edible plants have anti-*Bacillus anthracis* activity. The MHA plates showing ZOI produced by the aqueous plant extracts in agar well diffusion assay (AWDA) - **(A)** Bael (*Aegle marmelos* (L.) Correa) leaf: BE; Daruharidra (*Berberis asiatica* Roxb. ex DC) leaf: DA.; Guava (*Psidium guajava* L.) leaf: GU; common grass (*Cynodon dactylon* (L.) Pers.): GR; Mango (*Mangifera indica* L.) leaf: MA; Neem (*Azadirachta indica* A. Juss.) leaf: NE; Shetuta (*Morus indica* L.) leaf: SH; **(B)** Coriander (*Coriandrum sativum* L.) leaf: CO; Garlic (*Allium sativum* L.) bulb: GA; Ginger (*Zingiber officinale* Roscoe) rhizome: GI; Onion (*Allium cepa* L.) bulb: ON; Tulsi (*Ocimum sanctum* L.) leaf: TU; Turmeric (*Curcuma longa* L.) rhizome: TR; Ram Tulsi (*Ocimum gratissimum* L.) leaf: RTU. Antibiotic Rifampicin and ultrapure water were used as positive control (+ Ctrl) and solvent/diluent control (- Ctrl), respectively. The Aqueous Garlic (*Allium sativum L.*) Extract (AGE) appeared to be the most potent based upon the ZOI produced (see Sup. Table 2). **(C)** The supplementation of exponentially growing culture with 0-3.6% w/v of AGE inhibited its growth (OD_600_) in a dose-dependent manner. The values shown are the average of three independent experiments (n=3) and the error bars represent standard error of the mean. **(D)** Gram-stained exponentially growing *B. anthracis* cells (0% w/v AGE).

### 3.2 Aqueous Garlic (*Allium sativum* L.) Extract (AGE) inhibits the growth of *Bacillus anthracis* in a dose-dependent manner

The supplementation of MHB medium with different concentrations of AGE (0-3.6% w/v) inhibited the growth of exponentially growing *B. anthracis* culture in a dose-dependent manner (Figure 1C). The supplementation of MHB at concentrations ≥1.9% w/v of AGE completely inhibited growth of *B. anthracis* while the lower concentrations retarded/delayed the growth in a dose-dependent manner. The corresponding MIC for *B. anthracis* ≤20mg/ml, was lower than that reported previously for Gram-positive bacterial strains, *i.e.,* range 142.7–35.7 mg/ml, but fell in the MIC range of 35.7–1.1mg/ml reported for the Gram-negative strains (Duke *et al*., 2008 and references therein). It could be because of the general more sensitive nature of *B. anthracis* as also observed with its sensitivity towards various antimicrobials.

### 3.3 AGE induces morphological changes in *B. anthracis*

To assess the possible mechanism of anti-*B. anthracis* activity the morphology of *B. anthracis* cells growing in cultures supplemented with 0% w/v AGE (control) and 1.9% w/v AGE (treated) for different durations were examined using light microscopy and scanning electron microscopy. The control *B. anthracis* culture which was not exposed to AGE had unchanged morphology as expected (Figure 2 A top part, control cells; see also Figure 1D). The culture exposed to 1.9% w/v AGE, a growth inhibitory concentration, had altered morphology (shorter chains) or as single cells (Figure 2A bottom part, treated cells) within 3 h of AGE exposure. This observation indicates potential disruption/dissolution or weakening of the cell walls by AGE exposure. Incubation for 8 h resulted in the appearance of debris (data not shown).

**Figure 2.**
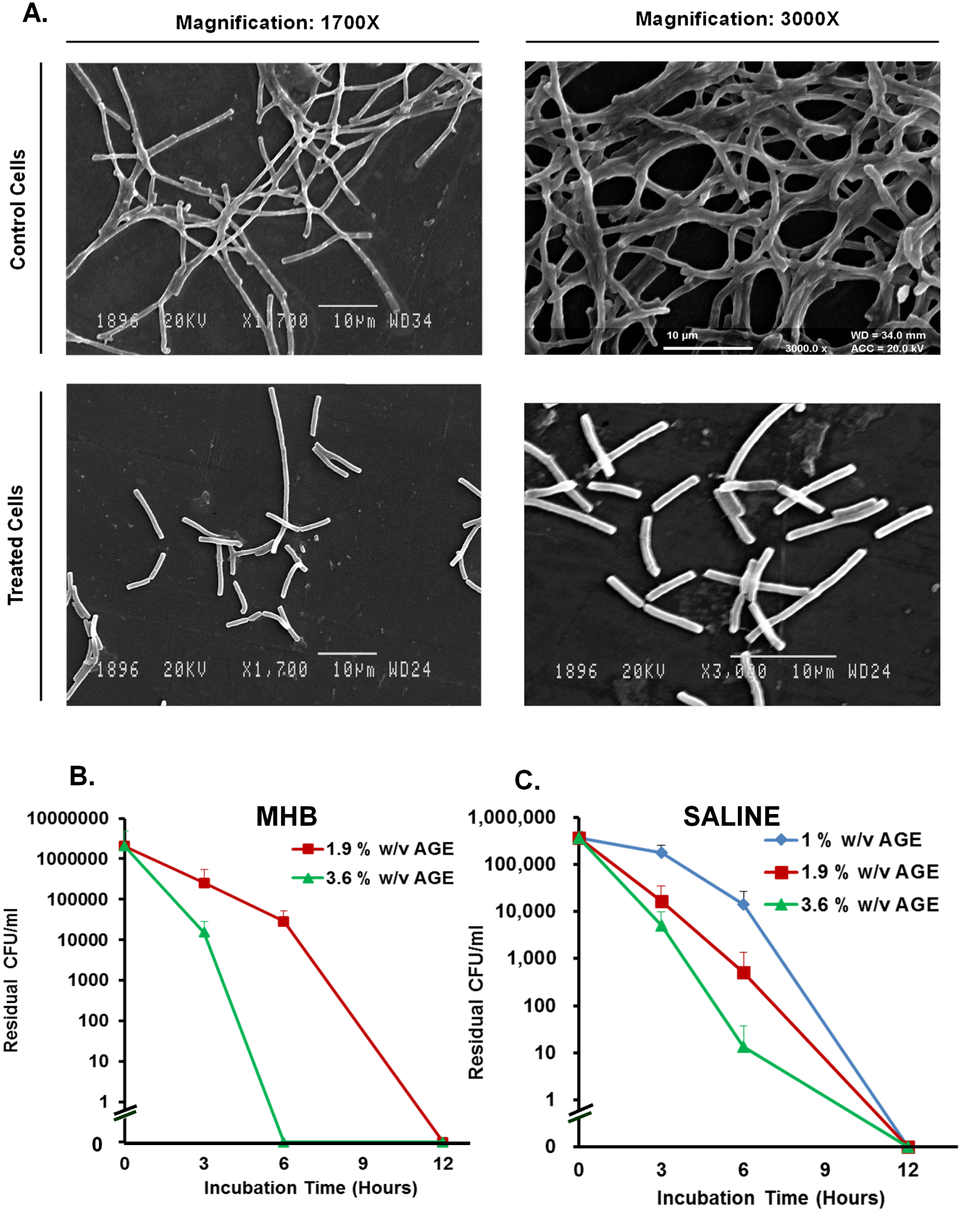
Aqueous Garlic (*Allium sativum* L.) Extract (AGE) kills *Bacillus anthracis* cells. **(A)** Scanning Electron Microscopy (SEM) image of the exponentially growing *B. anthracis* cells (top panel; control cells) and those exposed to 1.9% w/v AGE for 3 h (bottom panel; treated cells). Disintegration of longer chains and the appearance of smaller chains and single cells in the AGE exposed *B. anthracis* culture was observed (compare control cells in the top panel with AGE treated cells in the bottom panel). The exposure of exponentially growing *B. anthracis* cells to AGE (0 - 3.6% w/v at 37°C) in both MHB medium **(B)**, and normal saline **(C)**, decreased their viability (CFU/mL). The data shown is average of three experiments (n=3) and the error bars represent standard deviation.

### 3.4 AGE is bactericidal to vegetative *B. anthracis* cells

The bactericidal activity of AGE was assessed by exposing *B. anthracis* cells in the log phase of growth with different concentration of AGE in the presence of growth medium or saline followed by estimating the number of surviving colony forming units /ml (*i.e.,* CFU/ml) after different time intervals (0-12 h). The AGE at concentration 1.9% w/v and above was found to cause a decrease in the viability (CFU) of vegetative *B. anthracis* cells by as much as 6 logs (Figure 2B and C) within 6-12 h. However, 1 % w/v AGE which could only retard the growth in the presence of MHB medium (See Figure 1C; data not shown in Figure 2B) was found to kill *B. anthracis* cells in the absence of MHB (Figure 2C) indicating increased efficacy of AGE in killing *B. anthracis* cells in the absence of growth medium components.

### 3.5 AGE does not induce virulence plasmid pXO1 loss (plasmid curing) from *B. anthracis* Sterne strain

The colonies that developed on MHA plate on inoculation of *B. anthracis* culture exposed to sub-inhibitory concentration of AGE (1% w/v) for different durations (0 - 24 h), were examined for the presence of *pagA* gene - a virulence plasmid pXO1-borne gene, and *phoP* gene – a chromosome-borne gene using polymerase chain reaction (Sup. Figure 1). The exposure to a sub-inhibitory concentration of AGE for 24 h did not cause the loss of pXO1 plasmid as apparent from the specific amplification of the plasmid-borne *pagA* gene in 100% of the random colonies (8 out of 8) screened (Sup. Figure 1 A). The sampled colonies formed by the culture of *B. anthracis* cells exposed to 1% w/v AGE for different durations, *i.e.,* 0 - 24 h also consistently tested positive for the presence of *pagA* gene indicating retention of pXO1 plasmid and the inability of AGE to induce virulence plasmid pXO1 loss or curing (Sup. Figure 1 B). Similar results were obtained with colonies formed from residual surviving cells present in the *B. anthracis* culture exposed to growth inhibitory concentration of AGE, *i.e.,* >1.9% w/v for shorter durations (1 – 3 h; data not shown).

### 3.6 Anti-B*. anthracis* activity of AGE is stable near room temperature

The ability of the anti-*B. anthracis* activity of AGE to tolerate different temperatures was evaluated to assess its potential field use. Based upon the ZOI produced by AGE samples incubated at different temperatures (0-100°C) for different durations (0-15 days), it can be inferred that the anti-*B. anthracis* activity of AGE is quite stable around room temperature (20- 30°C), and also moderately stable at temperatures encountered near tropics during the summer season (see Figure 3A and 3B). It retained more than 80% of anti-*B. anthracis* activity on incubation at 40°C and 50°C for >1 day and 12 h, respectively. At 4°C incubation temperature, AGE did not show any loss in activity up to 15 days. However, on incubation at 100°C temperature, the anti-*B. anthracis* activity of AGE was lost within about 10 minutes (data not shown); suggesting at least S-allylcysteine (Kodera *et al*., 2002) which is reportedly quite stable at 100°C, would not be the bioactive constituent responsible for the anti-*B. anthracis* activity of AGE.

**Figure 3.**
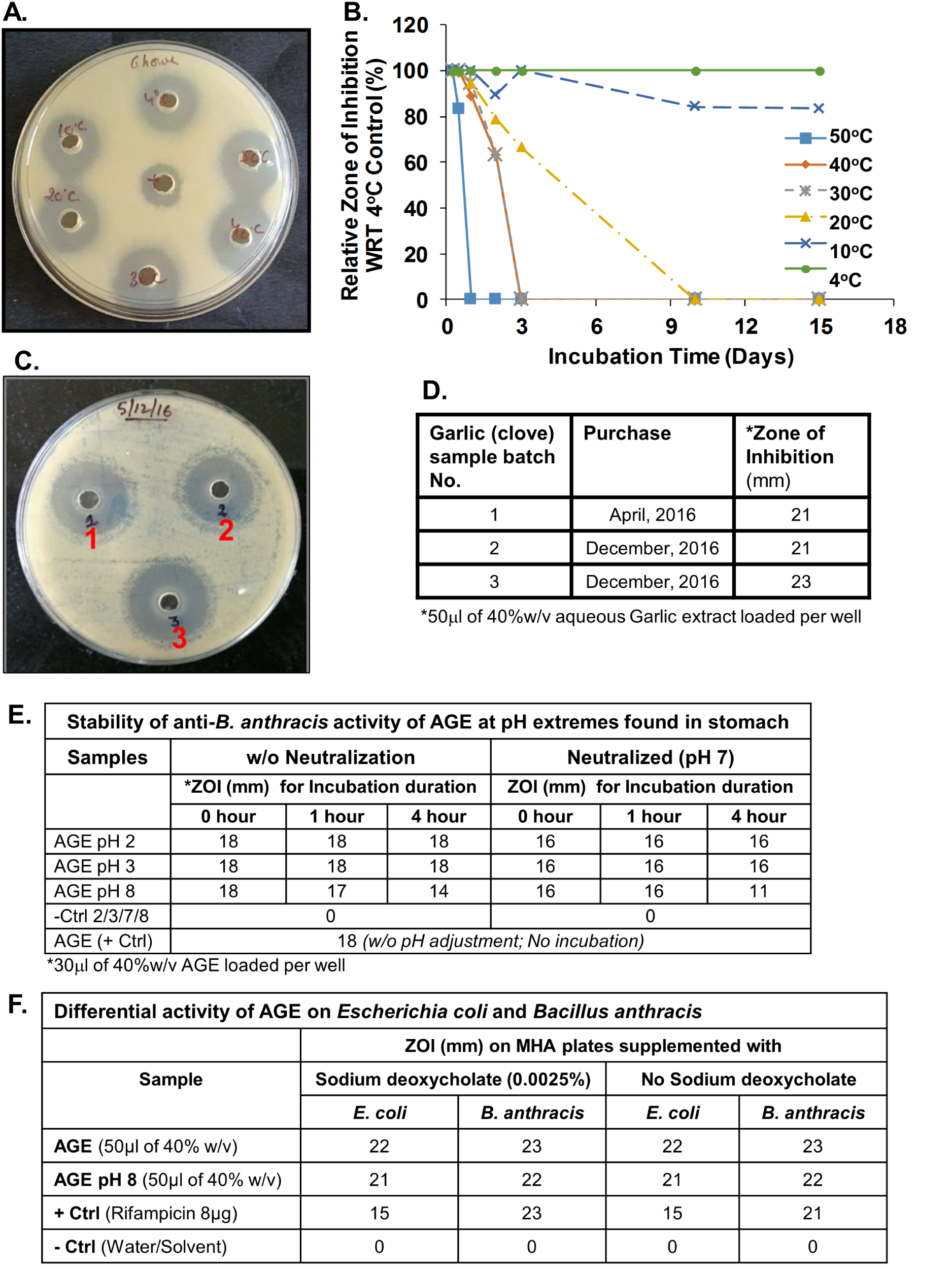
Characterization of Aqueous Garlic (*Allium sativum*) Extract (AGE). **(A-B)** Exposure of AGE (40% w/v) to higher temperature decreases its anti- *B. anthracis* activity. The result of an AWDA performed to assess the anti*-B. anthracis* activity remaining in AGE (50µl) after 6 h of incubation at 4 - 50°C temperature is shown in **(A)**. Antibiotic Rifampicin was used as positive control (+). **(B)** The relative ZOI produced by AGE samples incubated up to 15 days at temperatures 4-50°C. The data is presented as ZOI with respect to (WRT) that produced by AGE incubated at 4°C. The freshly prepared AGE appeared to be quite stable near room temperature. The AGE incubated at 40°C retained >80% activity even after incubation for 1 day. **(C)** Comparison of the anti-*B. anthracis* activity of AGE made from three different batches purchased from the local market of Chandigarh, India by AWDA. **(D)** Summary of the assay shown in (C). **(E)** Assessment of the effect of different pH conditions (*i.e.,* 2, 3 and 8) on AGE’s anti-*B. anthracis* activity. Aliquots of pH adjusted AGE (*i.e.,* pH 2, 3 and 8) were incubated for 0, 1 and 4 h at 37°C. The 30µl aliquots of the incubated samples with or without pH neutralization were assessed for the remaining/residual anti-*B. anthracis* activity by AWDA. (-Ctrl 2/3/7/8: Normal saline pH 2/3/7/8; +Ctrl: AGE; AGE pH 2/3/8: AGE samples with their pH adjusted to 2/3/8). **(F)** The relative sensitivity of *Escherichia coli* DH5α strain and *Bacillus anthracis* Sterne strain to AGE and the effect of the presence of bile salt sodium deoxycholate (0.0025%) on the AGE’s antimicrobial activity were assessed head to head on MHA plate by AWDA as indicated in section 2.7.3. The ZOI values reported are the average of three experiments (n=3).

### 3.7 Different batches of Garlic show similar anti-*B. anthracis* activity

The bioactive content of plant products are known to vary with origin, age and time, so for using any plant product for any particular purpose the plant or plant products from different origin or of age must consistently display desired characteristics within acceptable limits. To this end, we evaluated garlic samples collected/bought from the local market over an extended period (April 2016 to December 2016) for anti- *B. anthracis* activity using AWDA. All three samples evaluated displayed robust and similar anti-*B. anthracis* activity as apparent from the production of 21-23 mm size ZOIs (Figure 3C and 3D).

### 3.8 Anti-*B. anthracis* activity in AGE relatively stable in the physiologically relevant conditions

The ability of AGE to tolerate different physiologically relevant pH conditions was evaluated to assess its potential application in anthrax control (Figure 3E). Based upon the ZOI produced by pH adjusted AGE samples that were incubated for 1h and 4-h at 37°C, the anti-*B. anthracis* activity of AGE appeared to be quite stable at pH 2, 3 and 8 (AGE pH 2/3/8). However, the activity remaining in the samples tested without (w/o) any pH neutralization seemed to be more than those neutralized, i.e., adjusted to 7, before activity testing. We do not know the reason for this observation but speculate that the observed variation in AGE activity pre- and post-neutralization may be due to change in the solubility of antimicrobial compounds. The presence of bile salts, such as sodium deoxycholate, also did not seem to antagonize the anti-*B. anthracis* activity of AGE when tested up to the final concentration of 0.0025% w/w (Figure 3F). Still higher concentrations were not tolerated by *B. anthracis* culture (data not shown). Additionally, head-to-head comparison of the antimicrobial activity of AGE towards *B. anthracis* and the laboratory strain of *E. coli* DH5α, as a surrogate for the commensal resident *E. coli*, indicated *B. anthracis* to be slightly more sensitive to AGE than the *E. coli*.

### 3.9 AGE does not antagonize the activity of antibiotics recommended for the treatment of possible *Bacillus anthracis* exposure

Antibiotics prescribed for anthrax treatment, Amoxicillin (Am), Cefixime (Ce), Ciprofloxacin (C), Doxycycline (Dox), Levofloxacin (L), Penicillin (P), Rifampicin (R) and Tetracycline (T) (CDC, 2016; FDA, 2016) were assayed for possible interaction with AGE using AWDA. All the FDA approved antibiotics for anthrax inhibited the growth of *B. anthracis* as expected (Figure 4A and 4B). AGE did not appear to antagonize their activity. Further experimentation to estimate the MIC value along with fractional inhibitory concentration index (FICI) indicated a synergistic interaction of AGE with the antibiotics Penicillin, Rifampicin, Tetracycline while an indifferent interaction with the antibiotics Amoxicillin, Cefixime, Ciprofloxacin, Doxycycline, Levofloxacin (Figure 4C and Figure 4D). The MIC of AGE for *B. anthracis* was estimated to be 12.5 mg/mL, a value significantly lower than that reported for other Gram-positive bacterial strains (142.7–35.7 mg/ml) (Duke *et al*., 2008 and references therein).

**Figure 4.**
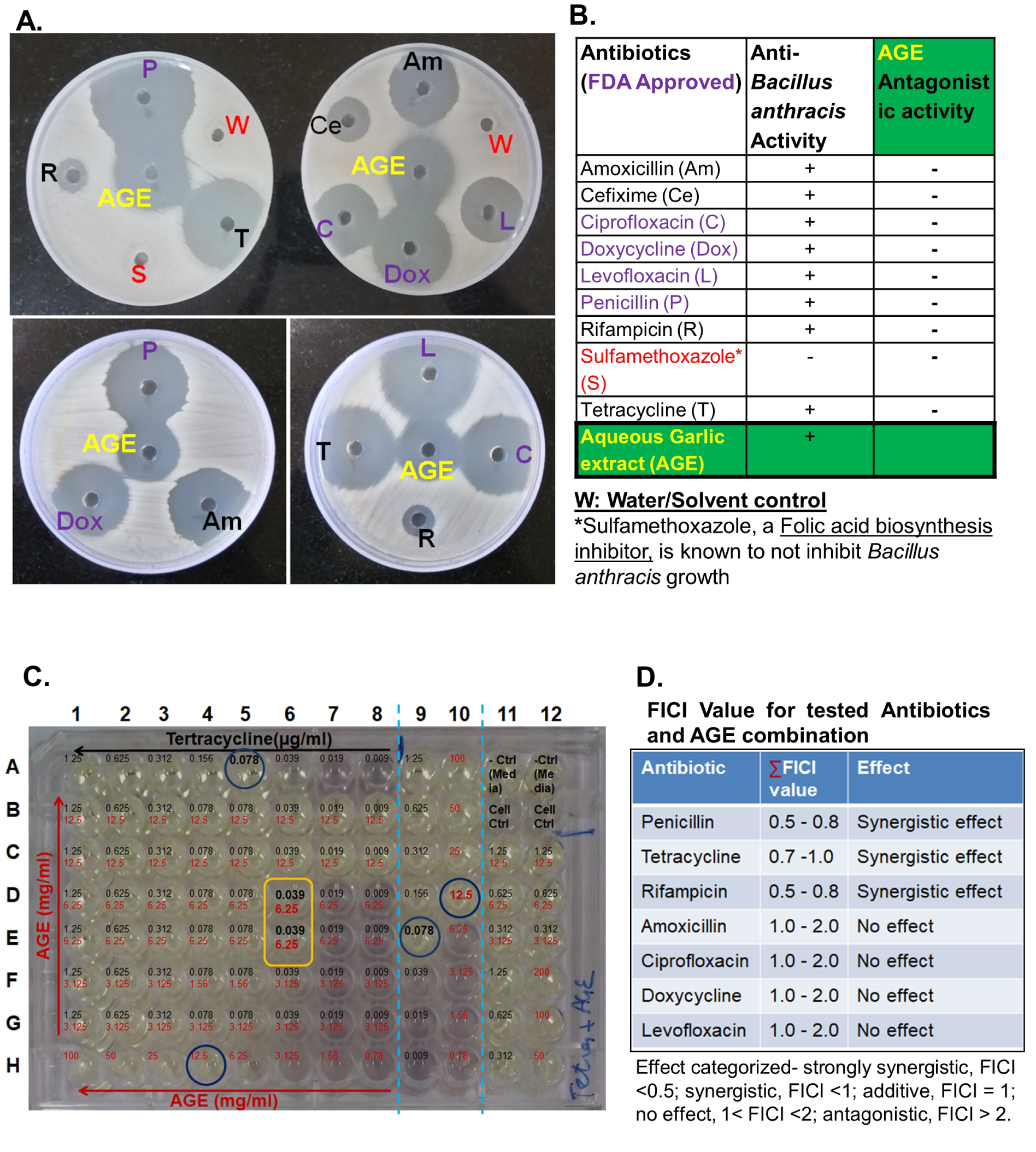
Aqueous Garlic (*Allium sativum*) Extract (AGE) does not antagonize the anti- *Bacillus anthracis* activity of the commonly employed antibiotics for anthrax control. (A) The antibiotics Amoxicillin (Am), Cefixime (Ce), Ciprofloxacin (C), Doxycycline (Dox), Levofloxacin (L), Penicillin (P), Rifampicin (R), Tetracycline (T) along with Sulfamethoxazole (S) - an antibiotic known to be ineffective against *B. anthracis,* were assayed for possible interaction with AGE using AWDA. The top and bottom panels are from an experiment performed with different concentration of the indicated test substances. Except Sulfamethoxazole, all antibiotics and AGE inhibited the growth of *B. anthracis* as expected. **(B)** Summary of the interaction of AGE with antibiotics in inhibiting *B. anthracis* growth as apparent from (A). Summary of the relative ability of different tested substances in inhibiting *B. anthracis* growth is provided in Sup. Figure 7. **(C)** A typical microdilution experiment for MIC and FICI value calculation is shown. The interaction between Tetracycline (*i.e.,* 1.25 - 0.009 µg/mL; black) and AGE (*i.e.,* 100 - 0.78 mg/mL; red) was assessed in the representative plate displayed. The concentration of tetracycline and AGE alone decreased from left to right (row A and H), while the concentration of AGE in combination with the indicated fixed concentration of antibiotic increased from bottom to top (row G to B) as indicated by the arrows. Column 9 and 10 are the duplicates of row A and H, respectively. The wells A11-12 are media control (-Ctrl) while B11-12 are cell control, *i.e., B. anthracis* cells without any antibiotics or AGE. Values indicated in the blue circle are MIC value of tetracycline (*i.e.,* 0.078 µg/mL) and AGE (*i.e.,* 12.5 mg/mL) in the rows A and H, respectively. Values in the yellow circle indicate the concentration of tetracycline and AGE in combination (*i.e.,* 0.039 µg/ml and 6.25 mg/ml respectively). (D) Fractional inhibitory concentration index (FICI) calculation for antbiotics and AGE combination. The values represent the range of three experiments performed independently.

### 3.10 Autobiography of AGE and Mass spectrometry of bioactive components

The separation of AGE constituents into bioactive entities was performed by coupling the analytical TLC separation (Supplementary Figure 2A) with bioautography (Sup. Figure 2B).The eluted potential bioactive components from the place corresponding to the bioactivity on the duplicate TLC chromatogram were subjeced to GC-MS for the detection of the ions belonging to potential bioactive compounds (Sup. Figure 3). The Toluene-Acetone (7:3) solvent system was found to be the best in separating two UV–fluorescent spots/components in AGE on silica TLC (Sup. Figure 2A, compare chromatogram No. 3 of bottom panel with others). The bioautography indicated UV-fluorescent spots to be inhibitory to *B. anthracis* growth (Sup. Figure 2B; encircled in white in *right panel: AGE TLC plate*). As expected, the negative controls used in the study did not inhibit the growth of *B. anthracis*. The UV-fluorescent spots observed in the AGE fractions that corresponded to anti-*B. anthracis* activity (growth inhibition) on bioautography were labeled as G1 and G2 (Sup. Figure 3A), isolated and processed for GC-MS analysis. The ZOI observed on bioautography covered both G1 and G2 spots, so one cannot be concluded as better than the other. The calculated retardation factor (Rf) for G1 and G2 was 0.7 and 0.6 respectively. The GC-MS profile of G1 and G2 are shown in Sup. Figure 3B. The list of potential bioactive compound hits as ascertained using the NIST 2.0 library is provided in Table 1 and the complete report can be found in the supplementary information (Sup. Figure 4 & 5). The ions generated from bioactive fraction of AGE did not seem to indicate the presence of the well-known bioactive organosulfur compounds reported in Garlic, *e.g.,* alliin, S-allyl-L-cysteine, γ- glutamyl-S-allyl-L-cysteine, allicin *etc.* (Duke *et al*., 2002; Goncagul and Ayaz, 2010; Rana *et al*., 2011; Sharifi-Rad *et al*., 2016; U.S. Department of Agriculture, 1992-2016). However, other class of compounds with potential antimicrobial activity were identified, *e.g.*, phthalic acid derivatives, acid esters, phenyl group-containing compounds, steroids etc. (Table 1 & 2; Sup. Figure 4 & 5; Perfluorotributylamine peaks coming from column were not included in Table 1). The absence of the well-known antibacterial compounds in our bioactive fraction could be the result of our use of water as extractant and the methodology adopted that would have led to the reduced extraction, decomposition and/or loss of the previously reported bioactive compounds (Kim, *et al*., 2005). The GC-MS analysis of the butanolic extract of Garlic was performed to assess the validity of this speculation (Sup. Figure 3C). The butanolic extract indeed contained well-known allyl derivatives as well as multiple S-containing compounds (Table 1, Sup. Figure 6) supporting the speculation.

**Table 1.**
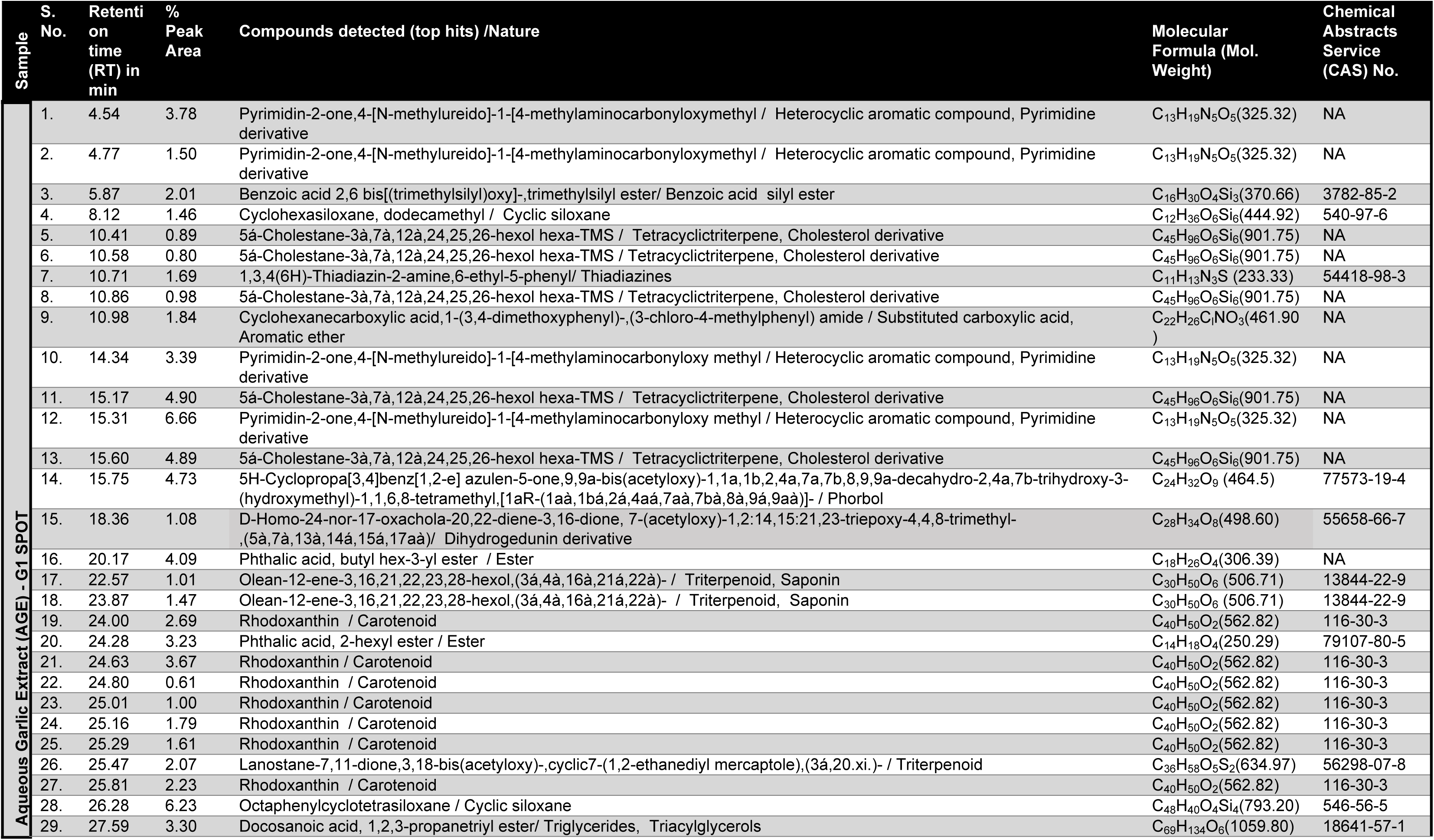
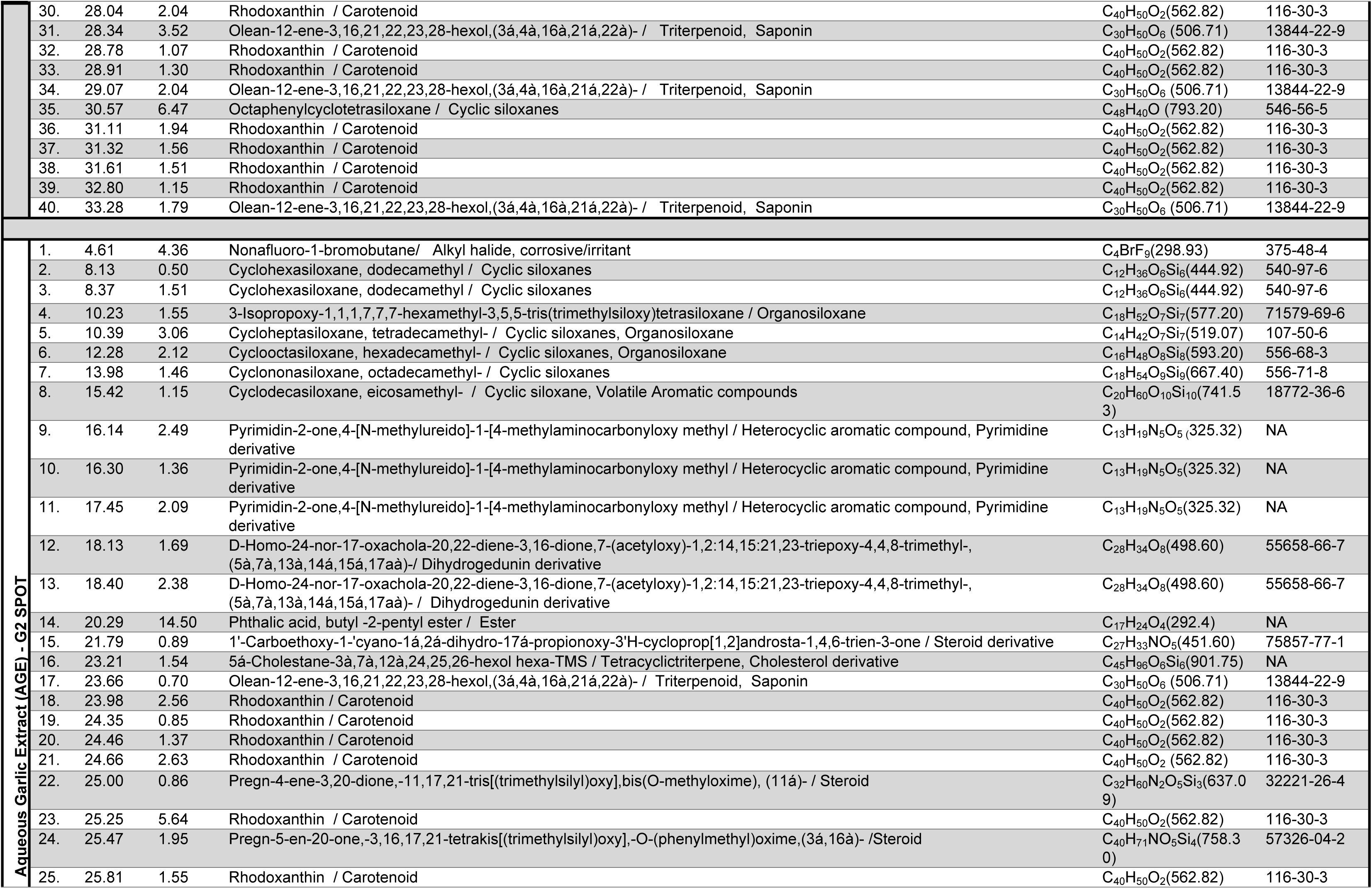
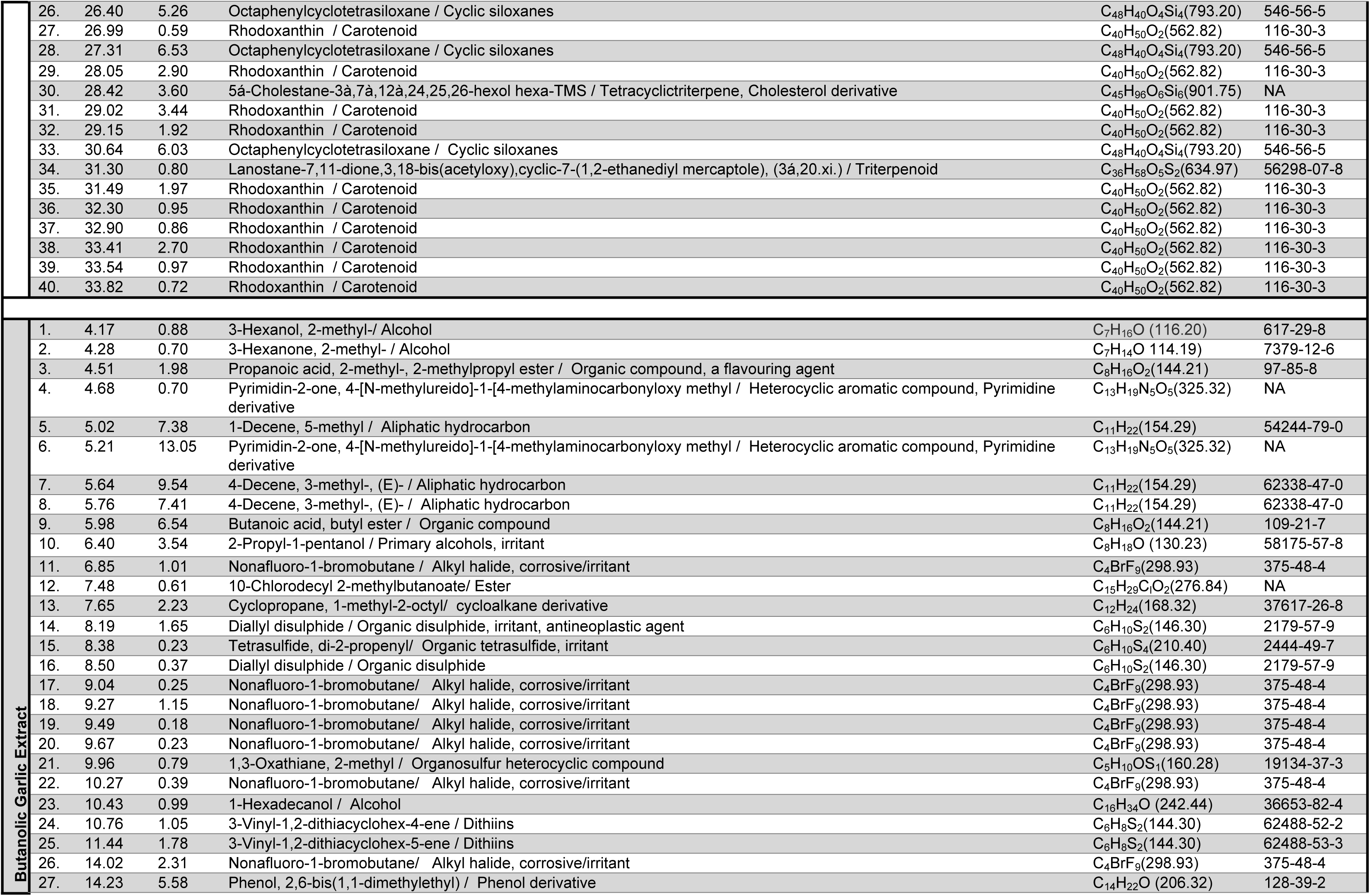
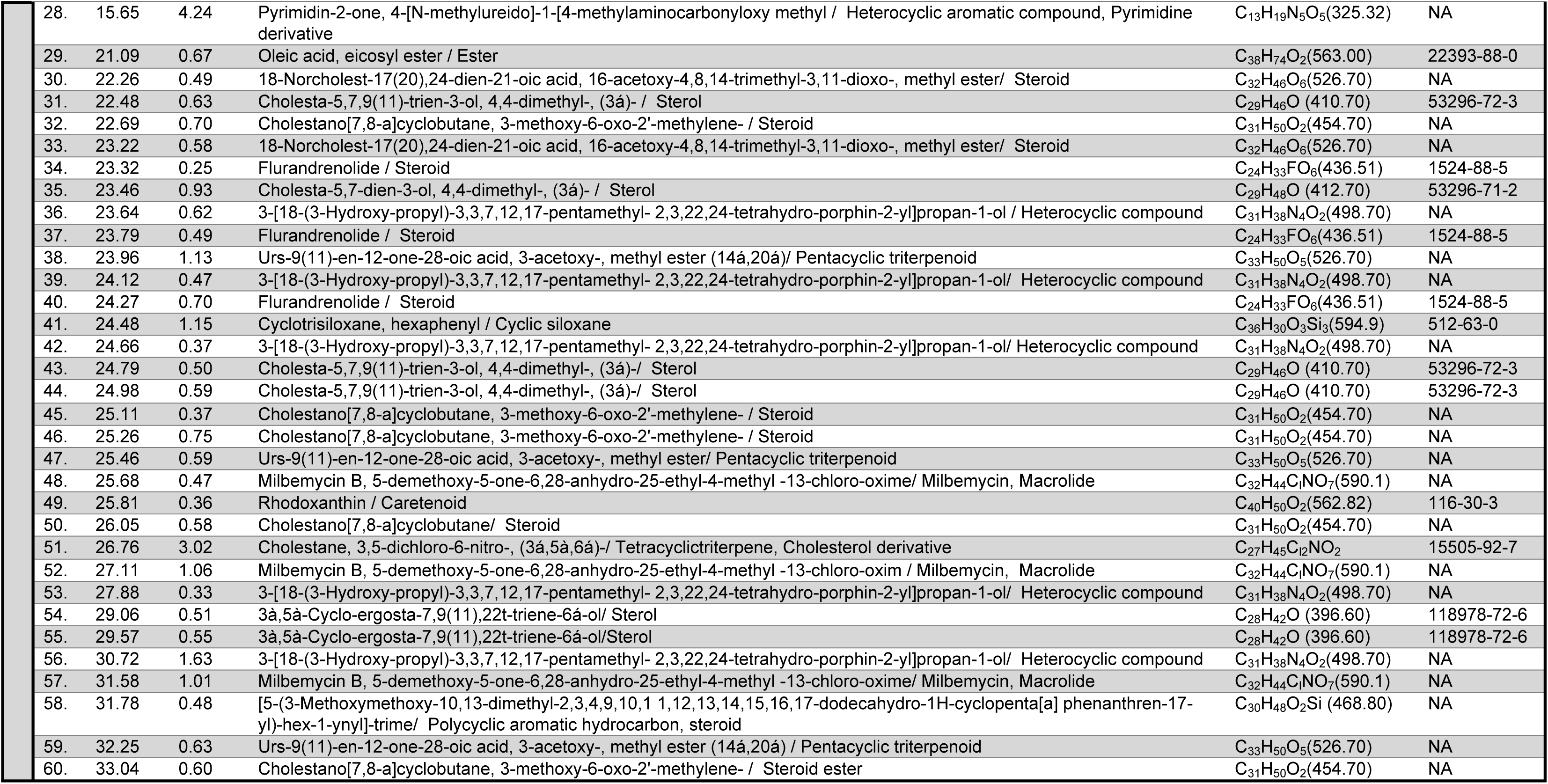
List of compounds predicted in bioactive fractions G1 and G2 of AGE and the butanolic extract of garlic. The ion fragments generated during GC-MS (Thermo Scientific TSQ 8000 Gas Chromatograph - Mass Spectrometer) analysis (Sup. Figure 3) were searched in NIST 2.0 library. Compounds predicted in the bioactive fractions with potential antimicrobial activity included phthalic acid derivatives, acid esters, phenyl group containing compounds, steroids etc (see Sup Figure. 4 and 5). The presence of various allyl derivatives as well as multiple S-containing compounds was indicated in the butanolic extract of garlic (see Sup. Figure 6).

**Table 2.**
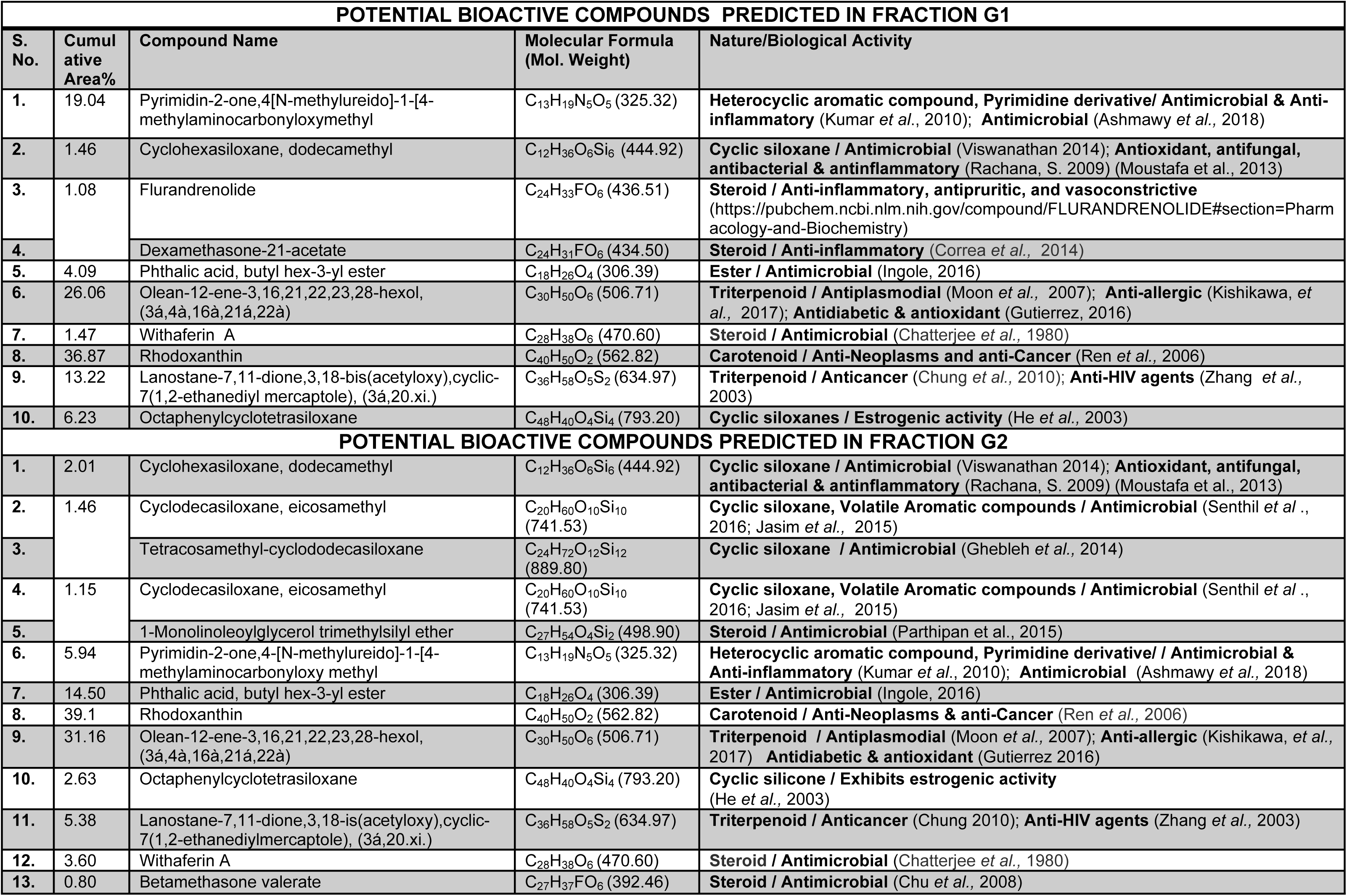
List of potentially bioactive compounds predicted in bioactive fractions G1 and G2 of AGE with relevant activity and references

| POTENTIAL BIOACTIVE COMPOUNDS PREDICTED IN FRACTION G1 |  |  |  |  |
| --- | --- | --- | --- | --- |
| S. No. | Cumulative Area% | Compound Name | Molecular Formula (Mol. Weight) | Nature/Biological Activity |
| 1. | 19.04 | Pyrimidin-2-one,4-[N-methylureido]-1-[4-methylaminocarbonyloxymethyl | C<sub>13</sub>H<sub>19</sub>N<sub>5</sub>O<sub>5</sub> (325.32) | **Heterocyclic aromatic compound, Pyrimidine derivative/ Antimicrobial & Anti-inflammatory** (Kumar *et al.*, 2010); **Antimicrobial** (Ashmawy *et al.*, 2018) |
| 2. | 1.46 | Cyclohexasiloxane, dodecamethyl | C<sub>12</sub>H<sub>36</sub>O<sub>6</sub>Si<sub>6</sub> (444.92) | **Cyclic siloxane / Antimicrobial** (Viswanathan 2014); **Antioxidant, antifungal, antibacterial & antiinflammatory** (Rachana, S. 2009) (Moustafa *et al.*, 2013) |
| 3. | 1.08 | Flurandrenolide | C<sub>24</sub>H<sub>33</sub>FO<sub>6</sub> (436.51) | **Steroid / Anti-inflammatory, antipruritic, and vasoconstrictive** (https://pubchem.ncbi.nlm.nih.gov/compound/FLURANDRENOLIDE#section=Pharmacology-and-Biochemistry) |
| 4. |  | Dexamethasone-21-acetate | C<sub>24</sub>H<sub>31</sub>FO<sub>6</sub> (434.50) | **Steroid / Anti-inflammatory** (Correa *et al.*, 2014) |
| 5. | 4.09 | Phthalic acid, butyl hex-3-yl ester | C<sub>18</sub>H<sub>26</sub>O<sub>4</sub> (306.39) | **Ester / Antimicrobial** (Ingole, 2016) |
| 6. | 26.06 | Olean-12-ene-3,16,21,22,23,28-hexol, (3á,4à,16à,21á,22à) | C<sub>30</sub>H<sub>50</sub>O<sub>6</sub> (506.71) | **Triterpenoid / Antiplasmodial** (Moon *et al.*, 2007); **Anti-allergic** (Kishikawa, *et al.*, 2017); **Antidiabetic & antioxidant** (Gutierrez, 2016) |
| 7. | 1.47 | Withaferin A | C<sub>28</sub>H<sub>38</sub>O<sub>6</sub> (470.60) | **Steroid / Antimicrobial** (Chatterjee *et al.*, 1980) |
| 8. | 36.87 | Rhodoxanthin | C<sub>40</sub>H<sub>50</sub>O<sub>2</sub> (562.82) | **Carotenoid / Anti-Neoplasms and anti-Cancer** (Ren *et al.*, 2006) |
| 9. | 13.22 | Lanostane-7,11-dione,3,18-bis(acetyloxy),cyclic-7(1,2-ethanediyl mercaptol), (3á,20.xi.) | C<sub>36</sub>H<sub>58</sub>O<sub>5</sub>S<sub>2</sub> (634.97) | **Triterpenoid / Anticancer** (Chung *et al.*, 2010); **Anti-HIV agents** (Zhang *et al.*, 2003) |
| 10. | 6.23 | Octaphenylcyclotetrasiloxane | C<sub>48</sub>H<sub>40</sub>O<sub>4</sub>Si<sub>4</sub> (793.20) | **Cyclic siloxanes / Estrogenic activity** (He *et al.*, 2003) |
| POTENTIAL BIOACTIVE COMPOUNDS PREDICTED IN FRACTION G2 |  |  |  |  |
| 1. | 2.01 | Cyclohexasiloxane, dodecamethyl | C<sub>12</sub>H<sub>36</sub>O<sub>6</sub>Si<sub>6</sub> (444.92) | **Cyclic siloxane / Antimicrobial** (Viswanathan 2014); **Antioxidant, antifungal, antibacterial & antiinflammatory** (Rachana, S. 2009) (Moustafa *et al.*, 2013) |
| 2. | 1.46 | Cyclodecasiloxane, eicosamethyl | C<sub>20</sub>H<sub>60</sub>O<sub>10</sub>Si<sub>10</sub> (741.53) | **Cyclic siloxane, Volatile Aromatic compounds / Antimicrobial** (Senthil *et al.*, 2016; Jasim *et al.*, 2015) |
| 3. |  | Tetracosamethyl-cyclododecasiloxane | C<sub>24</sub>H<sub>72</sub>O<sub>12</sub>Si<sub>12</sub> (889.80) | **Cyclic siloxane / Antimicrobial** (Ghebleh *et al.*, 2014) |
| 4. | 1.15 | Cyclodecasiloxane, eicosamethyl | C<sub>20</sub>H<sub>60</sub>O<sub>10</sub>Si<sub>10</sub> (741.53) | **Cyclic siloxane, Volatile Aromatic compounds / Antimicrobial** (Senthil *et al.*, 2016; Jasim *et al.*, 2015) |
| 5. |  | 1-Monolinoleoylglycerol trimethylsilyl ether | C<sub>27</sub>H<sub>54</sub>O<sub>4</sub>Si<sub>2</sub> (498.90) | **Steroid / Antimicrobial** (Parthipan *et al.*, 2015) |
| 6. | 5.94 | Pyrimidin-2-one,4-[N-methylureido]-1-[4-methylaminocarbonyloxy methyl | C<sub>13</sub>H<sub>19</sub>N<sub>5</sub>O<sub>5</sub> (325.32) | **Heterocyclic aromatic compound, Pyrimidine derivative/ Antimicrobial & Anti-inflammatory** (Kumar *et al.*, 2010); **Antimicrobial** (Ashmawy *et al.*, 2018) |
| 7. | 14.50 | Phthalic acid, butyl hex-3-yl ester | C<sub>18</sub>H<sub>26</sub>O<sub>4</sub> (306.39) | **Ester / Antimicrobial** (Ingole, 2016) |
| 8. | 39.1 | Rhodoxanthin | C<sub>40</sub>H<sub>50</sub>O<sub>2</sub> (562.82) | **Carotenoid / Anti-Neoplasms & anti-Cancer** (Ren *et al.*, 2006) |
| 9. | 31.16 | Olean-12-ene-3,16,21,22,23,28-hexol, (3á,4à,16à,21á,22à) | C<sub>30</sub>H<sub>50</sub>O<sub>6</sub> (506.71) | **Triterpenoid / Antiplasmodial** (Moon *et al.*, 2007); **Anti-allergic** (Kishikawa, *et al.*, 2017) **Antidiabetic & antioxidant** (Gutierrez 2016) |
| 10. | 2.63 | Octaphenylcyclotetrasiloxane | C<sub>48</sub>H<sub>40</sub>O<sub>4</sub>Si<sub>4</sub> (793.20) | **Cyclic silicone / Exhibits estrogenic activity** (He *et al.*, 2003) |
| 11. | 5.38 | Lanostane-7,11-dione,3,18-is(acetyloxy),cyclic-7(1,2-ethanediylmercaptol), (3á,20.xi.) | C<sub>36</sub>H<sub>58</sub>O<sub>5</sub>S<sub>2</sub> (634.97) | **Triterpenoid / Anticancer** (Chung 2010); **Anti-HIV agents** (Zhang *et al.*, 2003) |
| 12. | 3.60 | Withaferin A | C<sub>28</sub>H<sub>38</sub>O<sub>6</sub> (470.60) | **Steroid / Antimicrobial** (Chatterjee *et al.*, 1980) |
| 13. | 0.80 | Betamethasone valerate | C<sub>27</sub>H<sub>37</sub>FO<sub>6</sub> (392.46) | **Steroid / Antimicrobial** (Chu *et al.*, 2008) |

The identity of the vouchered Garlic sample used in the study was also verified by sequencing of the *rbcL* gene fragment amplified from the Garlic sample DNA. The sequence of the amplified *rbcL* gene fragment (submitted to GenBank MH020774) displayed more than 99% identity to the gene bank deposited *rbcL* gene sequences of *Allium sativum* L., confirming the authenticity of the Garlic sample.

## 4. Discussion

Anthrax remains a disease of greater concern in areas having larger share of livestock but relatively poor veterinary healthcare infrastructure, *e.g.,* Indian subcontinent and Africa (Shadomy *et al*., 2016; Turnbull, 2008). Its reemergence in the areas previously declared anthrax free is also on the rise (Shadomy *et al*., 2016). It remains one of the top ten killer diseases of the livestock in India (Rahman, 2012). The condition in other regions of Indian subcontinent could be similar. The discovery of new antimicrobials (Blaskovich *et al*., 2017) or agents that may affect the pathogenicity of organisms by modulating the quorum sensing (Bhardwaj *et al*., 2013; Ta and Arnason, 2015) or cause loss of virulence plasmids would be useful in controlling the diseases (Dastidar *et al*., 2013; Molnar *et al*., 1992; Spengler *et al*., 2006). Plants are an important source of antimicrobial compounds for controlling diseases (Cowan, 1999; Dastidar *et al*., 2013; Imam *et al*., 2016; Mahady, 2005; Ta and Arnason, 2015). Identification of edible plants that may have anti-*B. anthracis* activity would have a direct impact on the prevention of anthrax and decreasing its occurrence in the poorer endemic regions.

The extracts from a number of plants, mostly employing organic solvent extraction, have been indicated in the literature to have anti-*B. anthracis* activity (Akinpelu *et al*., 2008; Elisha *et al*., 2016; Mbwambo *et al*., 2011; Moshi and Mbwambo, 2005; Taher *et al*., 2012). In the current study, several edible plants commonly found in the Indian subcontinent (Duke *et al*., 2002; U.S. Department of Agriculture, 1992-2016) were evaluated for anti-*B. anthracis* activity. We had consciously chosen water as the extractant for its expected ease of use in the field as compared to any other extractant. A few plants displayed water-extractable anti-*B. anthracis* activity including Garlic that had been indicated previously (Sasaki and Kita, 2003). Among the tested plant extracts, the aqueous extract from Garlic (AGE) was found to be the most potent that reproducibly inhibited *B. anthracis* growth. However, the extract employed in the current study seemed to be relatively less active as compared to that used by Sasaki and Kita, 2003. One of the reasons could be the difference in the bioactive content present in different preparations being compared, *i.e.,* fresh garlic *vs.* dried powder and the methodology employed. Several plants such as Turmeric, Onion and Tulsi, which are variously attributed to have potent antimicrobial/bactericidal activity against many pathogens were found to be relatively inactive in the form tested, while Neem and Mango seemed to be moderately active (Duke *et al*., 2002; U.S. Department of Agriculture, 1992-2016). In a previous study by Jagtap *et al.,* 2016, the Neem extract and the Black Mulberry extract were shown to have potent anti-*B. anthracis* activity when tested up to 24 h of incubation. The use of about six-fold less concentrated aqueous extracts in the current study could be the reason for our observations. Turmeric and Bermuda grass extracts did not show any anti-*B. anthracis* activity in our assays. However, the turmeric derived curcumin and its derivatives (Antonelli, *et al*., 2014) as well as a protein from Bermuda grass (Arora *et al*., 2005) had been previously shown to inhibit the activity of anthrax toxin molecules. Recently, Gul and Bakht, 2015, had also shown the differential antibacterial activity extraction in the turmeric extracts when water, methanol, hexane and ethyl acetate were used as the extractants. The reduced extraction of the supposed bioactive secondary metabolites may be one of the possible reason for the inactivity of the aqueous turmeric extract in our assays.

The vegetative cells of the *B. anthracis* are known to be responsible for the pathogenesis primarily through the production of the anthrax toxins. Any agent that may kill vegetative cells of *B. anthracis* (Committee on Prepositioned Medical Countermeasures for the Public, 2012; Omotade *et al*., 2014) or that do not allow germination of spores are potent anti- anthrax control agents (Akoachere *et al*., 2007). Among the tested aqueous extracts, the AGE (≥1.9% w/v) had displayed robust killing of the vegetative cells of *B. anthracis.* It decreased the cell counts by more than 6 logs within 6-12 h of exposure, indicating its potential suitability for anthrax control.

Virulence plasmids, pXO1 and pXO2 are responsible for the pathogenicity of *B. anthracis.* Curing of these plasmids makes them avirulent (Kaur *et al*., 2013; Levy *et al*., 2012). Some small molecules such as acridine dyes are known to cause plasmid loss at sub-inhibitory concentration making virulent strains non-pathogenic (Molnar *et al*., 1992; Mbwambo *et al*., 2011; Taher *et al*., 2012). The ability of any safer anti-*B. anthracis* agent that could effect the curing of the virulence plasmid(s) at sub-inhibitory concentration would further increase its usefulness in anthrax prevention. However, AGE was not found to promote plasmid curing both at sub-inhibitory and inhibitory concentrations, thus, negating any such prospect for Garlic.

Bioactive content identification or characterization of any plant origin product is required to credibly correlate the constituent and their effect to recommend its preventive or curative use (Bhardwaj *et al*., 2013; Cowan, 1999; Friedman, 2015; Guil-Guerrero *et al*., 2016). The attempt to characterize the bioactive constituents present in AGE with potential anti-*B. anthracis* activity revealed the presence of bioactive compounds previously reported in other plants (see Table 2). The predominant antimicrobial species included: Phthalic acid, butyl hex-3-yl ester (Ingole, 2016); Pyrimidin-2-one,4-[N-methylureido]-1-[4-methylaminocarbonyloxymethy (Kumar et al., 2010; Barreto et al., 2010; Ashmawy et al., 2018); Withaferin A (Chatterjee et al., 1980); Cyclodecasiloxane, eicosamethyl and Tetracosamethyl-cyclododecasiloxane (Senthil et al ., 2016; Jasim et al., 2015; Ghebleh et al., 2014); Cyclohexasiloxane, dodecamethyl (Viswanathan 2014; Moustafa et al., 2013); Betamethasone valerate (Chu et al., 2008) and 1- Monolinoleoylglycerol trimethylsilyl ether (Parthipan et al., 2015). The antimicrobial activity of the indicated compounds had been variously characterized in different target organisms including the multidrug-resistant clinical isolates (Kumar et al., 2010; Barreto et al., 2010; Ashmawy et al., 2018). Other compounds with relevant activities included - Olean-12-ene- 3,16,21,22,23,28-hexol, (3á,4à,16à,21á,22à) – a triterpenoid with antiplasmodial activity (Moon et al., 2007); Lanostane-7,11-dione,3,18-bis(acetyloxy),cyclic-7(1,2-ethanediylmercaptole), (3á,20.xi.) - an anticancer and anti-HIV agent; and Rhodoxanthin - a carotenoid with cancer-preventive compound (Chung et al., 2010; Ren et al., 2006). There were also many other compounds with anti-inflammatory, anti-pruritic, anti-diabetic, anti-allergic or vasoconstrictive actions (Table 3).The observed anti-*B. anthracis* activity of the AGE could be the result of one or more compounds predicted by GC-MS analysis.

Although garlic is widely used in the food in the cooked form, its intake in the uncooked raw form is relatively limited. The characterization of thermal stability of the anti-*B. anthracis* activity present in AGE indicated it to be relatively stable at room temperature but quite unstable at 100°C - the temperature generally achieved during cooking. Antimicrobial agents when present in combination(s) may interact synergistically or antagonistically (Acar, 2000; Weiss *et al*., 2015; Wolfart *et al*., 2006). This kind of information is needed for the successful usage of any new antimicrobial agent. When we tested the AGE’s nature of interaction with antibiotics approved (FDA) for anthrax prevention, it did not seem to antagonize their activity.

Immediate future work may be focused on testing the efficacy of AGE alone or in combination with other common food ingredients in decreasing the *B. anthracis* load in the spiked animal meat, and the characterization of the bioactive fraction(s). Although AGE did not seem to promote virulence plasmid loss in our assays, ascertaining its effect on the expression of virulence factors (*e.g.*, Protective Antigen, Lethal Factor, Edema Factor), and the quorum sensing would remain a priority to further explore its potential utility in the anthrax control. The evaluation of Garlic to prevent the occurrence of gastrointestinal anthrax and the assessment of AGE as an injectable for therapy - as it is well tolerated in the model animals (HMPC, 2016), would be some other areas of active exploration. Further work needs to be undertaken to explore the possibility of employing Garlic or AGE to decrease the incidences of anthrax in endemic areas.

## 5. Conclusion

The comparative evaluation of the aqueous extracts of fourteen common edible plants, which are traditionally employed for treating diarrhea, stomachache and anthrax-like symptoms (Supplementary Table 1), suggested that the anti-*B. anthracis* activity of the aqueous extract of *Allium sativum* L. (Garlic) cloves/bulbs to be superior to that displayed by the extracts of *Azadirachta indica* A. Juss. (Neem) and *Mangifera indica* L. leaves. Interestingly, the aqueous extract of *Curcuma longa* L. (Turmeric) rhizome that is indicated for anthrax control/treatment in traditional medicine, did not show anti-*B. anthracis* activity under our assay conditions. The aqueous garlic extract (AGE) displayed acceptable stability under conditions relevant to its use, such as exposure to higher temperature (upto 40°C), extremes of pH (2-8) and bile salts. It did not antagonize the activity of different FDA approved antibiotics recommended for anthrax control, suggesting its compatibility with available preventive and therapeutic medication. The characterization of the bioactive fraction of AGE indicated the presence of phthalic acid derivatives, acid esters, phenyl group containing compounds, steroids etc. as potential anti- microbial constituents. However, additional exploratory work needs to be undertaken to further characterize the bioactive molecules present in AGE, determine their mode of action(s) and ascertain the possible preventive application of AGE in controlling the anthrax incidences in the remote endemic areas.

## Acknowledgement

The work was supported by a research grant ‘Ramalingaswami Fellowship (BT/RLF/Re-Entry/50/2011)’ from DBT, India to SS. Partial funding support to the laboratory of SS from DST-PURSE through Panjab University, Chandigarh is also duly acknowledged. The research funding agencies had no role in study design, data collection and analysis, decision to publish, or preparation of the manuscript.

## Author Contributions and email addresses

Conceived and designed the experiments: SS; Performed the experiments: RK, AT; Analyzed the data: RK, AT, SS; Contributed reagents/materials/analysis tools/discussion: RK, AT, MM, IKM, RB and SS; Wrote the paper: SS.

---

**Supplementary Figure 1. Aqueous Garlic Extract (AGE) at sub-inhibitory concentration does not promote virulence plasmid ‘pXO1’ loss from *B. anthracis* Sterne strain. (A)** The PCR analysis of eight random colonies growing from 1% w/v AGE exposed 24 h old culture for the presence of *pagA* (pXO1-borne) and *phoP* (genome-borne) genes using gene-specific primers are shown. All colonies examined retained the pXO1 plasmid as suggested by PCR amplification of the *pagA* gene. Genomic DNA from *B. anthracis* Sterne strain was used as positive control and that of *Escherichia coli* DH5α strain as a negative control for the PCR based analysis. **(B)** Summary of the virulence plasmid pXO1 loss assay. *B. anthracis* cells seem to retain plasmid pXO1 on AGE exposure at all the time points tested (0 - 24 h).

**Supplementary Figure 2. Optimization of Thin Layer Chromatography (TLC) conditions in combination with bioautography to fractionate bioactive components present in Aqueous Garlic Extract (AGE)**. **(A)** Different solvent systems were evaluated for their ability to separate the components present in AGE. The thin layer of silica (TLC plate) was spotted with AGE followed by the development of chromatogram using the indicated solvents. The fluorescent spots fractionated in AGE chromatograms were visualized under UV-light. Some solvent combinations were found to better resolve the fluorescent spots than others (compare chromatogram no. 3 with others). **(B)** The bioautography of AGE components separated on silica TLC using solvent system Toluene: Acetone (7:3) showed inhibition of the growth of *Bacillus anthracis* culture at specific position (encircled in white in *right panel: Aqueous Garlic Extract or AGE TLC plate; chromatogram labeled 2*) that correlated with UV-fluorescent spots visible in the upper middle part of the chromatogram 3 in (A) while the TLC plate that had only extractant water spotted and treated the same way (Left panel: Control TLC plate; *chromatogram labeled 1*), did not generate any zone of growth inhibition (ZOI).

**Supplementary Figure 3: GC-MS analysis of the TLC separated bioactive fraction of Aqueous Garlic (*Allium sativum)* Extract (AGE) to identify potential bioactive compounds.** The UV- fluorescent bands in the AGE chromatogram (Silica TLC) that corresponded with anti-*Bacillus anthracis* activity (growth inhibition) on the bioautogram were labeled G1 and G2 **(A)**, isolated and processed for GC-MS analysis on Thermo Scientific TSQ 8000 Gas Chromatograph - Mass Spectrometer, using latest NIST 2.0 Library (**B and Sup. Figure 4 and 5**). The compounds with potential antimicrobial activity, *e.g.,* phthalic acid derivatives, acid esters, phenyl group-containing compounds, steroids, were detected in the bioactive fractions (Table 2**, Sup. Figure 4 and 5)**. However, the GC-MS analysis of butanolic extract of garlic **(C)** showed the presence of various allyl derivatives as well as multiple S-containing compounds (Table 1**, Sup. Figure 6)** as expected.

**Supplementary Figure 4. GC-MS profile of bioactive G1 spot from AGE and the list of compound hits**

**Supplementary Figure 5. GC-MS profile of bioactive G2 spot from AGE and the list of compound hits**

**Supplementary Figure 6. GC-MS profile of butanolic extract of Garlic and the list of compound hits**

**Supplementary Figure 7. Zone of inhibition (ZOI) of antibiotics used in anthrax control and AGE**

**Supplementary Table 1.**
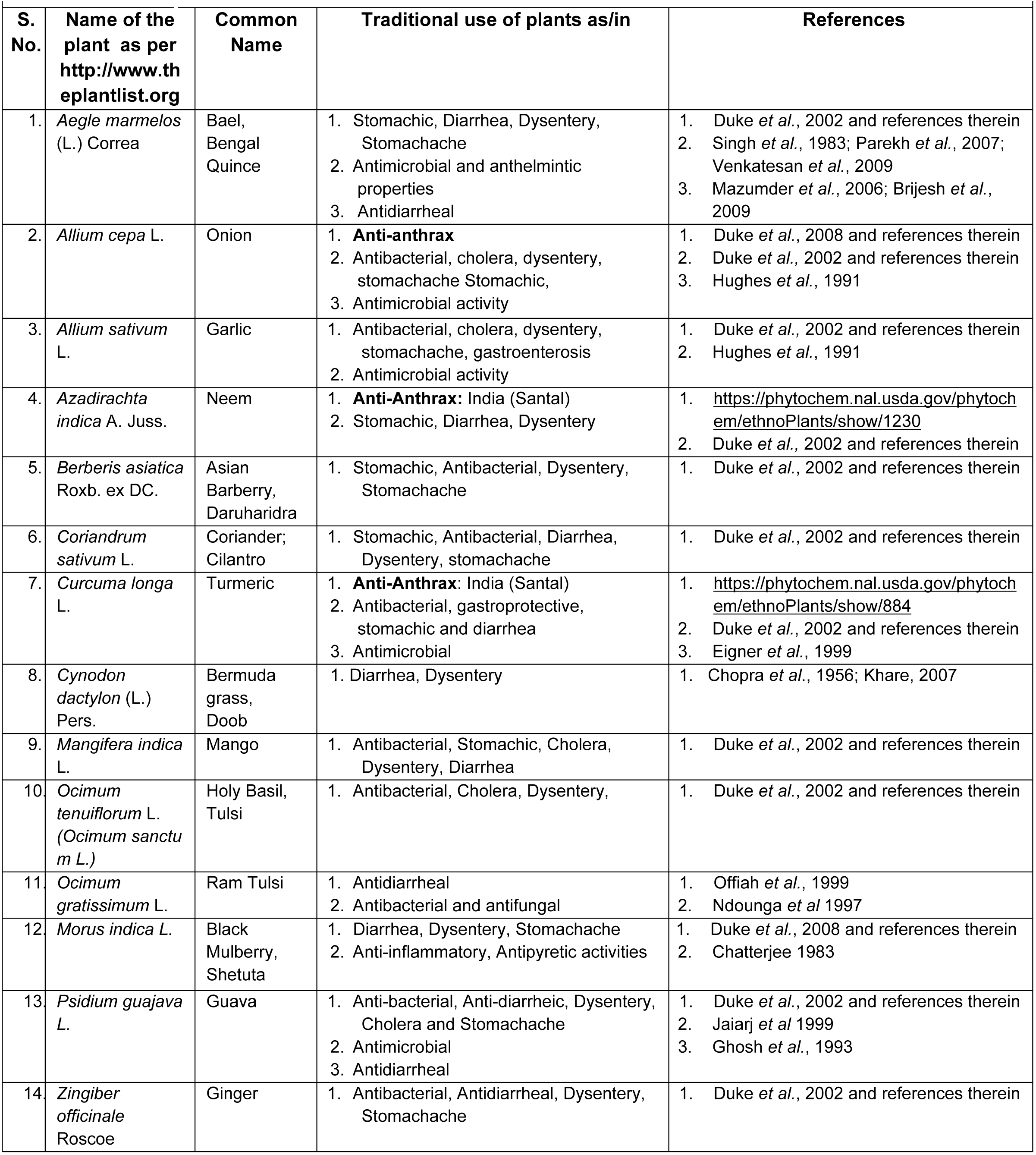
Traditional use of plants as indicated in the literature

**Supplementary Table 2.**
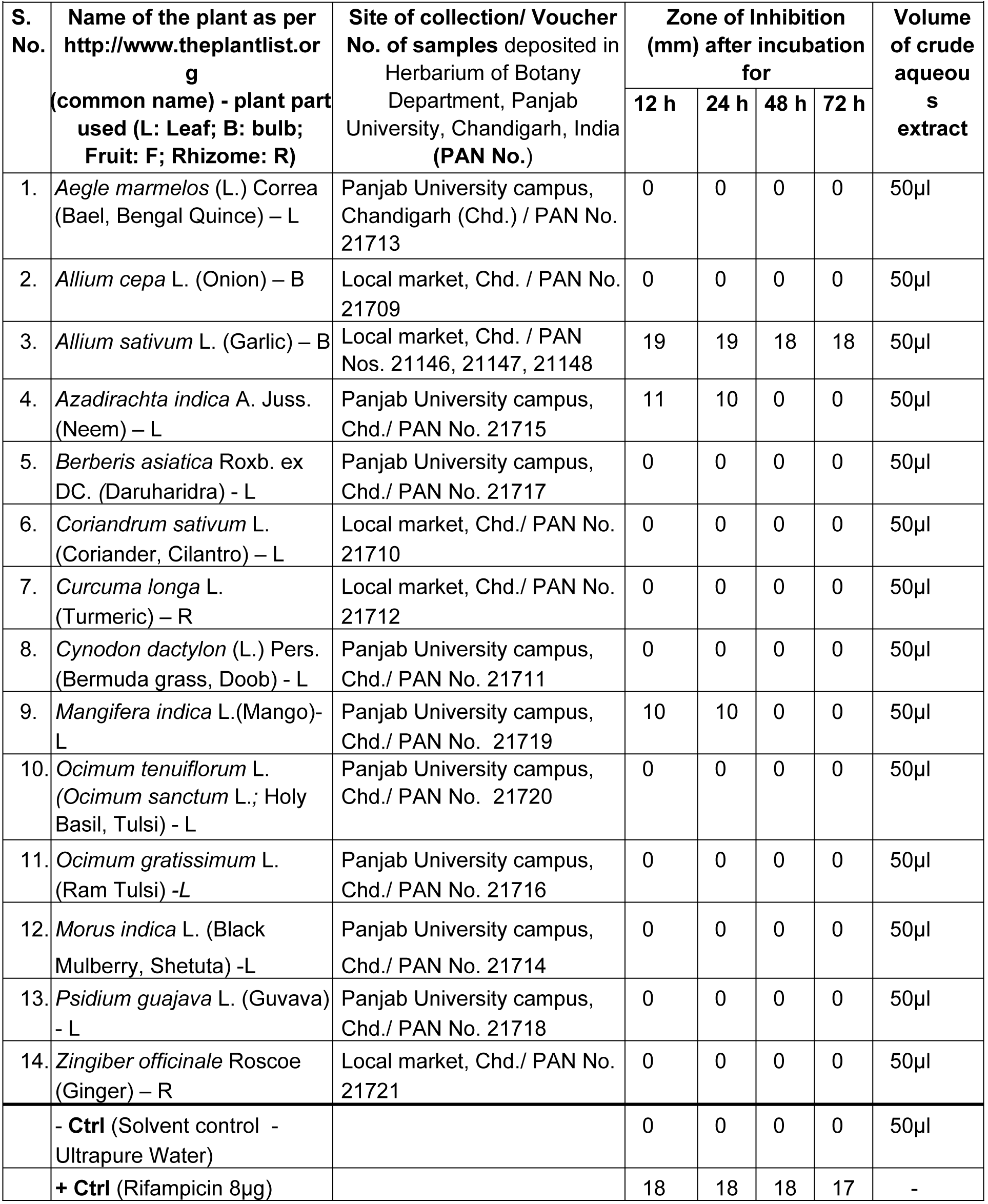
The aqueous extracts of different plants (40% w/v) differentially inhibited the growth of *B. anthracis* in agar-well diffusion assay (AWDA). The data shown are from one representative experiment. Antibiotic Rifampicin and ultrapure water were used as positive control (+ Ctrl) and solvent/diluent or negative control (- Ctrl). Note: Garlic displayed the highest concentration of water-soluble *anti-B. anthracis* activity constituents among the tested plants.

## Notes

### Competing Interest Statement

The authors have declared no competing interest.

### Summary of Updates

1)Graphical Overview of the manuscript included 2)Figures revised and updated. FICI value estimate for synergy with antibiotics used in anthrax control provided. 3)Table 1. List of compounds predicted in bioactive fractions G1 and G2 of AGE and the butanolic extract of garlic. -UPDATED 4) Table 2. Lists potentially bioactive compounds predicted in bioactive fractions G1 and G2 of AGE with relevant activity and references

