## Supplemental Files for "Common Garlic (*Allium sativum* L.) has Potent Anti-*Bacillus anthracis* Activity"

**Supplementary Table 1. Traditional use of plants as indicated in the literature**

**Supplementary Table 2.** The aqueous extracts of different plants (40% w/v) differentially inhibited the growth of *B. anthracis* in agar-well diffusion assay (AWDA). The data shown are from one representative experiment. Antibiotic Rifampicin and ultrapure water were used as positive control (+ Ctrl) and solvent/diluent or negative control (- Ctrl). Note: Garlic displayed the highest concentration of water-soluble *anti-B. anthracis* activity constituents among the tested plants.

**Supplementary Figure 1. Aqueous Garlic Extract (AGE) at sub-inhibitory concentration does not promote virulence plasmid 'pXO1' loss from *B. anthracis* Sterne strain. (A)** The PCR analysis of eight random colonies growing from 1% w/v AGE exposed 24 h old culture for the presence of *pagA* (pXO1-borne) and *phoP* (genome-borne) genes using gene-specific primers are shown. All colonies examined retained the pXO1 plasmid as suggested by PCR amplification of the *pagA* gene. Genomic DNA from *B. anthracis* Sterne strain was used as positive control and that of *Escherichia coli* DH5α strain as a negative control for the PCR based analysis. **(B)** Summary of the virulence plasmid pXO1 loss assay. *B. anthracis* cells seem to retain plasmid pXO1 on AGE exposure at all the time points tested (0 - 24 h).

**Supplementary Figure 2. Optimization of Thin Layer Chromatography (TLC) conditions in combination with bioautography to fractionate bioactive components present in Aqueous Garlic Extract (AGE)**. **(A)** Different solvent systems were evaluated for their ability to separate the components present in AGE. The thin layer of silica (TLC plate) was spotted with AGE followed by the development of chromatogram using the indicated solvents. The fluorescent spots fractionated in AGE chromatograms were visualized under UV-light. Some solvent combinations were found to better resolve the fluorescent spots than others (compare chromatogram no. 3 with others). **(B)** The bioautography of AGE components separated on silica TLC using solvent system Toluene: Acetone (7:3) showed inhibition of the growth of *Bacillus anthracis* culture at specific position (encircled in white in *right panel: Aqueous Garlic Extract or AGE TLC plate; chromatogram labeled 2*) that correlated with UV-fluorescent spots visible in the upper middle part of the chromatogram 3 in (A) while the TLC plate that had only extractant water spotted and treated the same way (Left panel: Control TLC plate; *chromatogram labeled 1*), did not generate any zone of growth inhibition (ZOI).

**Supplementary Figure 3: GC-MS analysis of the TLC separated bioactive fraction of Aqueous Garlic (*Allium sativum)* Extract (AGE) to identify potential bioactive compounds.** The UV- fluorescent bands in the AGE chromatogram (Silica TLC) that corresponded with anti-*Bacillus anthracis* activity (growth inhibition) on the bioautogram were labeled G1 and G2 **(A)**, isolated and processed for GC-MS analysis on Thermo Scientific TSQ 8000 Gas Chromatograph - Mass Spectrometer, using latest NIST 2.0 Library (**B and Sup. Figure 4 and 5**). The compounds with potential antimicrobial activity, *e.g.,* phthalic acid derivatives, acid esters, phenyl group-containing compounds, steroids, were detected in the bioactive fractions (**Table 2, Sup. Figure 4 and 5**). However, the GC-MS analysis of butanolic extract of garlic **(C)** showed the presence of various allyl derivatives as well as multiple S-containing compounds (**Table 1, Sup. Figure 6**) as expected.

**Supplementary Figure 4. GC-MS profile of bioactive G1 spot from AGE and the list of compound hits**

**Supplementary Figure 5. GC-MS profile of bioactive G2 spot from AGE and the list of compound hits**

**Supplementary Figure 6. GC-MS profile of butanolic extract of Garlic and the list of compound hits**

**Supplementary Figure 7. Zone of inhibition (ZOI) of antibiotics used in anthrax control and AGE**

A.

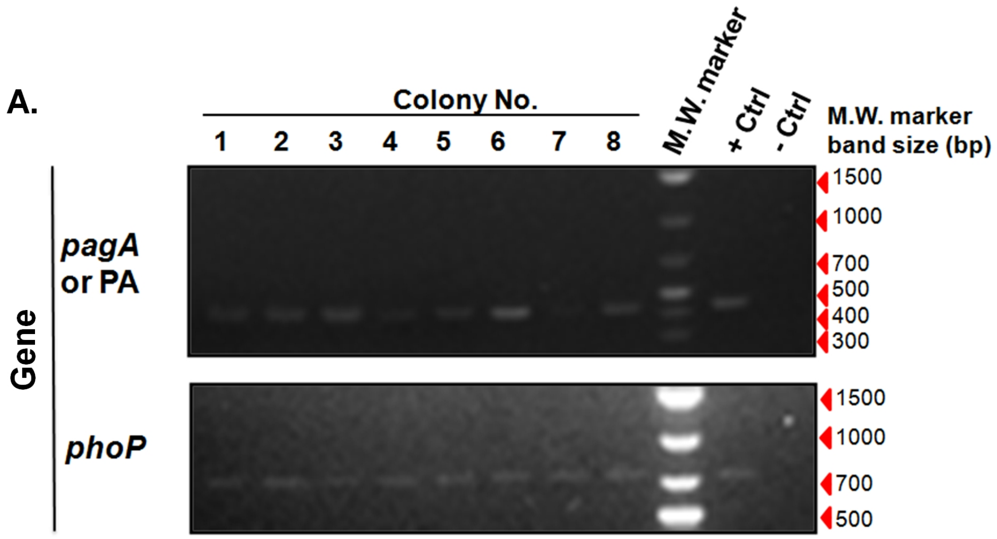


#### B. Summary of Virulence Plasmid Loss Assay

| *Genes* tested (borne) | *Bacillus anthracis* cells exposed to AGE (hrs) | | | | | [1]+ve control | [2]-ve control |
|---|---|---|---|---|---|---|---|
| | 0 hr | 3 hr | 6 hr | 12hr | 24hr | | |
| ***pagA* or *PA* gene (pXO1) detected in no. of colonies/ Total no. of colonies tested** | **10/10** | **10/10** | **9/10** | **10/10** | **8/8** | **1/1** | **0/1** |
| ***phoP* gene (Chromosome) detected in no. of colonies/ Total no. of colonies tested** | **10/10** | **10/10** | **10/10** | **10/10** | **8/8** | **1/1** | **0/1** |

[1]Untreated *Bacillus anthracis* Sterne Strain
[2] *Escherichia coli DH5*$\alpha$

**Supplementary Figure 1. Aqueous Garlic Extract (AGE) at sub-inhibitory concentration does not promote virulence plasmid 'pXO1' loss from B. anthracis Sterne strain.**

**A.**

**Thin Layer Chromatography (TLC) fractionation of AGE**

| CHROMATOGRAM NO. | SOLVENT SYSTEM EMPLOYED FOR DEVELOPING AGE CHROMATOGRAM |
|---|---|
| 1 | CHLOROFORM:METHANOL (7: 3) |
| 2 | TOLUENE:ACETIC ACID (7:3) |
| 3 | TOLUENE:ACETONE (7:3) |
| 4 | HEXANE:ETHYL ACETATE (5:5) |
| 5 | ETHANOL:WATER (5:5) |
| 6 | ETHYL ACETATE:ISOPROPANOL:ACETIC ACID:WATER (16 : 2 : 1: 1) |
| 7 | NEGATIVE CONTROL (without extract) developed with TOLUENE:ACETIC ACID (7:3) |

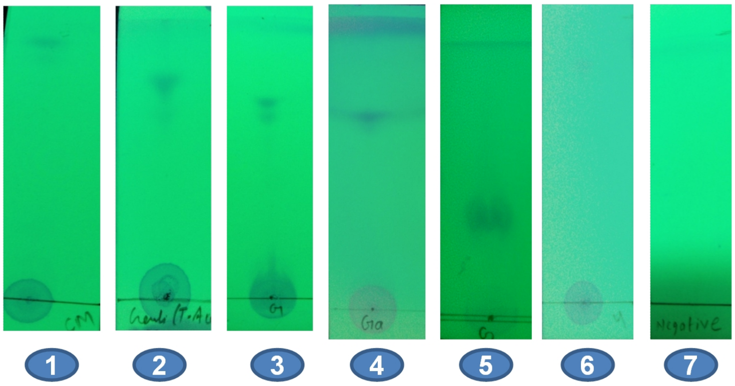


**Chromatogram No.**

**B.**

**Control TLC Plate** **AGE TLC Plate**

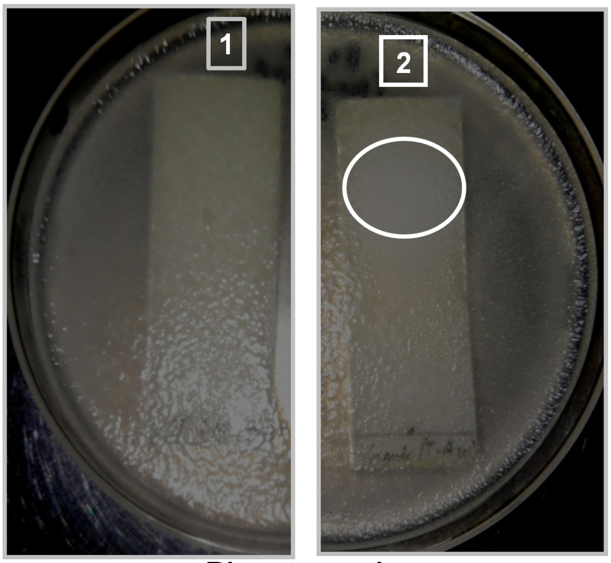


**Bioautography**

**Supplementary Figure 2. Optimization of Thin Layer Chromatography (TLC) conditions in combination with bioautography to fractionate bioactive components present in Aqueous Garlic Extract (AGE).**

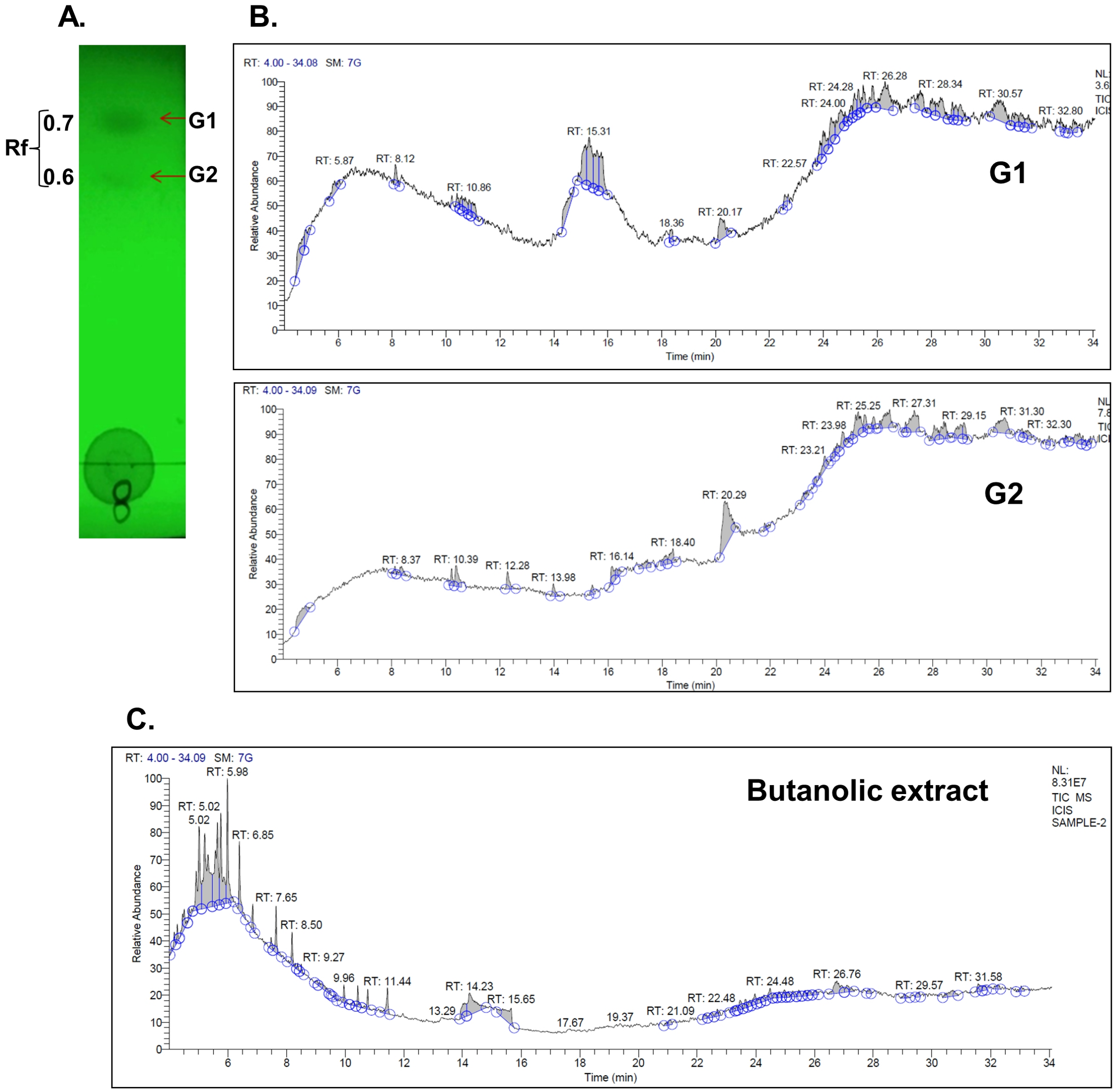


**Supplementary Figure 3: GC-MS analysis of the TLC separated bioactive fraction of Aqueous Garlic (Allium sativum) Extract (AGE) to identify potential bioactive compounds.**

**Supplementary Figure 4. GC-MS profile of bioactive G1 spot from AGE and the list of compound hits**

CIL/ SAIF Panjab University Chandigarh

#### Sample Header

| | |
|---|---|
| Data File: | 1 |
| Original Data Path: | C:\GCMS-data\YEAR 2016\NOV\24 |
| Sample Type: | Unknown |
| Sample ID: | 1 |
| Sample Name: | |
| Acquisition Date: | 11/24/16 11:10:32 AM |
| Run Time(min): | 30.08 |
| Injection Volume(μl): | 1.00 |
| Scans: | 3581 |
| Low Mass(m/z): | 100 |
| High Mass(m/z): | 800 |
| Instrument Method: | C:\GCMS-data\instrument method\GERNAL-gcms-METHOD.meth |

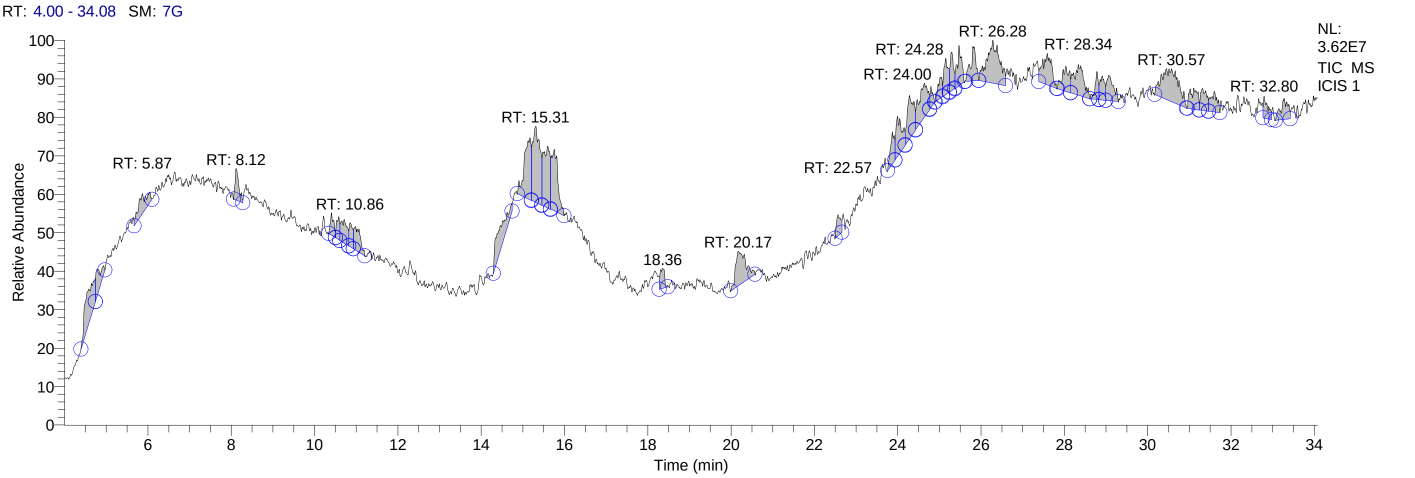


#### Qual Peak Table

| RT | Peak Area | Area % | Peak Height |
|---|---|---|---|
| 4.54 | 50215465.79 | 3.78 | 3459029.01 |
| 4.77 | 19868622.03 | 1.50 | 2602203.60 |
| 5.87 | 26690462.79 | 2.01 | 1897500.40 |
| 8.12 | 19422410.96 | 1.46 | 2991066.34 |
| 10.41 | 11770572.82 | 0.89 | 2115650.07 |
| 10.58 | 10609622.73 | 0.80 | 2159841.02 |
| 10.71 | 22480579.67 | 1.69 | 2238591.31 |
| 10.86 | 12950805.48 | 0.98 | 2251129.90 |
| 10.98 | 24378258.27 | 1.84 | 2224717.32 |
| 14.34 | 44930710.44 | 3.39 | 2688254.91 |
| 15.17 | 64982419.78 | 4.90 | 5865729.42 |
| 15.31 | 88326910.80 | 6.66 | 7131681.61 |
| 15.60 | 64921386.55 | 4.89 | 5848119.36 |
| 15.75 | 62800563.55 | 4.73 | 5817789.03 |
| 18.36 | 14363245.10 | 1.08 | 1878579.09 |
| 20.17 | 54257463.14 | 4.09 | 3269071.38 |
| 22.57 | 13419170.74 | 1.01 | 1990180.73 |
| 23.87 | 19510674.16 | 1.47 | 3051204.17 |
| 24.00 | 35743472.56 | 2.69 | 3780112.14 |
| 24.28 | 42914206.77 | 3.23 | 4107692.11 |
| 24.63 | 48696955.45 | 3.67 | 3206897.18 |
| 24.80 | 8111362.98 | 0.61 | 1938725.44 |
| 25.01 | 13276406.50 | 1.00 | 1963586.13 |
| 25.16 | 23757754.92 | 1.79 | 3450475.51 |
| 25.29 | 21330505.87 | 1.61 | 3651104.69 |
| 25.47 | 27463247.82 | 2.07 | 3796576.61 |
| 25.81 | 29584362.85 | 2.23 | 3244752.23 |
| 26.28 | 82695470.00 | 6.23 | 4021840.37 |

### CIL/ SAIF Panjab University Chandigarh

| RT | Peak Area | Area % | Peak Height |
|---|---|---|---|
| 27.59 | 43793903.44 | 3.30 | 2953061.54 |
| 28.04 | 27030052.36 | 2.04 | 2382500.78 |
| 28.34 | 46713122.64 | 3.52 | 2895070.96 |
| 28.78 | 14206210.68 | 1.07 | 2597907.35 |
| 28.91 | 17279140.31 | 1.30 | 2249040.41 |
| 29.07 | 27121123.14 | 2.04 | 2397996.13 |
| 30.57 | 85918529.48 | 6.47 | 3088105.04 |
| 31.11 | 25722608.39 | 1.94 | 1999242.91 |
| 31.32 | 20694823.42 | 1.56 | 2080524.93 |
| 31.61 | 20077020.33 | 1.51 | 1925625.82 |
| 32.80 | 15284543.14 | 1.15 | 2077950.48 |
| 33.28 | 23818010.81 | 1.79 | 2019691.86 |

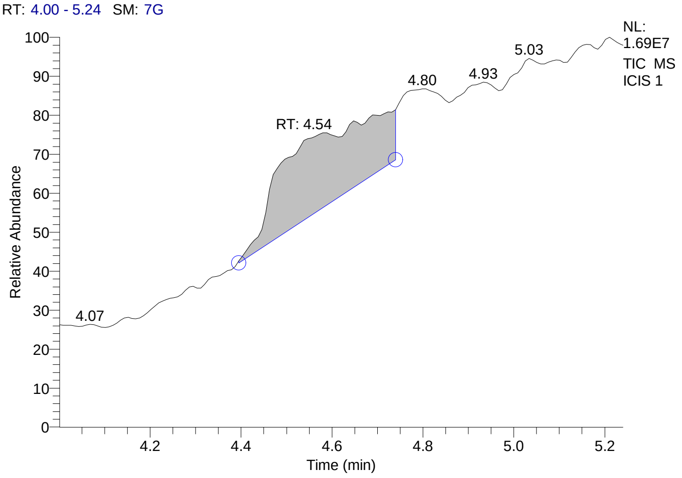


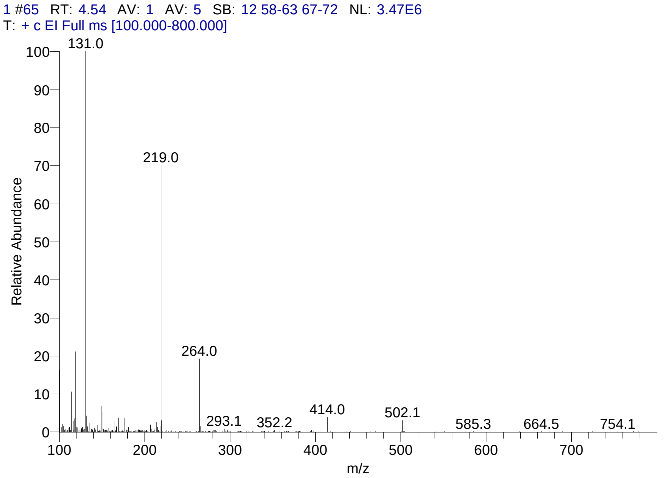


#### Library Search Results Table

| Compound Name | RT | Molecular Formula | Cas # |
|---|---|---|---|
| Perfluorotributylamine | 4.54 | C12F27N | 311-89-7 |
| Perfluoro(dibutylmethylamine) | 4.54 | C9F21N | 514-03-4 |
| Pyrimidin-2-one, 4-[N-methylureido]-1-[4-methylaminocarbonyloxy methyl | 4.54 | C13H19N5O5 | NA |

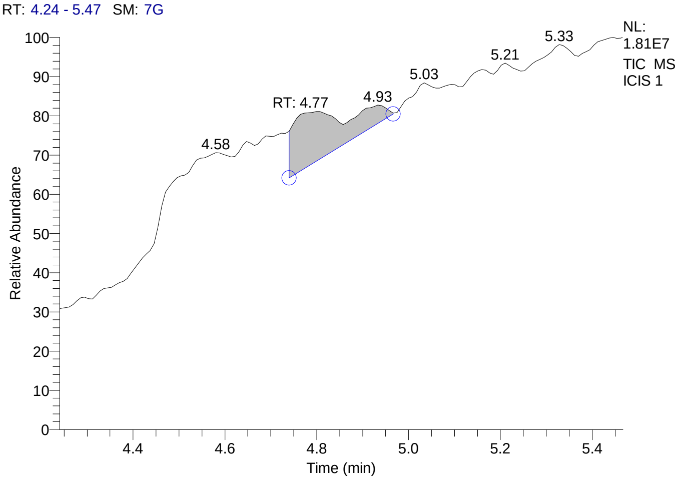


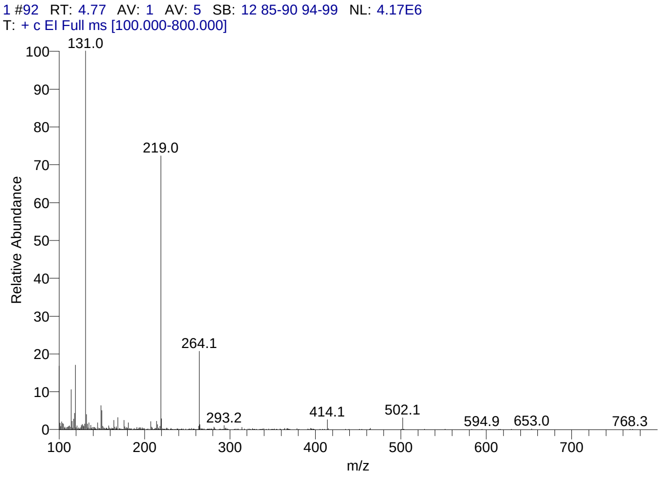

### CIL/ SAIF Panjab University Chandigarh

#### Library Search Results Table

| Compound Name | RT | Molecular Formula | Cas # |
|---|---|---|---|
| Perfluorotributylamine | 4.77 | C12F27N | 311-89-7 |
| Perfluoro(dibutylmethylamine) | 4.77 | C9F21N | 514-03-4 |
| Pyrimidin-2-one, 4-[N-methylureido]-1-[4-methylaminocarbonyloxy methyl | 4.77 | C13H19N5O5 | NA |

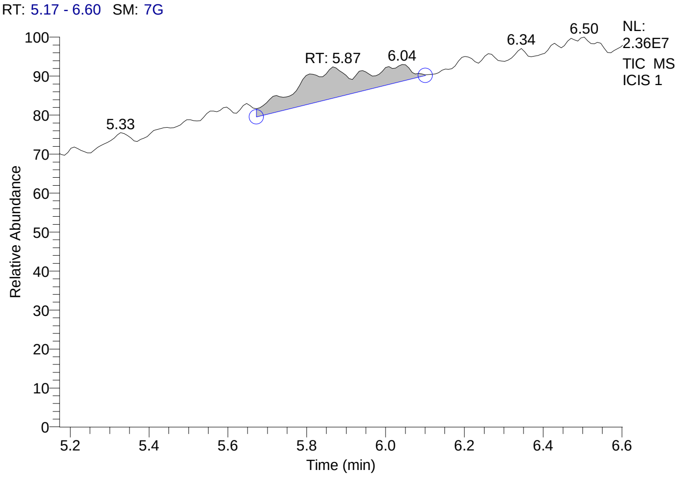


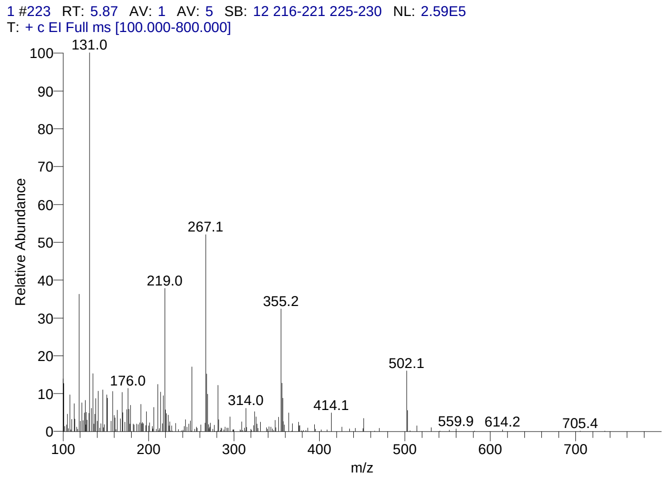


#### Library Search Results Table

| Compound Name | RT | Molecular Formula | Cas # |
|---|---|---|---|
| Benzoic acid, 2,6-bis[(trimethylsilyl)oxy]-, trimethylsilyl ester | 5.87 | C16H30O4Si3 | 3782-85-2 |
| Cyclopentasiloxane, decamethyl- | 5.87 | C10H30O5Si5 | 541-02-6 |
| Benzoic acid, 2,5-bis(trimethylsiloxy)-, trimethylsilyl ester | 5.87 | C16H30O4Si3 | 3618-20-0 |

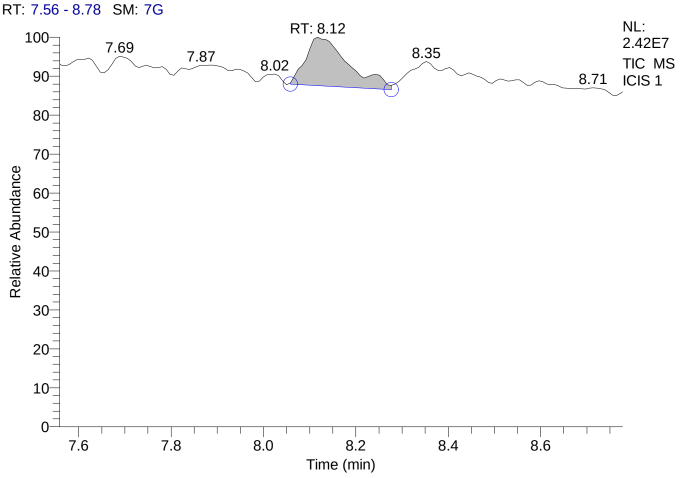


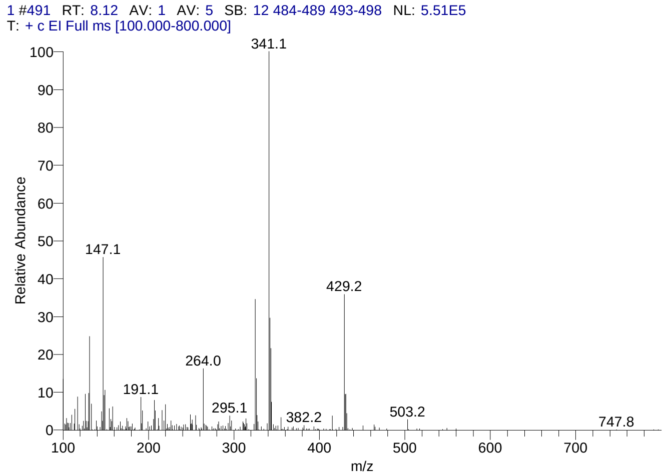


#### Library Search Results Table

| Compound Name | RT | Molecular Formula | Cas # |
|---|---|---|---|
| Cyclohexasiloxane, dodecamethyl- | 8.12 | C12H36O6Si6 | 540-97-6 |
| Fluoren-9-ol, 3,6-dimethoxy-9-(2-phenylethynyl)- | 8.12 | C23H18O3 | NA |
| Heptasiloxane, 1,1,3,3,5,5,7,7,9,9,11,11,13,13-tetradecamethyl- | 8.12 | C14H44O6Si7 | 19095-23-9 |

### CIL/ SAIF Panjab University Chandigarh

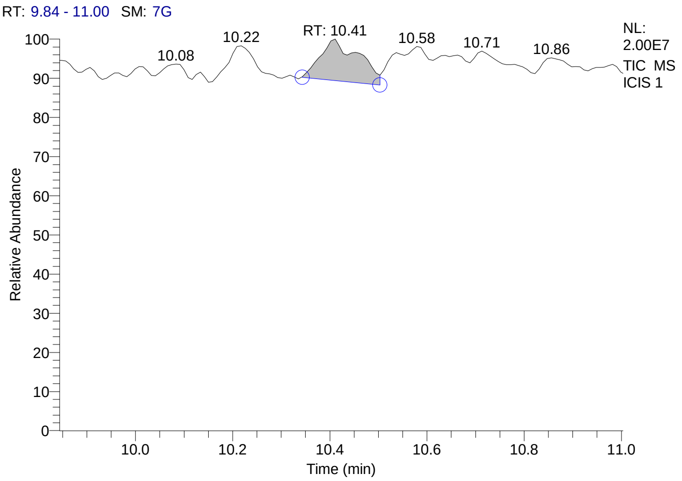


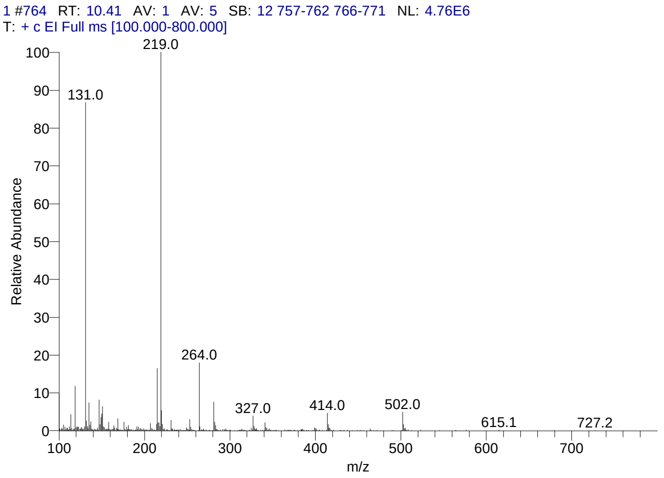


#### Library Search Results Table

| Compound Name | RT | Molecular Formula | Cas # |
|---|---|---|---|
| Perfluorotributylamine | 10.41 | C12F27N | 311-89-7 |
| 5á-Cholestane-3à,7à,12à,24,25,26-hexol hexa-TMS | 10.41 | C45H96O6Si6 | NA |
| Pyrimidin-2-one, 4-[N-methylureido]-1-[4-methylaminocarbonyloxy methyl | 10.41 | C13H19N5O5 | NA |

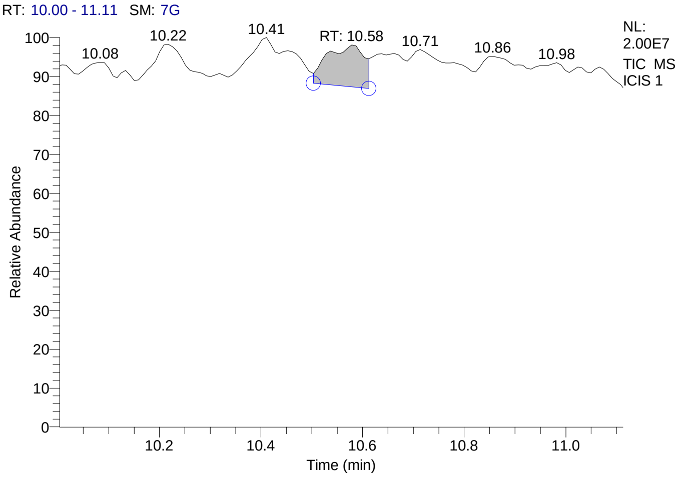


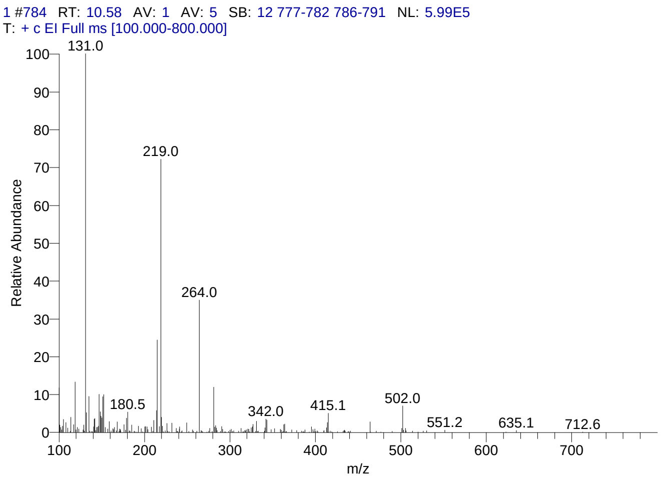


#### Library Search Results Table

| Compound Name | RT | Molecular Formula | Cas # |
|---|---|---|---|
| Perfluorotributylamine | 10.58 | C12F27N | 311-89-7 |
| 5á-Cholestane-3à,7à,12à,24,25,26-hexol hexa-TMS | 10.58 | C45H96O6Si6 | NA |
| Dodecanoic acid, tricosafluoro- | 10.58 | C12HF23O2 | 307-55-1 |

### CIL/ SAIF Panjab University Chandigarh

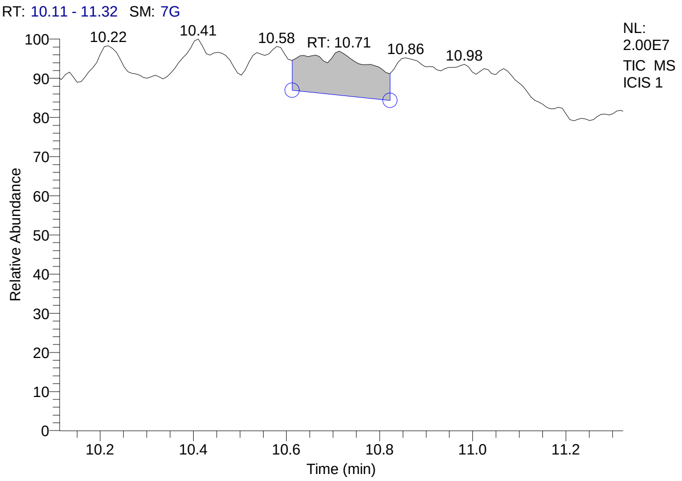


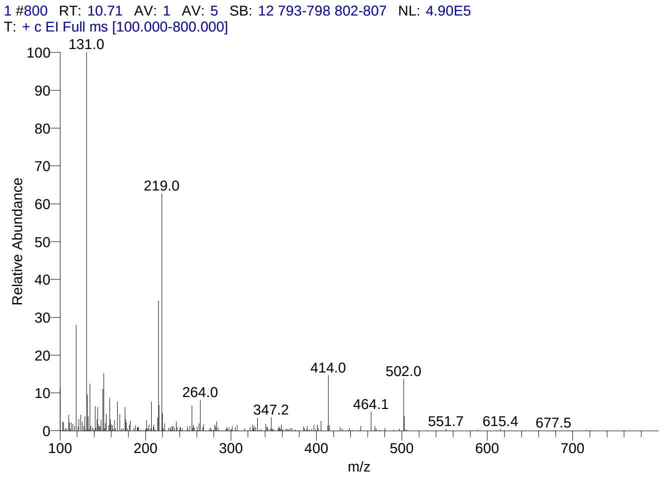


#### Library Search Results Table

| Compound Name | RT | Molecular Formula | Cas # |
|---|---|---|---|
| Perfluorotributylamine | 10.71 | C12F27N | 311-89-7 |
| 1,3,4(6H)- Thiadiazin-2-amine, 6-ethyl-5-phenyl- | 10.71 | C11H13N3S | 54418-98-3 |
| Dodecanoic acid, tricosafluoro- | 10.71 | C12HF23O2 | 307-55-1 |

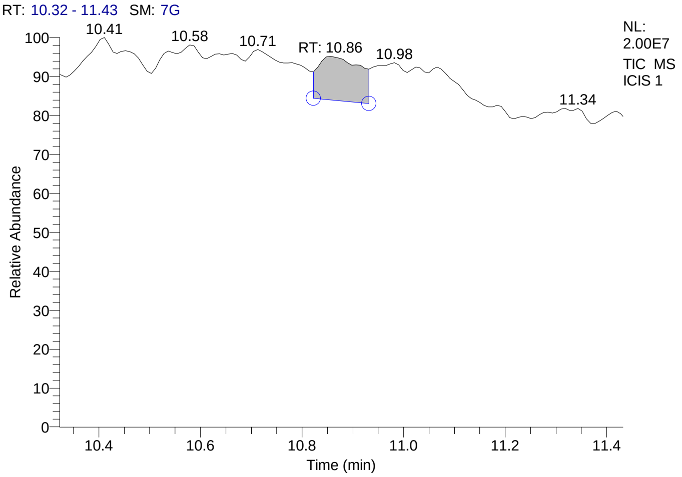


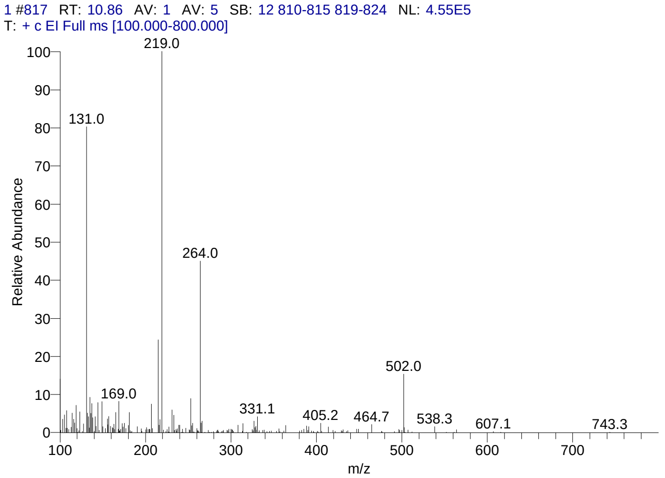


#### Library Search Results Table

| Compound Name | RT | Molecular Formula | Cas # |
|---|---|---|---|
| Perfluorotributylamine | 10.86 | C12F27N | 311-89-7 |
| 5á-Cholestane-3à,7à,12à,24,25,26-hexol hexa-TMS | 10.86 | C45H96O6Si6 | NA |
| Pyrimidin-2-one, 4-[N-methylureido]-1-[4-methylaminocarbonyloxy methyl | 10.86 | C13H19N5O5 | NA |

### CIL/ SAIF Panjab University Chandigarh

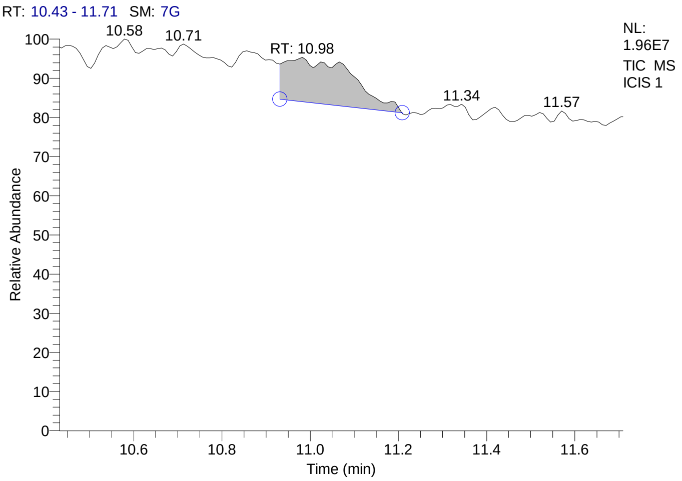


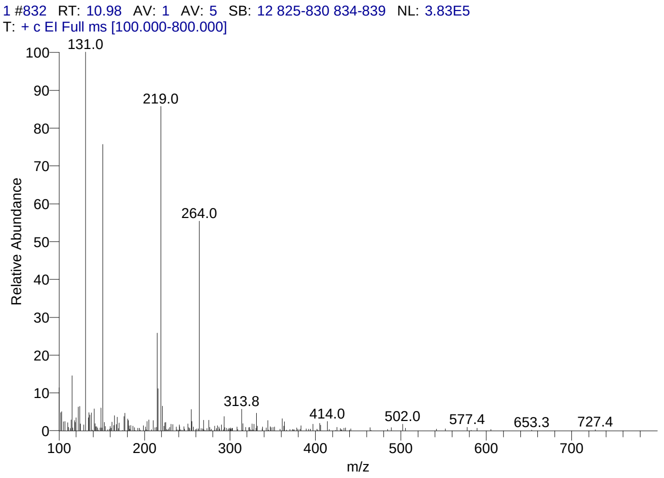


#### Library Search Results Table

| Compound Name | RT | Molecular Formula | Cas # |
|---|---|---|---|
| Cyclohexanecarboxylic acid, 1-(3,4-dimethoxyphenyl)-, (3-chloro-4-methylphenyl)amide | 10.98 | C22H26ClNO3 | NA |
| 4H-Cyclopropa[5',6']benz[1',2':7,8]azuleno[5,6-b]oxiren-4-one, 8-(acetyloxy)-1,1a,1b,1c,2a,3,3a,6a,6b,7,8,8a-dodecahydro-3a,6b,8a-trihydroxy-2a-(hydroxymethyl)-1,1,5,7-tetramethyl-, (1aà,1bá,1cá,2aá,3aá,6aà,6bà,7à,8á,8aà)- | 10.98 | C22H30O8 | 77646-23-2 |
| Pyrimidin-2-one, 4-[N-methylureido]-1-[4-methylaminocarbonyloxymethyl | 10.98 | C13H19N5O5 | NA |

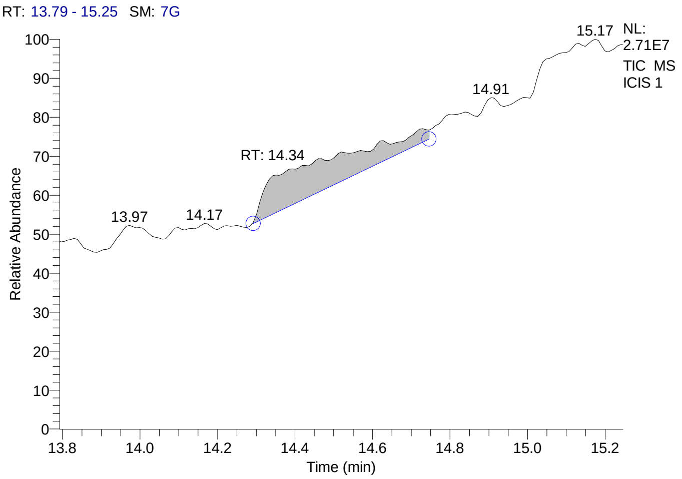


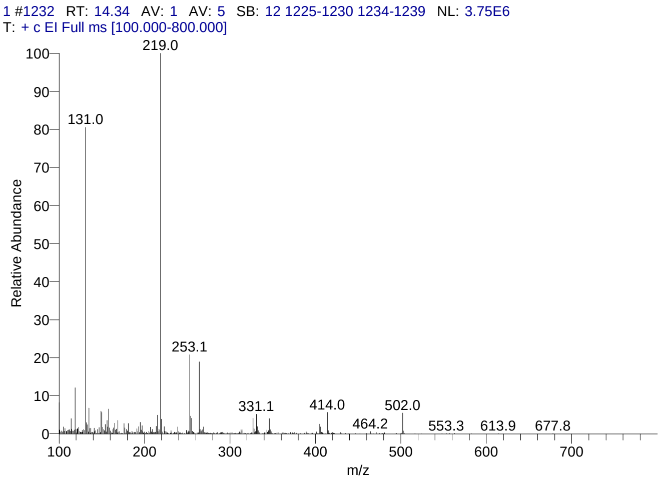


#### Library Search Results Table

| Compound Name | RT | Molecular Formula | Cas # |
|---|---|---|---|
| Perfluorotributylamine | 14.34 | C12F27N | 311-89-7 |
| Pyrimidin-2-one, 4-[N-methylureido]-1-[4-methylaminocarbonyloxymethyl | 14.34 | C13H19N5O5 | NA |
| 5á-Cholestane-3à,7à,12à,24,25,26-hexol hexa-TMS | 14.34 | C45H96O6Si6 | NA |

### CIL/ SAIF Panjab University Chandigarh

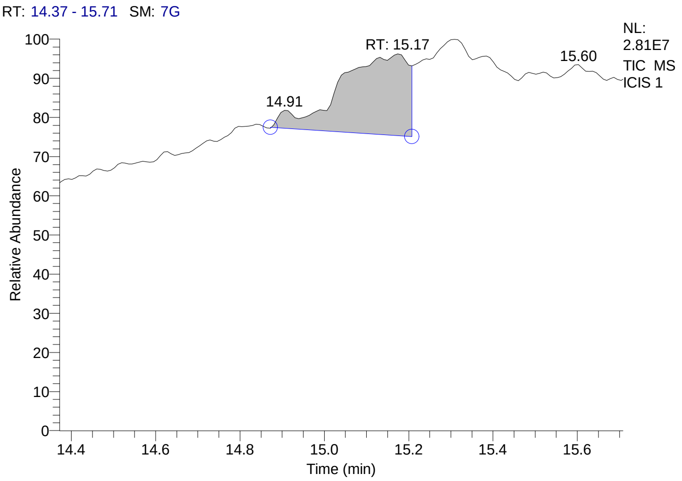


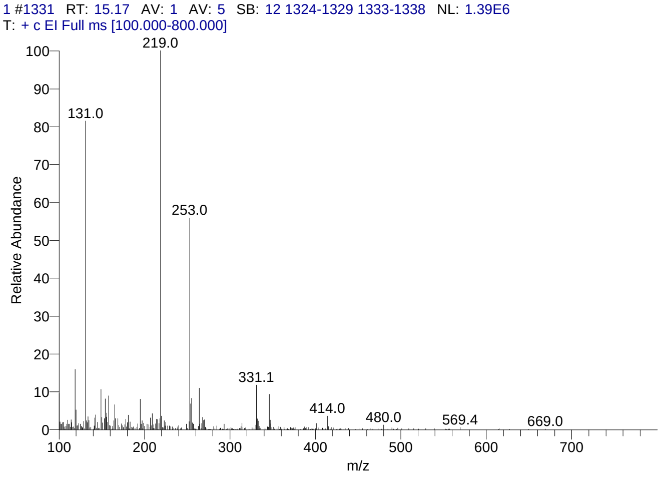


#### Library Search Results Table

| Compound Name | RT | Molecular Formula | Cas # |
|---|---|---|---|
| 5á-Cholestane-3à,7à,12à,24,25,26-hexol hexa-TMS | 15.17 | C45H96O6Si6 | NA |
| 4-Fluoro-2-nitroaniline, 5-[4-(pyrrolidin-1-yl)carbonylmethylpiperazin-1-yl]- | 15.17 | C16H22FN5O3 | NA |
| Cholestan-26-oic acid, 3,7,12,24-tetrakis(acetyloxy)-, methyl ester, (3à,5á,7à,12à)- | 15.17 | C36H56O10 | 60354-51-0 |

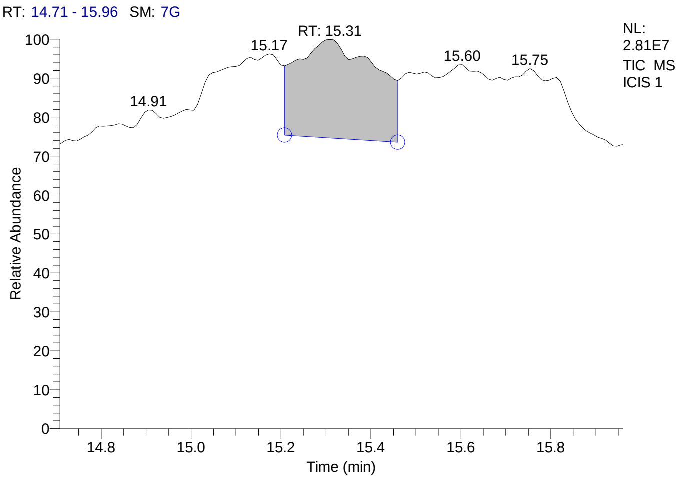


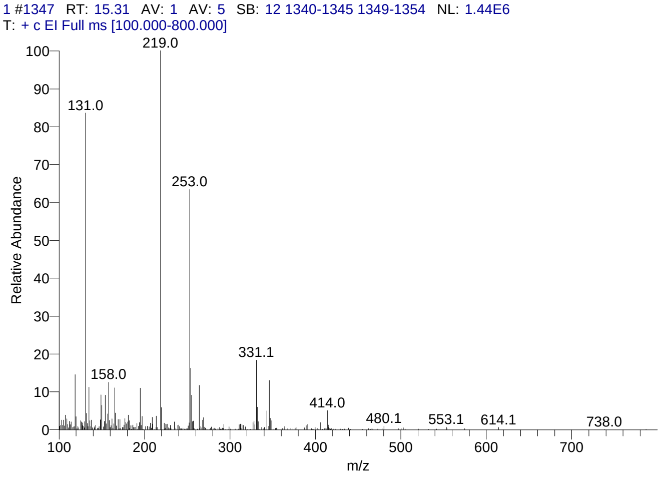


#### Library Search Results Table

| Compound Name | RT | Molecular Formula | Cas # |
|---|---|---|---|
| Pyrimidin-2-one, 4-[N-methylureido]-1-[4-methylaminocarbonyloxy methyl | 15.31 | C13H19N5O5 | NA |
| 4-Fluoro-2-nitroaniline, 5-[4-(pyrrolidin-1-yl)carbonylmethylpiperazin-1-yl]- | 15.31 | C16H22FN5O3 | NA |
| Cholestan-26-oic acid, 3,7,12,24-tetrakis(acetyloxy)-, methyl ester, (3à,5á,7à,12à)- | 15.31 | C36H56O10 | 60354-51-0 |

### CIL/ SAIF Panjab University Chandigarh

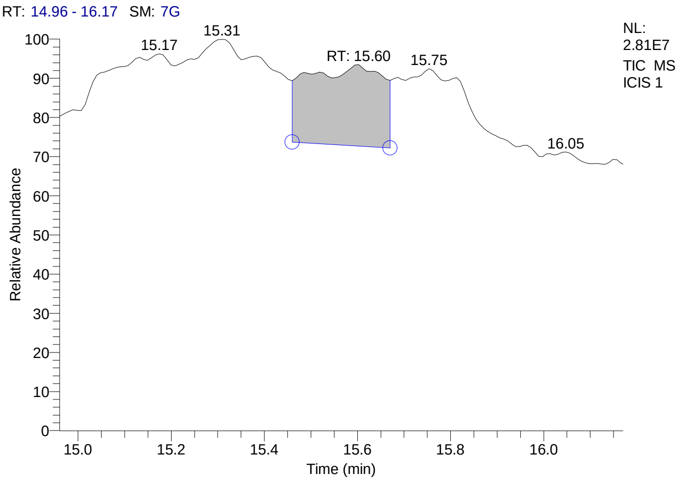


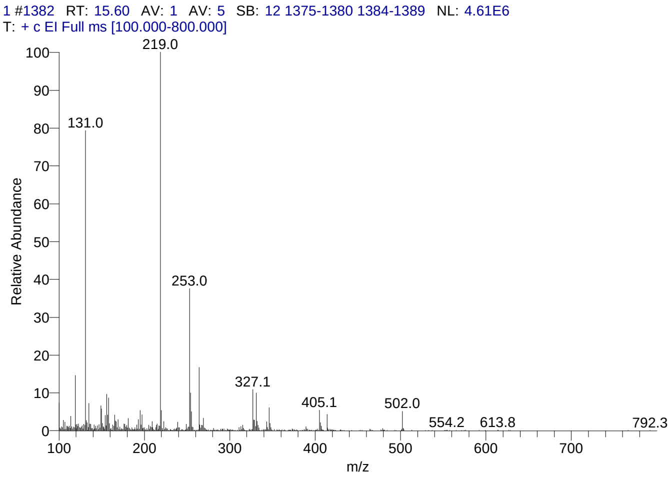


#### Library Search Results Table

| Compound Name | RT | Molecular Formula | Cas # |
|---|---|---|---|
| 5á-Cholestane-3à,7à,12à,24,25,26-hexol hexa-TMS | 15.60 | C45H96O6Si6 | NA |
| Cholestane-3,7,12,25-tetrol, tetraacetate, (3à,5á,7à,12à)- | 15.60 | C35H56O8 | 60354-50-9 |
| Cholestan-26-oic acid, 3,7,12,24-tetrakis(acetyloxy)-, methyl ester, (3à,5á,7à,12à)- | 15.60 | C36H56O10 | 60354-51-0 |

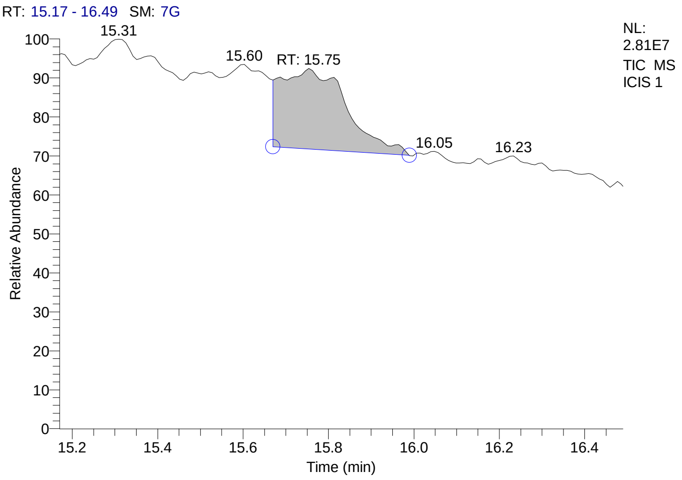


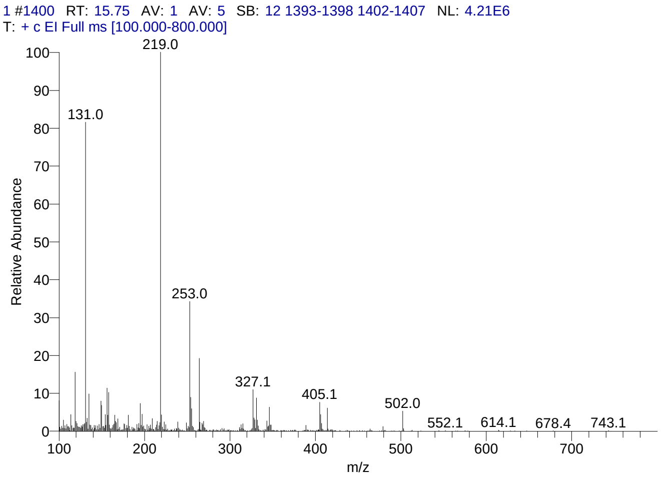


#### Library Search Results Table

| Compound Name | RT | Molecular Formula | Cas # |
|---|---|---|---|
| 5H-Cyclopropa[3,4]benz[1,2-e]azulen-5-one, 9,9a-bis(acetyloxy)-1,1a,1b,2,4a,7a,7b,8,9,9a-decahydro-2,4a,7b-trihydroxy-3-(hydroxymethyl)-1,1,6,8-tetramethyl-, [1aR-(1aà,1bá,2á,4aá,7aà,7bà,8à,9á,9aà)]- | 15.75 | C24H32O9 | 77573-19-4 |
| 5H-Cyclopropa[3,4]benz[1,2-e]azulen-5-one, 2,9,9a-tris(acetyloxy)-3-[(acetyloxy)methyl]-1,1a,1b,2,3,4,4a,7a,7b,8,9,9a-dodecahydro-3,4a,7b-trihydroxy-1,1,6,8-tetramethyl-, [1aR-(1aà,1bá,2à,3á,4aá,7aà,7bà,8à,9á,9aà)]- | 15.75 | C28H38O12 | 77646-81-2 |
| 3,19:5,6-Diepoxyandrostane, 17-acetoxy-4,4-dimethyl-3á-methoxy- | 15.75 | C24H36O5 | NA |

### CIL/ SAIF Panjab University Chandigarh

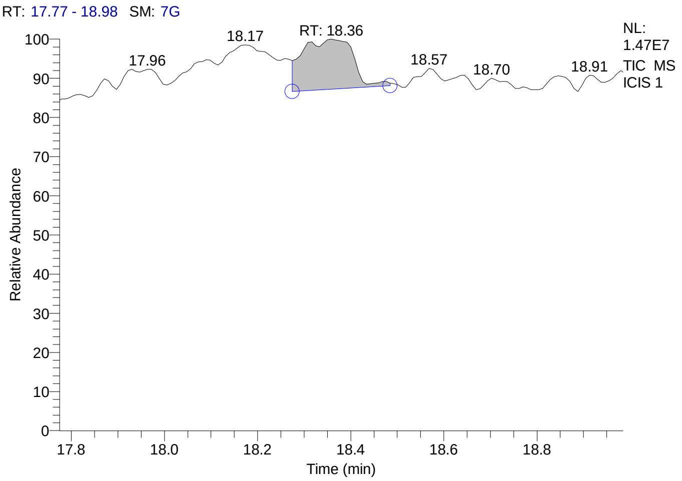


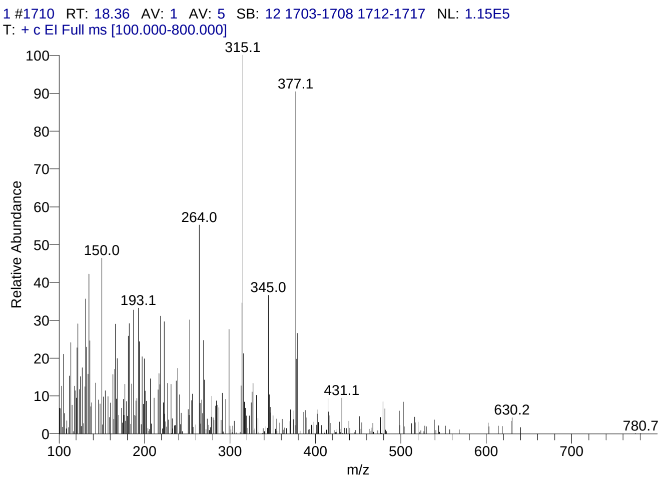


#### Library Search Results Table

| Compound Name | RT | Molecular Formula | Cas # |
|---|---|---|---|
| D-Homo-24-nor-17-oxachola-20,22-diene-3,16-dione, 7-(acetyloxy)-1,2:14,15:21,23-triepoxy-4,4,8-trimethyl-, (5à,7à,13à,14á,15á,17aà)- | 18.36 | C28H34O8 | 55658-66-7 |
| Flurandrenolide | 18.36 | C24H33FO6 | 1524-88-5 |
| Dexamethasone-21-acetate | 18.36 | C24H31FO6 | 1177-87-3 |

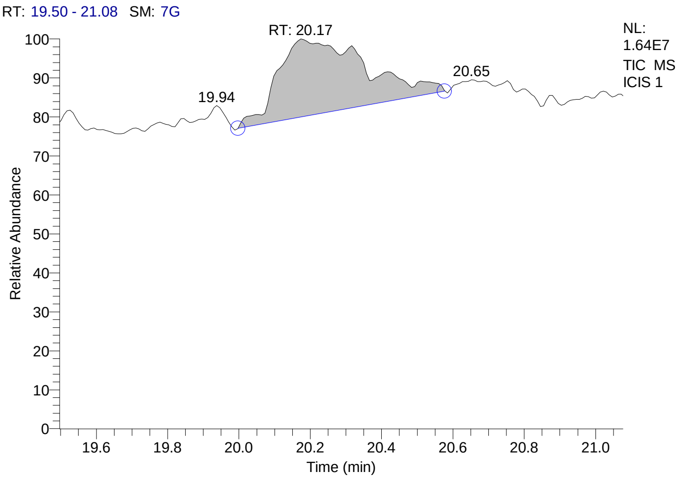


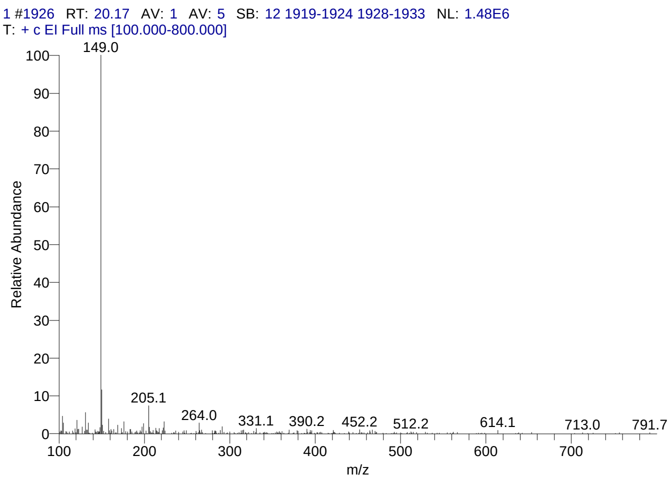


#### Library Search Results Table

| Compound Name | RT | Molecular Formula | Cas # |
|---|---|---|---|
| Phthalic acid, butyl hex-3-yl ester | 20.17 | C18H26O4 | NA |
| Phthalic acid, butyl dodecyl ester | 20.17 | C24H38O4 | NA |
| Phthalic acid, butyl 4-octyl ester | 20.17 | C20H30O4 | NA |

### CIL/ SAIF Panjab University Chandigarh

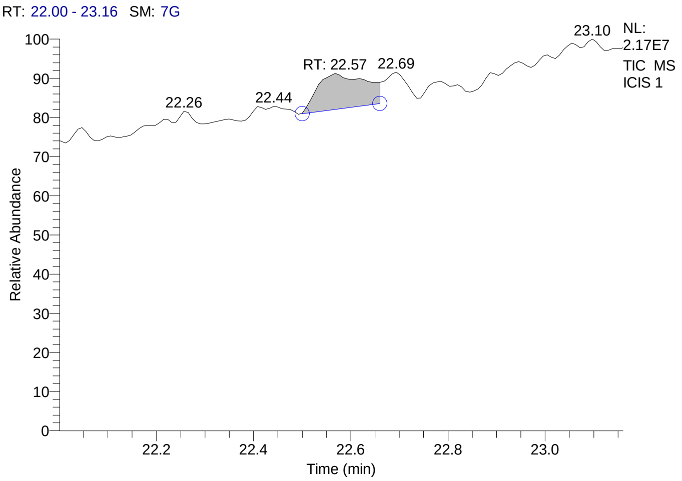


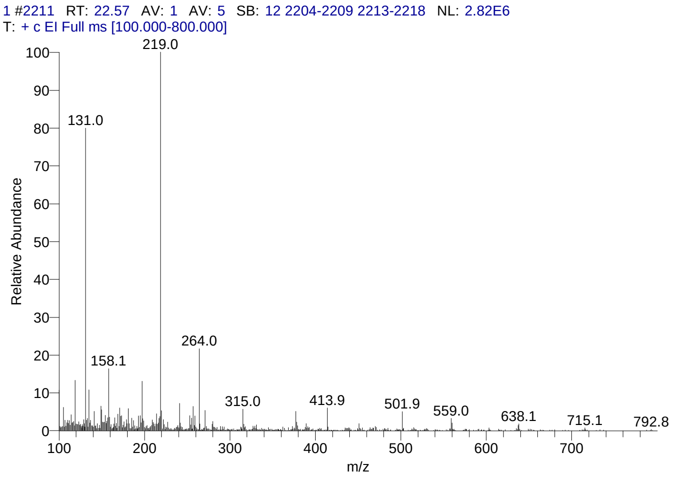


#### Library Search Results Table

| Compound Name | RT | Molecular Formula | Cas # |
|---|---|---|---|
| Olean-12-ene-3,16,21,22,23,28-hexol, (3á,4à,16à,21á,22à)- | 22.57 | C30H50O6 | 13844-22-9 |
| 5á-Cholestane-3à,7à,12à,24,25,26-hexol hexa-TMS | 22.57 | C45H96O6Si6 | NA |
| Olean-12-ene-3,16,21,22,23,28-hexol, (3á,4á,16à,21á,22à)- | 22.57 | C30H50O6 | 20853-07-0 |

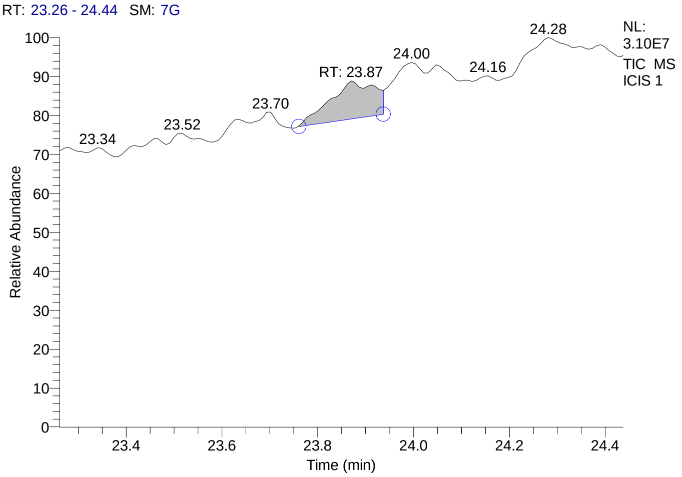


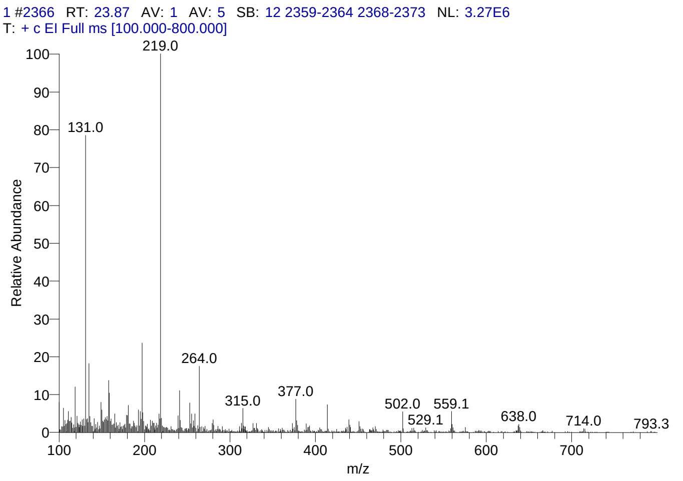


#### Library Search Results Table

| Compound Name | RT | Molecular Formula | Cas # |
|---|---|---|---|
| Olean-12-ene-3,16,21,22,23,28-hexol, (3á,4à,16à,21á,22à)- | 23.87 | C30H50O6 | 13844-22-9 |
| Withaferin A | 23.87 | C28H38O6 | 5119-48-2 |
| Lycophyll | 23.87 | C40H56O2 | 19891-75-9 |

### CIL/ SAIF Panjab University Chandigarh

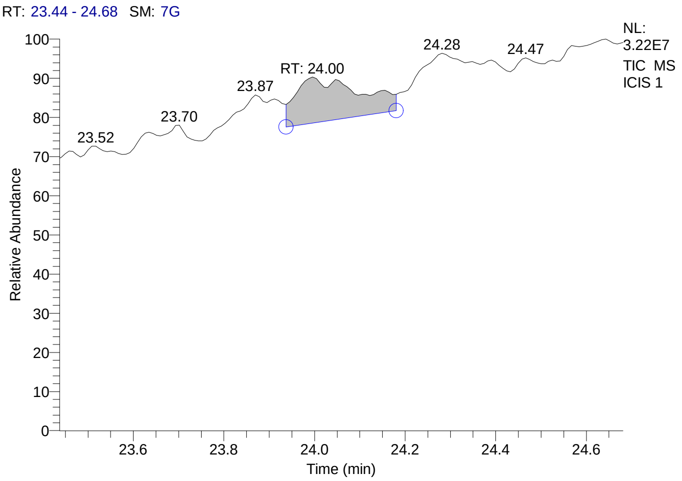


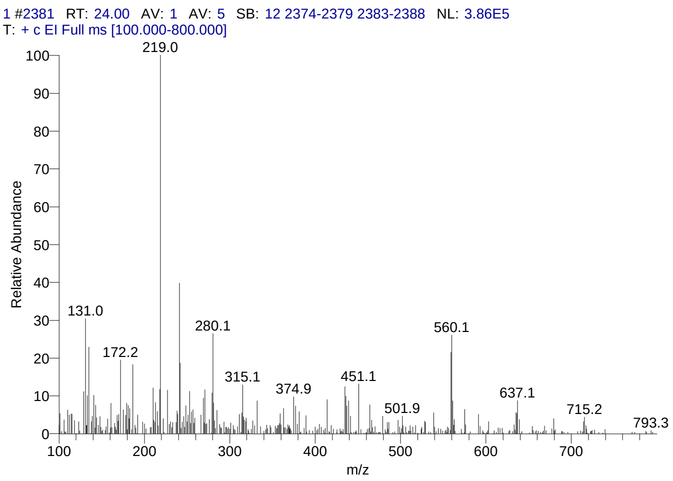


#### Library Search Results Table

| Compound Name | RT | Molecular Formula | Cas # |
|---|---|---|---|
| Rhodoxanthin | 24.00 | C40H50O2 | 116-30-3 |
| Lanostane-7,11-dione, 3,18-bis(acetyloxy)-, cyclic 7-(1,2-ethanediyl mercaptole), (3á,20.xi.)- | 24.00 | C36H58O5S2 | 56298-07-8 |
| Hematoporphyrin | 24.00 | C34H38N4O6 | 14459-29-1 |

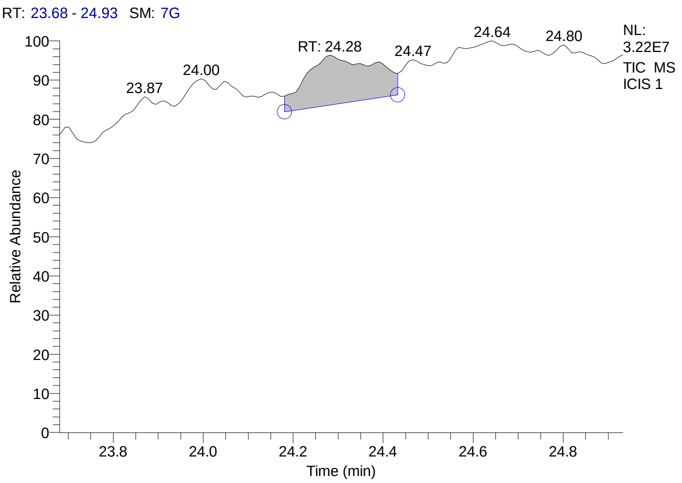


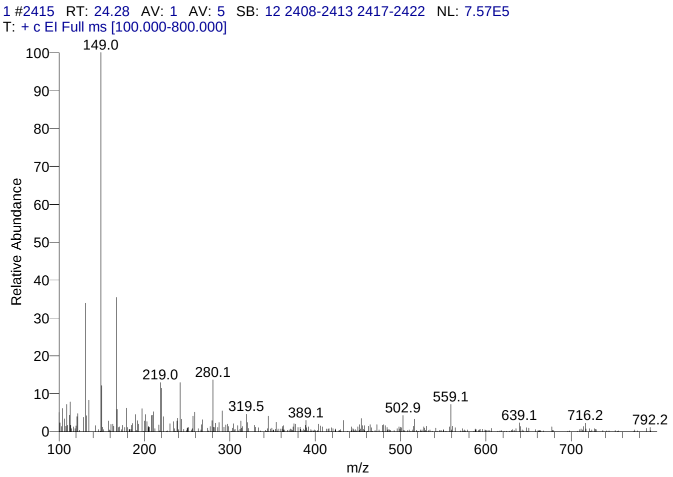


#### Library Search Results Table

| Compound Name | RT | Molecular Formula | Cas # |
|---|---|---|---|
| Phthalic acid, 2-hexyl ester | 24.28 | C14H18O4 | 79107-80-5 |
| Phthalic acid, 3,3-dimethylbut-2-yl pentyl ester | 24.28 | C19H28O4 | NA |
| 2-(Isobutoxycarbonyl)benzoic acid | 24.28 | C12H14O4 | NA |

### CIL/ SAIF Panjab University Chandigarh

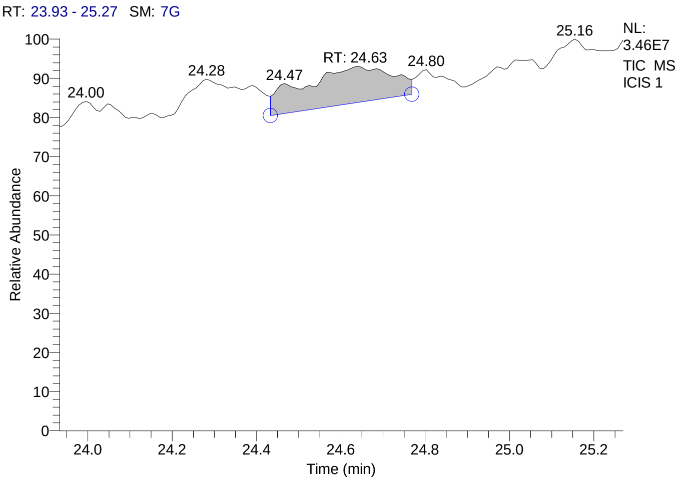


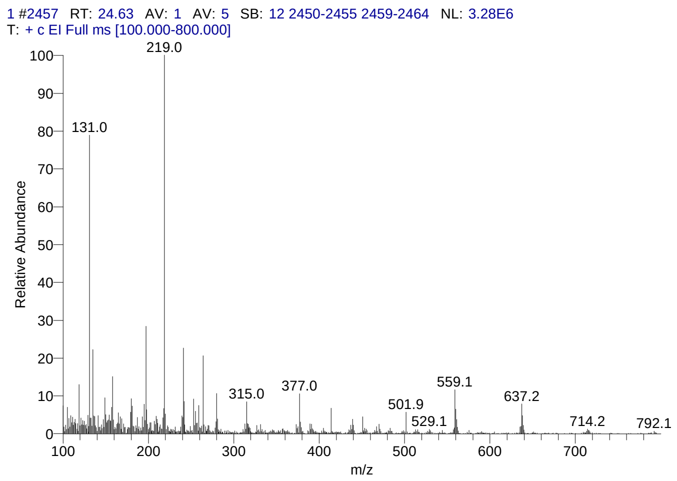


#### Library Search Results Table

| Compound Name | RT | Molecular Formula | Cas # |
|---|---|---|---|
| Rhodoxanthin | 24.63 | C40H50O2 | 116-30-3 |
| Olean-12-ene-3,16,21,22,23,28-hexol, (3á,4à,16à,21á,22à)- | 24.63 | C30H50O6 | 13844-22-9 |
| 4aà,4bá-Gibbane-1à,10á-dicarboxylic acid, 4a-formyl-7-hydroxy-1-methyl-8-methylene-, dimethyl ester | 24.63 | C22H30O6 | 6980-45-6 |

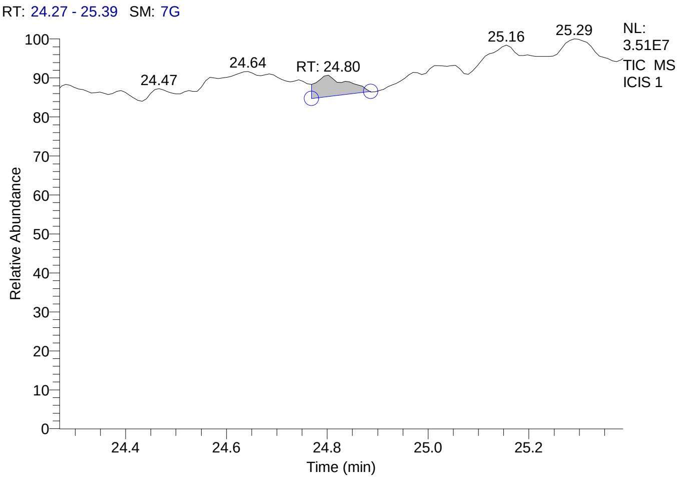


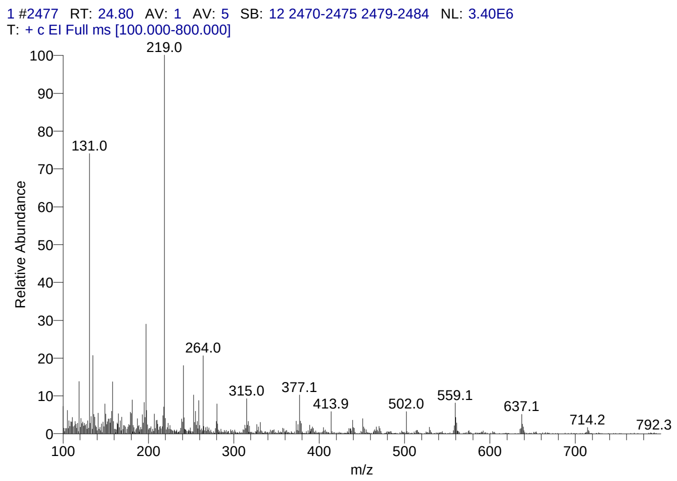


#### Library Search Results Table

| Compound Name | RT | Molecular Formula | Cas # |
|---|---|---|---|
| Rhodoxanthin | 24.80 | C40H50O2 | 116-30-3 |
| Olean-12-ene-3,16,21,22,23,28-hexol, (3á,4à,16à,21á,22à)- | 24.80 | C30H50O6 | 13844-22-9 |
| 4aà,4bá-Gibbane-1à,10á-dicarboxylic acid, 4a-formyl-7-hydroxy-1-methyl-8-methylene-, dimethyl ester | 24.80 | C22H30O6 | 6980-45-6 |

### CIL/ SAIF Panjab University Chandigarh

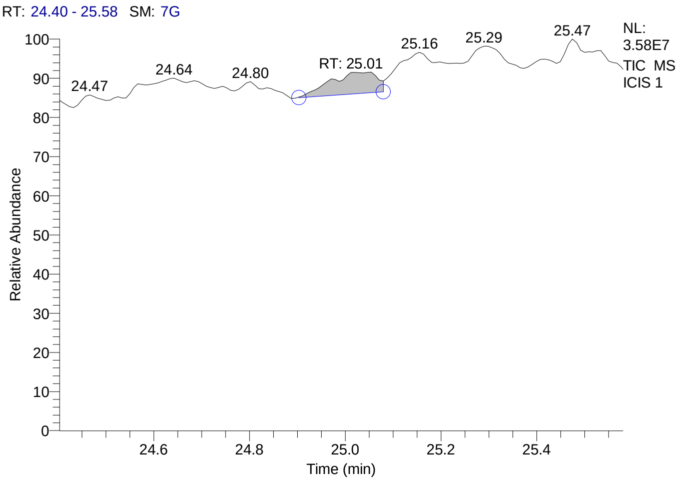


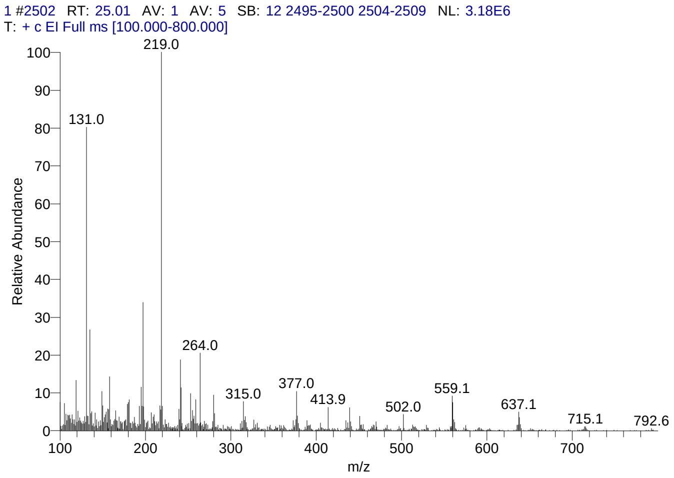


#### Library Search Results Table

| Compound Name | RT | Molecular Formula | Cas # |
|---|---|---|---|
| Rhodoxanthin | 25.01 | C40H50O2 | 116-30-3 |
| Olean-12-ene-3,16,21,22,23,28-hexol, (3á,4à,16à,21á,22à)- | 25.01 | C30H50O6 | 13844-22-9 |
| 4aà,4bá-Gibbane-1à,10á-dicarboxylic acid, 4a-formyl-7-hydroxy-1-methyl-8-methylene-, dimethyl ester | 25.01 | C22H30O6 | 6980-45-6 |

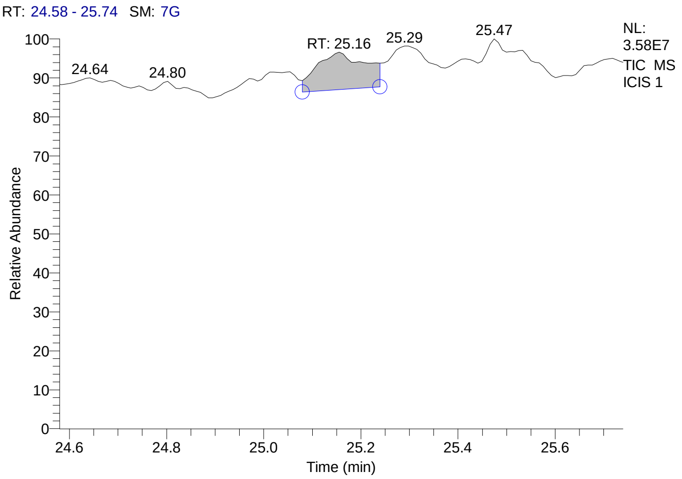


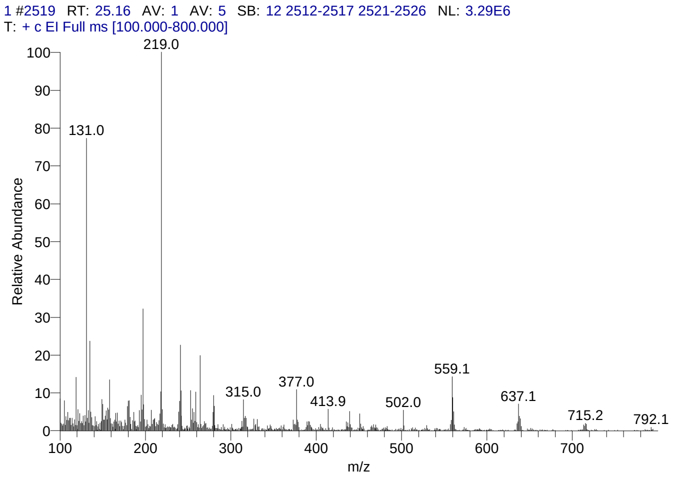


#### Library Search Results Table

| Compound Name | RT | Molecular Formula | Cas # |
|---|---|---|---|
| Rhodoxanthin | 25.16 | C40H50O2 | 116-30-3 |
| Olean-12-ene-3,16,21,22,23,28-hexol, (3á,4à,16à,21á,22à)- | 25.16 | C30H50O6 | 13844-22-9 |
| 4aà,4bá-Gibbane-1à,10á-dicarboxylic acid, 4a-formyl-7-hydroxy-1-methyl-8-methylene-, dimethyl ester | 25.16 | C22H30O6 | 6980-45-6 |

### CIL/ SAIF Panjab University Chandigarh

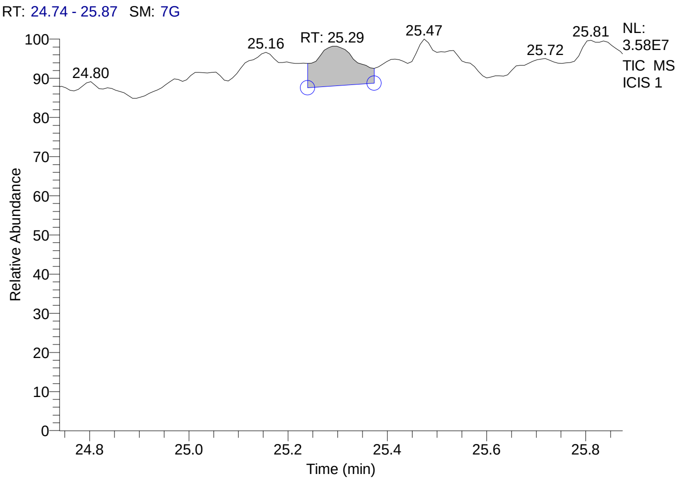


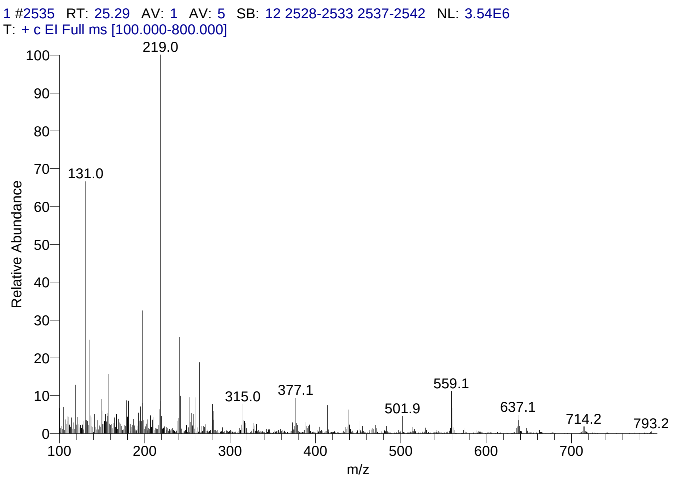


#### Library Search Results Table

| Compound Name | RT | Molecular Formula | Cas # |
|---|---|---|---|
| Rhodoxanthin | 25.29 | C40H50O2 | 116-30-3 |
| Olean-12-ene-3,16,21,22,23,28-hexol, (3á,4à,16à,21á,22à)- | 25.29 | C30H50O6 | 13844-22-9 |
| 4aà,4bá-Gibbane-1à,10á-dicarboxylic acid, 4a-formyl-7-hydroxy-1-methyl-8-methylene-, dimethyl ester | 25.29 | C22H30O6 | 6980-45-6 |

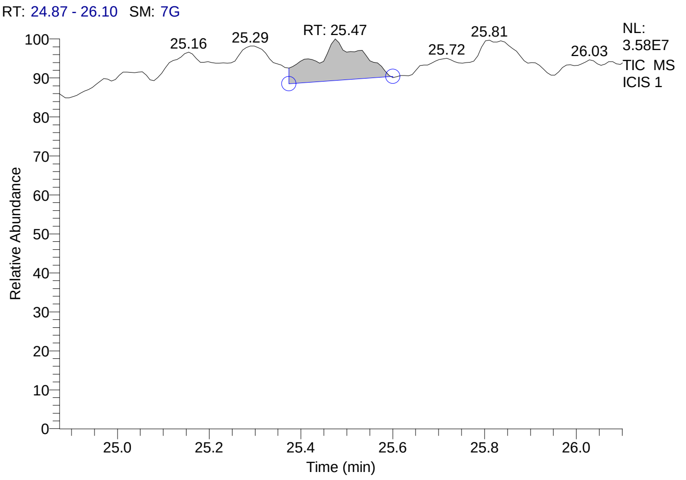


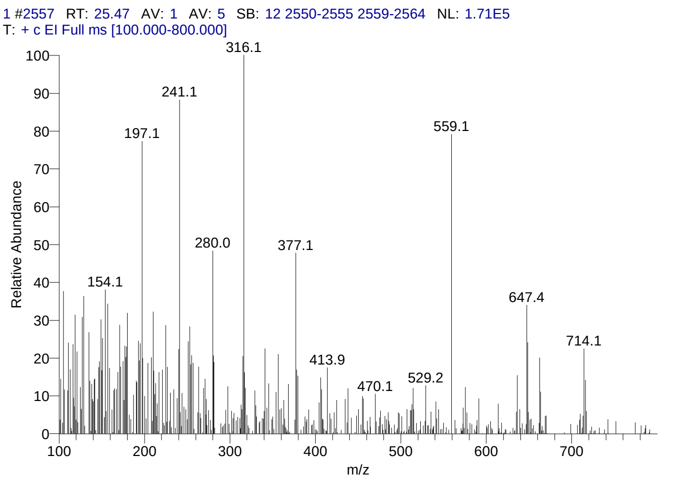


#### Library Search Results Table

| Compound Name | RT | Molecular Formula | Cas # |
|---|---|---|---|
| Lanostane-7,11-dione, 3,18-bis(acetyloxy)-, cyclic 7-(1,2-ethanediyl mercaptole), (3á,20.xi.)- | 25.47 | C36H58O5S2 | 56298-07-8 |
| Pregn-4-ene-3,11,20-trione, 6,17,21-tris[(trimethylsilyl)oxy]-, 3,20-bis(O-methyloxime), (6á)- | 25.47 | C32H58N2O6Si3 | 57326-06-4 |
| [(1H)-Pyrrole-3-propanoic acid, 2-ethoxycarbonyl-4-ethoxycarbonylmethyl]-5,5'-met hylene, bis-, diethyl ester | 25.47 | C33H46N2O12 | 116377-74-3 |

### CIL/ SAIF Panjab University Chandigarh

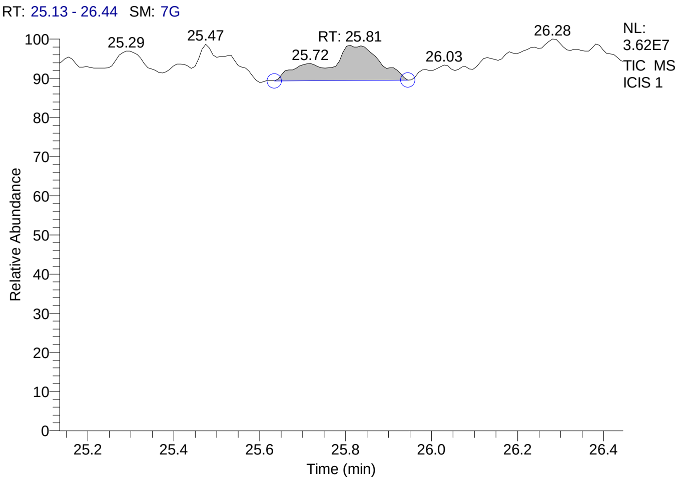


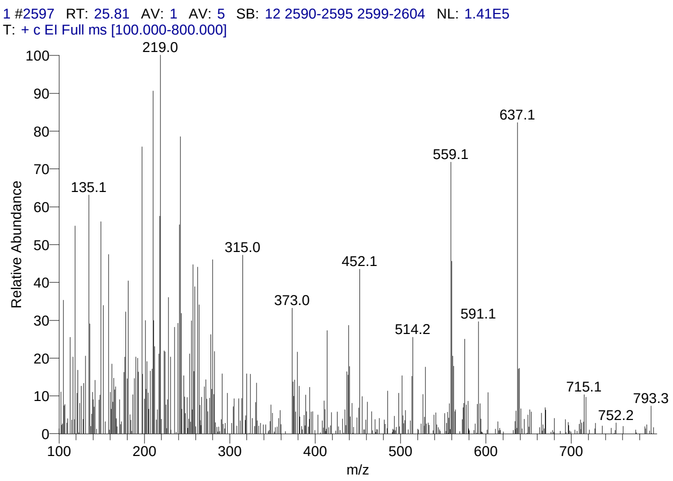


#### Library Search Results Table

| Compound Name | RT | Molecular Formula | Cas # |
|---|---|---|---|
| Rhodoxanthin | 25.81 | C40H50O2 | 116-30-3 |
| Lanostane-7,11-dione, 3,18-bis(acetyloxy)-, cyclic 7-(1,2-ethanediyl mercaptole), (3á,20.xi.)- | 25.81 | C36H58O5S2 | 56298-07-8 |
| Pregn-4-en-18-al, 3,20-bis(methoxyimino)-11,21-bis[(trimethylsilyl)oxy]-, O-methyloxime, (11á,17à)- | 25.81 | C30H53N3O5Si2 | 69854-81-5 |

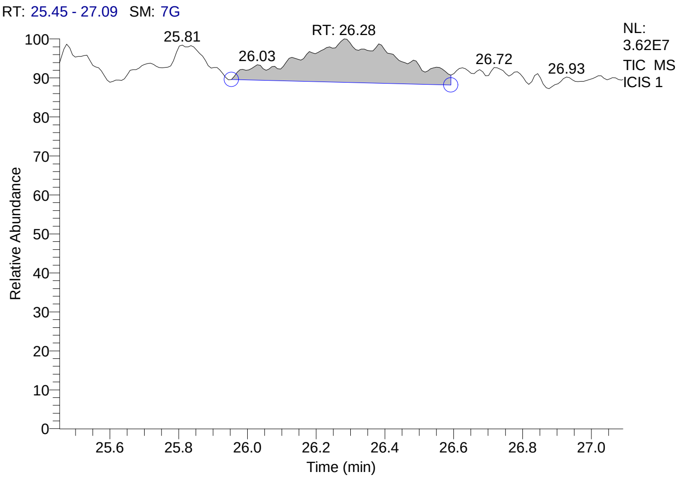


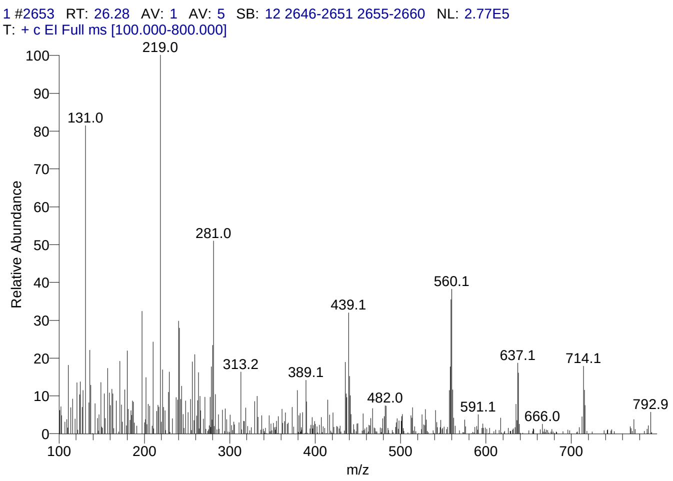


#### Library Search Results Table

| Compound Name | RT | Molecular Formula | Cas # |
|---|---|---|---|
| Octaphenylcyclotetrasiloxane | 26.28 | C48H40O4Si4 | 546-56-5 |
| Rhodoxanthin | 26.28 | C40H50O2 | 116-30-3 |
| Lanostane-7,11-dione, 3,18-bis(acetyloxy)-, cyclic 7-(1,2-ethanediyl mercaptole), (3á,20.xi.)- | 26.28 | C36H58O5S2 | 56298-07-8 |

### CIL/ SAIF Panjab University Chandigarh

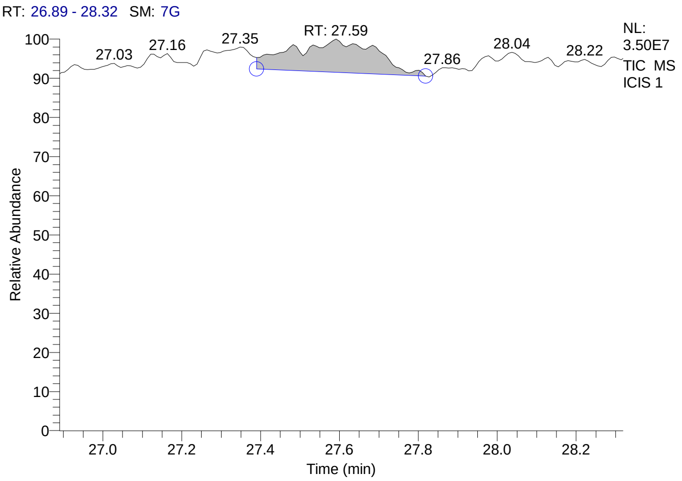


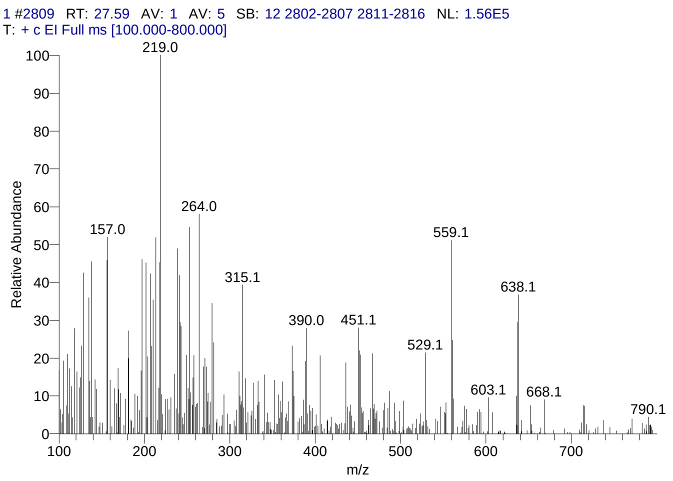


#### Library Search Results Table

| Compound Name | RT | Molecular Formula | Cas # |
|---|---|---|---|
| Docosanoic acid, 1,2,3-propanetriyl ester | 27.59 | C69H134O6 | 18641-57-1 |
| 19-Norpregna-1,3,5,7,9-pentaen-21-al, 3,17-bis[(trimethylsilyl)oxy]-, O-methyloxime, (17à)- | 27.59 | C27H41NO3Si2 | 74299-04-0 |
| Pregn-4-ene-3,20-dione, 11,17,21-tris[(trimethylsilyl)oxy]-, bis(O-methyloxime), (11á)- | 27.59 | C32H60N2O5Si3 | 32221-26-4 |

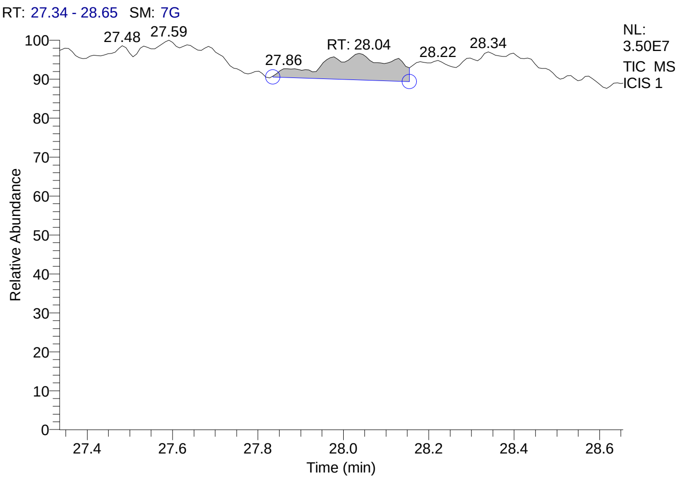


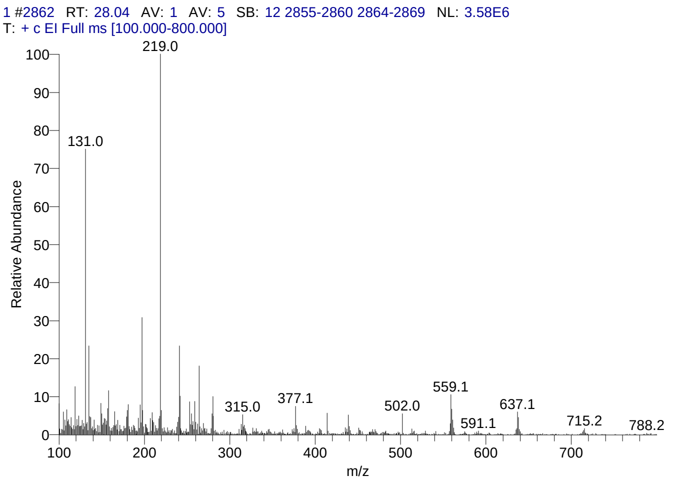


#### Library Search Results Table

| Compound Name | RT | Molecular Formula | Cas # |
|---|---|---|---|
| Rhodoxanthin | 28.04 | C40H50O2 | 116-30-3 |
| Olean-12-ene-3,16,21,22,23,28-hexol, (3á,4à,16à,21á,22à)- | 28.04 | C30H50O6 | 13844-22-9 |
| 4aà,4bá-Gibbane-1à,10á-dicarboxylic acid, 4a-formyl-7-hydroxy-1-methyl-8-methylene-, dimethyl ester | 28.04 | C22H30O6 | 6980-45-6 |

### CIL/ SAIF Panjab University Chandigarh

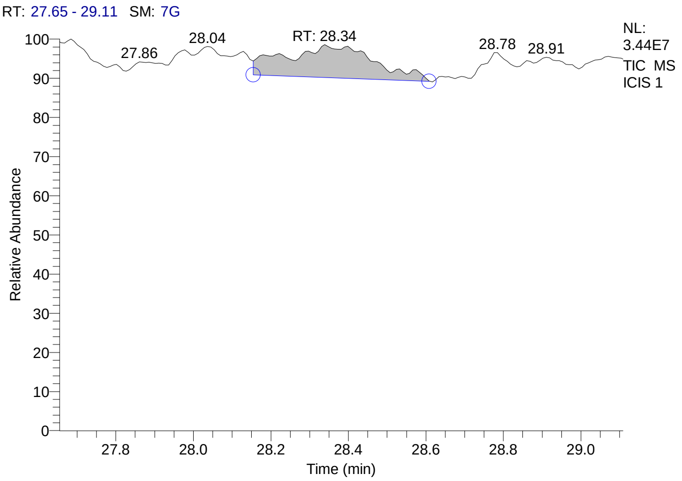


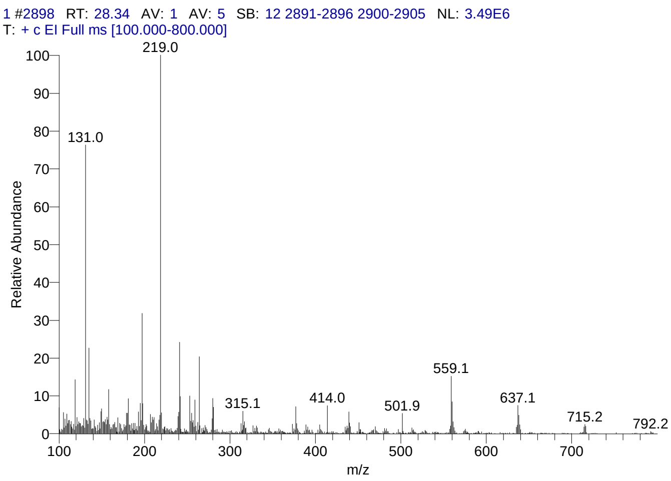


#### Library Search Results Table

| Compound Name | RT | Molecular Formula | Cas # |
|---|---|---|---|
| Olean-12-ene-3,16,21,22,23,28-hexol, (3á,4à,16à,21á,22à)- | 28.34 | C30H50O6 | 13844-22-9 |
| 4aà,4bá-Gibbane-1à,10á-dicarboxylic acid, 4a-formyl-7-hydroxy-1-methyl-8-methylene-, dimethyl ester | 28.34 | C22H30O6 | 6980-45-6 |
| Olean-12-ene-3,16,21,22,23,28-hexol, (3á,4á,16à,21á,22à)- | 28.34 | C30H50O6 | 20853-07-0 |

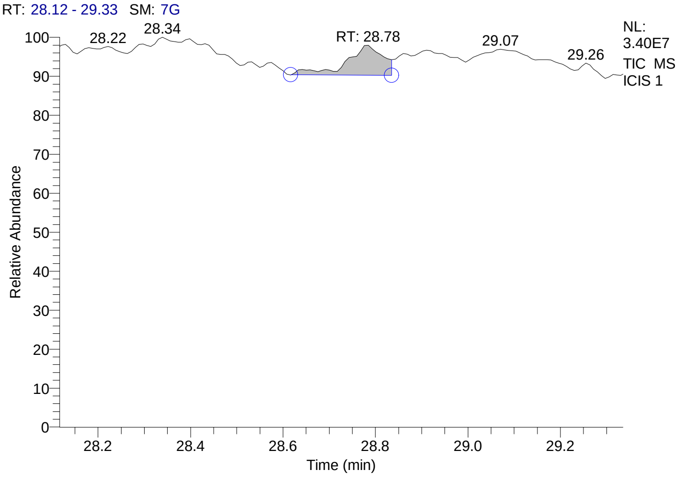


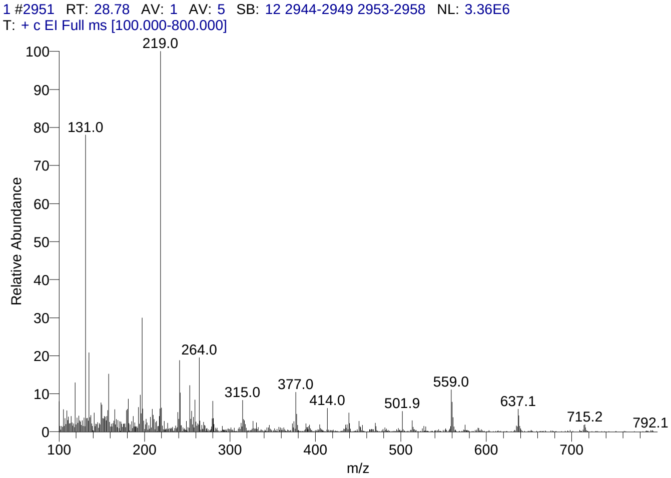


#### Library Search Results Table

| Compound Name | RT | Molecular Formula | Cas # |
|---|---|---|---|
| Rhodoxanthin | 28.78 | C40H50O2 | 116-30-3 |
| Olean-12-ene-3,16,21,22,23,28-hexol, (3á,4à,16à,21á,22à)- | 28.78 | C30H50O6 | 13844-22-9 |
| 4aà,4bá-Gibbane-1à,10á-dicarboxylic acid, 4a-formyl-7-hydroxy-1-methyl-8-methylene-, dimethyl ester | 28.78 | C22H30O6 | 6980-45-6 |

### CIL/ SAIF Panjab University Chandigarh

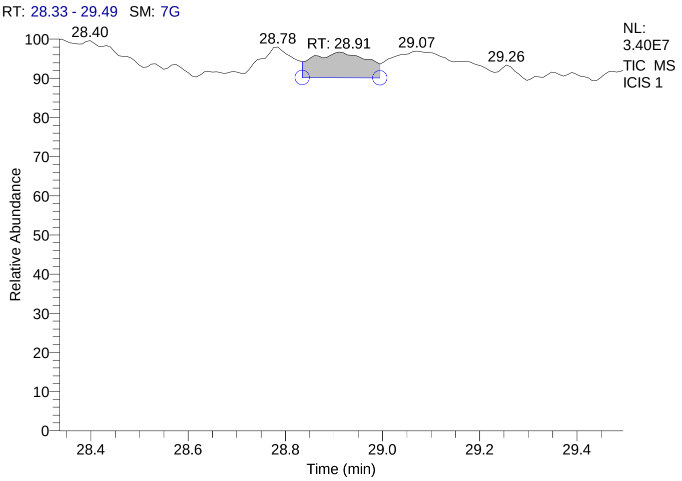


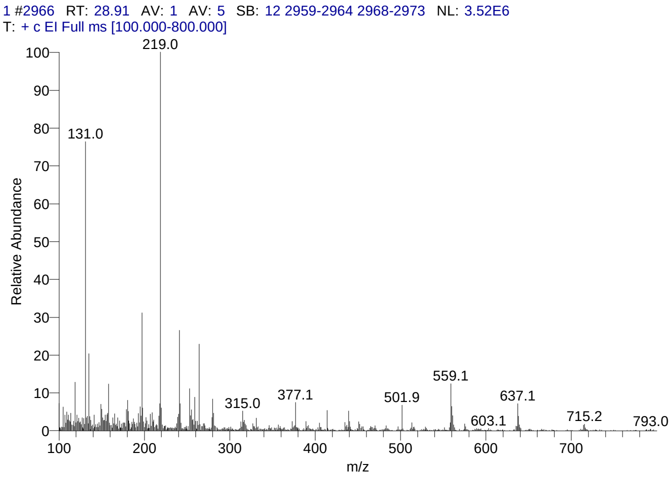


#### Library Search Results Table

| Compound Name | RT | Molecular Formula | Cas # |
|---|---|---|---|
| Rhodoxanthin | 28.91 | C40H50O2 | 116-30-3 |
| 4aà,4bá-Gibbane-1à,10á-dicarboxylic acid, 4a-formyl-7-hydroxy-1-methyl-8-methylene-, dimethyl ester | 28.91 | C22H30O6 | 6980-45-6 |
| Olean-12-ene-3,16,21,22,23,28-hexol, (3á,4à,16à,21á,22à)- | 28.91 | C30H50O6 | 13844-22-9 |

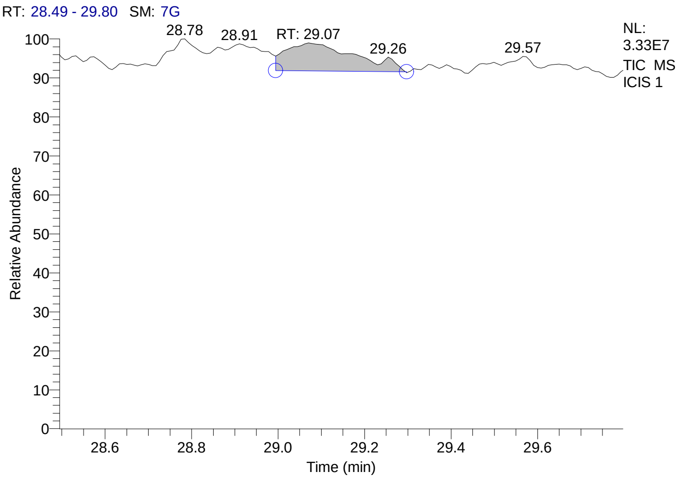


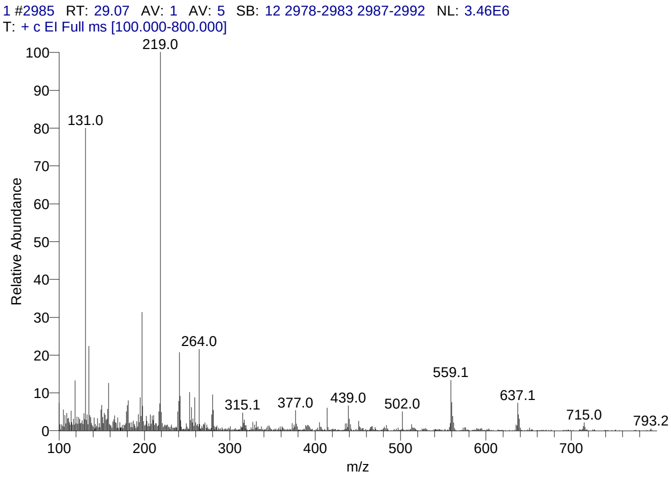


#### Library Search Results Table

| Compound Name | RT | Molecular Formula | Cas # |
|---|---|---|---|
| Olean-12-ene-3,16,21,22,23,28-hexol, (3á,4à,16à,21á,22à)- | 29.07 | C30H50O6 | 13844-22-9 |
| 4aà,4bá-Gibbane-1à,10á-dicarboxylic acid, 4a-formyl-7-hydroxy-1-methyl-8-methylene-, dimethyl ester | 29.07 | C22H30O6 | 6980-45-6 |
| Olean-12-ene-3,16,21,22,23,28-hexol, (3á,4á,16à,21á,22à)- | 29.07 | C30H50O6 | 20853-07-0 |

### CIL/ SAIF Panjab University Chandigarh

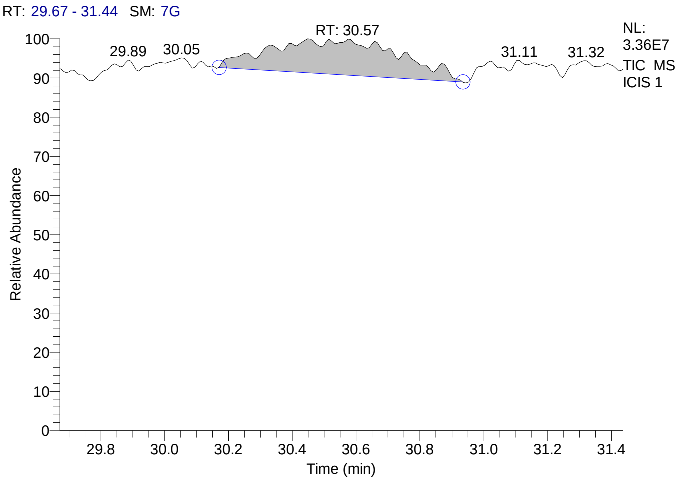


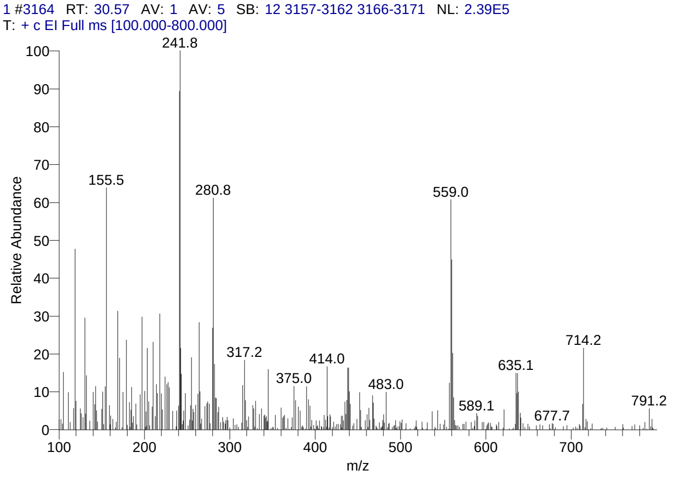


#### Library Search Results Table

| Compound Name | RT | Molecular Formula | Cas # |
|---|---|---|---|
| Octaphenylcyclotetrasiloxane | 30.57 | $C_{48}H_{40}O_4Si_4$ | 546-56-5 |
| Rhodoxanthin | 30.57 | $C_{40}H_{50}O_2$ | 116-30-3 |
| Thiophene, 2,3,5-tris(diphenylphosphino)- | 30.57 | $C_{40}H_{31}P_3S$ | NA |

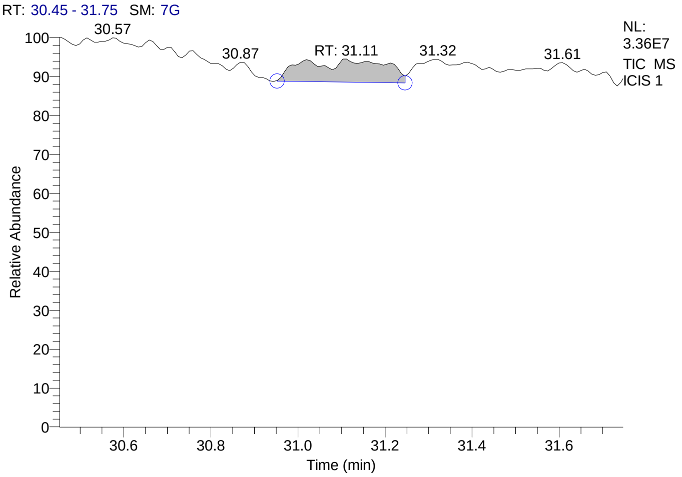


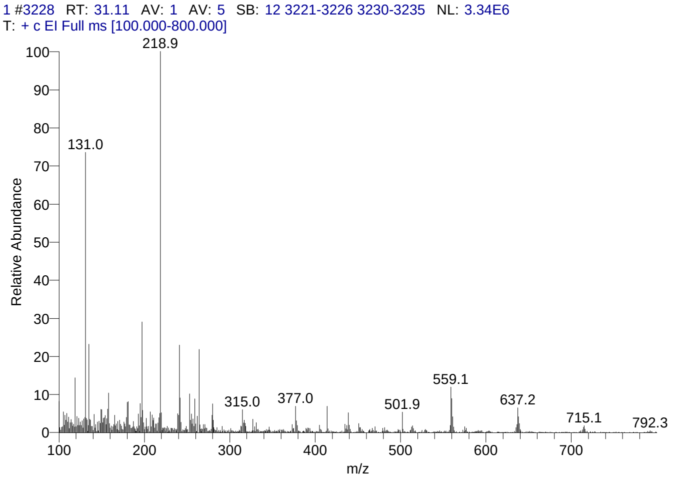


#### Library Search Results Table

| Compound Name | RT | Molecular Formula | Cas # |
|---|---|---|---|
| Rhodoxanthin | 31.11 | $C_{40}H_{50}O_2$ | 116-30-3 |
| Olean-12-ene-3,16,21,22,23,28-hexol, (3á,4à,16à,21á,22à)- | 31.11 | $C_{30}H_{50}O_6$ | 13844-22-9 |
| 4aà,4bá-Gibbane-1à,10á-dicarboxylic acid, 4a-formyl-7-hydroxy-1-methyl-8-methylene-, dimethyl ester | 31.11 | $C_{22}H_{30}O_6$ | 6980-45-6 |

### CIL/ SAIF Panjab University Chandigarh

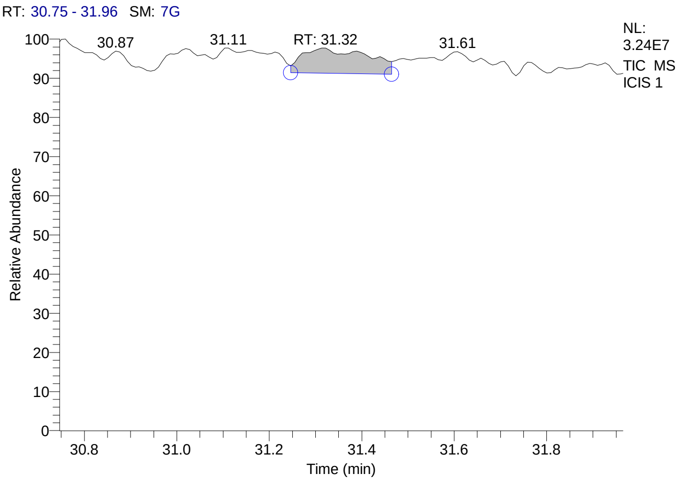


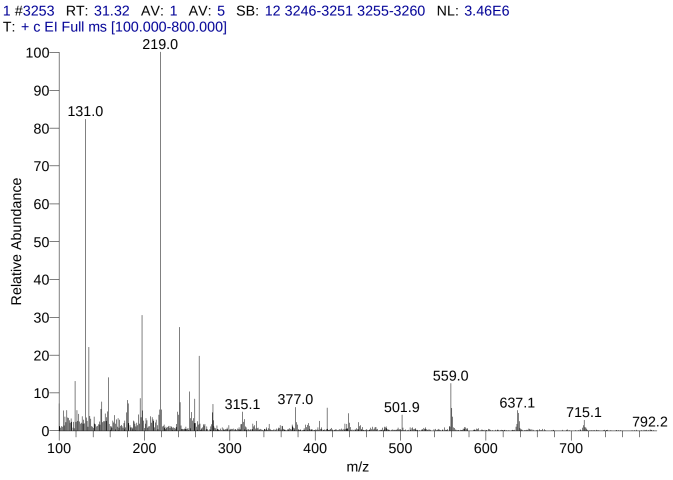


#### Library Search Results Table

| Compound Name | RT | Molecular Formula | Cas # |
|---|---|---|---|
| Rhodoxanthin | 31.32 | C40H50O2 | 116-30-3 |
| 4aà,4bá-Gibbane-1à,10á-dicarboxylic acid, 4a-formyl-7-hydroxy-1-methyl-8-methylene-, dimethyl ester | 31.32 | C22H30O6 | 6980-45-6 |
| Olean-12-ene-3,16,21,22,23,28-hexol, (3á,4à,16à,21á,22à)- | 31.32 | C30H50O6 | 13844-22-9 |

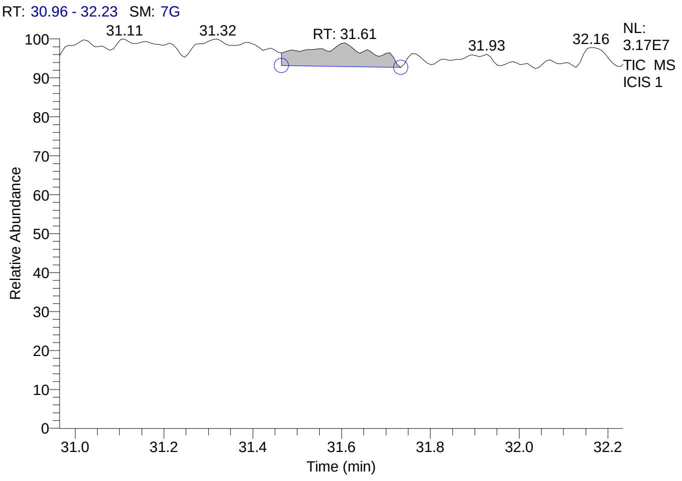


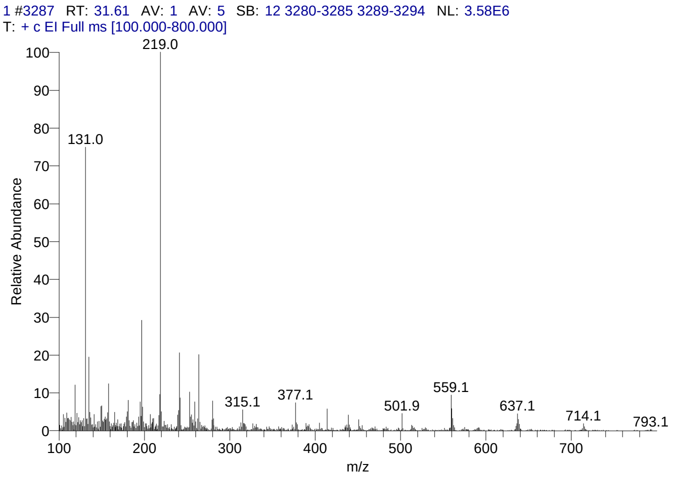


#### Library Search Results Table

| Compound Name | RT | Molecular Formula | Cas # |
|---|---|---|---|
| Rhodoxanthin | 31.61 | C40H50O2 | 116-30-3 |
| 4aà,4bá-Gibbane-1à,10á-dicarboxylic acid, 4a-formyl-7-hydroxy-1-methyl-8-methylene-, dimethyl ester | 31.61 | C22H30O6 | 6980-45-6 |
| Olean-12-ene-3,16,21,22,23,28-hexol, (3á,4à,16à,21á,22à)- | 31.61 | C30H50O6 | 13844-22-9 |

### CIL/ SAIF Panjab University Chandigarh

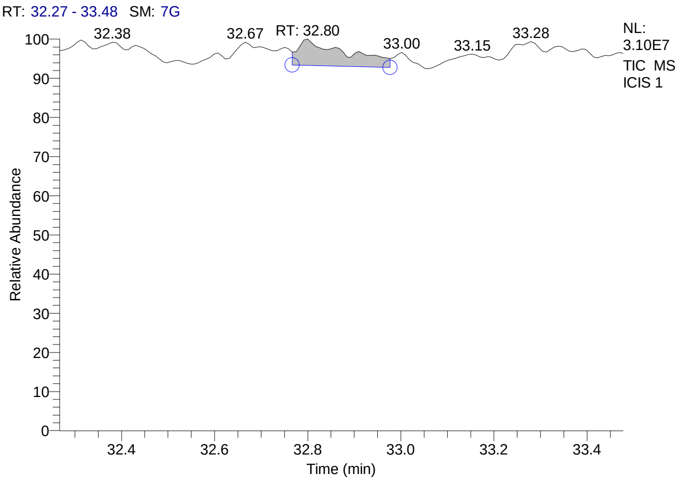


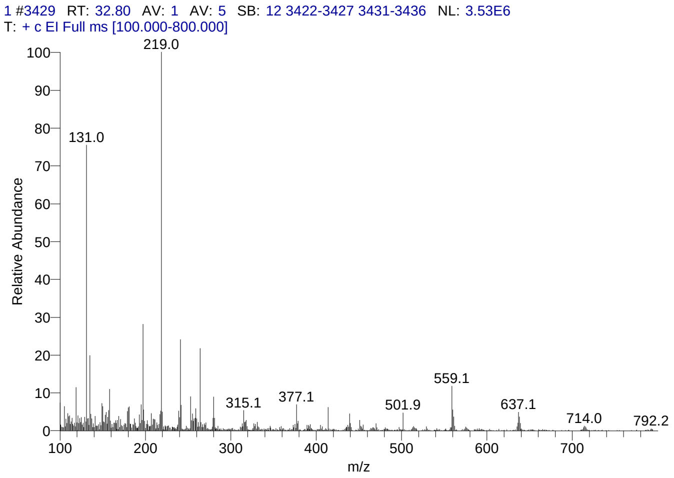


#### Library Search Results Table

| Compound Name | RT | Molecular Formula | Cas # |
|---|---|---|---|
| Rhodoxanthin | 32.80 | C40H50O2 | 116-30-3 |
| 4aà,4bá-Gibbane-1à,10á-dicarboxylic acid, 4a-formyl-7-hydroxy-1-methyl-8-methylene-, dimethyl ester | 32.80 | C22H30O6 | 6980-45-6 |
| Olean-12-ene-3,16,21,22,23,28-hexol, (3á,4à,16à,21á,22à)- | 32.80 | C30H50O6 | 13844-22-9 |

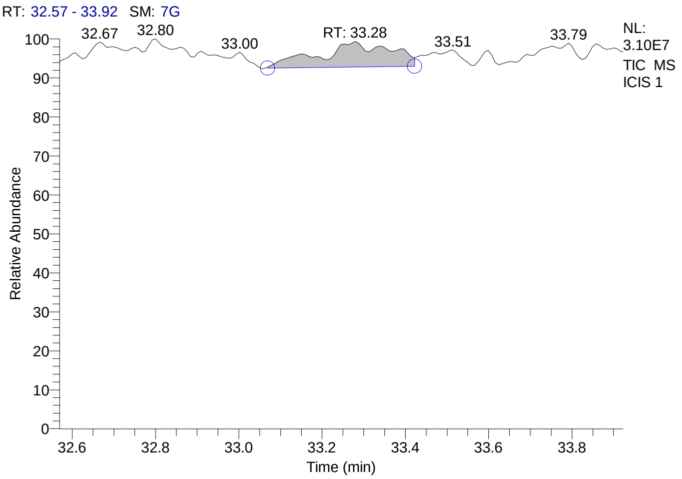


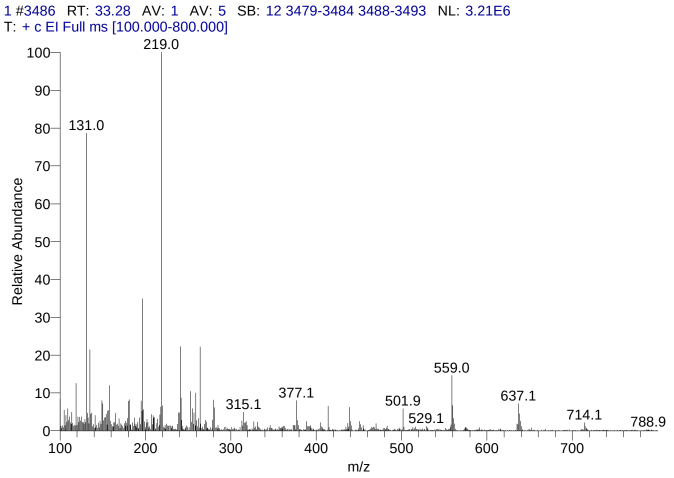


#### Library Search Results Table

| Compound Name | RT | Molecular Formula | Cas # |
|---|---|---|---|
| Olean-12-ene-3,16,21,22,23,28-hexol, (3á,4à,16à,21á,22à)- | 33.28 | C30H50O6 | 13844-22-9 |
| Olean-12-ene-3,16,21,22,28-pentol, 21-(2-methyl-2-butenoate), [3á,16à,21á(Z),22à]- | 33.28 | C35H56O6 | 20089-98-9 |
| 4aà,4bá-Gibbane-1à,10á-dicarboxylic acid, 4a-formyl-7-hydroxy-1-methyl-8-methylene-, dimethyl ester | 33.28 | C22H30O6 | 6980-45-6 |

**Supplementary Figure 5. GC-MS profile of bioactive G2 spot from AGE and the list of compound hits**

CIL/ SAIF Panjab University Chandigarh

#### Sample Header

| | |
|---|---|
| Data File: | 2 |
| Original Data Path: | C:\GCMS-data\YEAR 2016\NOV\24 |
| Sample Type: | Unknown |
| Sample ID: | 1 |
| Sample Name: | |
| Acquisition Date: | 11/24/16 11:47:34 AM |
| Run Time(min): | 30.09 |
| Injection Volume(µl): | 1.00 |
| Scans: | 3582 |
| Low Mass(m/z): | 100 |
| High Mass(m/z): | 800 |
| Instrument Method: | C:\GCMS-data\instrument method\GERNAL-gcms-METHOD.meth |

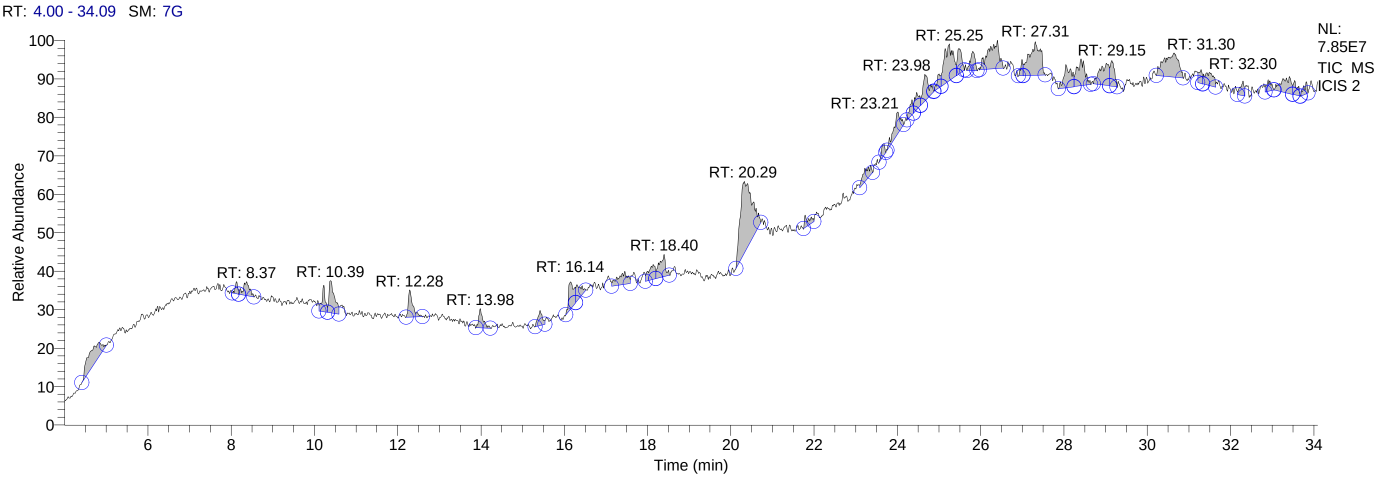


#### Qual Peak Table

| RT | Peak Area | Area % | Peak Height |
|---|---|---|---|
| 4.61 | 84959861.20 | 4.36 | 3857568.85 |
| 8.13 | 9792217.67 | 0.50 | 2471416.09 |
| 8.37 | 29366956.69 | 1.51 | 2826441.04 |
| 10.23 | 30199788.31 | 1.55 | 5411253.56 |
| 10.39 | 59572913.51 | 3.06 | 6513334.31 |
| 12.28 | 41284173.74 | 2.12 | 5542808.33 |
| 13.98 | 28329958.91 | 1.46 | 3954956.86 |
| 15.42 | 22430036.25 | 1.15 | 3057366.71 |
| 16.14 | 48562549.29 | 2.49 | 5578246.09 |
| 16.30 | 26380102.94 | 1.36 | 3398502.43 |
| 17.45 | 40607395.80 | 2.09 | 2645618.55 |
| 18.13 | 32948828.05 | 1.69 | 3016375.34 |
| 18.40 | 46259258.61 | 2.38 | 4535805.73 |
| 20.29 | 282220349.71 | 14.50 | 14729031.97 |
| 21.79 | 17337457.00 | 0.89 | 2512011.05 |
| 23.21 | 29927403.99 | 1.54 | 2777269.50 |
| 23.66 | 13597644.69 | 0.70 | 2489592.03 |
| 23.98 | 49859738.54 | 2.56 | 4513767.26 |
| 24.35 | 16523862.92 | 0.85 | 3042634.12 |
| 24.46 | 26677003.07 | 1.37 | 3558685.79 |
| 24.66 | 51193247.52 | 2.63 | 5467028.86 |
| 25.00 | 16780540.83 | 0.86 | 2788860.87 |
| 25.25 | 109834380.16 | 5.64 | 7494012.68 |
| 25.47 | 37921277.62 | 1.95 | 5258389.78 |
| 25.81 | 30186552.88 | 1.55 | 3932872.79 |
| 26.40 | 102376007.06 | 5.26 | 5764109.32 |
| 26.99 | 11543226.16 | 0.59 | 3288640.37 |
| 27.31 | 127037857.39 | 6.53 | 6827937.38 |

### CIL/ SAIF Panjab University Chandigarh

| RT | Peak Area | Area % | Peak Height |
|---|---|---|---|
| 28.05 | 56533185.75 | 2.90 | 4759974.95 |
| 28.42 | 70009395.55 | 3.60 | 5477293.25 |
| 29.02 | 66888998.34 | 3.44 | 4504627.86 |
| 29.15 | 37384176.03 | 1.92 | 5241885.31 |
| 30.64 | 117310537.05 | 6.03 | 4921677.11 |
| 31.30 | 15583577.88 | 0.80 | 2851640.13 |
| 31.49 | 38323250.08 | 1.97 | 2675615.81 |
| 32.30 | 18535080.32 | 0.95 | 2956308.09 |
| 32.90 | 16791033.15 | 0.86 | 2322260.61 |
| 33.41 | 52510721.70 | 2.70 | 3496522.30 |
| 33.54 | 18883692.88 | 0.97 | 2920566.64 |
| 33.82 | 13988083.89 | 0.72 | 2427343.61 |

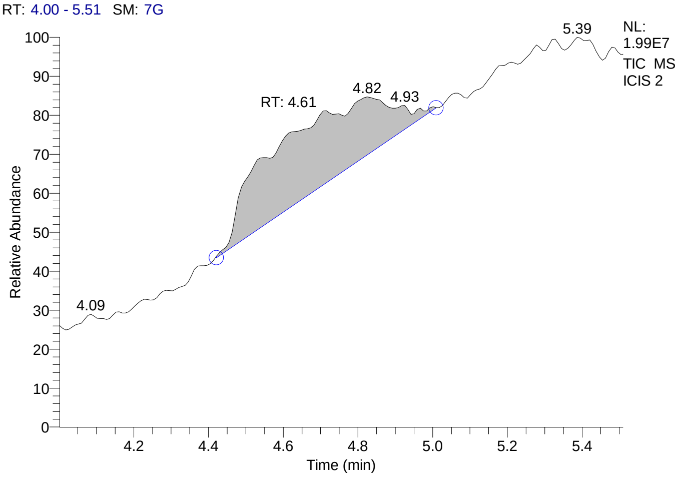


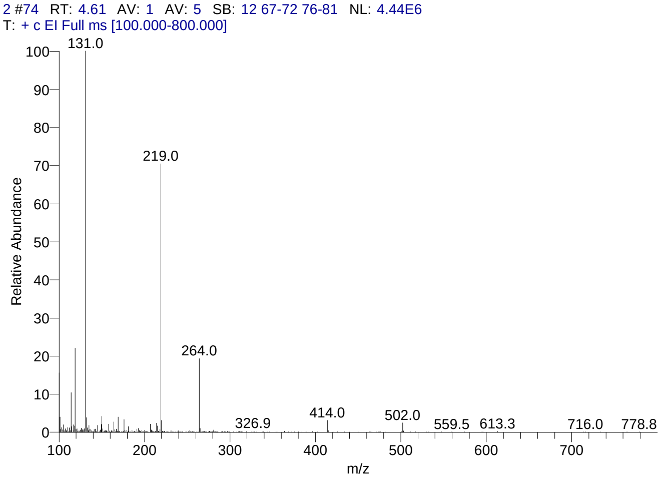


#### Library Search Results Table

| Compound Name | RT | Molecular Formula | Cas # |
|---|---|---|---|
| Perfluorotributylamine | 4.61 | C12F27N | 311-89-7 |
| Perfluoro(dibutylmethylamine) | 4.61 | C9F21N | 514-03-4 |
| Nonafluoro-1-bromobutane | 4.61 | C4BrF9 | 375-48-4 |

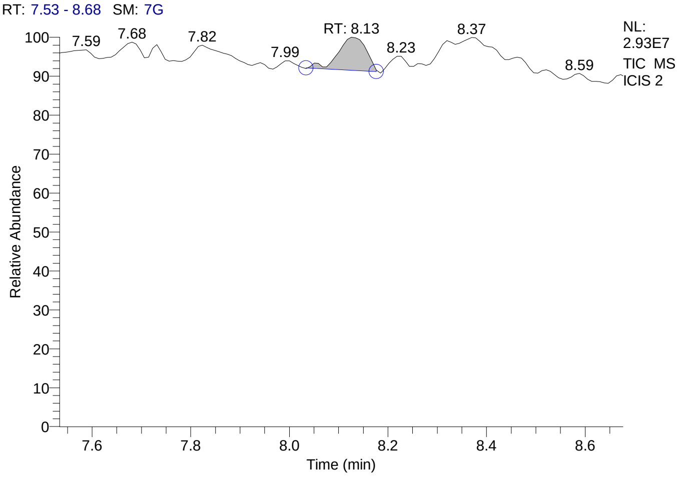


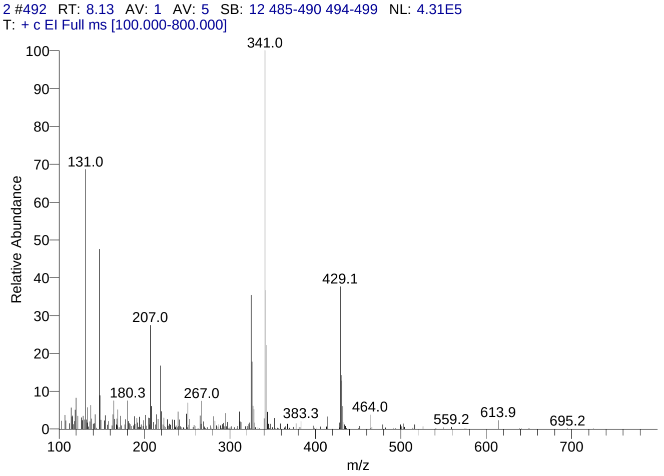

### CIL/ SAIF Panjab University Chandigarh

#### Library Search Results Table

| Compound Name | RT | Molecular Formula | Cas # |
|---|---|---|---|
| Cyclohexasiloxane, dodecamethyl- | 8.13 | C12H36O6Si6 | 540-97-6 |
| Heptasiloxane, 1,1,3,3,5,5,7,7,9,9,11,11,13,13-tetradecamethyl- | 8.13 | C14H44O6Si7 | 19095-23-9 |
| 2H-1,4-Benzodiazepin-2-one, 7-chloro-1,3-dihydro-5-phenyl-1-(trimethylsilyl)-3-[ (trimethylsilyl)oxy]- | 8.13 | C21H27ClN2O2Si2 | 55319-93-2 |

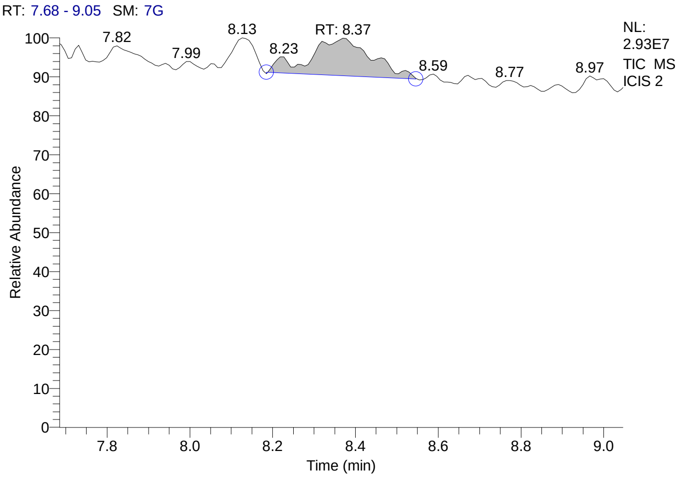


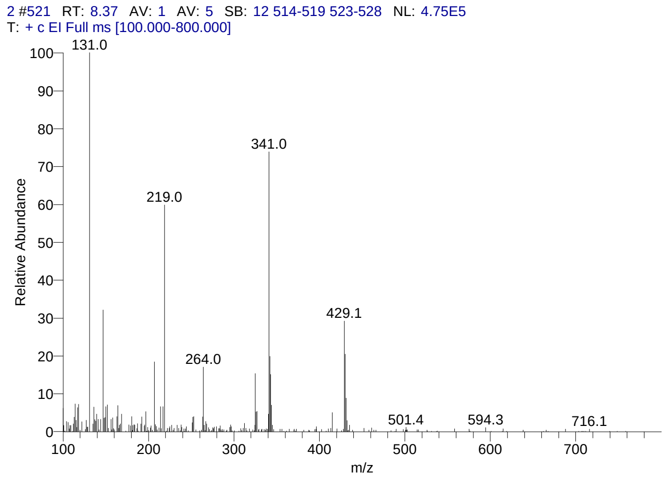


#### Library Search Results Table

| Compound Name | RT | Molecular Formula | Cas # |
|---|---|---|---|
| Cyclohexasiloxane, dodecamethyl- | 8.37 | C12H36O6Si6 | 540-97-6 |
| 2H-1,4-Benzodiazepin-2-one, 7-chloro-1,3-dihydro-5-phenyl-1-(trimethylsilyl)-3-[ (trimethylsilyl)oxy]- | 8.37 | C21H27ClN2O2Si2 | 55319-93-2 |
| 2-Amino-4-(4-bromo-phenyl)-6-methyl-5,6,7,8-tetra hydro-[1,6]naphthyridine-3-carbonitrile | 8.37 | C16H15BrN4 | 296770-49-5 |

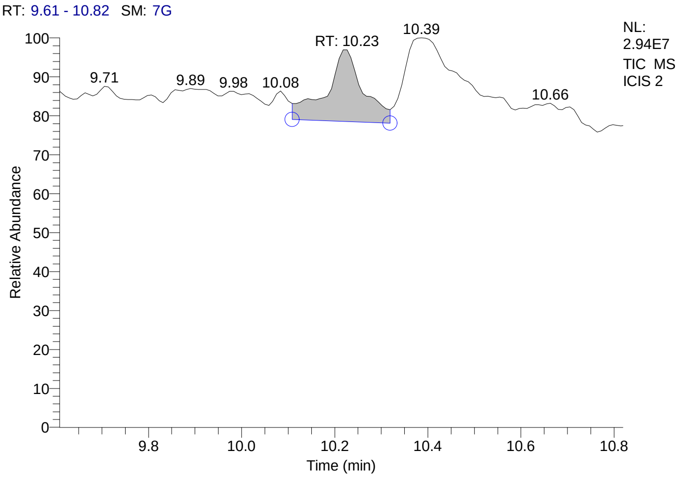


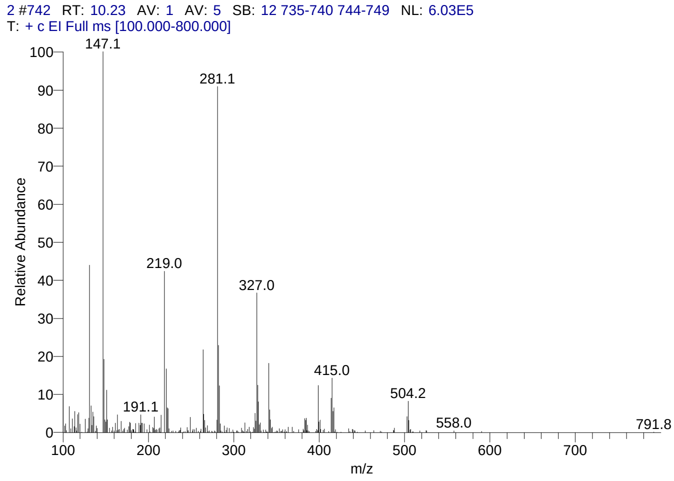


#### Library Search Results Table

| Compound Name | RT | Molecular Formula | Cas # |
|---|---|---|---|
| Cycloheptasiloxane, tetradecamethyl- | 10.23 | C14H42O7Si7 | 107-50-6 |

### CIL/ SAIF Panjab University Chandigarh

| Compound Name | RT | Molecular Formula | Cas # |
|---|---|---|---|
| 3-Isopropoxy-1,1,1,7,7,7-hexamethyl-3,5,5-tris(trimethylsiloxy)tetrasiloxane | 10.23 | C18H52O7Si7 | 71579-69-6 |
| 3-Butoxy-1,1,1,7,7,7-hexamethyl-3,5,5-tris(trimethylsiloxy)tetrasiloxane | 10.23 | C19H54O7Si7 | 72439-84-0 |

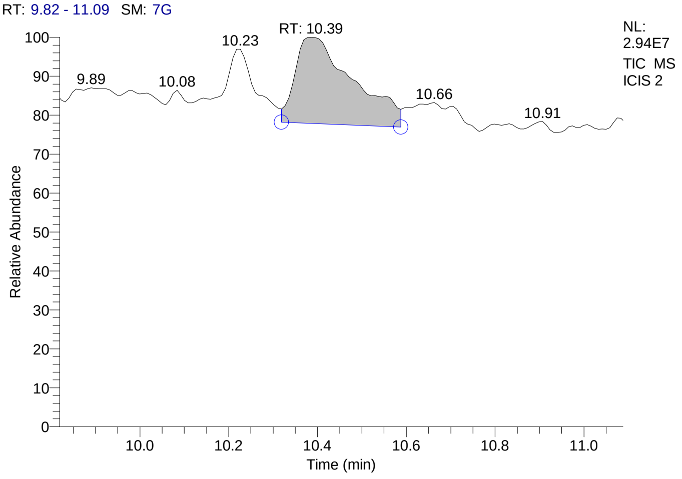


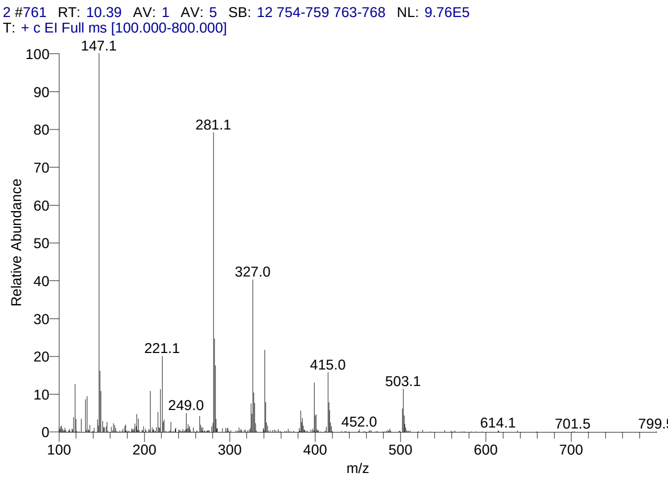


#### Library Search Results Table

| Compound Name | RT | Molecular Formula | Cas # |
|---|---|---|---|
| Cycloheptasiloxane, tetradecamethyl- | 10.39 | C14H42O7Si7 | 107-50-6 |
| 3-Isopropoxy-1,1,1,7,7,7-hexamethyl-3,5,5-tris(trimethylsiloxy)tetrasiloxane | 10.39 | C18H52O7Si7 | 71579-69-6 |
| 3-Butoxy-1,1,1,7,7,7-hexamethyl-3,5,5-tris(trimethylsiloxy)tetrasiloxane | 10.39 | C19H54O7Si7 | 72439-84-0 |

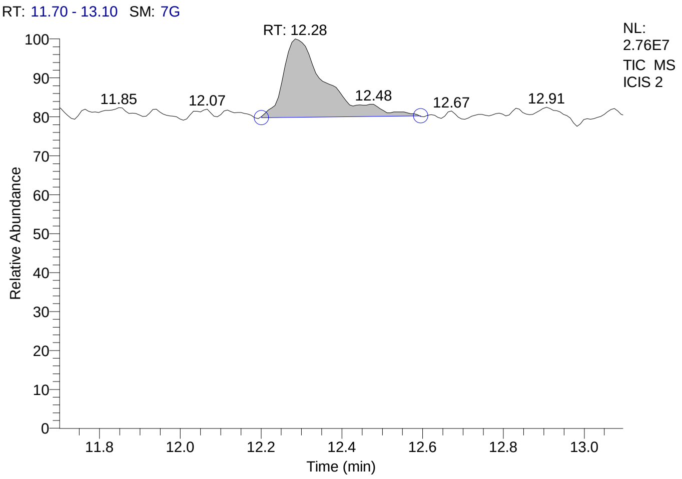


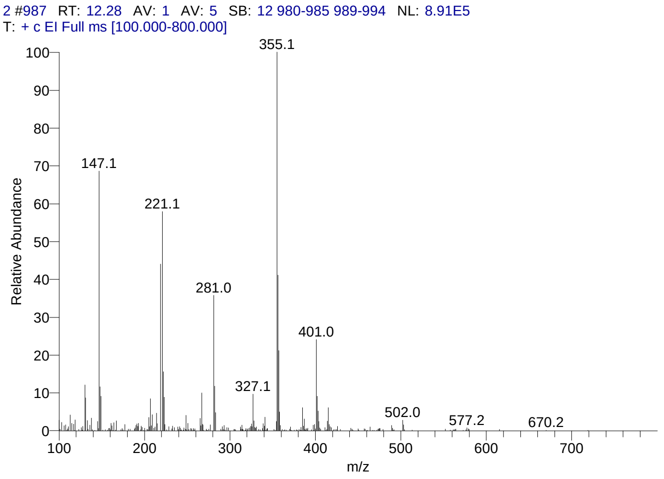


#### Library Search Results Table

| Compound Name | RT | Molecular Formula | Cas # |
|---|---|---|---|
| Cyclooctasiloxane, hexadecamethyl- | 12.28 | C16H48O8Si8 | 556-68-3 |
| Hexasiloxane, tetradecamethyl- | 12.28 | C14H42O5Si6 | 107-52-8 |
| Silane, [[4-[1,2-bis[(trimethylsilyl)oxy]ethyl]-1,2-phenylene]bis(oxy)]bis[trimethyl- | 12.28 | C20H42O4Si4 | 56114-62-6 |

### CIL/ SAIF Panjab University Chandigarh

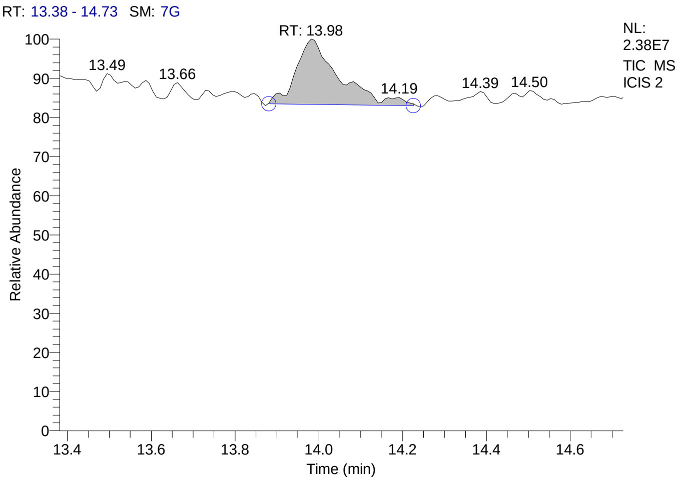


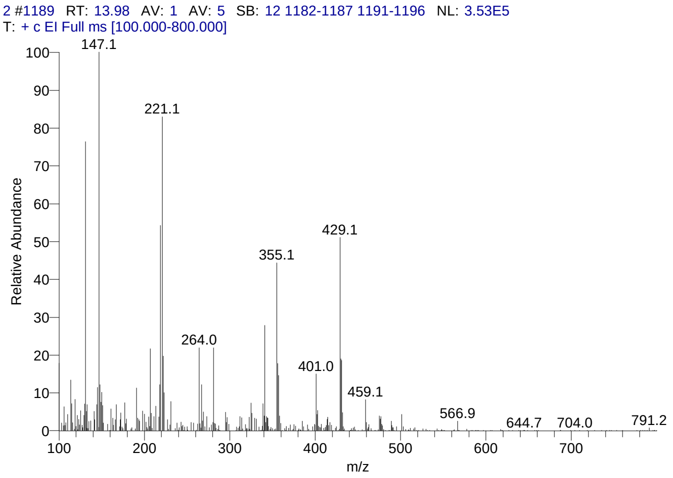


#### Library Search Results Table

| Compound Name | RT | Molecular Formula | Cas # |
|---|---|---|---|
| Cyclononasiloxane, octadecamethyl- | 13.98 | C18H54O9Si9 | 556-71-8 |
| Cyclodecasiloxane, eicosamethyl- | 13.98 | C20H60O10Si10 | 18772-36-6 |
| Tetracosamethyl-cyclododecasiloxane | 13.98 | C24H72O12Si12 | 18919-94-3 |

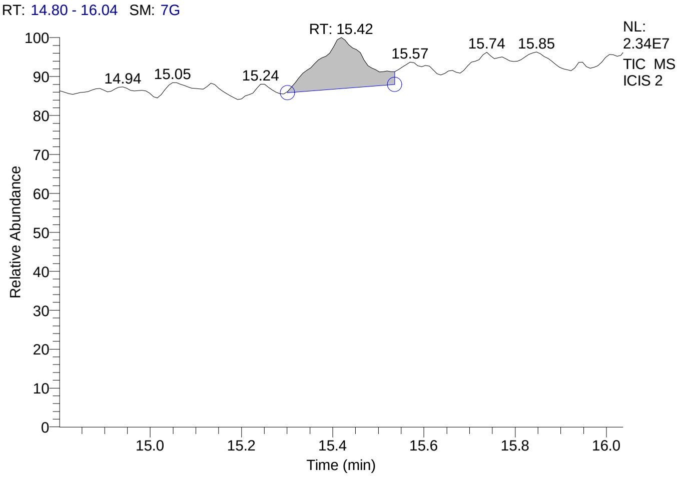


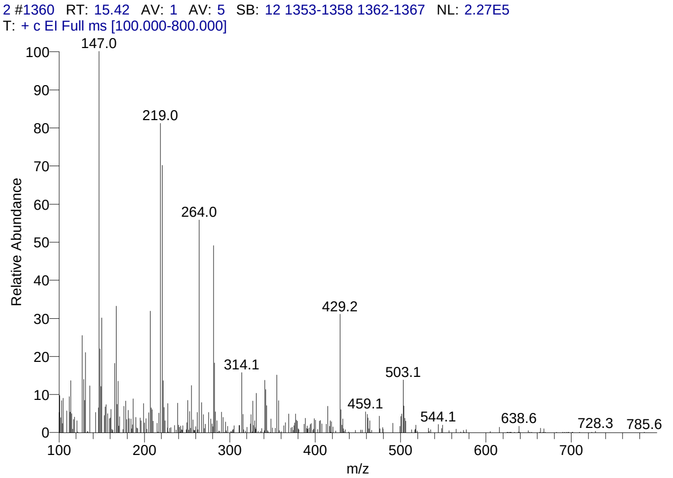


#### Library Search Results Table

| Compound Name | RT | Molecular Formula | Cas # |
|---|---|---|---|
| Cyclodecasiloxane, eicosamethyl- | 15.42 | C20H60O10Si10 | 18772-36-6 |
| Octasiloxane, 1,1,3,3,5,5,7,7,9,9,11,11,13,13,15,15-hexadecamethyl- | 15.42 | C16H50O7Si8 | 19095-24-0 |
| 1-Monolinoleoylglycerol trimethylsilyl ether | 15.42 | C27H54O4Si2 | 54284-45-6 |

### CIL/ SAIF Panjab University Chandigarh

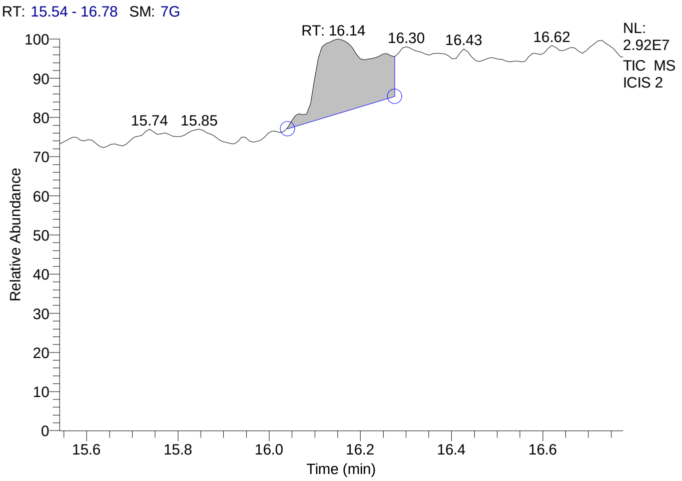


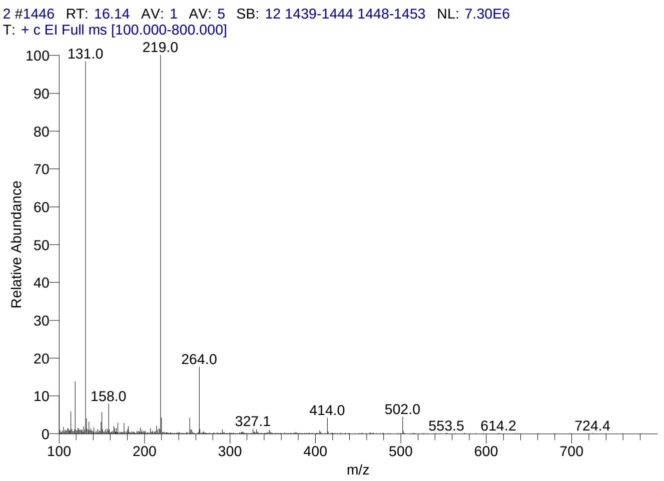


#### Library Search Results Table

| Compound Name | RT | Molecular Formula | Cas # |
|---|---|---|---|
| Perfluorotributylamine | 16.14 | C12F27N | 311-89-7 |
| Pyrimidin-2-one, 4-[N-methylureido]-1-[4-methylaminocarbonyloxy methyl | 16.14 | C13H19N5O5 | NA |
| Perfluoro(dibutylmethylamine) | 16.14 | C9F21N | 514-03-4 |

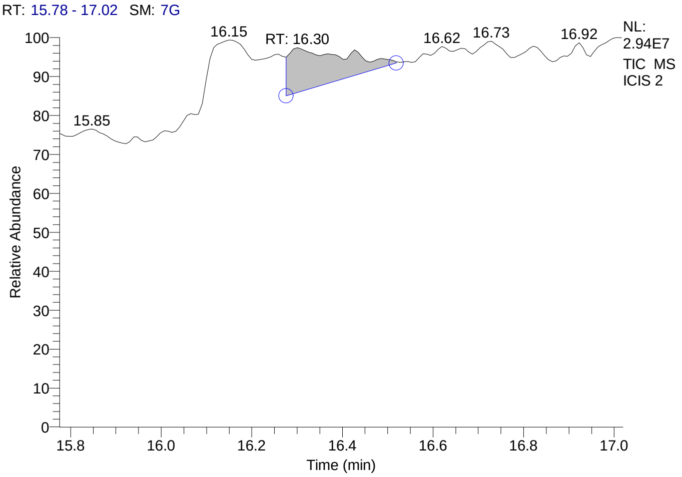


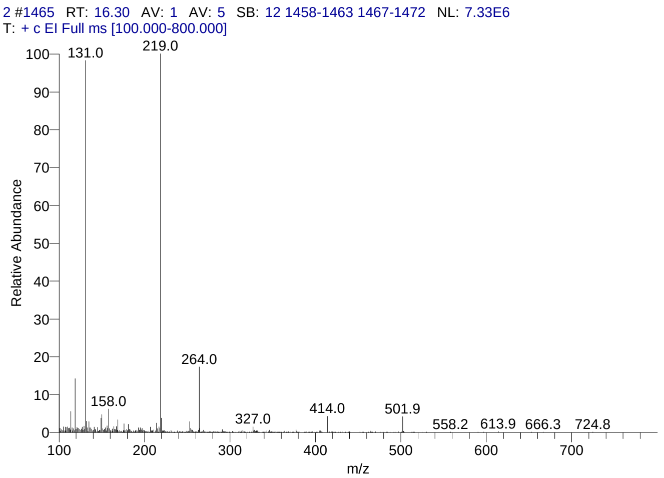


#### Library Search Results Table

| Compound Name | RT | Molecular Formula | Cas # |
|---|---|---|---|
| Perfluorotributylamine | 16.30 | C12F27N | 311-89-7 |
| Pyrimidin-2-one, 4-[N-methylureido]-1-[4-methylaminocarbonyloxy methyl | 16.30 | C13H19N5O5 | NA |
| Perfluoro(dibutylmethylamine) | 16.30 | C9F21N | 514-03-4 |

### CIL/ SAIF Panjab University Chandigarh

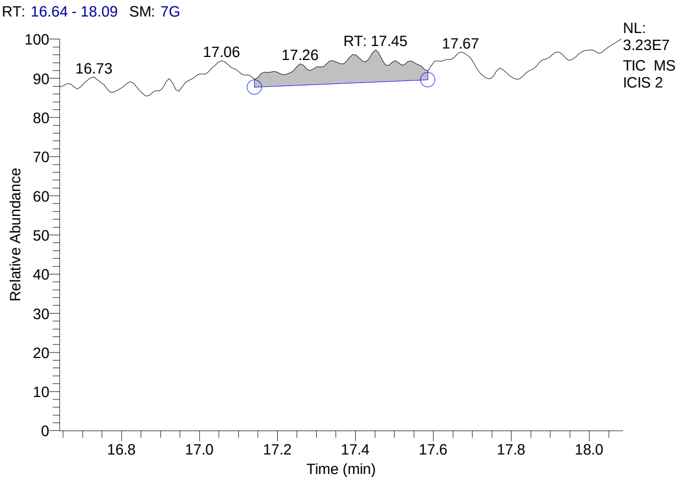


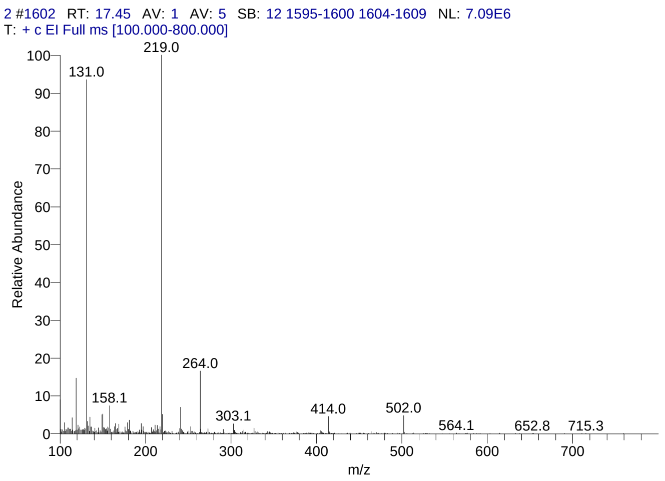


#### Library Search Results Table

| Compound Name | RT | Molecular Formula | Cas # |
|---|---|---|---|
| Perfluorotributylamine | 17.45 | C12F27N | 311-89-7 |
| Pyrimidin-2-one, 4-[N-methylureido]-1-[4-methylaminocarbonyloxy methyl | 17.45 | C13H19N5O5 | NA |
| 5á-Cholestane-3à,7à,12à,24,25,26-hexol hexa-TMS | 17.45 | C45H96O6Si6 | NA |

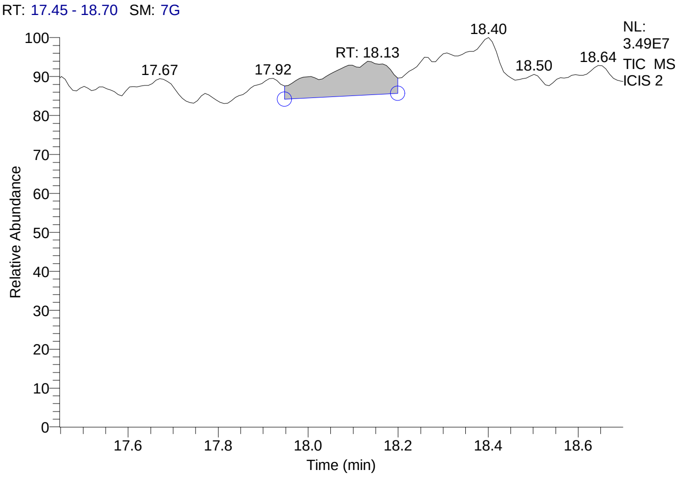


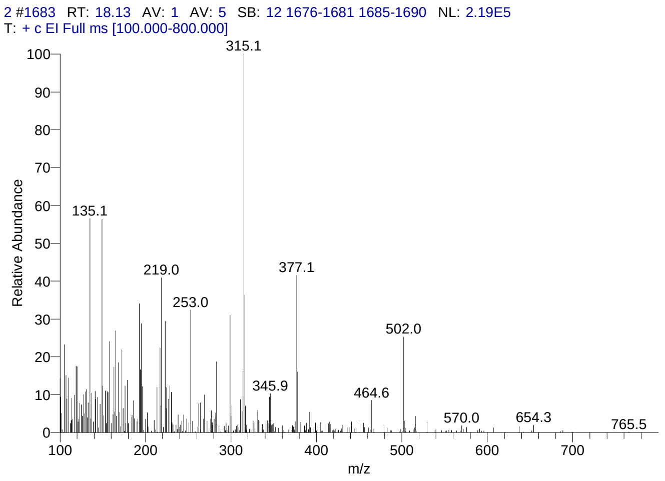


#### Library Search Results Table

| Compound Name | RT | Molecular Formula | Cas # |
|---|---|---|---|
| D-Homo-24-nor-17-oxachola-20,22-diene-3,16-dione, 7-(acetyloxy)-1,2:14,15:21,23-triepoxy-4,4,8-trimethyl-, (5à,7à,13à,14á,15á,17aà)- | 18.13 | C28H34O8 | 55658-66-7 |
| D-Homo-24-nor-17-oxachola-1,20,22-triene-3,7,16-dione, 14,15:21,23-diepoxy-4,4,8-trimethyl-, (5à,13à,14á,15á,17aà)- | 18.13 | C26H30O6 | 13072-74-7 |
| D-Homo-24-nor-17-oxachola-20,22-diene-3,16-dione, 1,2:14,15:21,23-triepoxy-7-hydroxy-4,4,8-trimethyl-, (5à,7à,13à,14á,15á,17aà)- | 18.13 | C26H32O7 | 35963-08-7 |

### CIL/ SAIF Panjab University Chandigarh

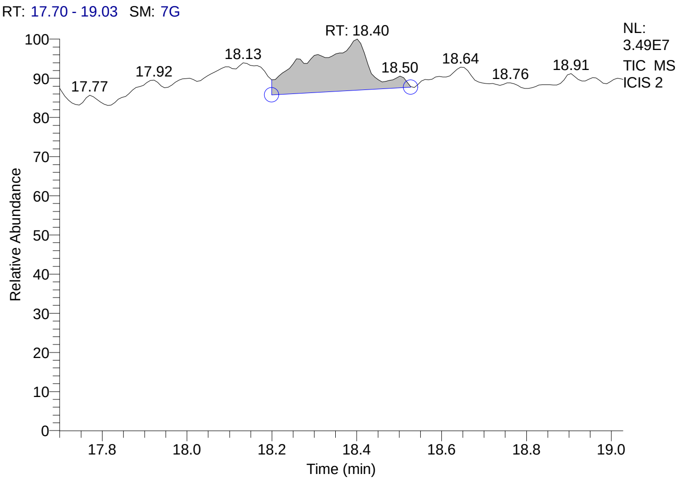


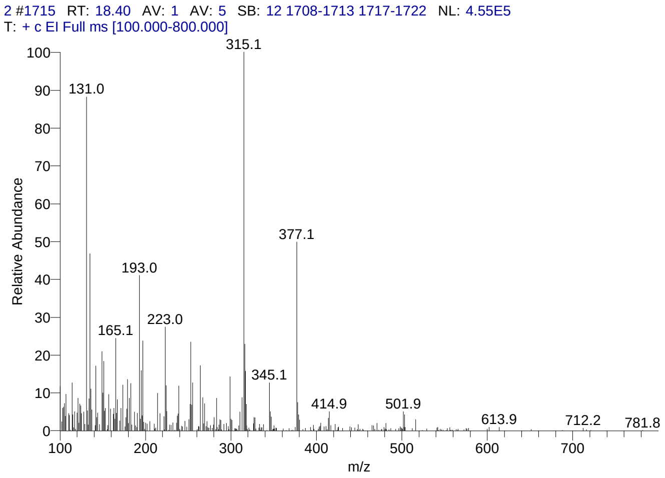


#### Library Search Results Table

| Compound Name | RT | Molecular Formula | Cas # |
|---|---|---|---|
| D-Homo-24-nor-17-oxachola-20,22-diene-3,16-dione, 7-(acetyloxy)-1,2:14,15:21,23-triepoxy-4,4,8-trimethyl-, (5à,7à,13à,14á,15á,17aà)- | 18.40 | C28H34O8 | 55658-66-7 |
| D-Homo-24-nor-17-oxachola-1,20,22-triene-3,7,16-dione, 14,15:21,23-diepoxy-4,4,8-trimethyl-, (5à,13à,14á,15á,17aà)- | 18.40 | C26H30O6 | 13072-74-7 |
| D-Homo-24-nor-17-oxachola-20,22-diene-3,16-dione, 1,2:14,15:21,23-triepoxy-7-hydroxy-4,4,8-trimethyl-, (5à,7à,13à,14á,15á,17aà)- | 18.40 | C26H32O7 | 35963-08-7 |

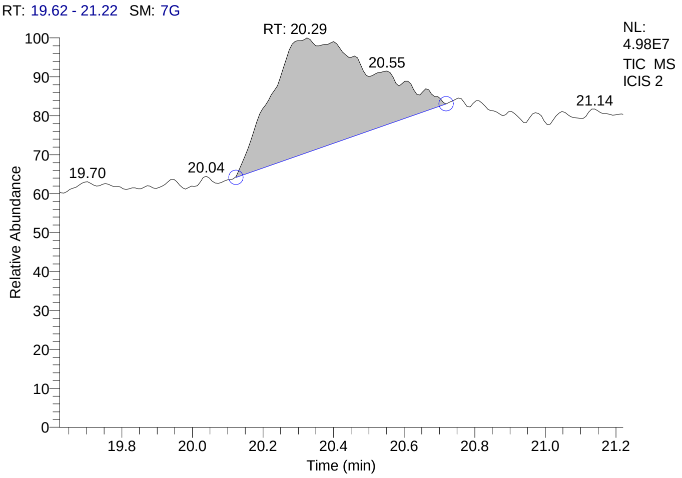


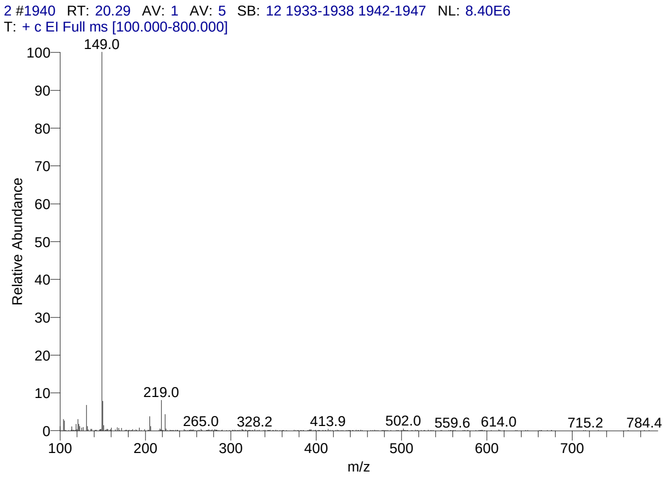


#### Library Search Results Table

| Compound Name | RT | Molecular Formula | Cas # |
|---|---|---|---|
| Phthalic acid, butyl 2-pentyl ester | 20.29 | C17H24O4 | NA |
| Phthalic acid, butyl hex-3-yl ester | 20.29 | C18H26O4 | NA |
| Phthalic acid, butyl hept-4-yl ester | 20.29 | C19H28O4 | NA |

### CIL/ SAIF Panjab University Chandigarh

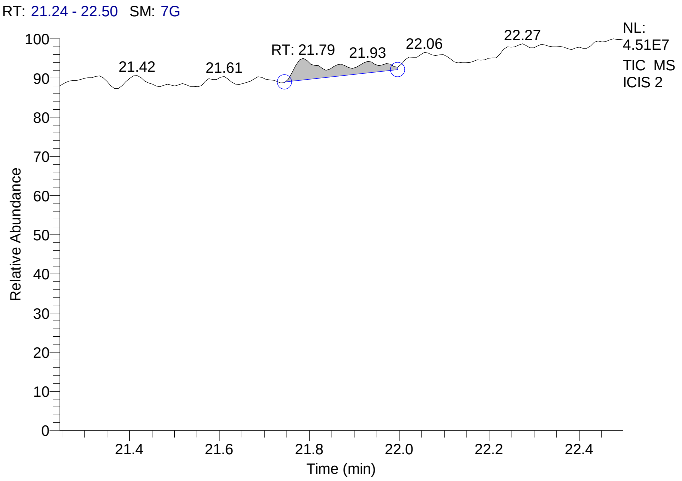


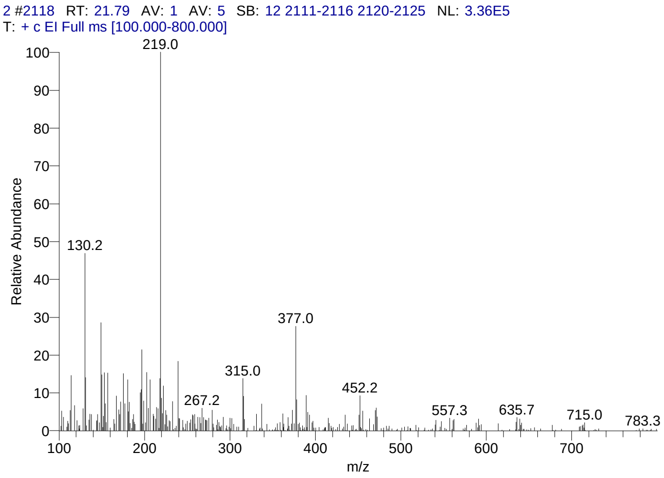


#### Library Search Results Table

| Compound Name | RT | Molecular Formula | Cas # |
|---|---|---|---|
| 1'-Carboethoxy-1'-cyano-1á,2á-dihydro-17á-propionoxy-3'H-cycloprop[1,2]androsta-1,4,6-trien-3-one | 21.79 | C27H33NO5 | 75857-77-1 |
| Acetic acid, 17-[1-hydroxy-1,5-dimethyl-5-(tetrahydropyran-2-yloxy)-hex-2-ynyl]-10,13-dimethyl-11-oxohexadecahydrocyclopenta[a]phenanthr-3-yl (ester) | 21.79 | C34H52O6 | NA |
| 4-Normethyl-9,19-cyclolanoststan-7-one, 3-acetoxy- | 21.79 | C31H50O3 | NA |

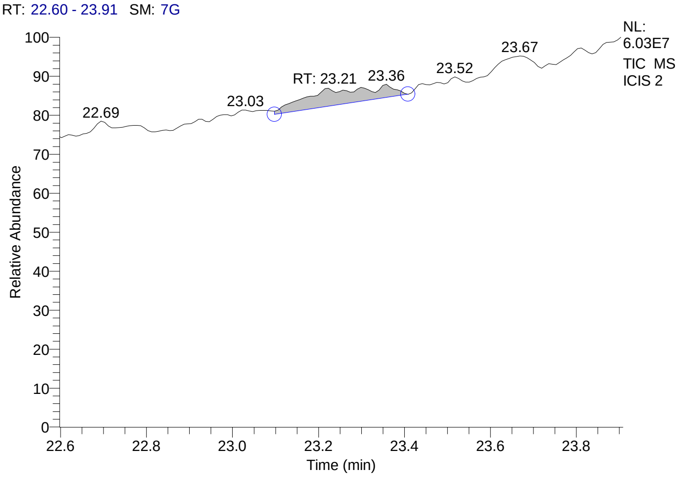


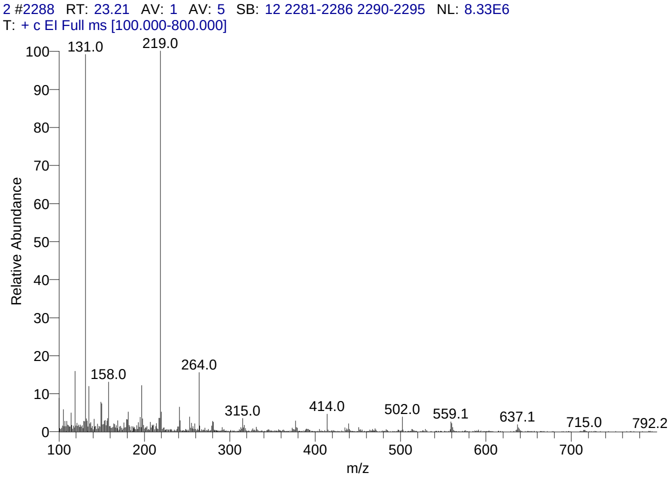


#### Library Search Results Table

| Compound Name | RT | Molecular Formula | Cas # |
|---|---|---|---|
| 5á-Cholestane-3à,7à,12à,24,25,26-hexol hexa-TMS | 23.21 | C45H96O6Si6 | NA |
| Rhodoxanthin | 23.21 | C40H50O2 | 116-30-3 |
| Olean-12-ene-3,16,21,22,23,28-hexol, (3á,4à,16à,21á,22à)- | 23.21 | C30H50O6 | 13844-22-9 |

### CIL/ SAIF Panjab University Chandigarh

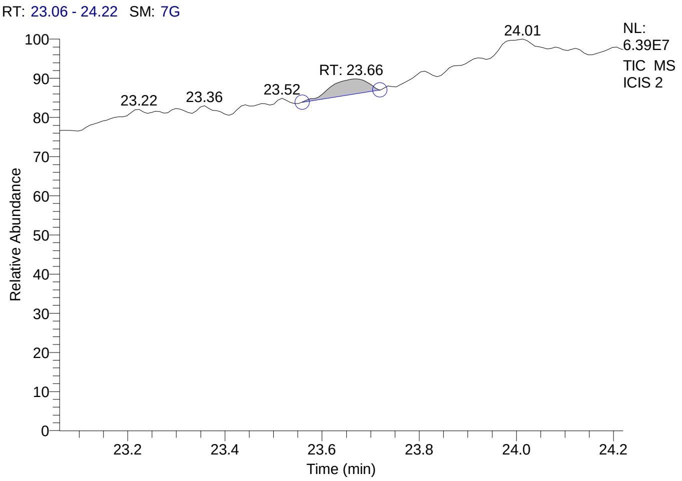


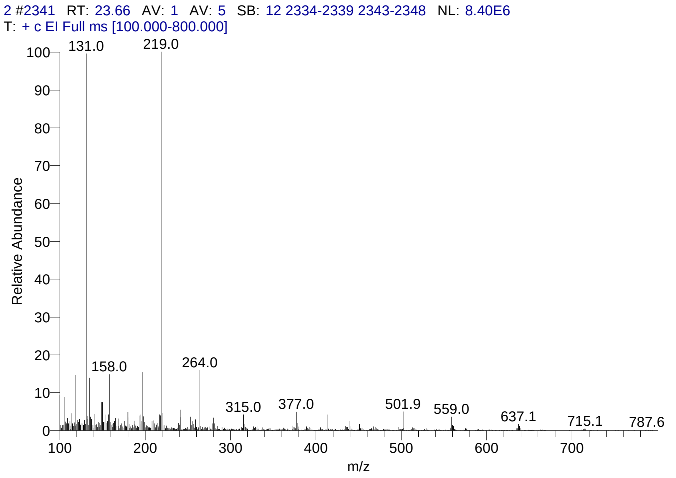


#### Library Search Results Table

| Compound Name | RT | Molecular Formula | Cas # |
|---|---|---|---|
| Olean-12-ene-3,16,21,22,23,28-hexol, (3á,4à,16à,21á,22à)- | 23.66 | C30H50O6 | 13844-22-9 |
| 5á-Cholestane-3à,7à,12à,24,25,26-hexol hexa-TMS | 23.66 | C45H96O6Si6 | NA |
| Olean-12-ene-3,16,21,22,23,28-hexol, (3á,4á,16à,21á,22à)- | 23.66 | C30H50O6 | 20853-07-0 |

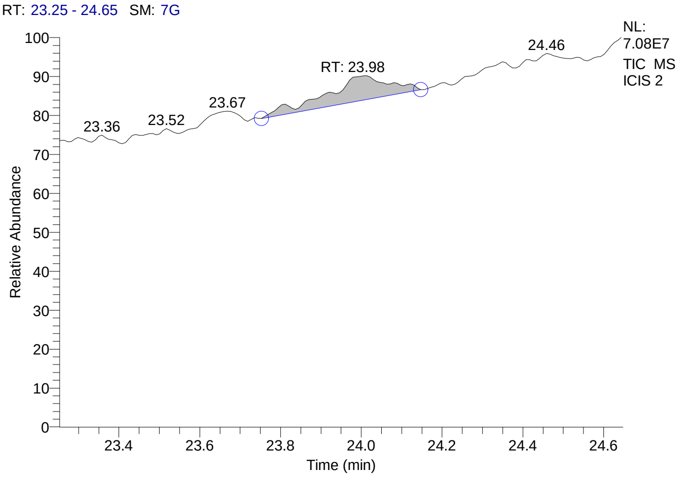


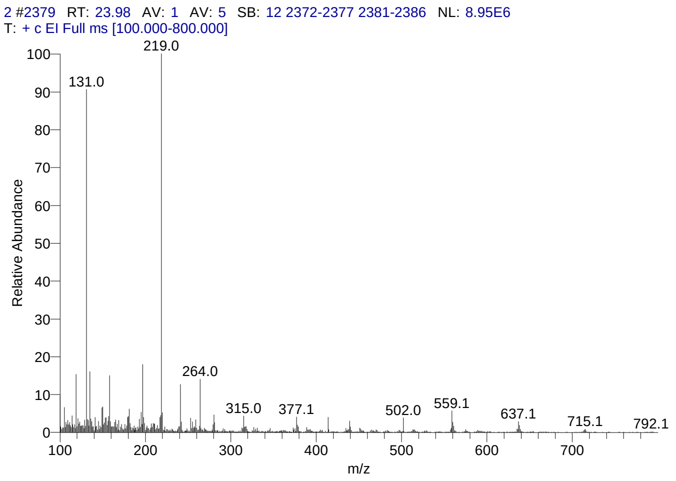


#### Library Search Results Table

| Compound Name | RT | Molecular Formula | Cas # |
|---|---|---|---|
| Rhodoxanthin | 23.98 | C40H50O2 | 116-30-3 |
| Olean-12-ene-3,16,21,22,23,28-hexol, (3á,4à,16à,21á,22à)- | 23.98 | C30H50O6 | 13844-22-9 |
| 5á-Cholestane-3à,7à,12à,24,25,26-hexol hexa-TMS | 23.98 | C45H96O6Si6 | NA |

### CIL/ SAIF Panjab University Chandigarh

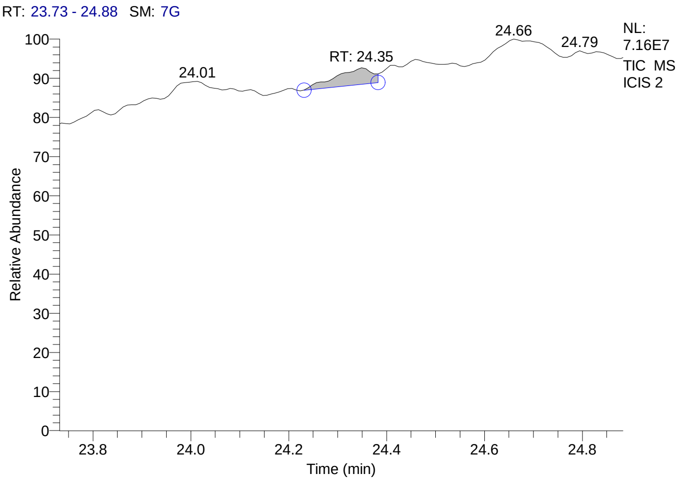


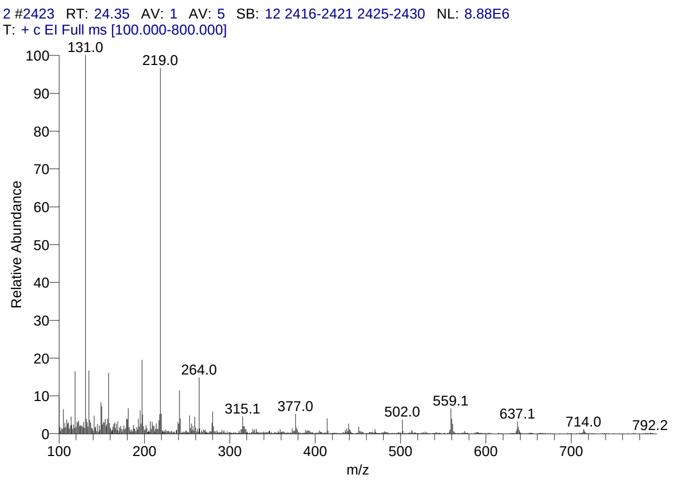


#### Library Search Results Table

| Compound Name | RT | Molecular Formula | Cas # |
|---|---|---|---|
| Rhodoxanthin | 24.35 | C40H50O2 | 116-30-3 |
| Olean-12-ene-3,16,21,22,23,28-hexol, (3á,4à,16à,21á,22à)- | 24.35 | C30H50O6 | 13844-22-9 |
| 5á-Cholestane-3à,7à,12à,24,25,26-hexol hexa-TMS | 24.35 | C45H96O6Si6 | NA |

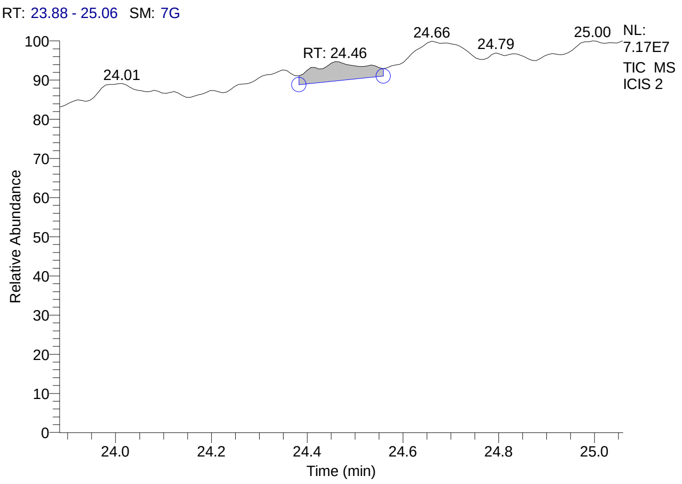


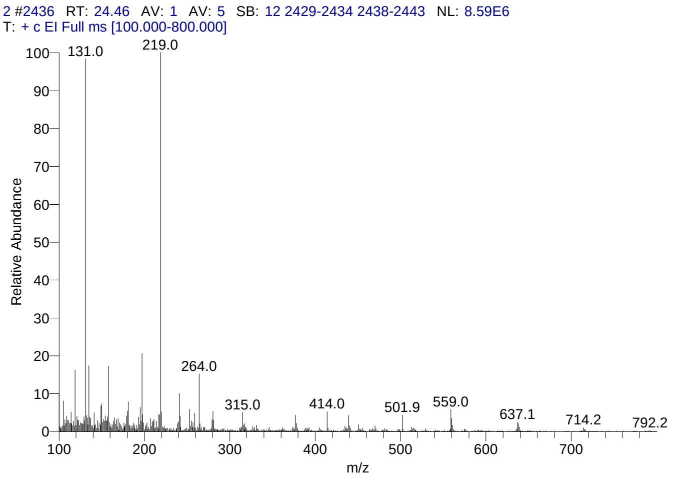


#### Library Search Results Table

| Compound Name | RT | Molecular Formula | Cas # |
|---|---|---|---|
| Rhodoxanthin | 24.46 | C40H50O2 | 116-30-3 |
| Olean-12-ene-3,16,21,22,23,28-hexol, (3á,4à,16à,21á,22à)- | 24.46 | C30H50O6 | 13844-22-9 |
| 5á-Cholestane-3à,7à,12à,24,25,26-hexol hexa-TMS | 24.46 | C45H96O6Si6 | NA |

### CIL/ SAIF Panjab University Chandigarh

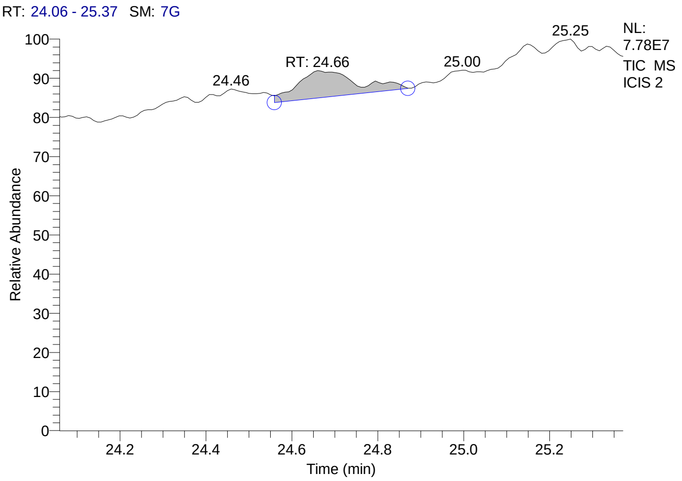


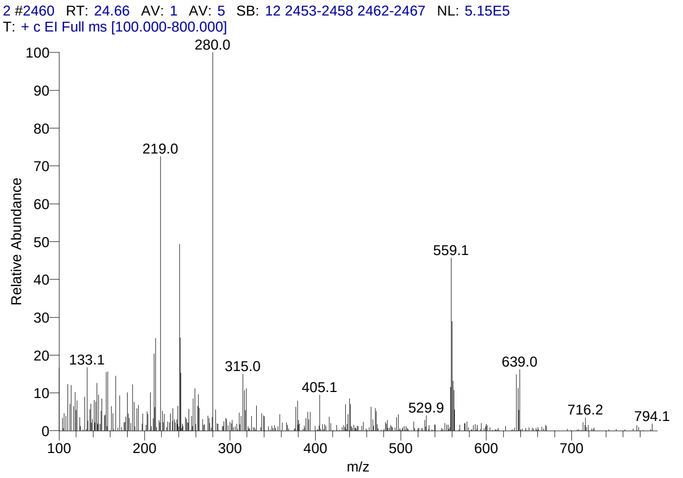


#### Library Search Results Table

| Compound Name | RT | Molecular Formula | Cas # |
|---|---|---|---|
| Rhodoxanthin | 24.66 | C40H50O2 | 116-30-3 |
| Octaphenylcyclotetrasiloxane | 24.66 | C48H40O4Si4 | 546-56-5 |
| Lanostane-7,11-dione, 3,18-bis(acetyloxy)-, cyclic 7-(1,2-ethanediyl mercaptole), (3á,20.xi.)- | 24.66 | C36H58O5S2 | 56298-07-8 |

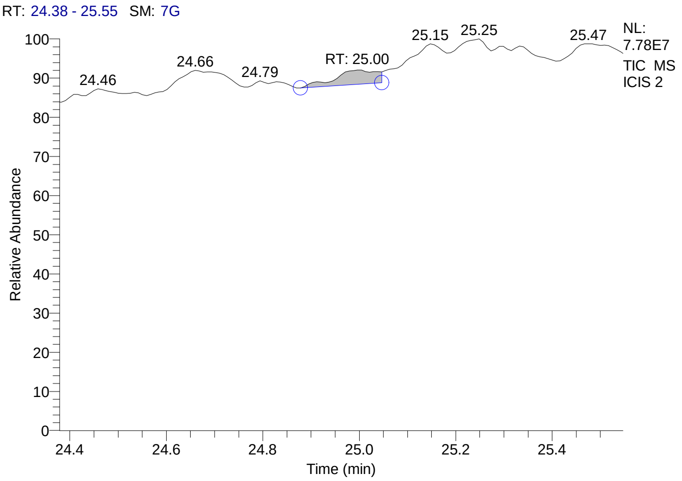


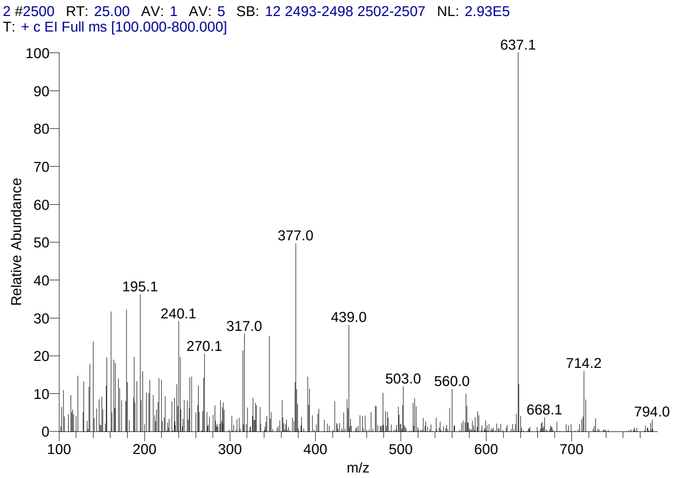


#### Library Search Results Table

| Compound Name | RT | Molecular Formula | Cas # |
|---|---|---|---|
| Pregn-4-ene-3,20-dione, 11,17,21-tris[(trimethylsilyl)oxy]-, bis(O-methyloxime), (11á)- | 25.00 | C32H60N2O5Si3 | 32221-26-4 |
| Rhodoxanthin | 25.00 | C40H50O2 | 116-30-3 |
| Pregn-5-en-20-one, 3,16,17,21-tetrakis[(trimethylsilyl)oxy]-, O-(phenylmethyl)oxime, (3á,16à)- | 25.00 | C40H71NO5Si4 | 57326-04-2 |

### CIL/ SAIF Panjab University Chandigarh

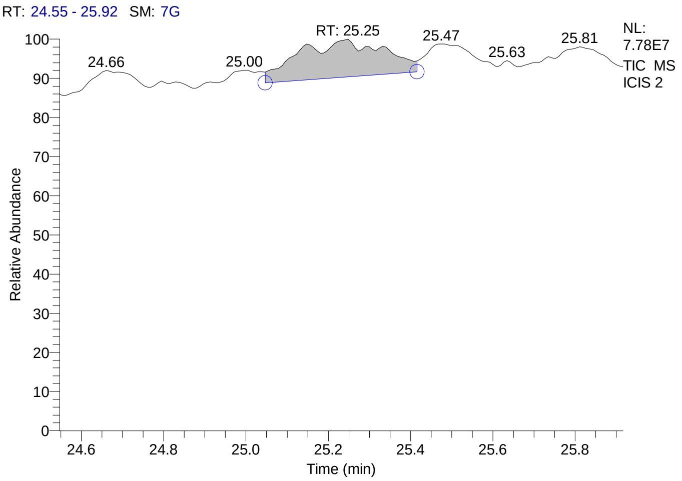


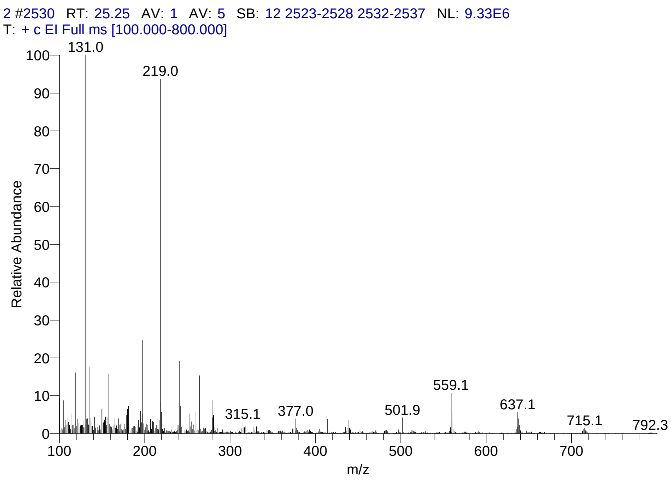


#### Library Search Results Table

| Compound Name | RT | Molecular Formula | Cas # |
|---|---|---|---|
| Rhodoxanthin | 25.25 | C40H50O2 | 116-30-3 |
| Olean-12-ene-3,16,21,22,23,28-hexol, (3á,4à,16à,21á,22à)- | 25.25 | C30H50O6 | 13844-22-9 |
| 4aà,4bá-Gibbane-1à,10á-dicarboxylic acid, 4a-formyl-7-hydroxy-1-methyl-8-methylene-, dimethyl ester | 25.25 | C22H30O6 | 6980-45-6 |

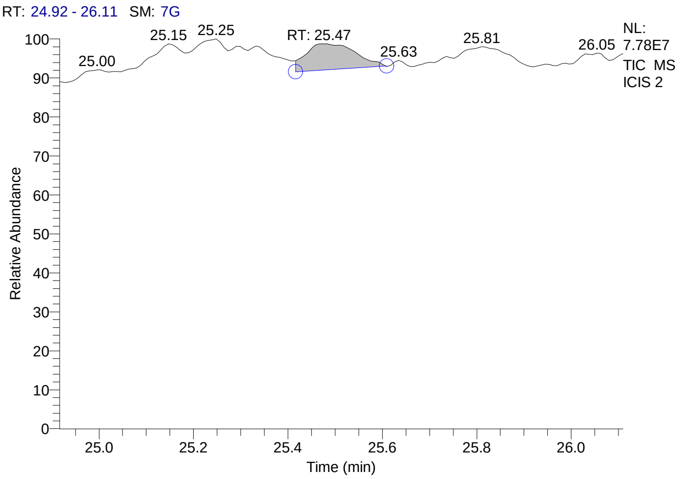


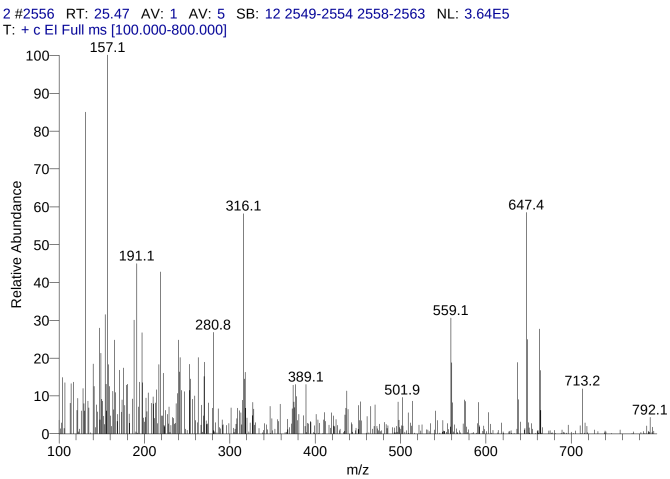


#### Library Search Results Table

| Compound Name | RT | Molecular Formula | Cas # |
|---|---|---|---|
| Pregn-5-en-20-one, 3,16,17,21-tetrakis[(trimethylsilyl)oxy]-, O-(phenylmethyl)oxime, (3á,16à)- | 25.47 | C40H71NO5Si4 | 57326-04-2 |
| Lanostane-7,11-dione, 3,18-bis(acetyloxy)-, cyclic 7-(1,2-ethanediyl mercaptole), (3á,20.xi.)- | 25.47 | C36H58O5S2 | 56298-07-8 |
| [(1H)-Pyrrole-3-propanoic acid, 2-ethoxycarbonyl-4-ethoxycarbonylmethyl]-5,5'-methylene, bis-, diethyl ester | 25.47 | C33H46N2O12 | 116377-74-3 |

### CIL/ SAIF Panjab University Chandigarh

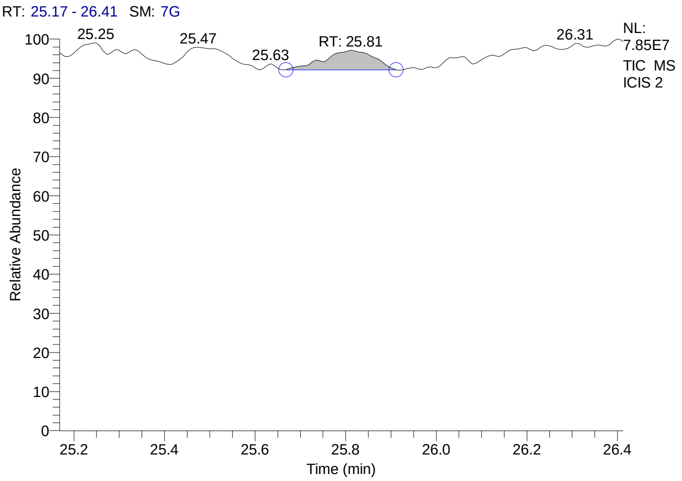


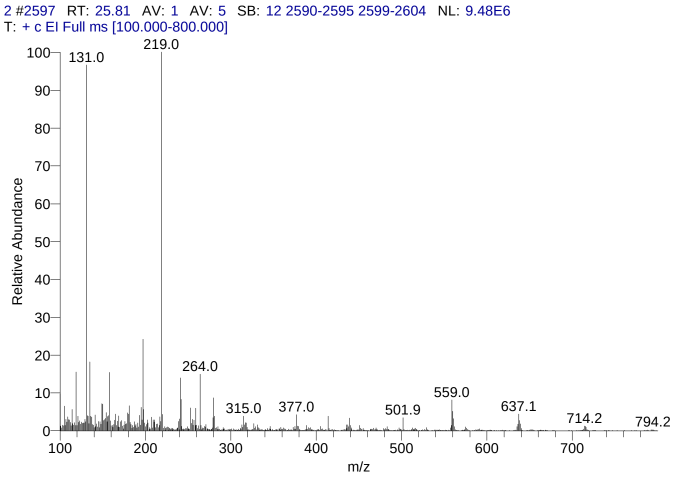


#### Library Search Results Table

| Compound Name | RT | Molecular Formula | Cas # |
|---|---|---|---|
| Rhodoxanthin | 25.81 | C40H50O2 | 116-30-3 |
| Olean-12-ene-3,16,21,22,23,28-hexol, (3á,4à,16à,21á,22à)- | 25.81 | C30H50O6 | 13844-22-9 |
| 5á-Cholestane-3à,7à,12à,24,25,26-hexol hexa-TMS | 25.81 | C45H96O6Si6 | NA |

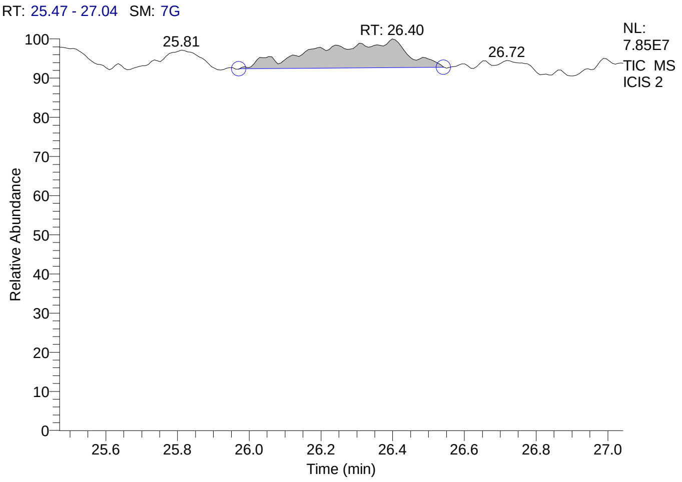


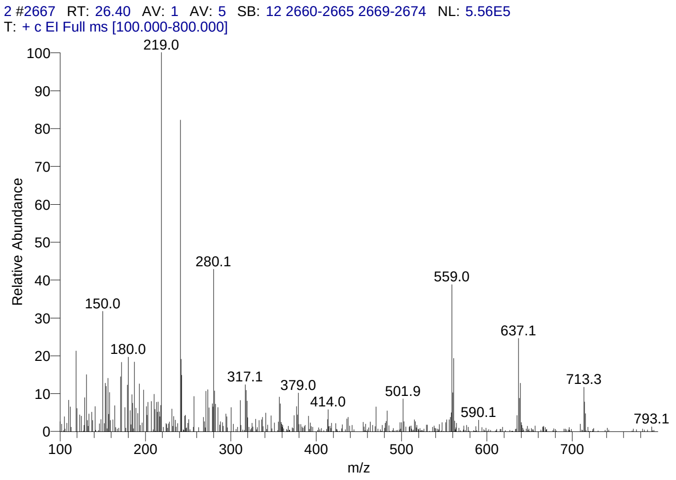


#### Library Search Results Table

| Compound Name | RT | Molecular Formula | Cas # |
|---|---|---|---|
| Octaphenylcyclotetrasiloxane | 26.40 | C48H40O4Si4 | 546-56-5 |
| Benzene, hexakis(1-bromoethyl)- | 26.40 | C18H24Br6 | 87773-60-2 |
| Pregn-4-en-18-al, 3,20-bis(methoxyimino)-11,21-bis[(trimethylsilyl)oxy]-, O-methyloxime, (11á,17à)- | 26.40 | C30H53N3O5Si2 | 69854-81-5 |

### CIL/ SAIF Panjab University Chandigarh

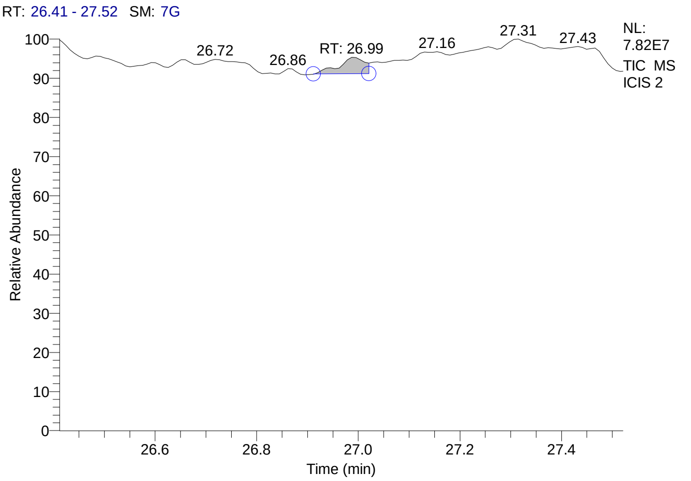


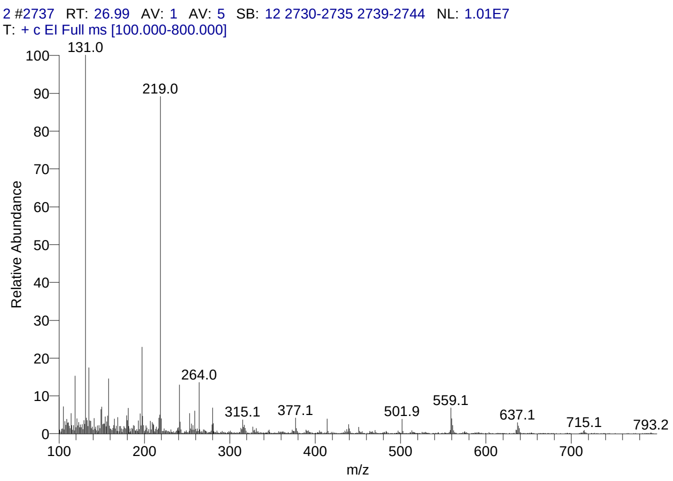


#### Library Search Results Table

| Compound Name | RT | Molecular Formula | Cas # |
|---|---|---|---|
| Rhodoxanthin | 26.99 | C40H50O2 | 116-30-3 |
| Olean-12-ene-3,16,21,22,23,28-hexol, (3á,4à,16à,21á,22à)- | 26.99 | C30H50O6 | 13844-22-9 |
| 5á-Cholestane-3à,7à,12à,24,25,26-hexol hexa-TMS | 26.99 | C45H96O6Si6 | NA |

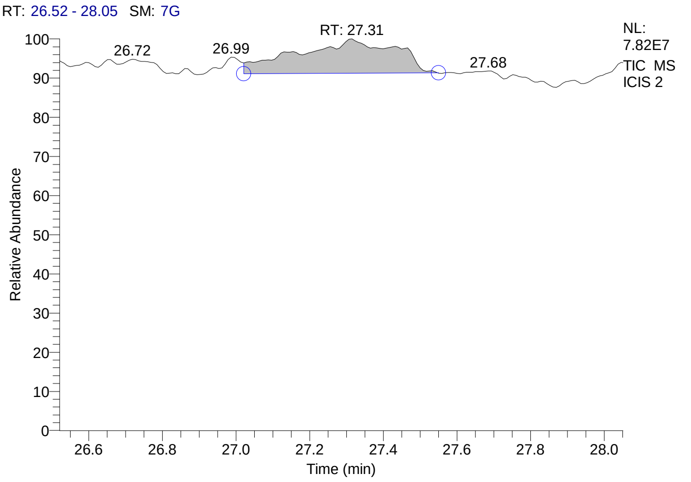


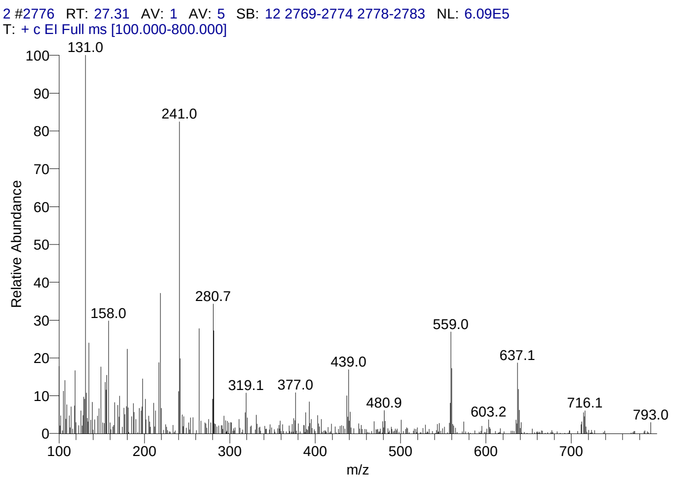


#### Library Search Results Table

| Compound Name | RT | Molecular Formula | Cas # |
|---|---|---|---|
| Octaphenylcyclotetrasiloxane | 27.31 | C48H40O4Si4 | 546-56-5 |
| Benzene, hexakis(1-bromoethyl)- | 27.31 | C18H24Br6 | 87773-60-2 |
| Pregn-4-ene-3,20-dione, 11,17,21-tris[(trimethylsilyl)oxy]-, bis(O-methyloxime), (11á)- | 27.31 | C32H60N2O5Si3 | 32221-26-4 |

### CIL/ SAIF Panjab University Chandigarh

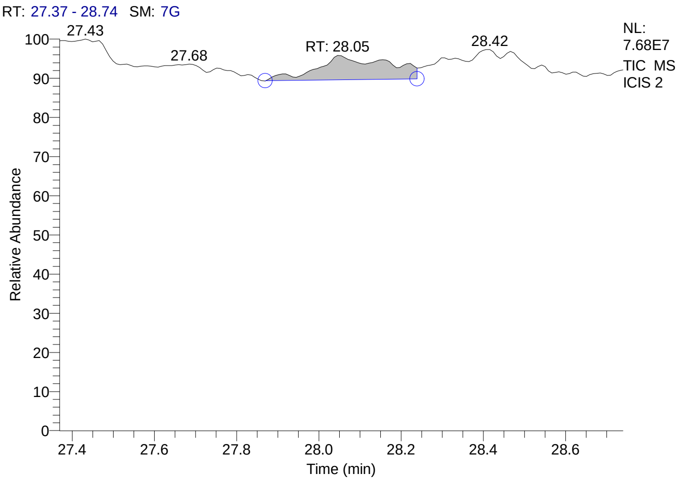


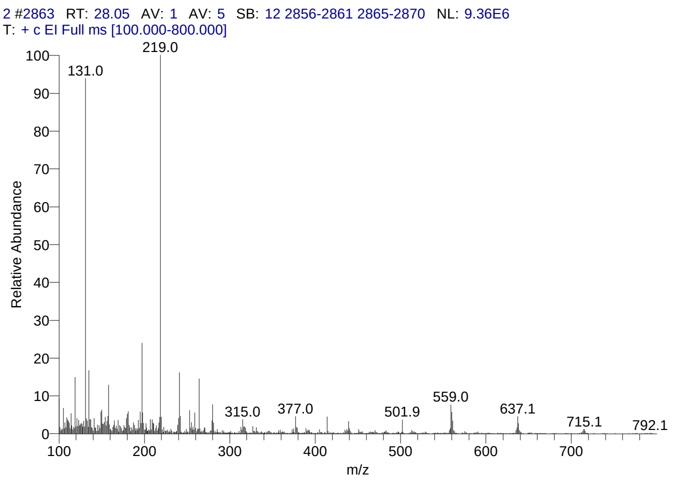


#### Library Search Results Table

| Compound Name | RT | Molecular Formula | Cas # |
|---|---|---|---|
| Rhodoxanthin | 28.05 | C40H50O2 | 116-30-3 |
| Olean-12-ene-3,16,21,22,23,28-hexol, (3á,4à,16à,21á,22à)- | 28.05 | C30H50O6 | 13844-22-9 |
| 5á-Cholestane-3à,7à,12à,24,25,26-hexol hexa-TMS | 28.05 | C45H96O6Si6 | NA |

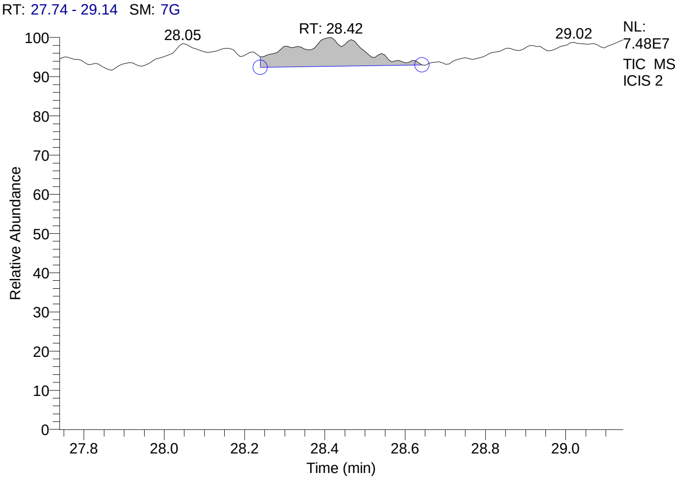


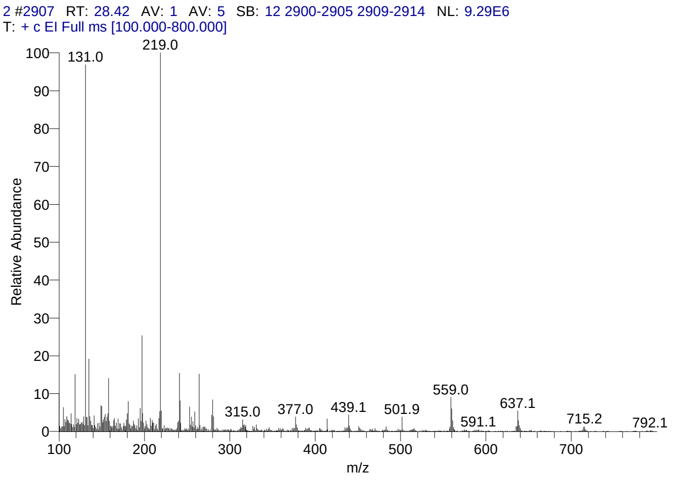


#### Library Search Results Table

| Compound Name | RT | Molecular Formula | Cas # |
|---|---|---|---|
| 5á-Cholestane-3à,7à,12à,24,25,26-hexol hexa-TMS | 28.42 | C45H96O6Si6 | NA |
| Olean-12-ene-3,16,21,22,23,28-hexol, (3á,4à,16à,21á,22à)- | 28.42 | C30H50O6 | 13844-22-9 |
| Withaferin A | 28.42 | C28H38O6 | 5119-48-2 |

### CIL/ SAIF Panjab University Chandigarh

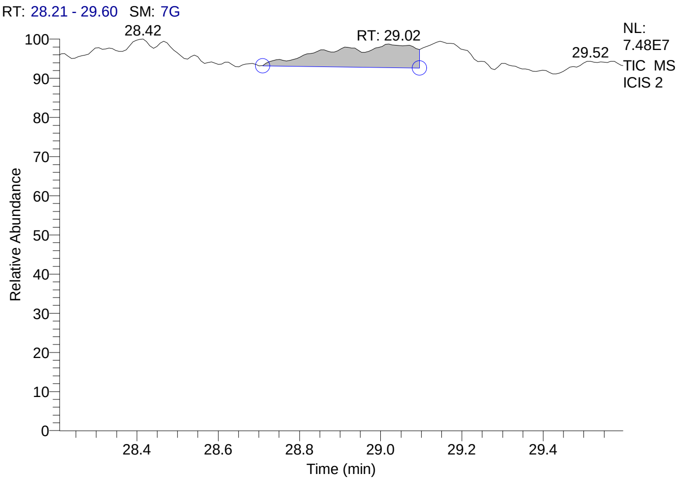


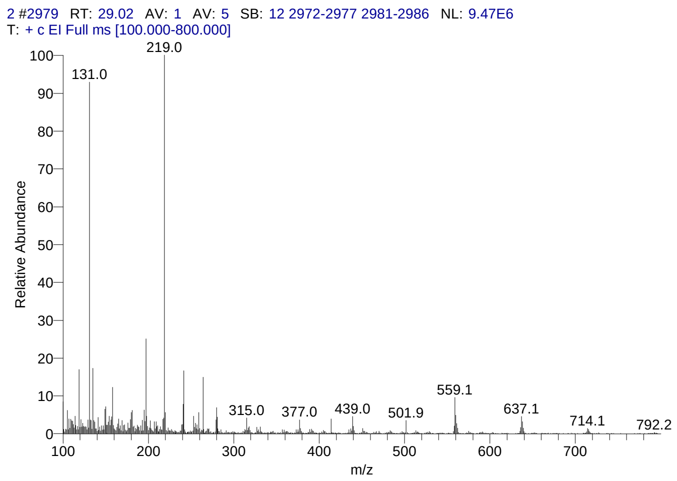


#### Library Search Results Table

| Compound Name | RT | Molecular Formula | Cas # |
|---|---|---|---|
| Rhodoxanthin | 29.02 | C40H50O2 | 116-30-3 |
| Olean-12-ene-3,16,21,22,23,28-hexol, (3á,4à,16à,21á,22à)- | 29.02 | C30H50O6 | 13844-22-9 |
| 5á-Cholestane-3à,7à,12à,24,25,26-hexol hexa-TMS | 29.02 | C45H96O6Si6 | NA |

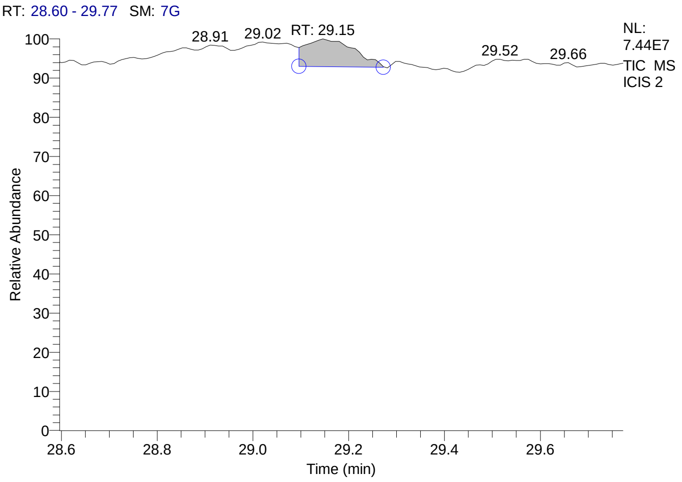


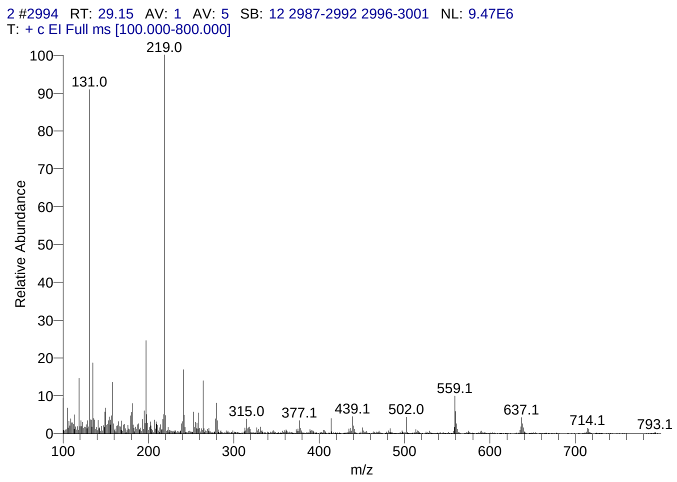


#### Library Search Results Table

| Compound Name | RT | Molecular Formula | Cas # |
|---|---|---|---|
| Rhodoxanthin | 29.15 | C40H50O2 | 116-30-3 |
| Olean-12-ene-3,16,21,22,23,28-hexol, (3á,4à,16à,21á,22à)- | 29.15 | C30H50O6 | 13844-22-9 |
| 5á-Cholestane-3à,7à,12à,24,25,26-hexol hexa-TMS | 29.15 | C45H96O6Si6 | NA |

### CIL/ SAIF Panjab University Chandigarh

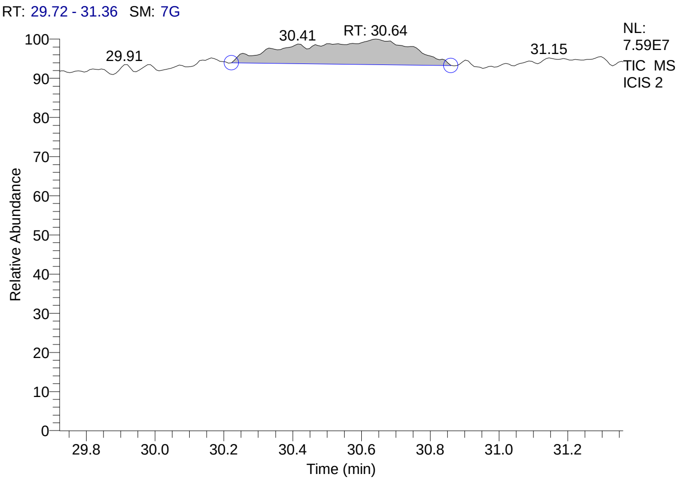


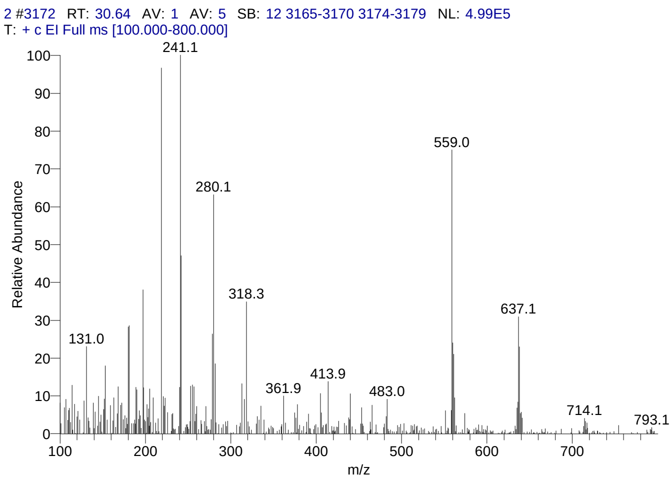


#### Library Search Results Table

| Compound Name | RT | Molecular Formula | Cas # |
|---|---|---|---|
| Octaphenylcyclotetrasiloxane | 30.64 | C48H40O4Si4 | 546-56-5 |
| Rhodoxanthin | 30.64 | C40H50O2 | 116-30-3 |
| Thiophene, 2,3,5-tris(diphenylphosphino)- | 30.64 | C40H31P3S | NA |

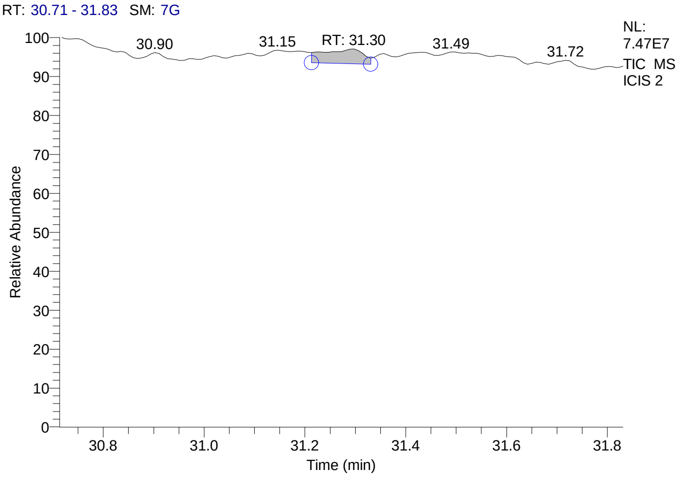


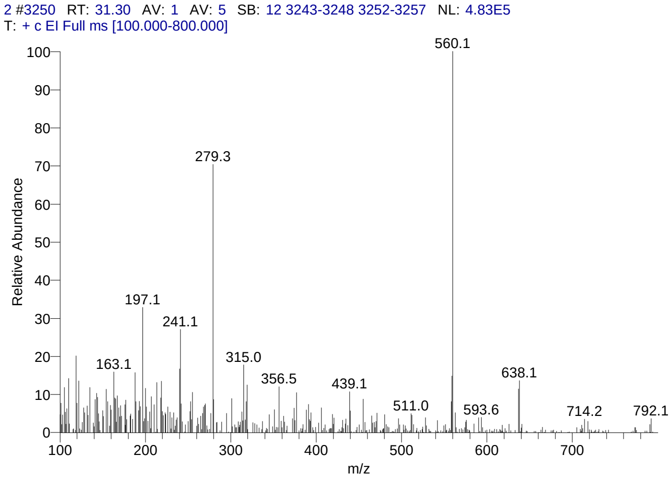


#### Library Search Results Table

| Compound Name | RT | Molecular Formula | Cas # |
|---|---|---|---|
| Lanostane-7,11-dione, 3,18-bis(acetyloxy)-, cyclic 7-(1,2-ethanediyl mercaptole), (3á,20.xi.)- | 31.30 | C36H58O5S2 | 56298-07-8 |
| Betamethasone valerate | 31.30 | C27H37FO6 | 2152-44-5 |
| Cholestano[7,8-a]cyclobutane, 3-methoxy-6-oxo-2'-methylene- | 31.30 | C31H50O2 | NA |

### CIL/ SAIF Panjab University Chandigarh

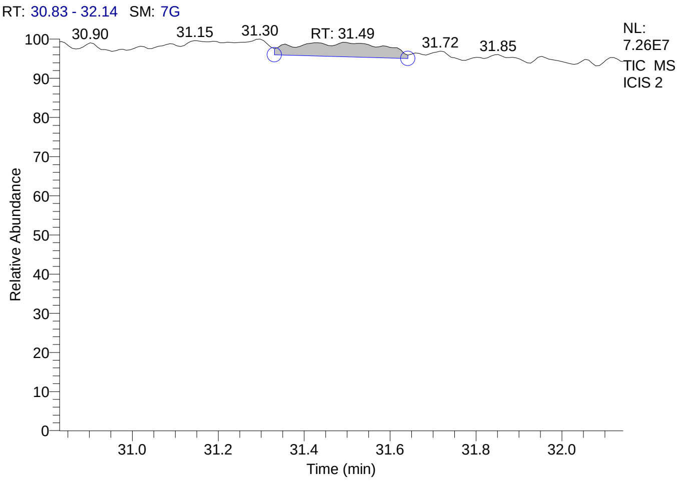


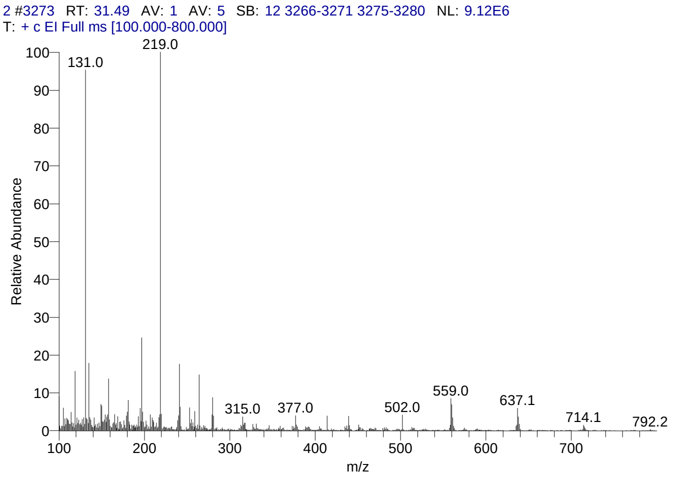


#### Library Search Results Table

| Compound Name | RT | Molecular Formula | Cas # |
|---|---|---|---|
| Rhodoxanthin | 31.49 | C40H50O2 | 116-30-3 |
| Olean-12-ene-3,16,21,22,23,28-hexol, (3á,4à,16à,21á,22à)- | 31.49 | C30H50O6 | 13844-22-9 |
| 5á-Cholestane-3à,7à,12à,24,25,26-hexol hexa-TMS | 31.49 | C45H96O6Si6 | NA |

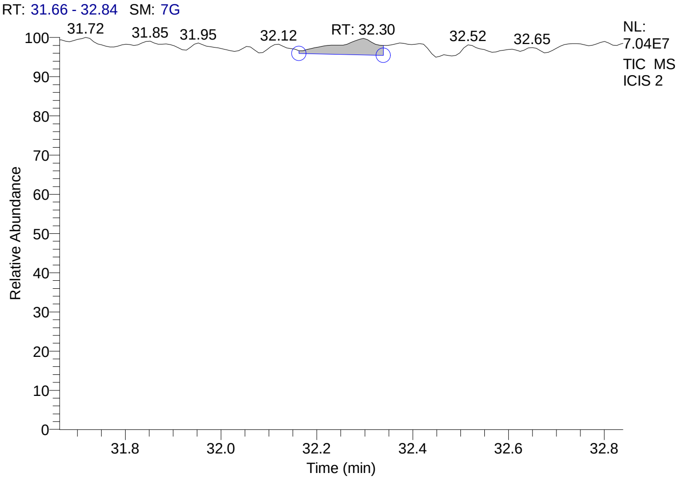


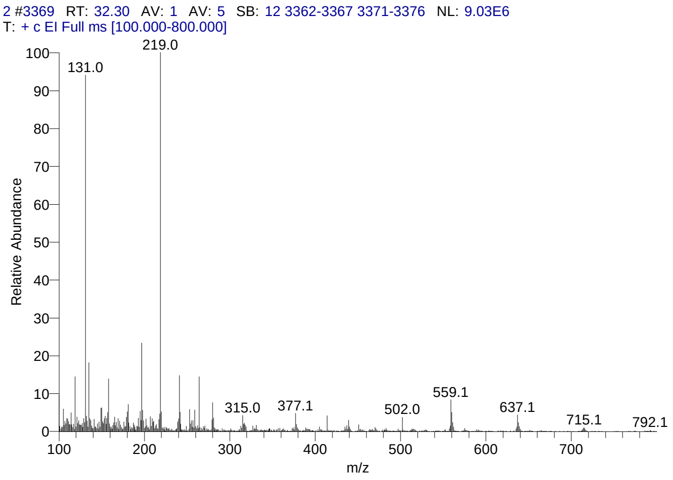


#### Library Search Results Table

| Compound Name | RT | Molecular Formula | Cas # |
|---|---|---|---|
| Rhodoxanthin | 32.30 | C40H50O2 | 116-30-3 |
| Olean-12-ene-3,16,21,22,23,28-hexol, (3á,4à,16à,21á,22à)- | 32.30 | C30H50O6 | 13844-22-9 |
| 5á-Cholestane-3à,7à,12à,24,25,26-hexol hexa-TMS | 32.30 | C45H96O6Si6 | NA |

### CIL/ SAIF Panjab University Chandigarh

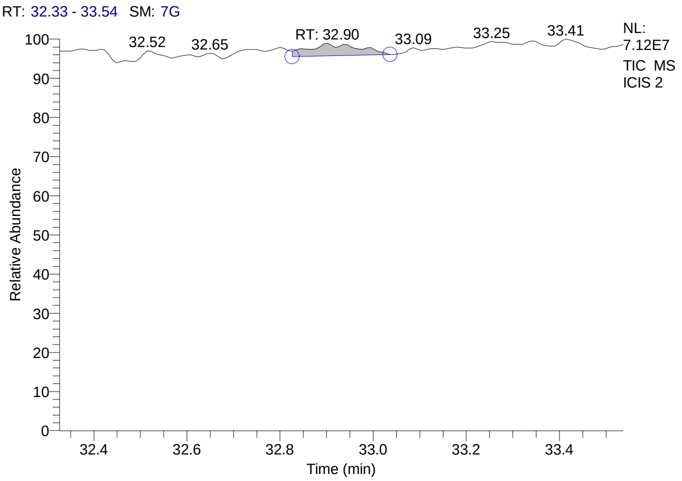


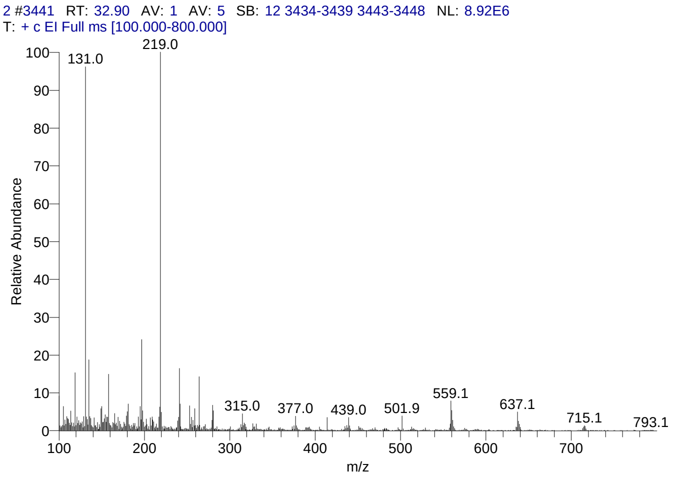


#### Library Search Results Table

| Compound Name | RT | Molecular Formula | Cas # |
|---|---|---|---|
| Rhodoxanthin | 32.90 | C40H50O2 | 116-30-3 |
| 5á-Cholestane-3à,7à,12à,24,25,26-hexol hexa-TMS | 32.90 | C45H96O6Si6 | NA |
| Olean-12-ene-3,16,21,22,23,28-hexol, (3á,4à,16à,21á,22à)- | 32.90 | C30H50O6 | 13844-22-9 |

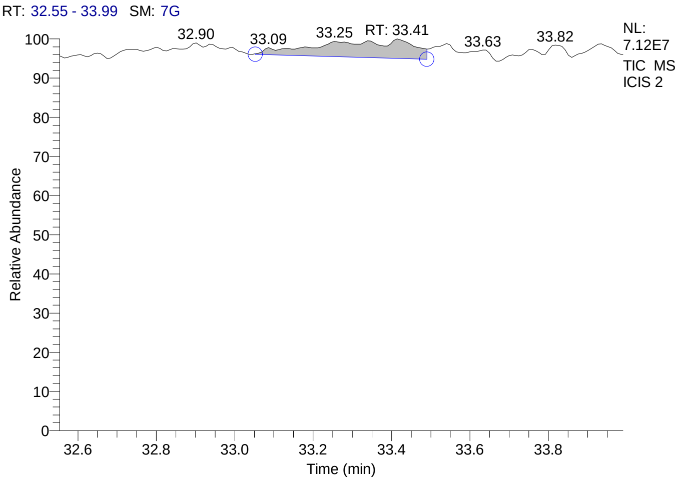


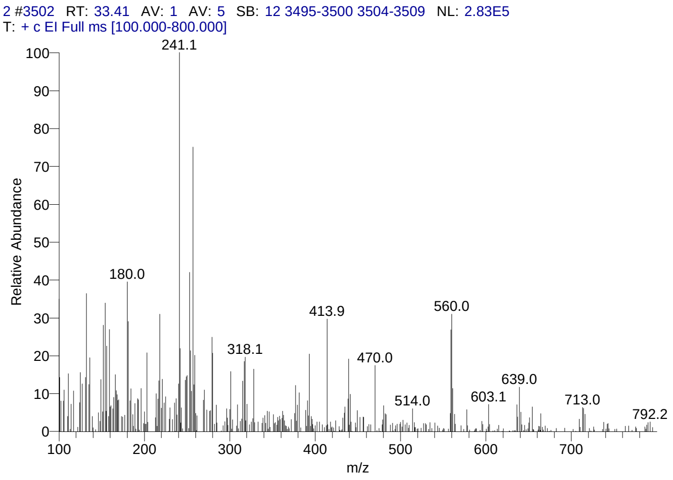


#### Library Search Results Table

| Compound Name | RT | Molecular Formula | Cas # |
|---|---|---|---|
| Rhodoxanthin | 33.41 | C40H50O2 | 116-30-3 |
| Pregn-5-en-20-one, 3,16,17,21-tetrakis[(trimethylsilyl)oxy]-, O-(phenylmethyl)oxime, (3á,16à)- | 33.41 | C40H71NO5Si4 | 57326-04-2 |
| Pregn-5-en-20-one, 3,11,17,21-tetrakis[(trimethylsilyl)oxy]-, O-methyloxime, (3á,11á)- | 33.41 | C34H67NO5Si4 | 57325-94-7 |

### CIL/ SAIF Panjab University Chandigarh

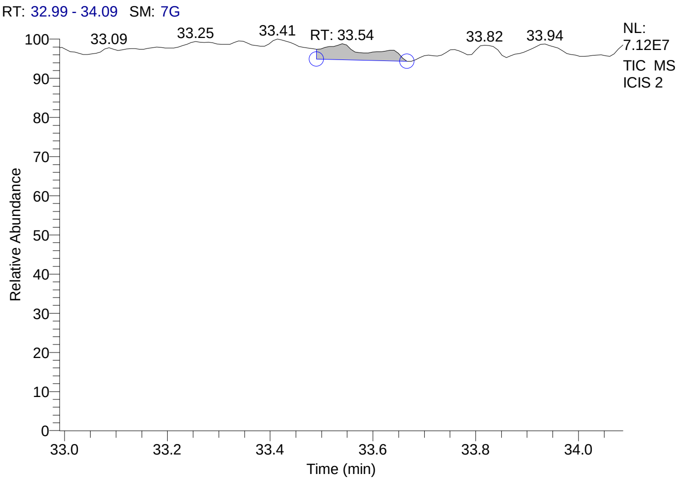


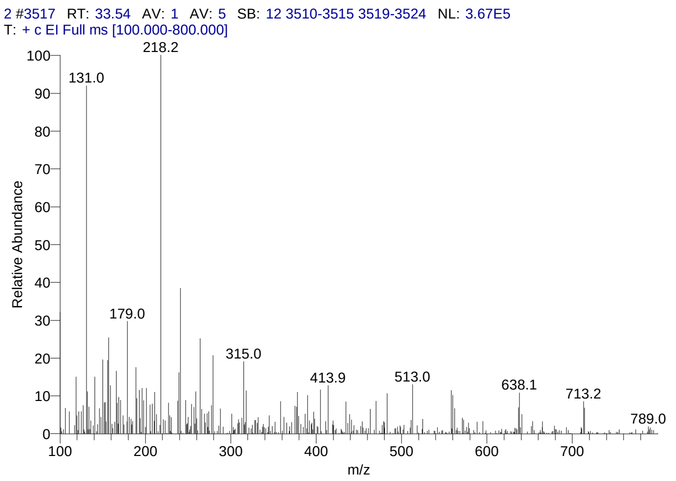


#### Library Search Results Table

| Compound Name | RT | Molecular Formula | Cas # |
|---|---|---|---|
| Rhodoxanthin | 33.54 | C40H50O2 | 116-30-3 |
| Pregn-4-ene-3,20-dione, 11,17,21-tris[(trimethylsilyl)oxy]-, bis(O-methyloxime), (11á)- | 33.54 | C32H60N2O5Si3 | 32221-26-4 |
| Benzene, hexakis(1-bromoethyl)- | 33.54 | C18H24Br6 | 87773-60-2 |

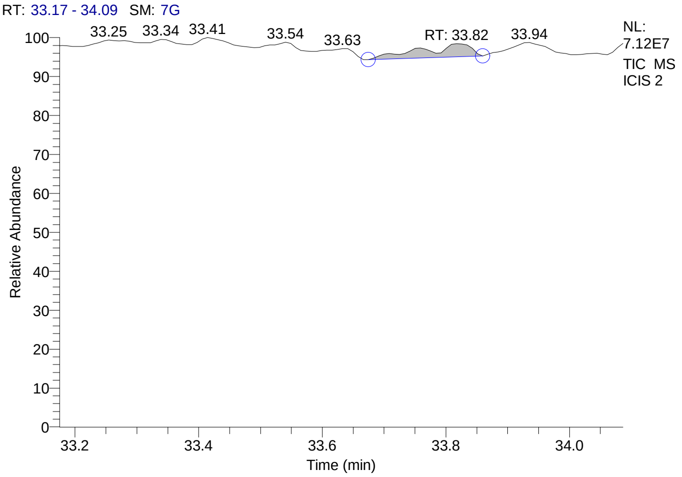


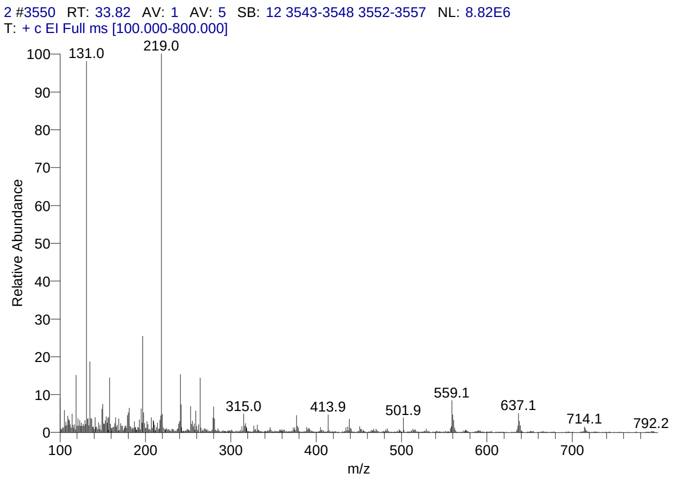


#### Library Search Results Table

| Compound Name | RT | Molecular Formula | Cas # |
|---|---|---|---|
| Rhodoxanthin | 33.82 | C40H50O2 | 116-30-3 |
| 5á-Cholestane-3à,7à,12à,24,25,26-hexol hexa-TMS | 33.82 | C45H96O6Si6 | NA |
| Olean-12-ene-3,16,21,22,23,28-hexol, (3á,4à,16à,21á,22à)- | 33.82 | C30H50O6 | 13844-22-9 |

**Supplementary Figure 6. GC-MS profile of butanolic extract of Garlic and the list of compound hits**

CIL/ SAIF Panjab University Chandigarh

#### Sample Header

| | |
|---|---|
| Data File: | SAMPLE-2 |
| Original Data Path: | C:\GCMS-DATA\YEAR 2016\DEC\16 |
| Sample Type: | Unknown |
| Sample ID: | 1 |
| Sample Name: | |
| Acquisition Date: | 12/16/16 01:18:47 PM |
| Run Time(min): | 30.09 |
| Injection Volume(µl): | 1.00 |
| Scans: | 3582 |
| Low Mass(m/z): | 50 |
| High Mass(m/z): | 700 |
| Instrument Method: | C:\GCMS-data\instrument method\GERNAL-gcms-METHOD.meth |

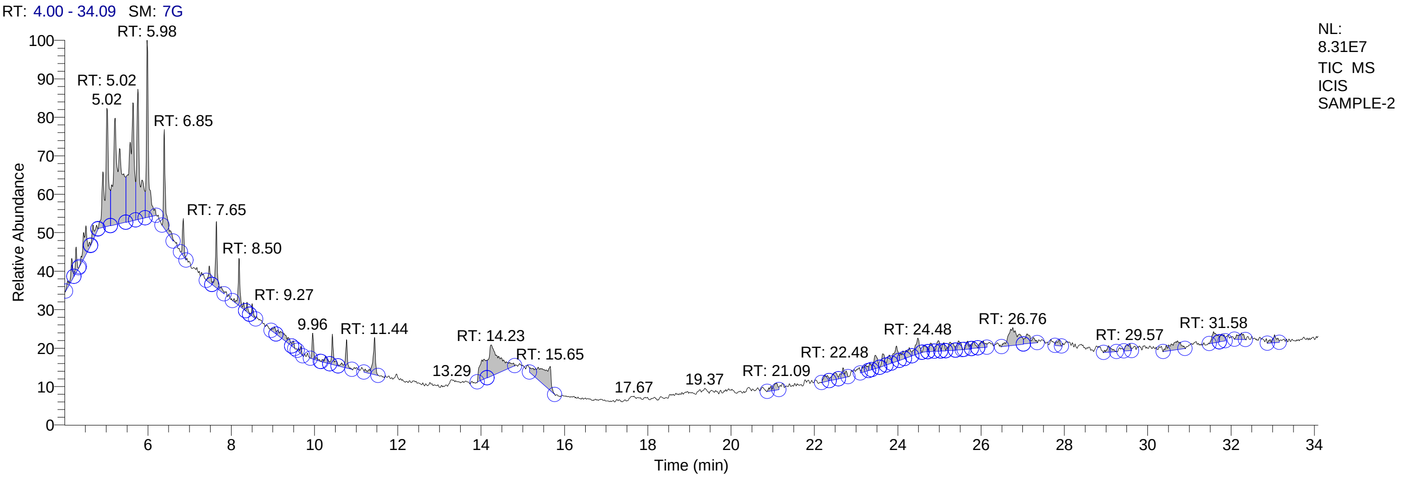


#### Qual Peak Table

| RT | Peak Area | Area % | Peak Height |
|---|---|---|---|
| 4.17 | 18654330.92 | 0.88 | 4690427.97 |
| 4.28 | 14738338.61 | 0.70 | 5597342.82 |
| 4.51 | 41828527.36 | 1.98 | 6199636.10 |
| 4.68 | 14876708.51 | 0.70 | 3306623.68 |
| 5.02 | 155841872.22 | 7.38 | 25497667.45 |
| 5.21 | 275705405.08 | 13.05 | 23017492.03 |
| 5.64 | 201590076.78 | 9.54 | 25408803.03 |
| 5.76 | 156618104.54 | 7.41 | 28139903.11 |
| 5.98 | 138129204.94 | 6.54 | 38244812.92 |
| 6.40 | 74805655.08 | 3.54 | 21342231.25 |
| 6.85 | 21356503.76 | 1.01 | 8091121.41 |
| 7.48 | 12960890.41 | 0.61 | 3606487.23 |
| 7.65 | 47181485.21 | 2.23 | 14333824.50 |
| 8.19 | 34830613.54 | 1.65 | 10219507.93 |
| 8.38 | 4923068.05 | 0.23 | 2085198.00 |
| 8.50 | 7758393.10 | 0.37 | 2638316.50 |
| 9.04 | 5354980.93 | 0.25 | 1336405.29 |
| 9.27 | 24225206.22 | 1.15 | 1421516.39 |
| 9.49 | 3892135.31 | 0.18 | 1378088.33 |
| 9.67 | 4943546.38 | 0.23 | 1571358.39 |
| 9.96 | 16686261.95 | 0.79 | 5570592.50 |
| 10.27 | 8275354.13 | 0.39 | 1395993.11 |
| 10.43 | 20867115.81 | 0.99 | 6543851.86 |
| 10.76 | 22281839.20 | 1.05 | 6157323.87 |
| 11.44 | 37680140.86 | 1.78 | 8056250.17 |
| 14.02 | 48859047.87 | 2.31 | 4359629.58 |
| 14.23 | 117896911.67 | 5.58 | 6721762.46 |
| 15.65 | 89564787.30 | 4.24 | 5202815.00 |

### CIL/ SAIF Panjab University Chandigarh

| RT | Peak Area | Area % | Peak Height |
|---|---|---|---|
| 21.09 | 14218529.34 | 0.67 | 1488437.88 |
| 22.26 | 10299705.20 | 0.49 | 1514971.71 |
| 22.48 | 13292703.72 | 0.63 | 1535488.81 |
| 22.69 | 14839703.14 | 0.70 | 2132461.22 |
| 23.22 | 12202606.99 | 0.58 | 1433750.15 |
| 23.32 | 5232442.65 | 0.25 | 1489429.89 |
| 23.46 | 19609476.51 | 0.93 | 2921430.48 |
| 23.64 | 13026932.90 | 0.62 | 2695068.99 |
| 23.79 | 10430362.98 | 0.49 | 1918498.00 |
| 23.96 | 23835788.59 | 1.13 | 3320121.61 |
| 24.12 | 9835456.26 | 0.47 | 1783289.93 |
| 24.27 | 14793081.98 | 0.70 | 1986237.26 |
| 24.48 | 24318959.94 | 1.15 | 3415770.63 |
| 24.66 | 7802411.36 | 0.37 | 1461945.55 |
| 24.79 | 10524759.92 | 0.50 | 1853833.50 |
| 24.98 | 12436750.66 | 0.59 | 2263942.76 |
| 25.11 | 7814512.17 | 0.37 | 1777012.01 |
| 25.26 | 15899563.90 | 0.75 | 1773077.74 |
| 25.46 | 12435521.92 | 0.59 | 1934122.49 |
| 25.68 | 9827429.54 | 0.47 | 1739196.34 |
| 25.81 | 7666288.74 | 0.36 | 1607253.52 |
| 26.05 | 12318053.92 | 0.58 | 1461469.64 |
| 26.76 | 63837310.07 | 3.02 | 3822169.67 |
| 27.11 | 22447863.90 | 1.06 | 2133776.32 |
| 27.88 | 6991590.90 | 0.33 | 1324475.72 |
| 29.06 | 10742796.27 | 0.51 | 1372026.86 |
| 29.57 | 11646727.81 | 0.55 | 1531248.36 |
| 30.72 | 34440627.10 | 1.63 | 1890793.53 |
| 31.58 | 21440862.98 | 1.01 | 2439429.43 |
| 31.78 | 10043541.49 | 0.48 | 1624306.77 |
| 32.25 | 13224673.88 | 0.63 | 1407962.24 |
| 33.04 | 12699333.57 | 0.60 | 1723518.75 |

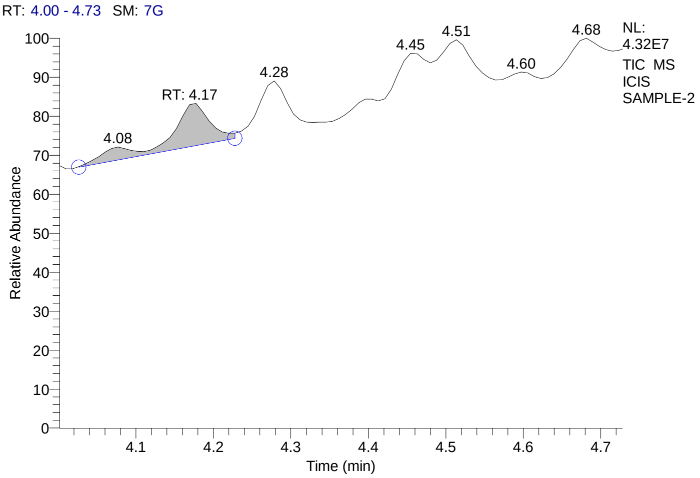


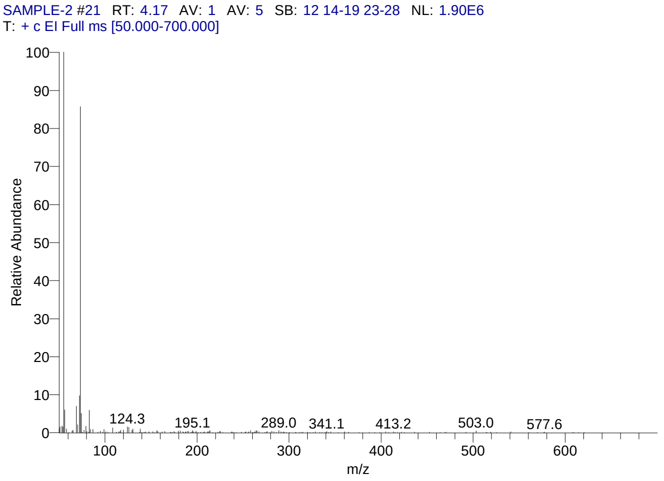


#### Library Search Results Table

| Compound Name | RT | Molecular Formula | Cas # |
|---|---|---|---|
| 3-Hexanol, 2-methyl- | 4.17 | C7H16O | 617-29-8 |
| 4-Heptanol | 4.17 | C7H16O | 589-55-9 |
| 3-Pentanol, 2,4-dimethyl- | 4.17 | C7H16O | 600-36-2 |

### CIL/ SAIF Panjab University Chandigarh

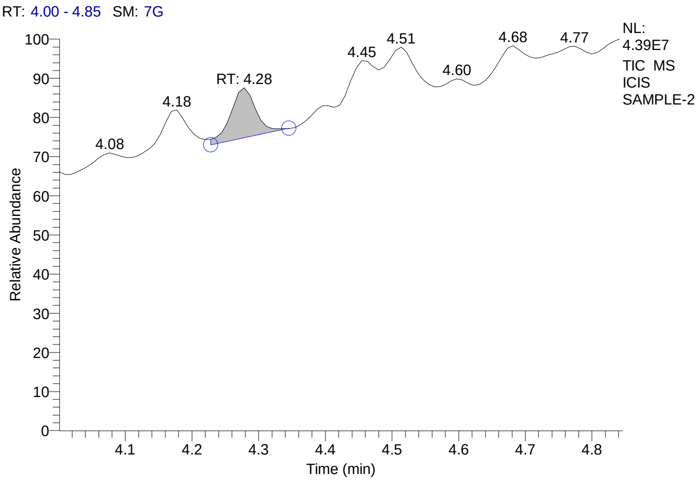


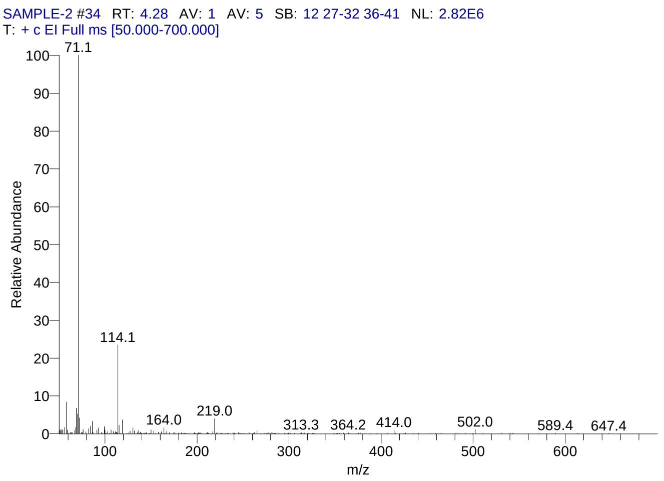


#### Library Search Results Table

| Compound Name | RT | Molecular Formula | Cas # |
|---|---|---|---|
| 3-Hexanone, 2-methyl- | 4.28 | C7H14O | 7379-12-6 |
| 3-Pentanone, 2,4-dimethyl- | 4.28 | C7H14O | 565-80-0 |
| 4-Heptanone | 4.28 | C7H14O | 123-19-3 |

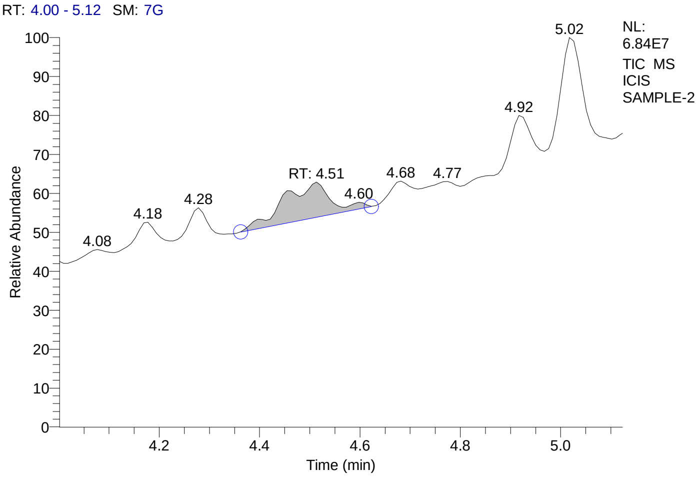


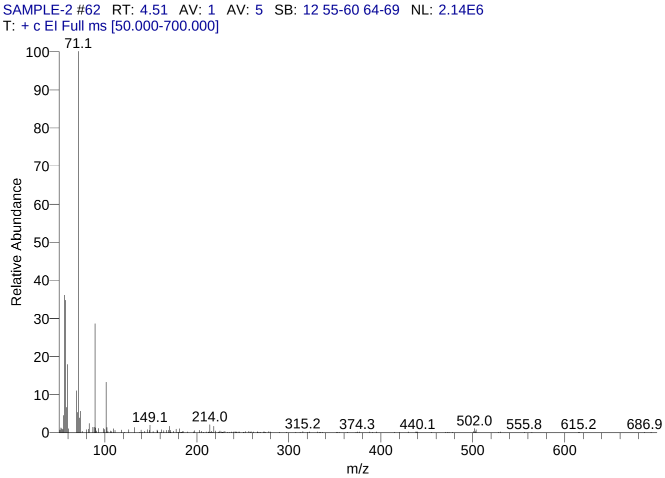


#### Library Search Results Table

| Compound Name | RT | Molecular Formula | Cas # |
|---|---|---|---|
| Propanoic acid, 2-methyl-, 2-methylpropyl ester | 4.51 | C8H16O2 | 97-85-8 |
| Butanoic acid, butyl ester | 4.51 | C8H16O2 | 109-21-7 |
| Propanoic acid, 2-methyl-, butyl ester | 4.51 | C8H16O2 | 97-87-0 |

### CIL/ SAIF Panjab University Chandigarh

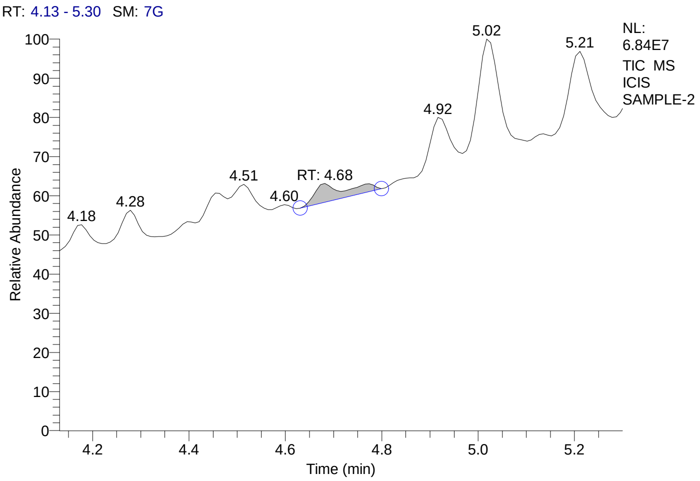


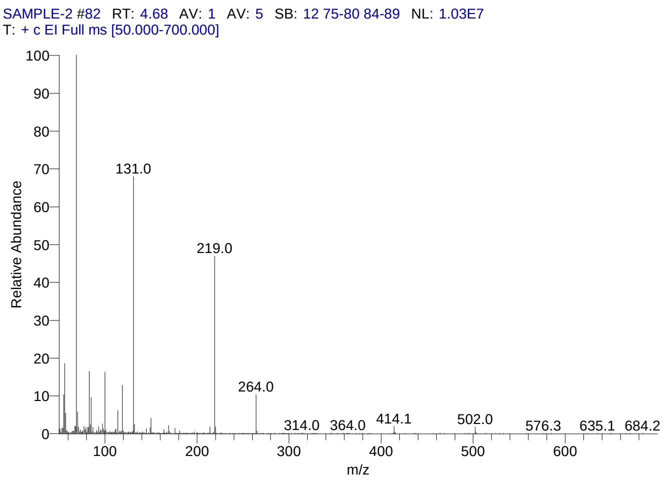


#### Library Search Results Table

| Compound Name | RT | Molecular Formula | Cas # |
|---|---|---|---|
| Perfluorotributylamine | 4.68 | C12F27N | 311-89-7 |
| Pyrimidin-2-one, 4-[N-methylureido]-1-[4-methylaminocarbonyloxy methyl | 4.68 | C13H19N5O5 | NA |
| Perfluoro(dibutylmethylamine) | 4.68 | C9F21N | 514-03-4 |

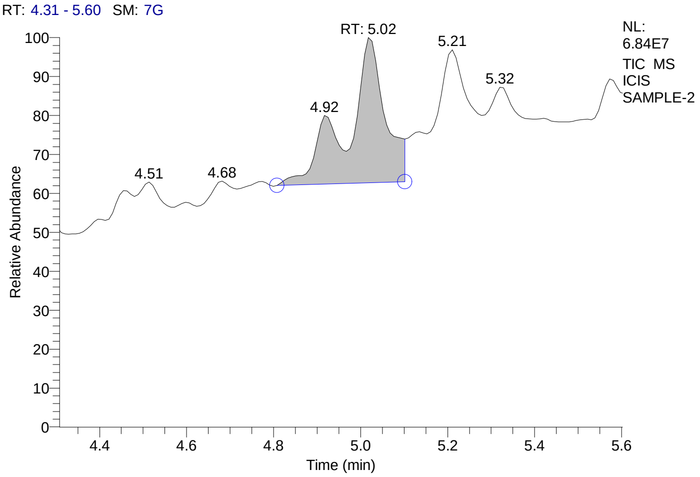


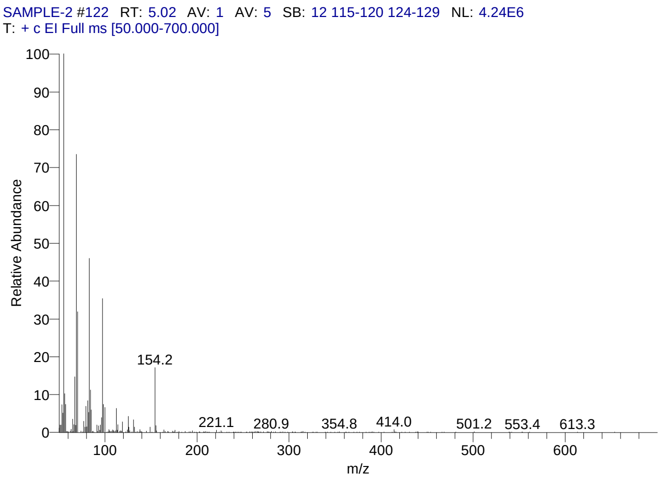


#### Library Search Results Table

| Compound Name | RT | Molecular Formula | Cas # |
|---|---|---|---|
| 1-Decene, 5-methyl- | 5.02 | C11H22 | 54244-79-0 |
| Cyclopropane, 1-butyl-1-methyl-2-propyl- | 5.02 | C11H22 | 41977-34-8 |
| 3-Methyldec-3-ene | 5.02 | C11H22 | 36969-75-2 |

### CIL/ SAIF Panjab University Chandigarh

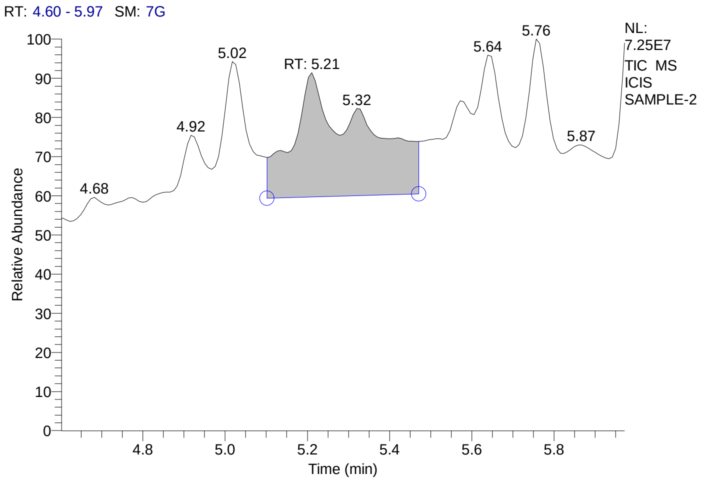


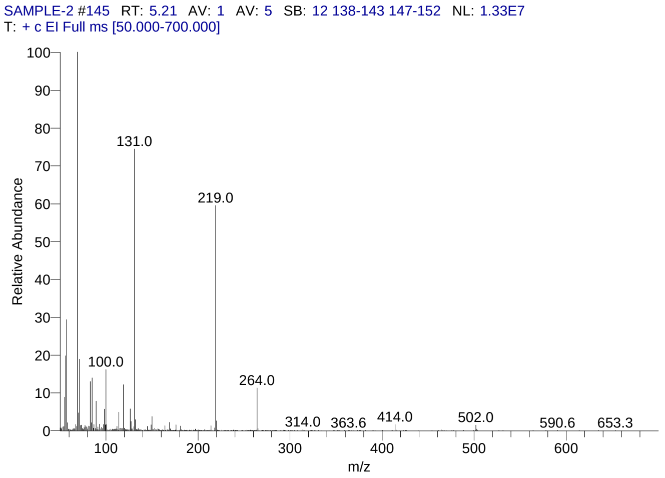


#### Library Search Results Table

| Compound Name | RT | Molecular Formula | Cas # |
|---|---|---|---|
| Perfluorotributylamine | 5.21 | C12F27N | 311-89-7 |
| Pyrimidin-2-one, 4-[N-methylureido]-1-[4-methylaminocarbonyloxy methyl | 5.21 | C13H19N5O5 | NA |
| Perfluoro(dibutylmethylamine) | 5.21 | C9F21N | 514-03-4 |

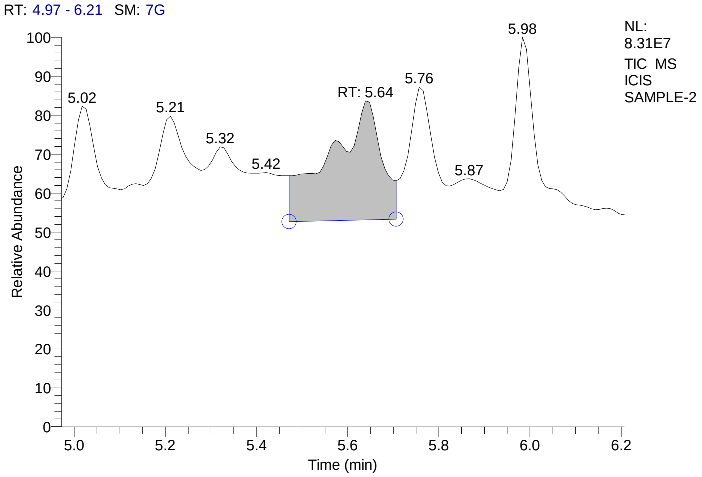


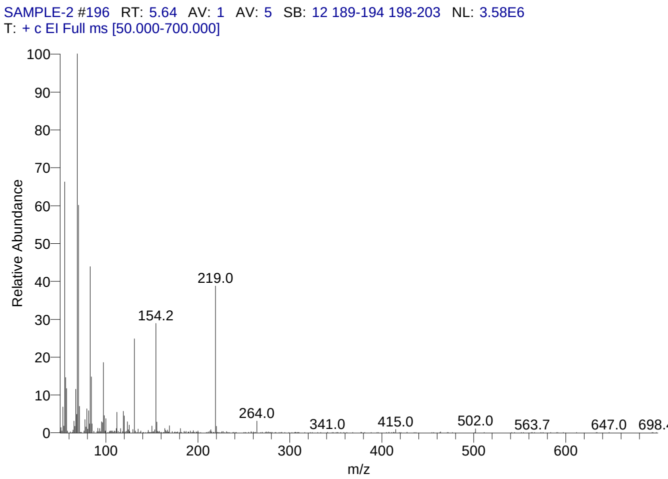


#### Library Search Results Table

| Compound Name | RT | Molecular Formula | Cas # |
|---|---|---|---|
| 4-Decene, 3-methyl-, (E)- | 5.64 | C11H22 | 62338-47-0 |
| 2-Octene, 2,3,7-trimethyl- | 5.64 | C11H22 | 33933-75-4 |
| 1-Decene, 5-methyl- | 5.64 | C11H22 | 54244-79-0 |

### CIL/ SAIF Panjab University Chandigarh

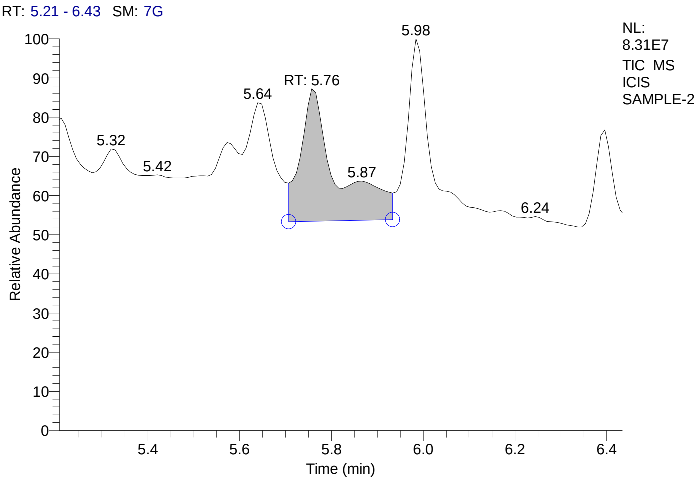


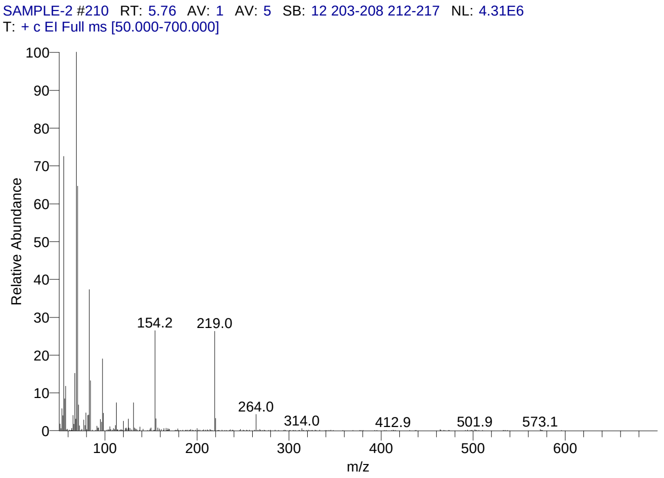


### Library Search Results Table

| Compound Name | RT | Molecular Formula | Cas # |
|---|---|---|---|
| 4-Decene, 3-methyl-, (E)- | 5.76 | C11H22 | 62338-47-0 |
| 2-Decene, 8-methyl-, (Z)- | 5.76 | C11H22 | 74630-25-4 |
| 3-Methyldec-3-ene | 5.76 | C11H22 | 36969-75-2 |

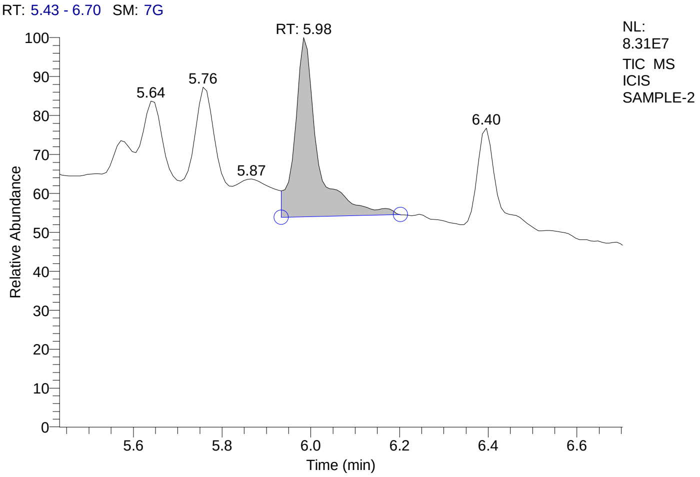


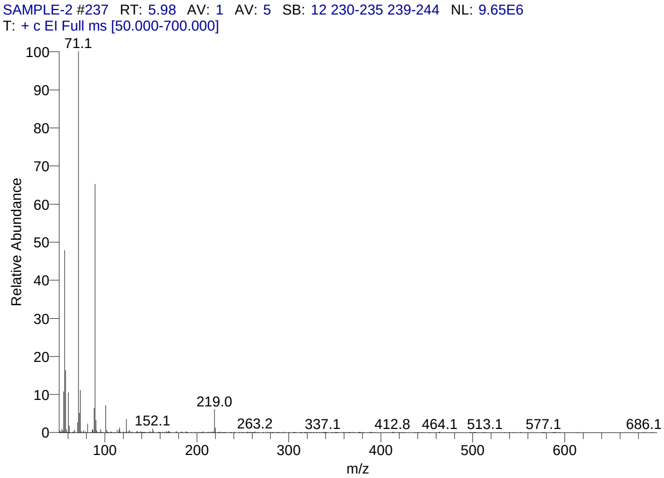


### Library Search Results Table

| Compound Name | RT | Molecular Formula | Cas # |
|---|---|---|---|
| Butanoic acid, butyl ester | 5.98 | C8H16O2 | 109-21-7 |
| Butanoic acid, 2-methylpropyl ester | 5.98 | C8H16O2 | 539-90-2 |
| Propanoic acid, 2-methyl-, 2-methylpropyl ester | 5.98 | C8H16O2 | 97-85-8 |

### CIL/ SAIF Panjab University Chandigarh

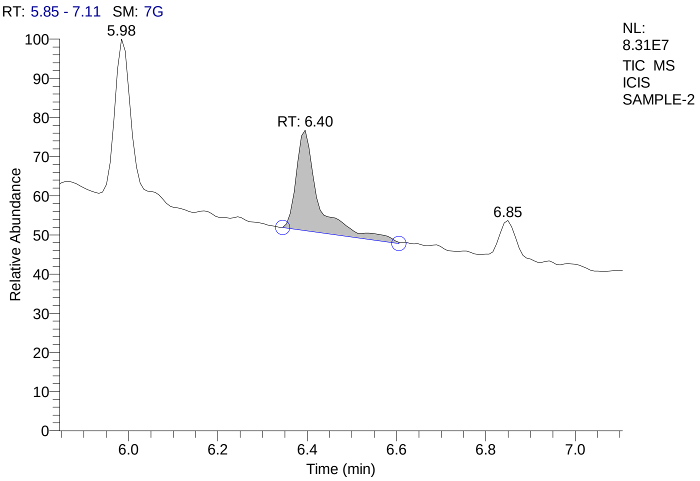


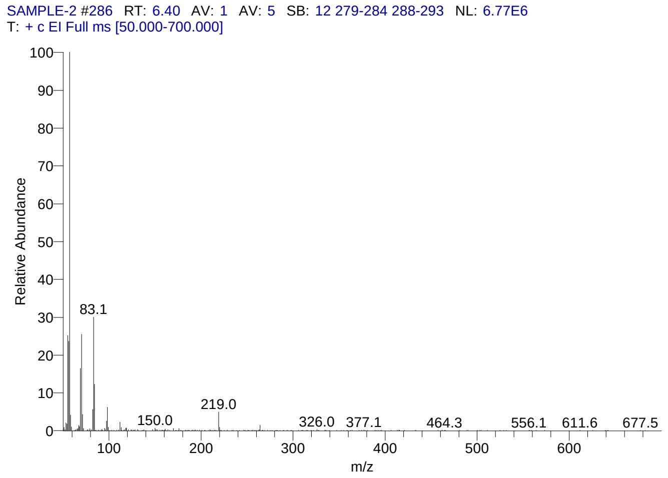


#### Library Search Results Table

| Compound Name | RT | Molecular Formula | Cas # |
|---|---|---|---|
| 2-Propyl-1-pentanol | 6.40 | C8H18O | 58175-57-8 |
| 1-Hexanol, 2-ethyl- | 6.40 | C8H18O | 104-76-7 |
| Formic acid, 2-ethylhexyl ester | 6.40 | C9H18O2 | NA |

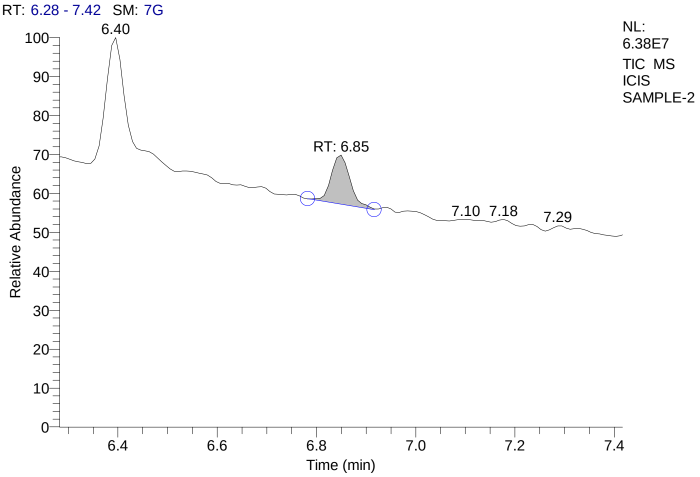


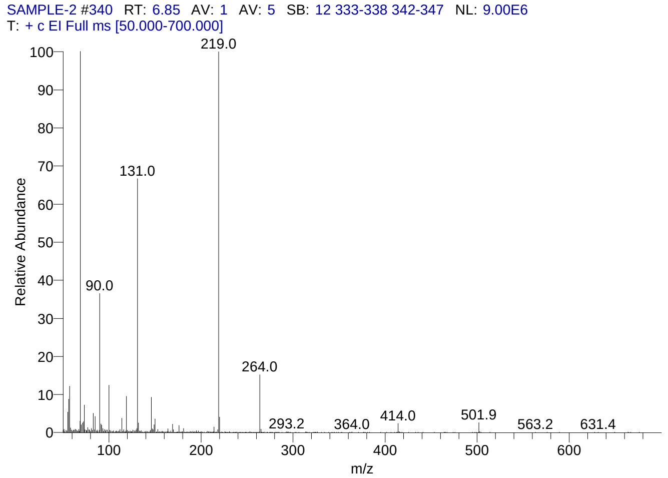


#### Library Search Results Table

| Compound Name | RT | Molecular Formula | Cas # |
|---|---|---|---|
| Perfluorotributylamine | 6.85 | C12F27N | 311-89-7 |
| Perfluoro(dibutylmethylamine) | 6.85 | C9F21N | 514-03-4 |
| Nonafluoro-1-bromobutane | 6.85 | C4BrF9 | 375-48-4 |

### CIL/ SAIF Panjab University Chandigarh

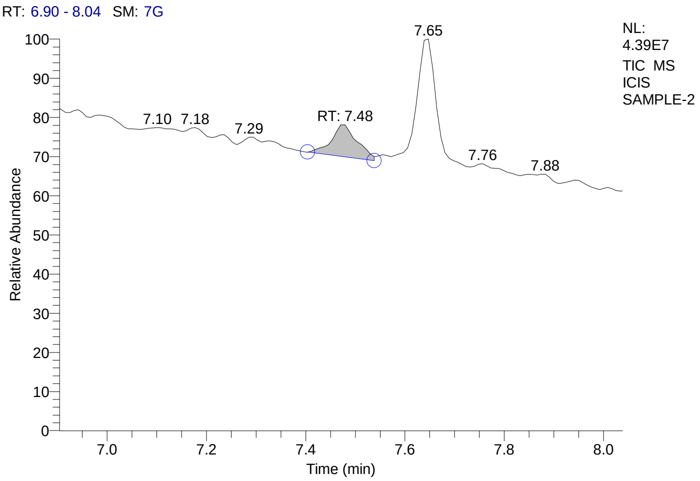


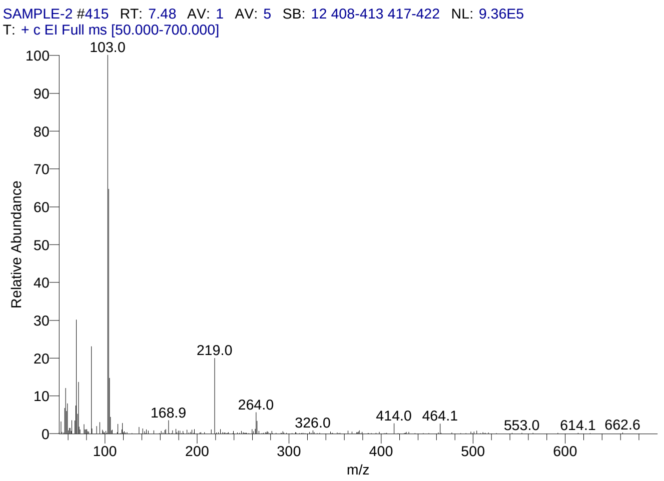


#### Library Search Results Table

| Compound Name | RT | Molecular Formula | Cas # |
|---|---|---|---|
| 10-Chlorodecyl 2-methylbutanoate | 7.48 | C15H29ClO2 | NA |
| Isovaleric acid, 8-chlorooctyl ester | 7.48 | C13H25ClO2 | NA |
| 8-Chlorooctyl 2-methylbutanoate | 7.48 | C13H25ClO2 | NA |

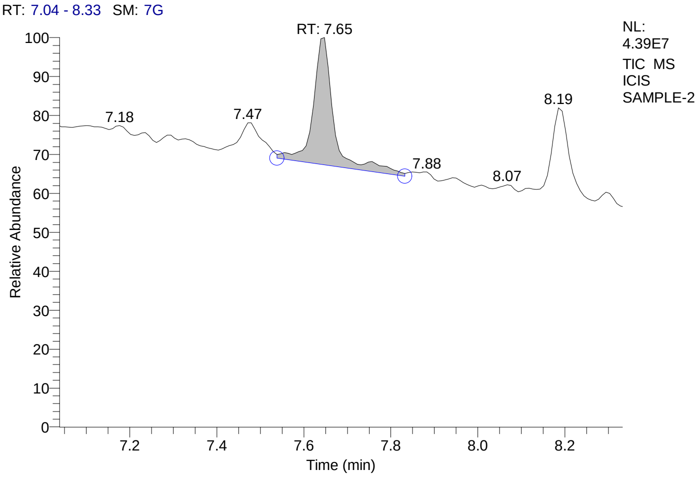


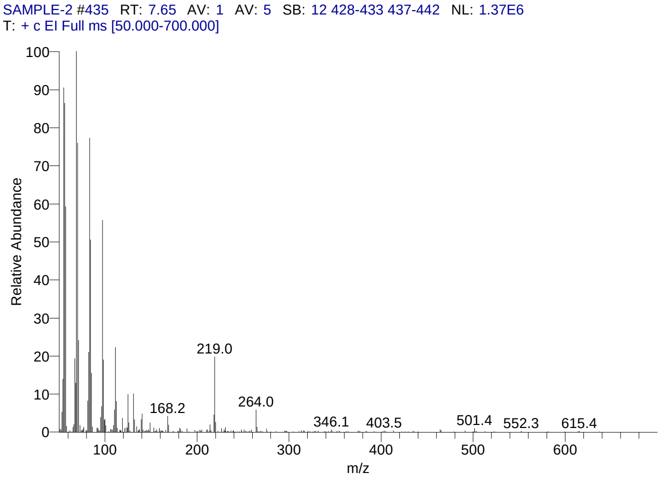


#### Library Search Results Table

| Compound Name | RT | Molecular Formula | Cas # |
|---|---|---|---|
| Cyclopropane, 1-methyl-2-octyl- | 7.65 | C12H24 | 37617-26-8 |
| Cyclopropane, 1-butyl-2-pentyl-, cis- | 7.65 | C12H24 | 74663-88-0 |
| 3-Tetradecene, (Z)- | 7.65 | C14H28 | 41446-67-7 |

### CIL/ SAIF Panjab University Chandigarh

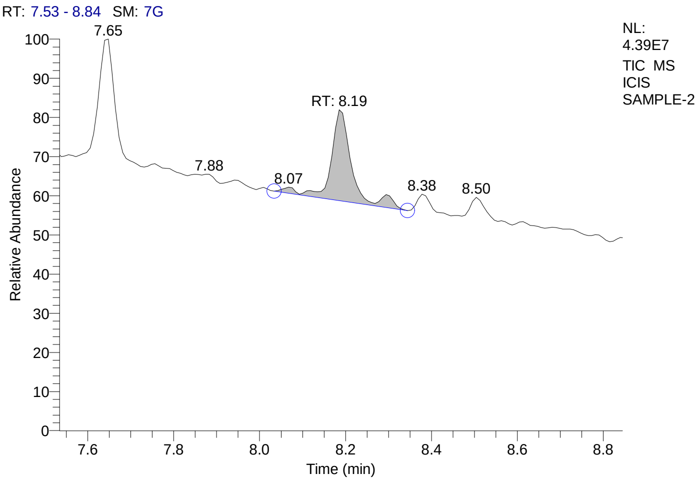


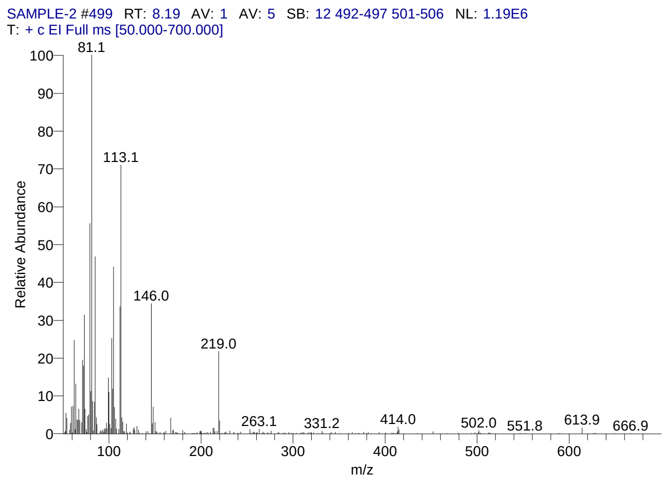


#### Library Search Results Table

| Compound Name | RT | Molecular Formula | Cas # |
|---|---|---|---|
| Diallyl disulphide | 8.19 | C6H10S2 | 2179-57-9 |
| Tetrasulfide, di-2-propenyl | 8.19 | C6H10S4 | 2444-49-7 |
| 2-Oxa-7-thiatricyclo[4.4.0.0(3,8)]decane | 8.19 | C8H12OS | 39637-14-4 |

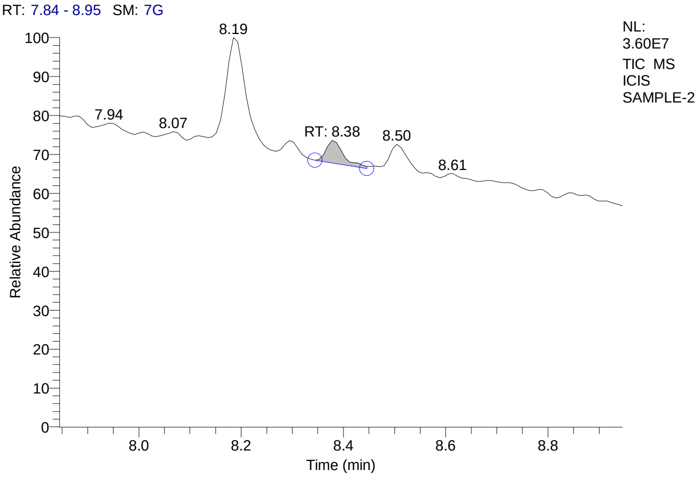


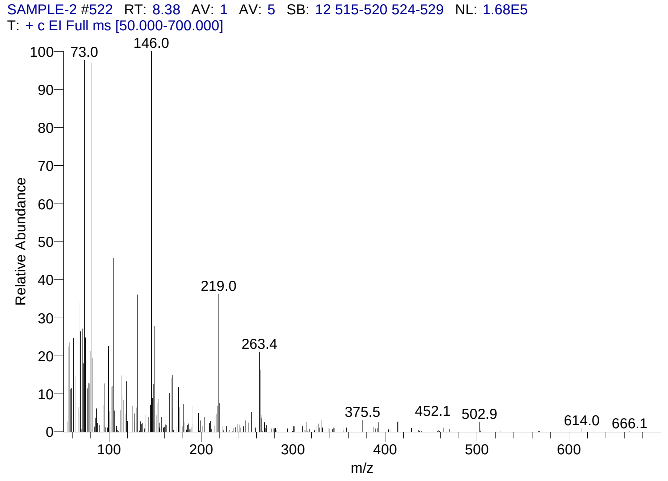


#### Library Search Results Table

| Compound Name | RT | Molecular Formula | Cas # |
|---|---|---|---|
| Tetrasulfide, di-2-propenyl | 8.38 | C6H10S4 | 2444-49-7 |
| Diallyl disulphide | 8.38 | C6H10S2 | 2179-57-9 |
| 9,10-Secocholesta-5,7,10(19)-triene-1,3-diol, 25-[(trimethylsilyl)oxy]-, (3á,5Z,7E)- | 8.38 | C30H52O3Si | 55759-94-9 |

### CIL/ SAIF Panjab University Chandigarh

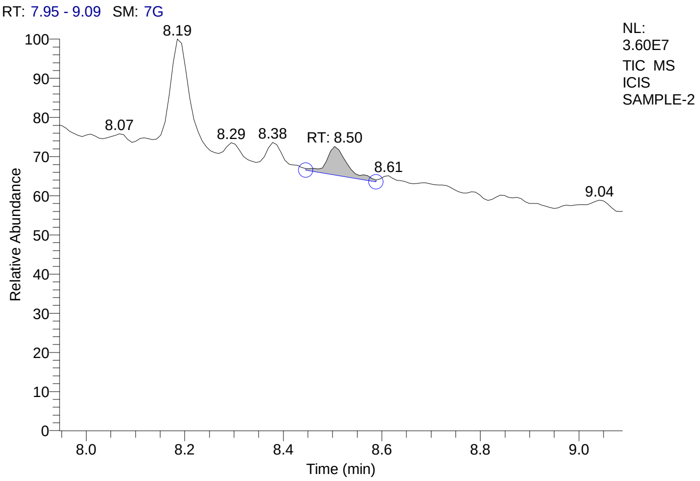


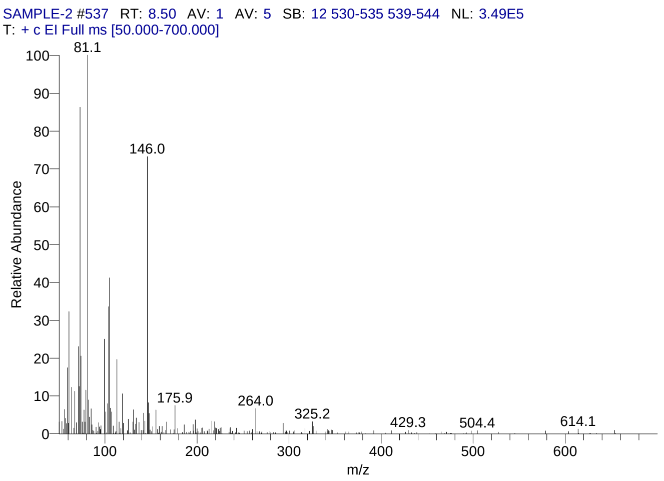


#### Library Search Results Table

| Compound Name | RT | Molecular Formula | Cas # |
|---|---|---|---|
| Diallyl disulphide | 8.50 | C6H10S2 | 2179-57-9 |
| Tetrasulfide, di-2-propenyl | 8.50 | C6H10S4 | 2444-49-7 |
| 2-Vinyl-1,3-dithiane | 8.50 | C6H10S2 | 61685-40-3 |

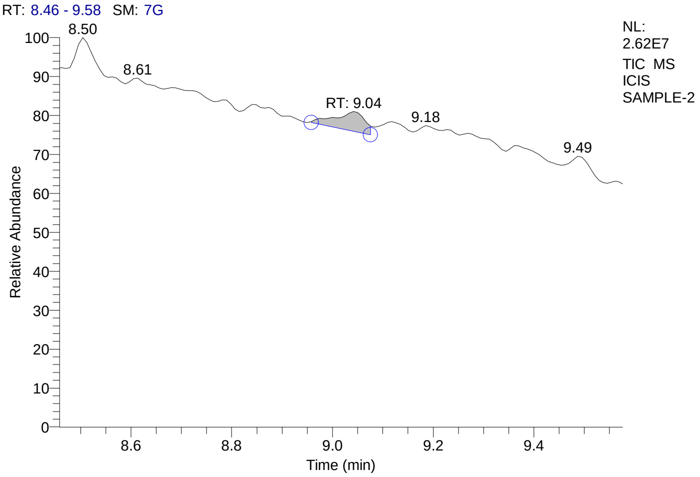


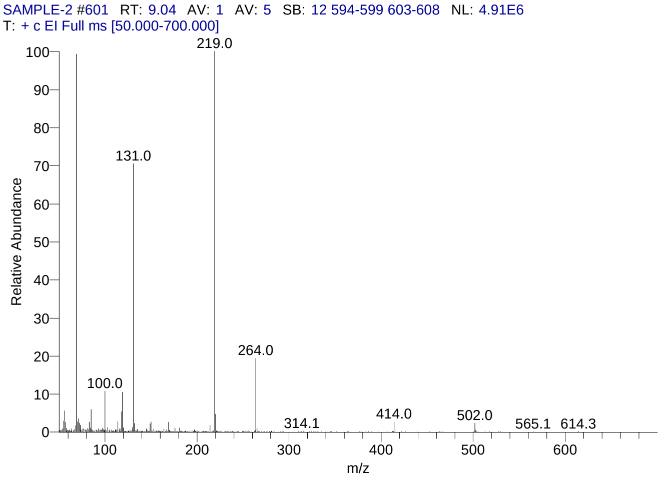


#### Library Search Results Table

| Compound Name | RT | Molecular Formula | Cas # |
|---|---|---|---|
| Perfluorotributylamine | 9.04 | C12F27N | 311-89-7 |
| Perfluoro(dibutylmethylamine) | 9.04 | C9F21N | 514-03-4 |
| Nonafluoro-1-bromobutane | 9.04 | C4BrF9 | 375-48-4 |

### CIL/ SAIF Panjab University Chandigarh

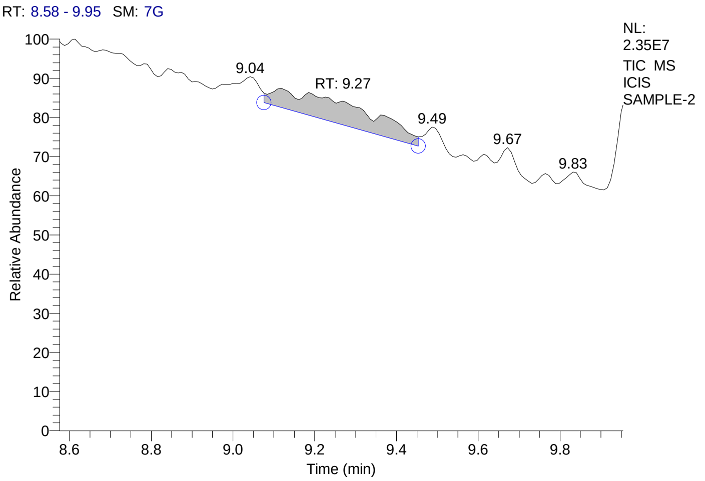


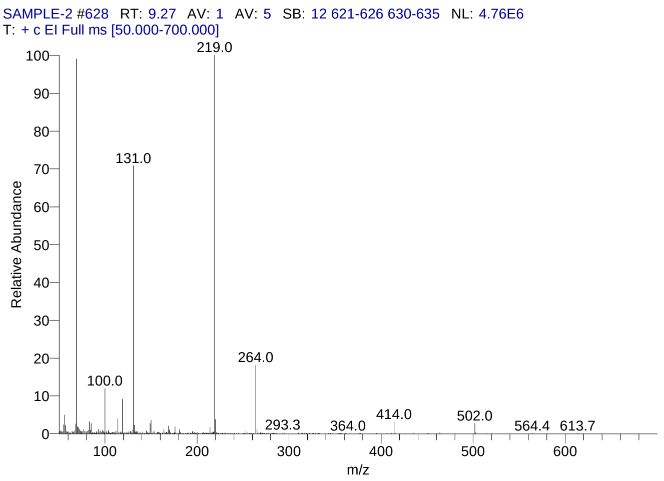


#### Library Search Results Table

| Compound Name | RT | Molecular Formula | Cas # |
|---|---|---|---|
| Perfluorotributylamine | 9.27 | C12F27N | 311-89-7 |
| Perfluoro(dibutylmethylamine) | 9.27 | C9F21N | 514-03-4 |
| Nonafluoro-1-bromobutane | 9.27 | C4BrF9 | 375-48-4 |

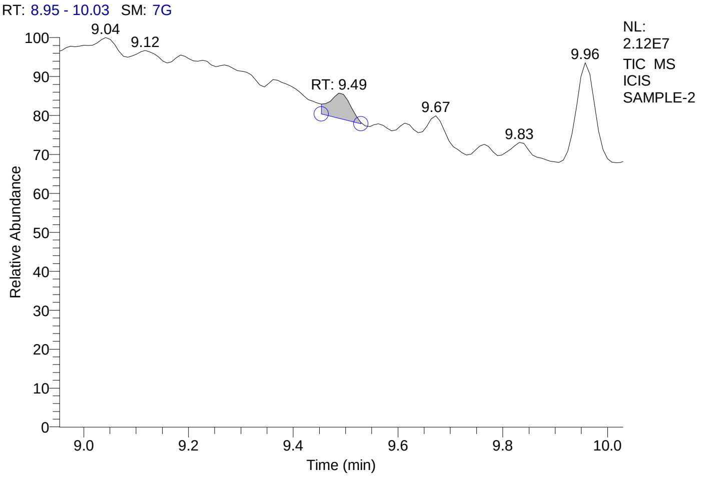


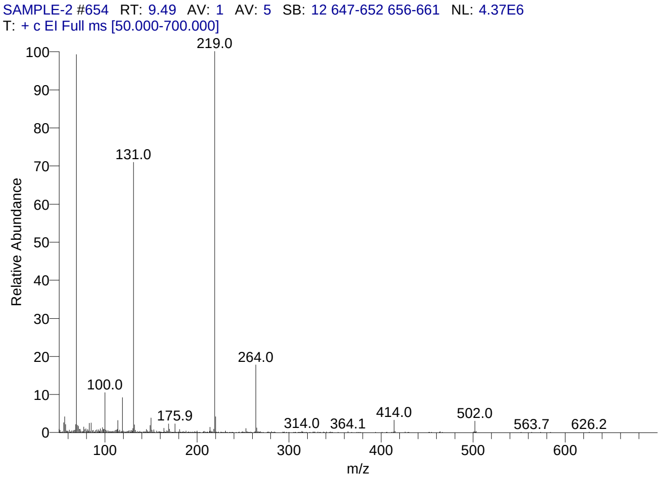


#### Library Search Results Table

| Compound Name | RT | Molecular Formula | Cas # |
|---|---|---|---|
| Perfluorotributylamine | 9.49 | C12F27N | 311-89-7 |
| Perfluoro(dibutylmethylamine) | 9.49 | C9F21N | 514-03-4 |
| Nonafluoro-1-bromobutane | 9.49 | C4BrF9 | 375-48-4 |

### CIL/ SAIF Panjab University Chandigarh

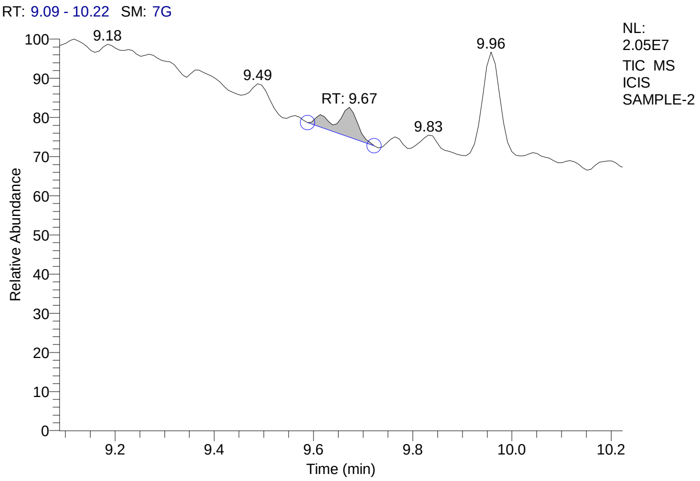


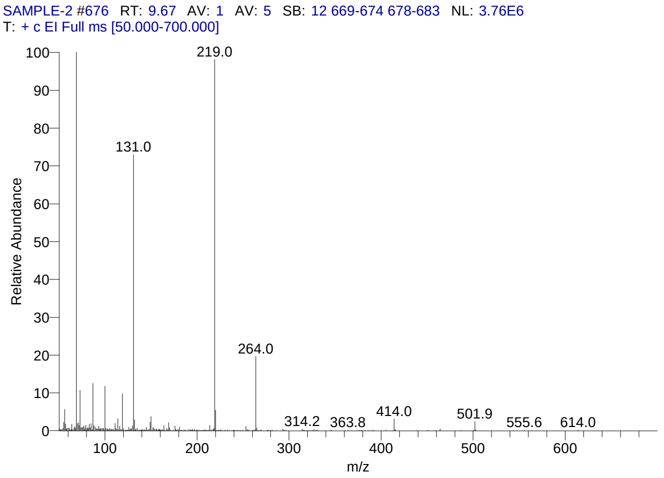


#### Library Search Results Table

| Compound Name | RT | Molecular Formula | Cas # |
|---|---|---|---|
| Perfluorotributylamine | 9.67 | C12F27N | 311-89-7 |
| Perfluoro(dibutylmethylamine) | 9.67 | C9F21N | 514-03-4 |
| Nonafluoro-1-bromobutane | 9.67 | C4BrF9 | 375-48-4 |

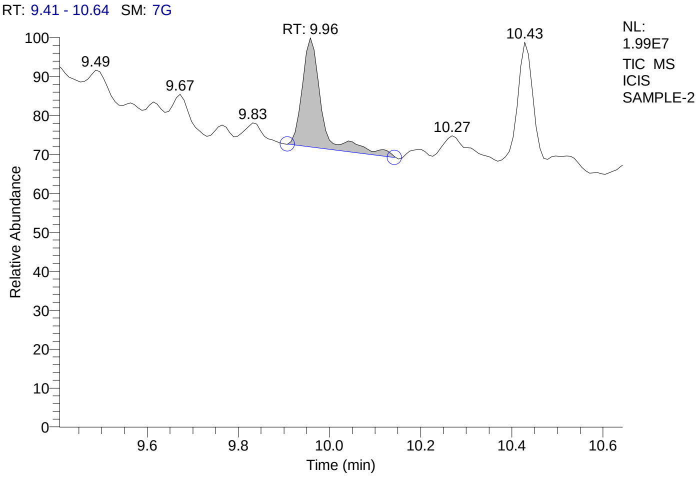


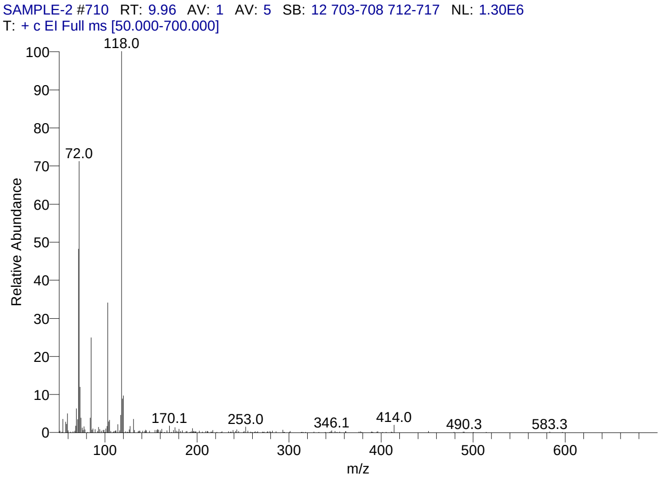


#### Library Search Results Table

| Compound Name | RT | Molecular Formula | Cas # |
|---|---|---|---|
| 1,3-Oxathiane, 2-methyl- | 9.96 | C5H10OS | 19134-37-3 |
| Butanethioic acid, S-methyl ester | 9.96 | C5H10OS | 2432-51-1 |
| 2-Phenylpropyl isobutyrate | 9.96 | C13H18O2 | 65813-53-8 |

### CIL/ SAIF Panjab University Chandigarh

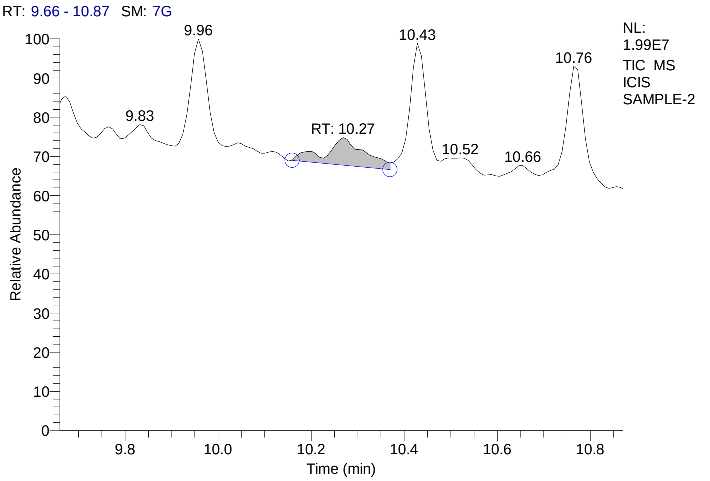


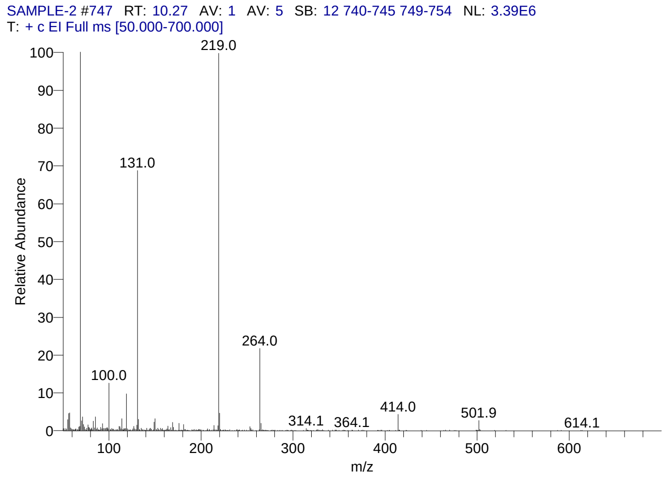


#### Library Search Results Table

| Compound Name | RT | Molecular Formula | Cas # |
|---|---|---|---|
| Perfluorotributylamine | 10.27 | C12F27N | 311-89-7 |
| Perfluoro(dibutylmethylamine) | 10.27 | C9F21N | 514-03-4 |
| Nonafluoro-1-bromobutane | 10.27 | C4BrF9 | 375-48-4 |

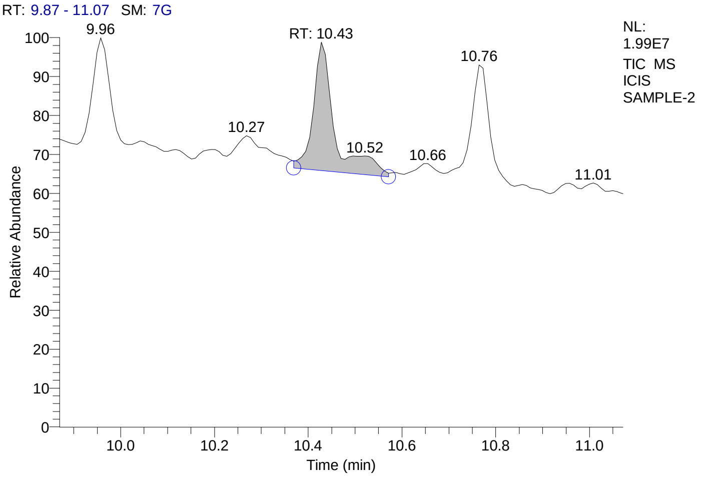


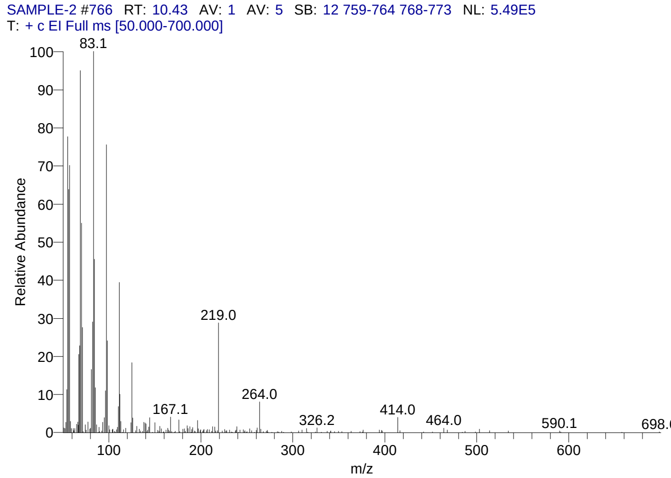


#### Library Search Results Table

| Compound Name | RT | Molecular Formula | Cas # |
|---|---|---|---|
| 1-Hexadecanol | 10.43 | C16H34O | 36653-82-4 |
| 2-Dodecanol | 10.43 | C12H26O | 10203-28-8 |
| Cyclotridecane | 10.43 | C13H26 | 295-02-3 |

### CIL/ SAIF Panjab University Chandigarh

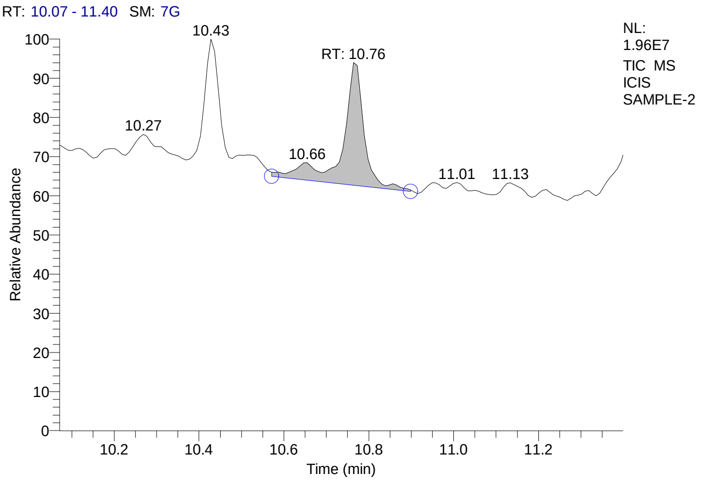


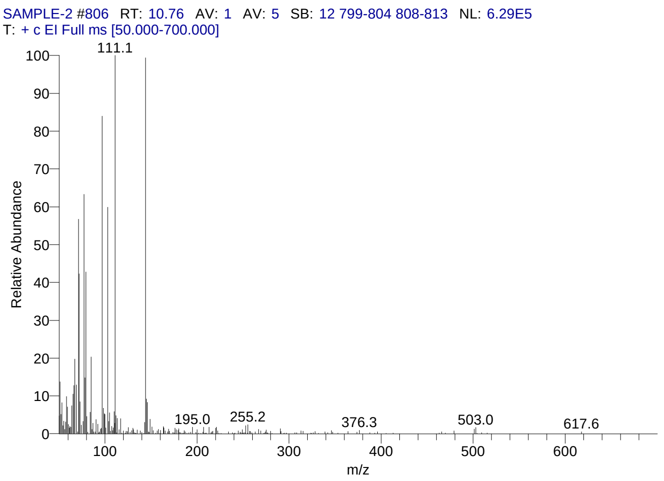


#### Library Search Results Table

| Compound Name | RT | Molecular Formula | Cas # |
|---|---|---|---|
| 3-Vinyl-1,2-dithiacyclohex-4-ene | 10.76 | C6H8S2 | 62488-52-2 |
| 3-Vinyl-1,2-dithiacyclohex-5-ene | 10.76 | C6H8S2 | 62488-53-3 |
| 1,2-Dithiol-1-ium, 3,4,5-trimethyl-, bromide | 10.76 | C6H9BrS2 | 55836-86-7 |

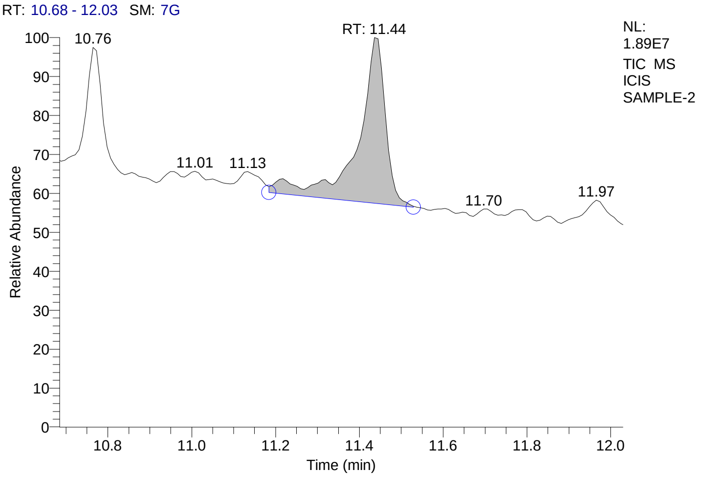


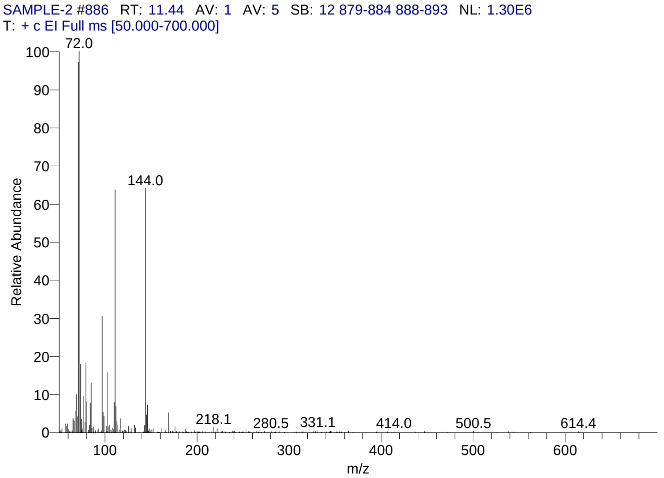


#### Library Search Results Table

| Compound Name | RT | Molecular Formula | Cas # |
|---|---|---|---|
| 3-Vinyl-1,2-dithiacyclohex-5-ene | 11.44 | C6H8S2 | 62488-53-3 |
| 3-Vinyl-1,2-dithiacyclohex-4-ene | 11.44 | C6H8S2 | 62488-52-2 |
| 1,2-Dithiol-1-ium, 3,4,5-trimethyl-, bromide | 11.44 | C6H9BrS2 | 55836-86-7 |

### CIL/ SAIF Panjab University Chandigarh

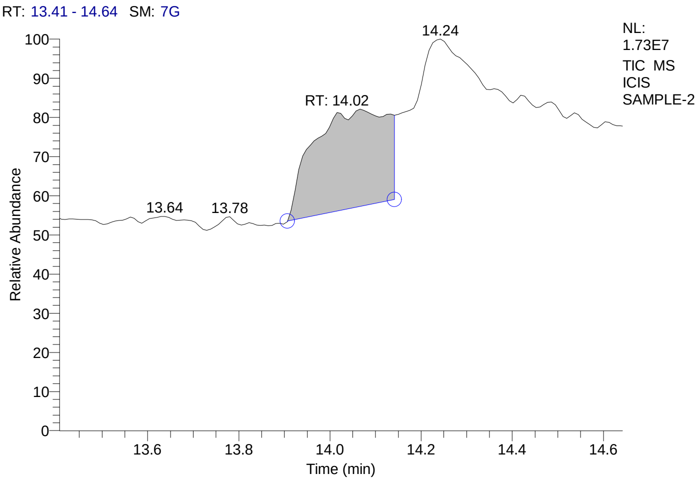


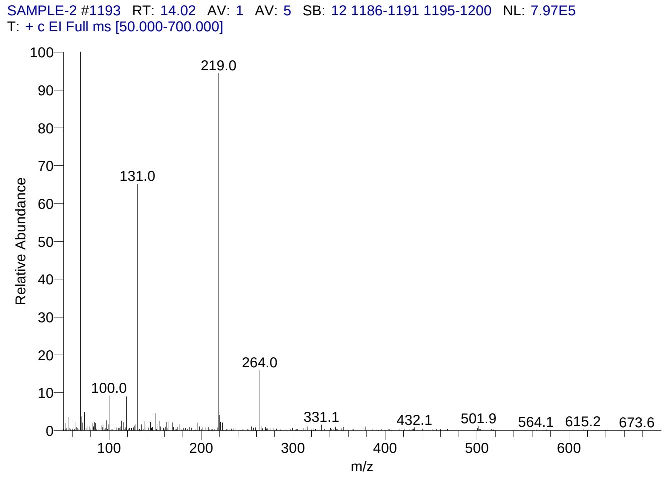


#### Library Search Results Table

| Compound Name | RT | Molecular Formula | Cas # |
|---|---|---|---|
| Perfluorotributylamine | 14.02 | C12F27N | 311-89-7 |
| Nonafluoro-1-bromobutane | 14.02 | C4BrF9 | 375-48-4 |
| Perfluoro(dibutylmethylamine) | 14.02 | C9F21N | 514-03-4 |

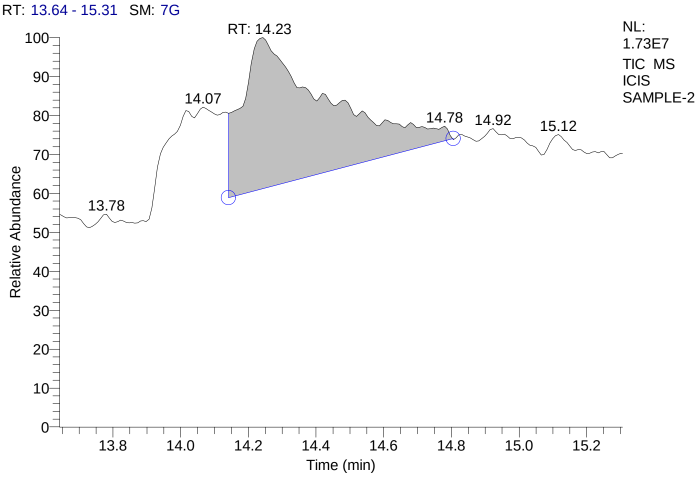


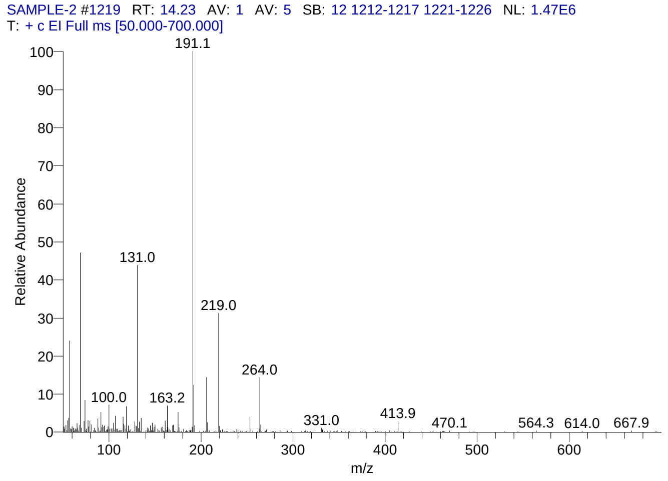


#### Library Search Results Table

| Compound Name | RT | Molecular Formula | Cas # |
|---|---|---|---|
| Phenol, 2,6-bis(1,1-dimethylethyl)- | 14.23 | C14H22O | 128-39-2 |
| Phenol, 2,4-bis(1,1-dimethylethyl)- | 14.23 | C14H22O | 96-76-4 |
| Phenol, 3,5-bis(1,1-dimethylethyl)- | 14.23 | C14H22O | 1138-52-9 |

### CIL/ SAIF Panjab University Chandigarh

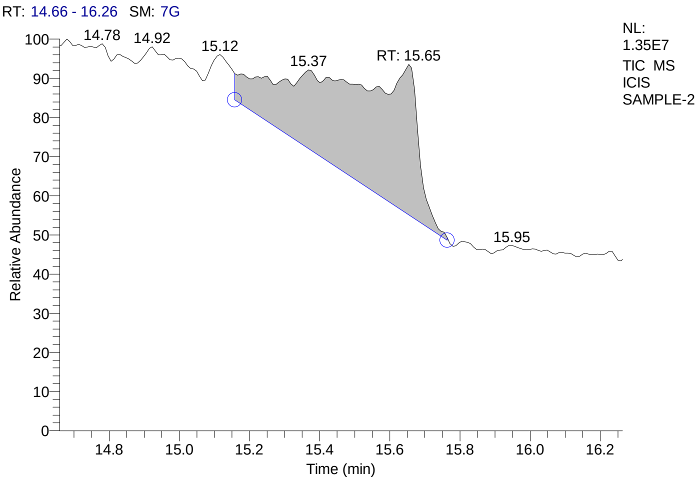


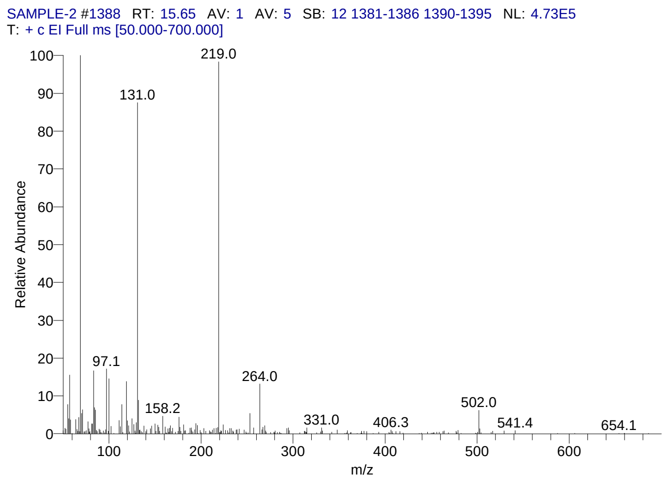


#### Library Search Results Table

| Compound Name | RT | Molecular Formula | Cas # |
|---|---|---|---|
| Perfluorotributylamine | 15.65 | C12F27N | 311-89-7 |
| Pyrimidin-2-one, 4-[N-methylureido]-1-[4-methylaminocarbonyloxy methyl | 15.65 | C13H19N5O5 | NA |
| 5á-Cholestane-3à,7à,12à,24,25,26-hexol hexa-TMS | 15.65 | C45H96O6Si6 | NA |

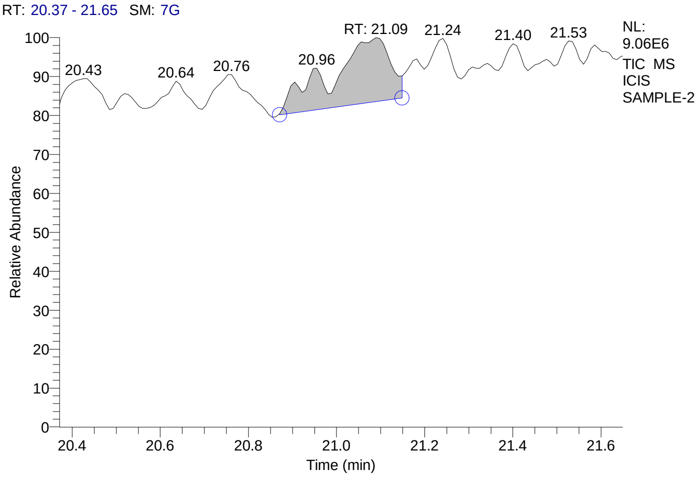


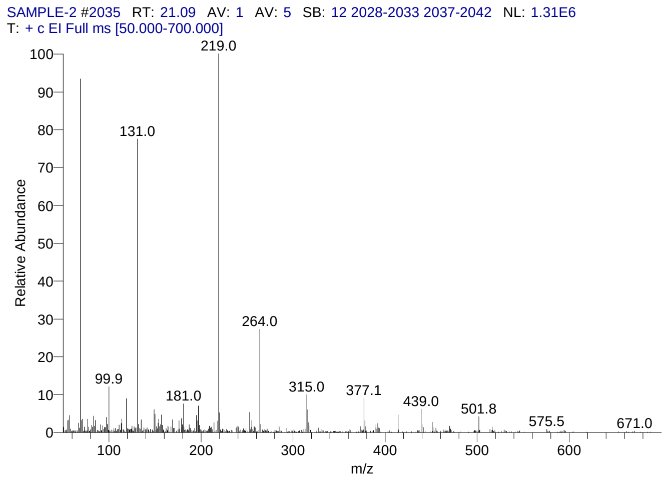


#### Library Search Results Table

| Compound Name | RT | Molecular Formula | Cas # |
|---|---|---|---|
| Perfluorotributylamine | 21.09 | C12F27N | 311-89-7 |
| Oleic acid, eicosyl ester | 21.09 | C38H74O2 | 22393-88-0 |
| 5á-Cholestane-3à,7à,12à,24,25,26-hexol hexa-TMS | 21.09 | C45H96O6Si6 | NA |

### CIL/ SAIF Panjab University Chandigarh

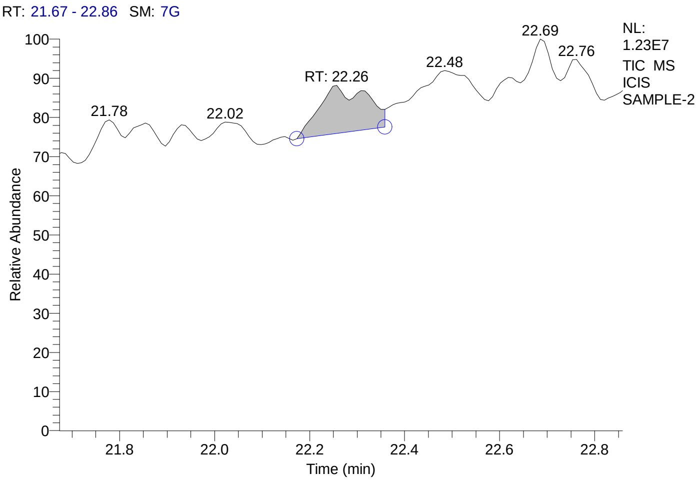


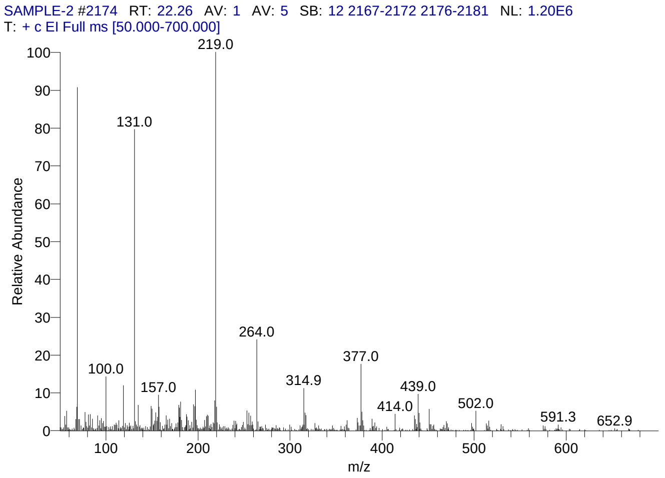


#### Library Search Results Table

| Compound Name | RT | Molecular Formula | Cas # |
|---|---|---|---|
| 18-Norcholest-17(20),24-dien-21-oic acid, 16-acetoxy-4,8,14-trimethyl-3,11-dioxo-, methyl ester | 22.26 | C32H46O6 | NA |
| Cholestano[7,8-a]cyclobutane, 3-methoxy-6-oxo-2'-methylene- | 22.26 | C31H50O2 | NA |
| Olean-12-ene-3,16,21,22,28-pentol, 21-(2-methyl-2-butenoate), [3á,16à,21á(Z),22à]- | 22.26 | C35H56O6 | 20089-98-9 |

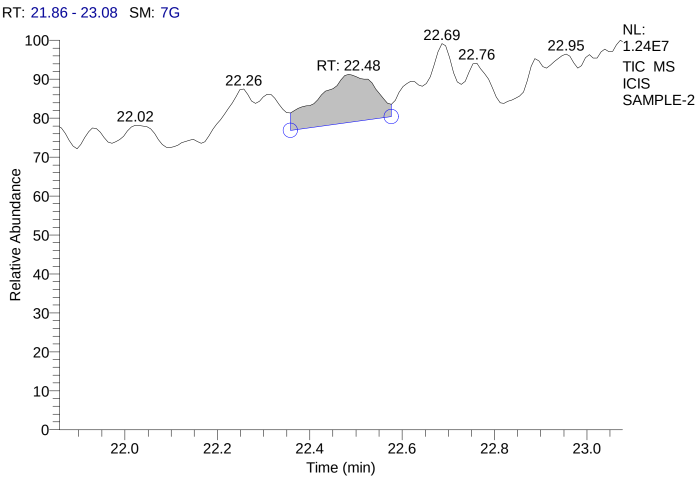


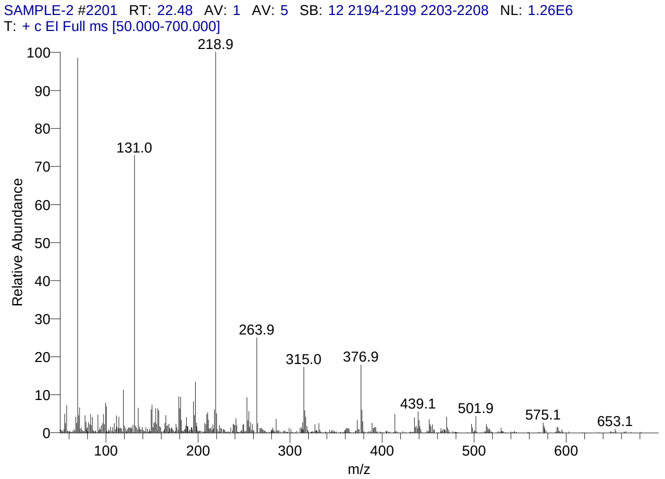


#### Library Search Results Table

| Compound Name | RT | Molecular Formula | Cas # |
|---|---|---|---|
| Cholesta-5,7,9(11)-trien-3-ol, 4,4-dimethyl-, (3á)- | 22.48 | C29H46O | 53296-72-3 |
| Oleic acid, eicosyl ester | 22.48 | C38H74O2 | 22393-88-0 |
| 3à,5à-Cyclo-ergosta-7,9(11),22t-triene-6á-ol | 22.48 | C28H42O | 118978-72-6 |

### CIL/ SAIF Panjab University Chandigarh

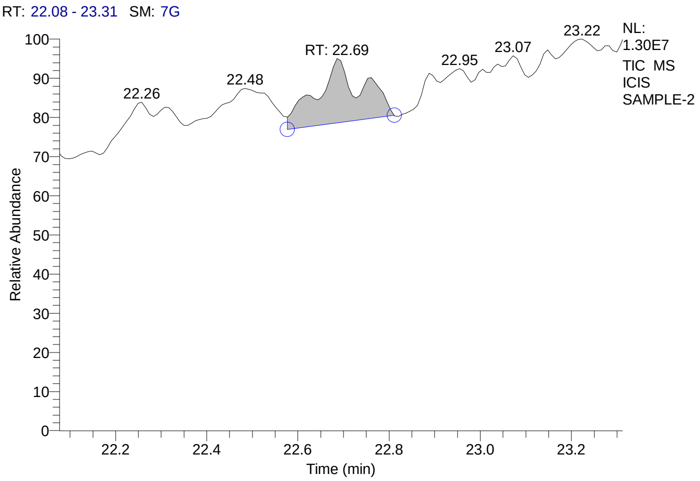


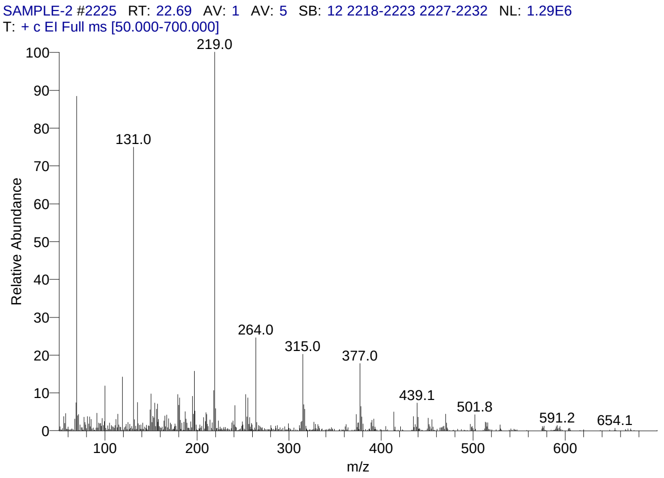


#### Library Search Results Table

| Compound Name | RT | Molecular Formula | Cas # |
|---|---|---|---|
| Cholestano[7,8-a]cyclobutane, 3-methoxy-6-oxo-2'-methylene- | 22.69 | C31H50O2 | NA |
| Cholesta-5,7,9(11)-trien-3-ol, 4,4-dimethyl-, (3á)- | 22.69 | C29H46O | 53296-72-3 |
| 1H-Cyclopropa[3,4]benz[1,2-e]azulene-5,7b,9,9a-tetrol, 1a,1b,4,4a,5,7a,8,9-octahydro-3-(hydroxymethyl)-1,1,6,8-tetramethyl-, 9,9a-diacetate, [1aR-(1aà,1bá,4aá,5á,7aà,7bà,8à,9á,9aà)]- | 22.69 | C24H34O7 | 77508-65-7 |

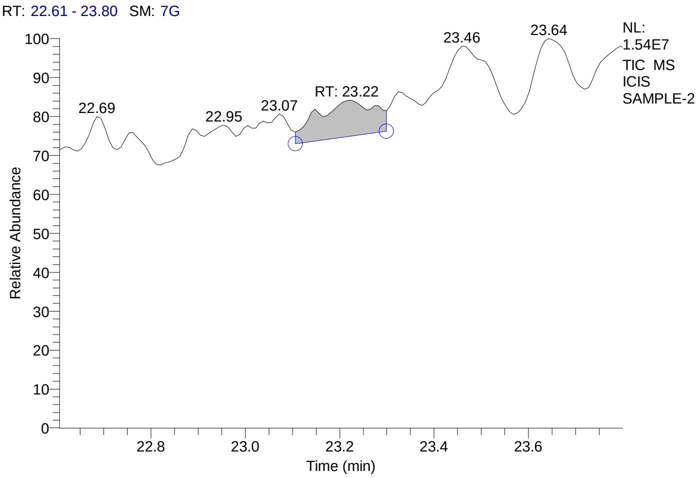


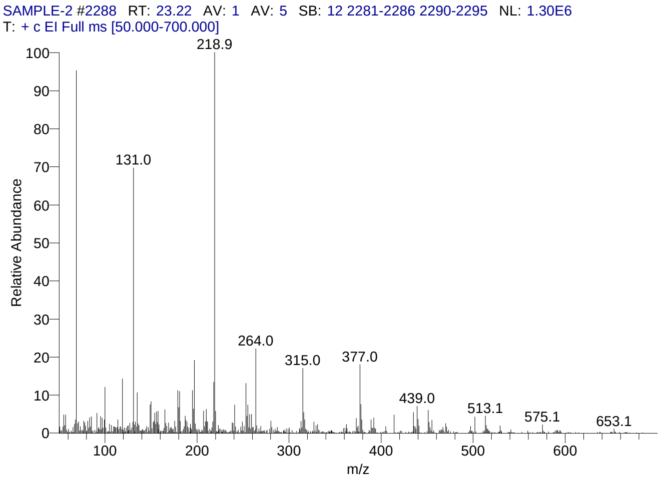


#### Library Search Results Table

| Compound Name | RT | Molecular Formula | Cas # |
|---|---|---|---|
| 18-Norcholest-17(20),24-dien-21-oic acid, 16-acetoxy-4,8,14-trimethyl-3,11-dioxo-, methyl ester | 23.22 | C32H46O6 | NA |
| Olean-12-ene-3,16,21,22,23,28-hexol, (3á,4à,16à,21á,22à)- | 23.22 | C30H50O6 | 13844-22-9 |
| Cholestano[7,8-a]cyclobutane, 3-methoxy-6-oxo-2'-methylene- | 23.22 | C31H50O2 | NA |

### CIL/ SAIF Panjab University Chandigarh

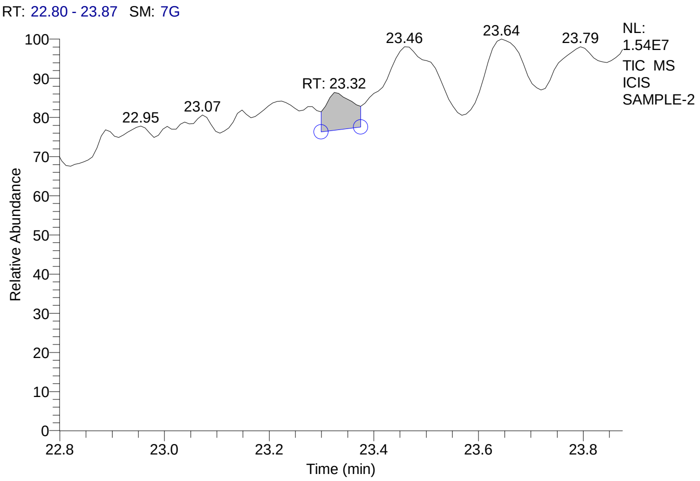


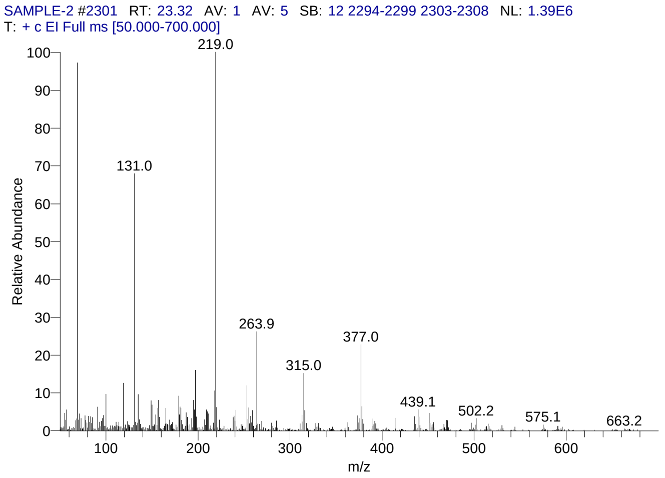


#### Library Search Results Table

| Compound Name | RT | Molecular Formula | Cas # |
|---|---|---|---|
| Flurandrenolide | 23.32 | C24H33FO6 | 1524-88-5 |
| 18-Norcholest-17(20),24-dien-21-oic acid, 16-acetoxy-4,8,14-trimethyl-3,11-dioxo-, methyl ester | 23.32 | C32H46O6 | NA |
| Oleic acid, eicosyl ester | 23.32 | C38H74O2 | 22393-88-0 |

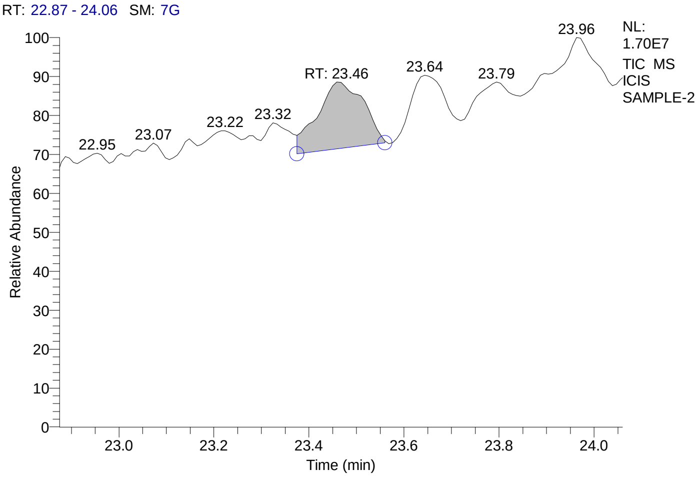


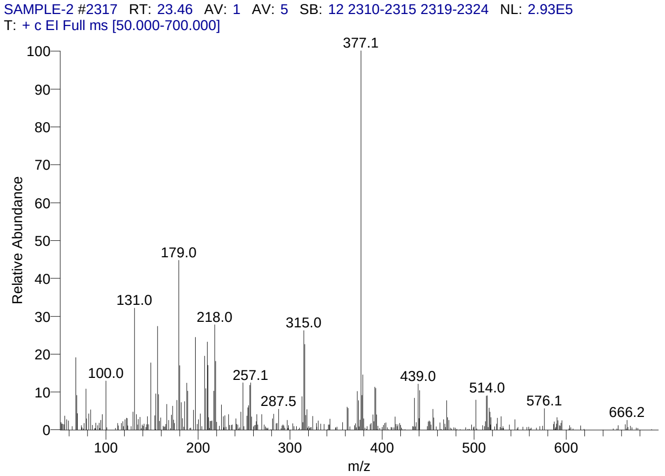


#### Library Search Results Table

| Compound Name | RT | Molecular Formula | Cas # |
|---|---|---|---|
| Cholesta-5,7-dien-3-ol, 4,4-dimethyl-, (3á)- | 23.46 | C29H48O | 53296-71-2 |
| 3-[18-(3-Hydroxy-propyl)-3,3,7,12,17-pentamethyl-2,3,22,24-tetrahydro-porphin-2-yl]propan-1-ol | 23.46 | C31H38N4O2 | NA |
| Cholesta-5,7,9(11)-trien-3-ol, 4,4-dimethyl-, (3á)- | 23.46 | C29H46O | 53296-72-3 |

### CIL/ SAIF Panjab University Chandigarh

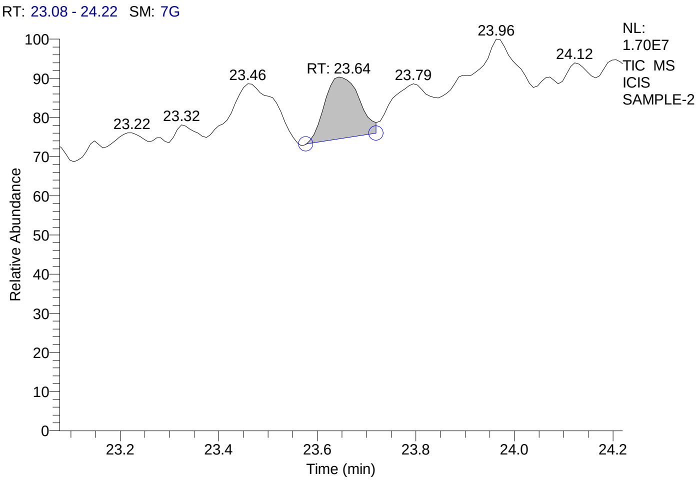


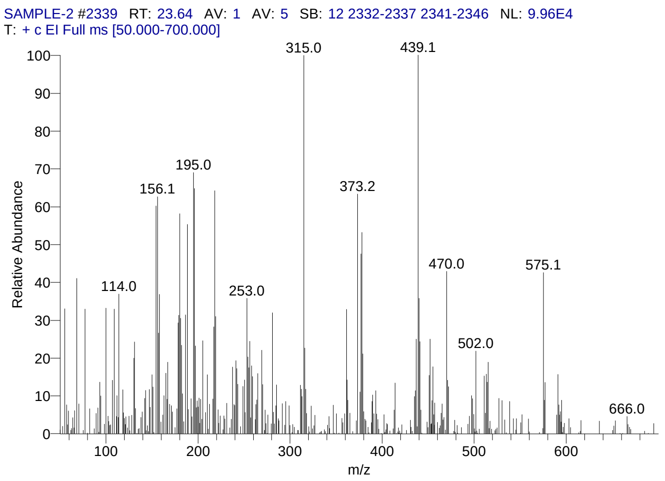


#### Library Search Results Table

| Compound Name | RT | Molecular Formula | Cas # |
|---|---|---|---|
| 3-[18-(3-Hydroxy-propyl)-3,3,7,12,17-pentamethyl-2,3,22,24-tetrahydro-porphin-2-yl]propan-1-ol | 23.64 | C31H38N4O2 | NA |
| Cholestano[7,8-a]cyclobutane, 3-methoxy-6-oxo-2'-methylene- | 23.64 | C31H50O2 | NA |
| Pregn-4-en-18-al, 3,20-bis(methoxyimino)-11,21-bis[(trimethylsilyl)oxy]-, O-methyloxime, (11á,17à)- | 23.64 | C30H53N3O5Si2 | 69854-81-5 |

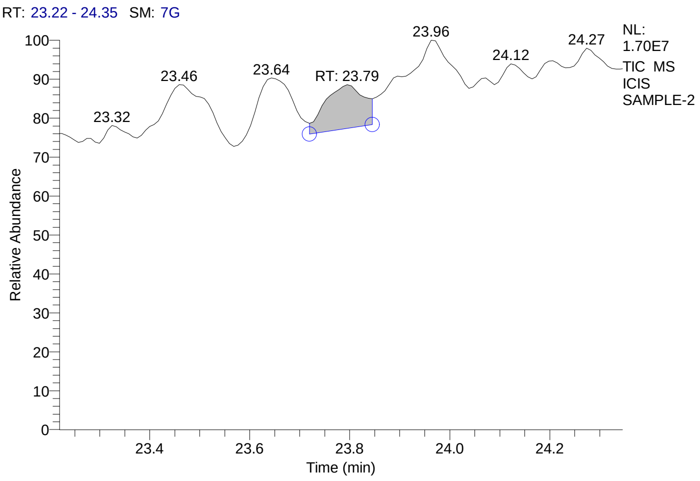


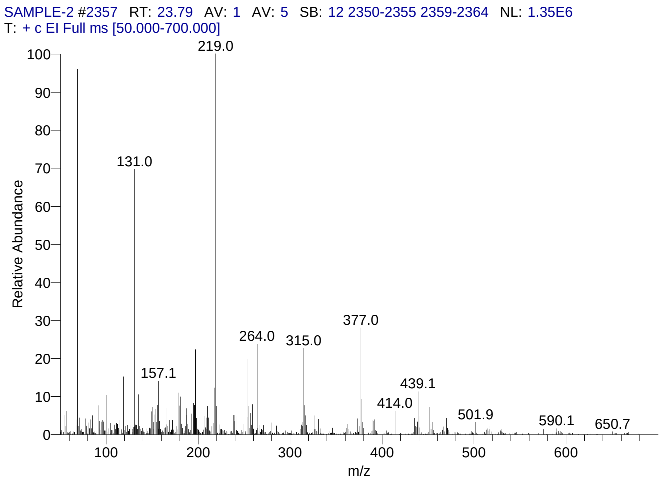


#### Library Search Results Table

| Compound Name | RT | Molecular Formula | Cas # |
|---|---|---|---|
| Flurandrenolide | 23.79 | C24H33FO6 | 1524-88-5 |
| Cholesta-5,7,9(11)-trien-3-ol, 4,4-dimethyl-, (3á)- | 23.79 | C29H46O | 53296-72-3 |
| 3à,5à-Cyclo-ergosta-7,9(11),22t-triene-6á-ol | 23.79 | C28H42O | 118978-72-6 |

### CIL/ SAIF Panjab University Chandigarh

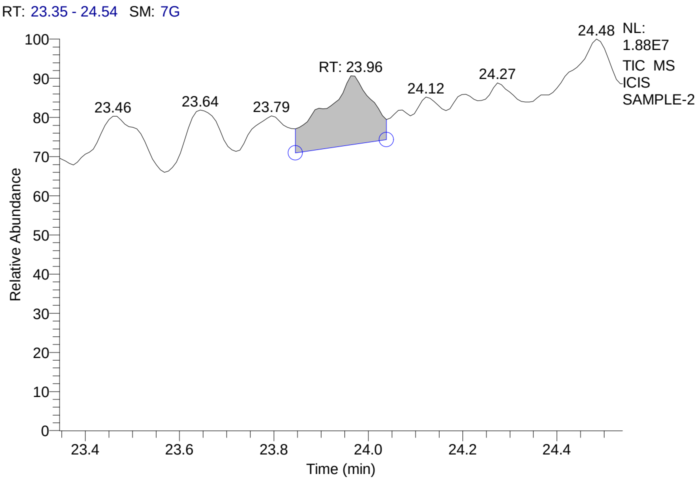


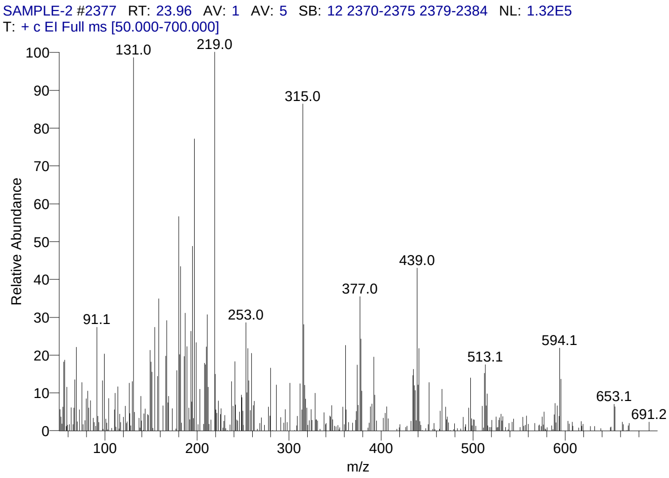


#### Library Search Results Table

| Compound Name | RT | Molecular Formula | Cas # |
|---|---|---|---|
| Urs-9(11)-en-12-one-28-oic acid, 3-acetoxy-, methyl ester (14á,20á) | 23.96 | C33H50O5 | NA |
| 3-[18-(3-Hydroxy-propyl)-3,3,7,12,17-pentamethyl-2,3,22,24-tetrahydro-porphin-2-yl]propan-1-ol | 23.96 | C31H38N4O2 | NA |
| 4'-Apo-á,.psi.-carotenoic acid, methyl ester | 23.96 | C36H48O2 | 5389-78-6 |

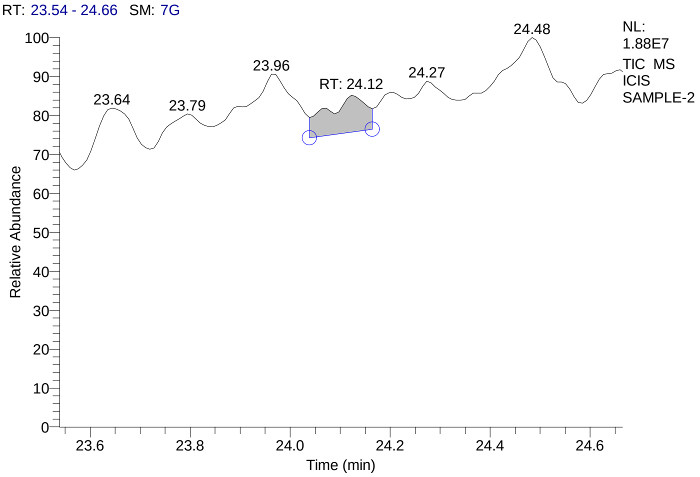


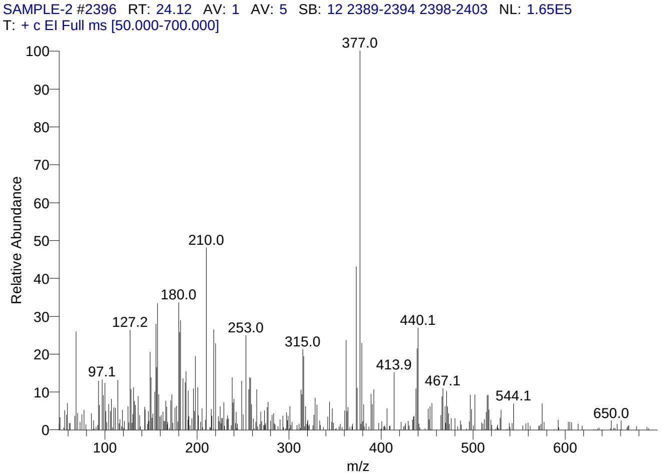


#### Library Search Results Table

| Compound Name | RT | Molecular Formula | Cas # |
|---|---|---|---|
| 3-[18-(3-Hydroxy-propyl)-3,3,7,12,17-pentamethyl-2,3,22,24-tetrahydro-porphin-2-yl]propan-1-ol | 24.12 | C31H38N4O2 | NA |
| 4,6-Bis(4-chloro-3-(trifluoromethyl)phenoxy)-2-(methylthio)pyrimidine | 24.12 | C19H10Cl2F6N2O2S | NA |
| D-Glucopyranoside, (3á,22à,25S)-22,25-epoxy-3-methoxyfurost-5-en-26-yl 2,3,4,6-tetra-O-methyl- | 24.12 | C38H62O9 | 56196-23-7 |

### CIL/ SAIF Panjab University Chandigarh

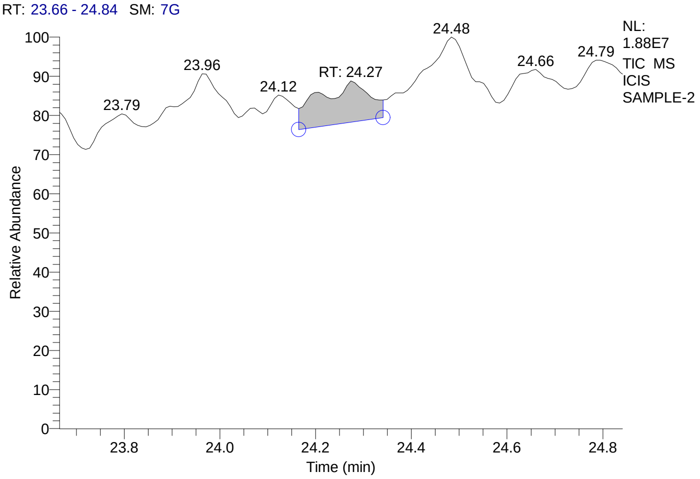


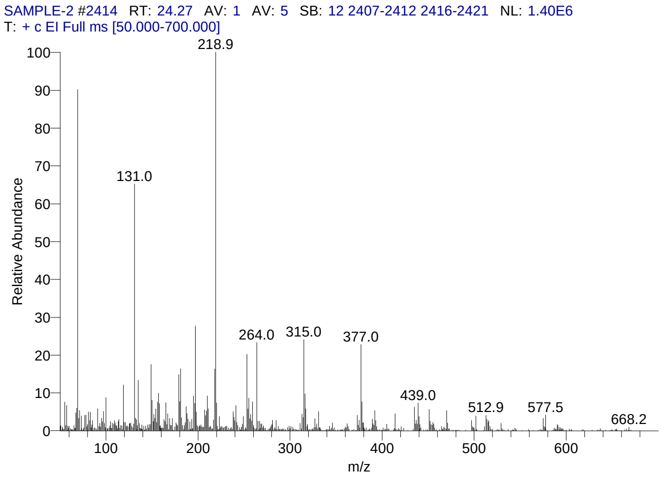


#### Library Search Results Table

| Compound Name | RT | Molecular Formula | Cas # |
|---|---|---|---|
| Flurandrenolide | 24.27 | C24H33FO6 | 1524-88-5 |
| Cholestano[7,8-a]cyclobutane, 3-methoxy-6-oxo-2'-methylene- | 24.27 | C31H50O2 | NA |
| Cholesta-5,7,9(11)-trien-3-ol, 4,4-dimethyl-, (3á)- | 24.27 | C29H46O | 53296-72-3 |

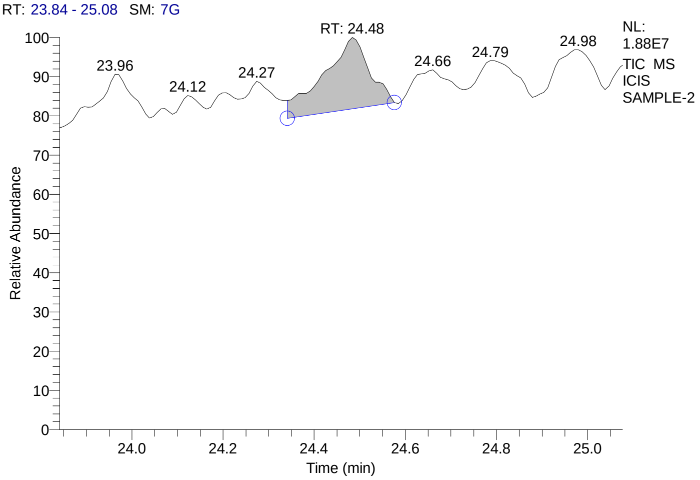


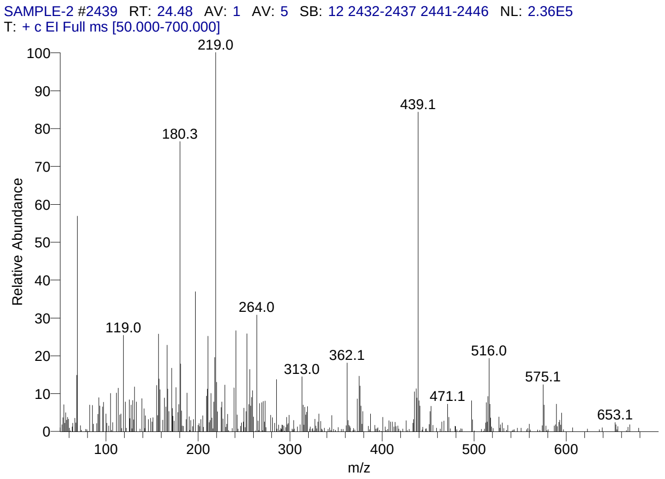


#### Library Search Results Table

| Compound Name | RT | Molecular Formula | Cas # |
|---|---|---|---|
| Cyclotrisiloxane, hexaphenyl- | 24.48 | C36H30O3Si3 | 512-63-0 |
| 5H-Cyclopropa(3,4)benz(1,2-e)azulen-5-one, 1,1a-à,1b-á,4,4a,7a-à,7b,8,9,9a-decahydro-7b-à,9-á, 9a-à-trihydroxy-3-hydroxymethyl-1,1,6,8-à-tetrameth yl-4a-methoxy-, 9,9a-didecanoate | 24.48 | C41H66O8 | 54870-24-5 |
| 3-[18-(3-Hydroxy-propyl)-3,3,7,12,17-pentamethyl-2,3,22,24-tetrahydro-porphin-2-yl]propan-1-ol | 24.48 | C31H38N4O2 | NA |

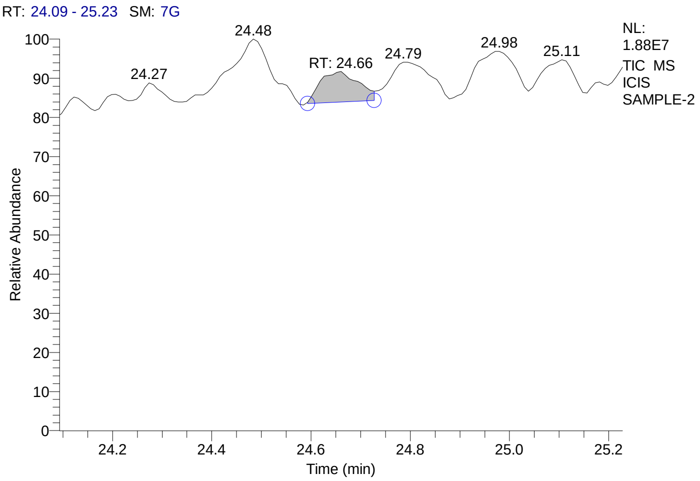


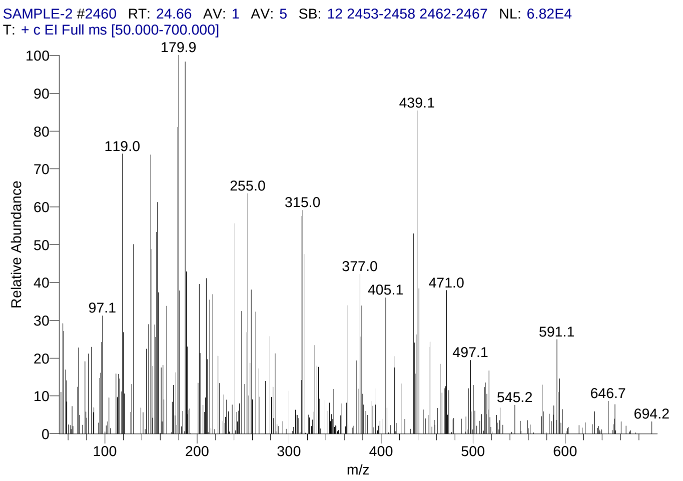


#### Library Search Results Table

| Compound Name | RT | Molecular Formula | Cas # |
|---|---|---|---|
| 3-[18-(3-Hydroxy-propyl)-3,3,7,12,17-pentamethyl-2,3,22,24-tetrahydro-porphin-2-yl]propan-1-ol | 24.66 | C31H38N4O2 | NA |
| Cholestano[7,8-a]cyclobutane, 3-methoxy-6-oxo-2'-methylene- | 24.66 | C31H50O2 | NA |
| Milbemycin B, 5-demethoxy-5-one-6,28-anhydro-25-ethyl-4-methyl-13-chloro-oxime | 24.66 | C32H44ClNO7 | NA |

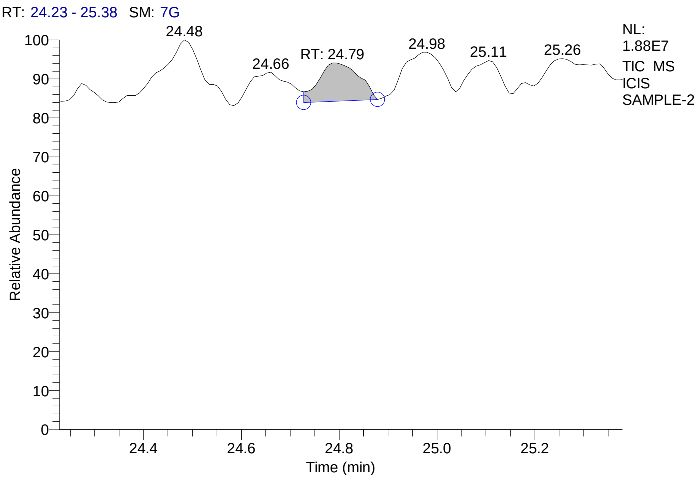


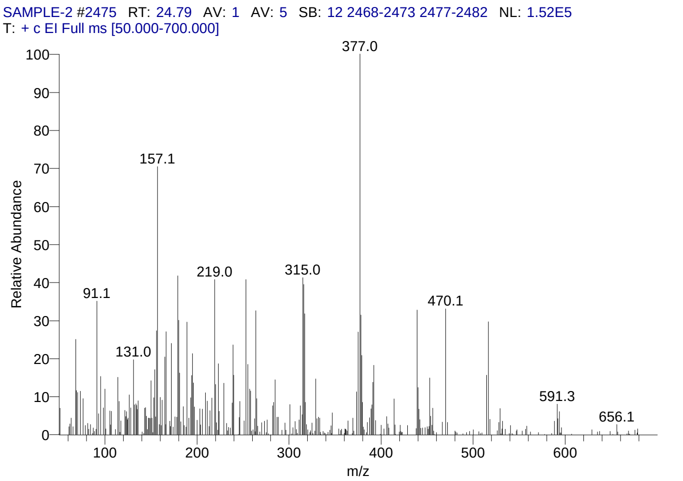


#### Library Search Results Table

| Compound Name | RT | Molecular Formula | Cas # |
|---|---|---|---|
| Cholesta-5,7,9(11)-trien-3-ol, 4,4-dimethyl-, (3á)- | 24.79 | C29H46O | 53296-72-3 |
| Flurandrenolide | 24.79 | C24H33FO6 | 1524-88-5 |
| Cyclotrisiloxane, 2,4,6-trimethyl-2,4,6-triphenyl- | 24.79 | C21H24O3Si3 | 546-45-2 |

### CIL/ SAIF Panjab University Chandigarh

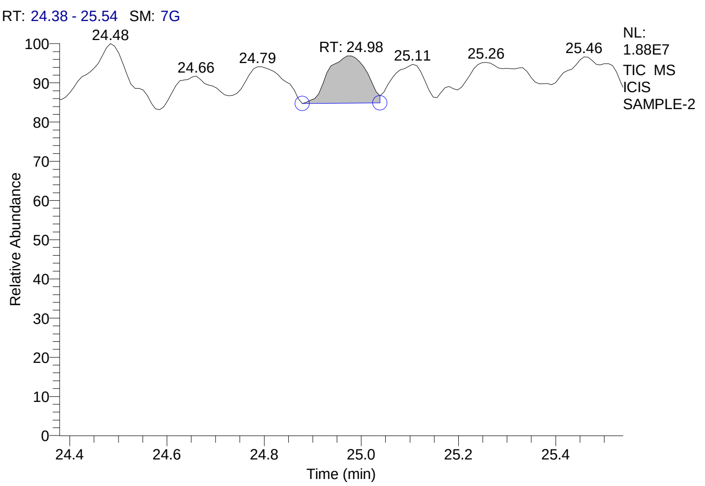


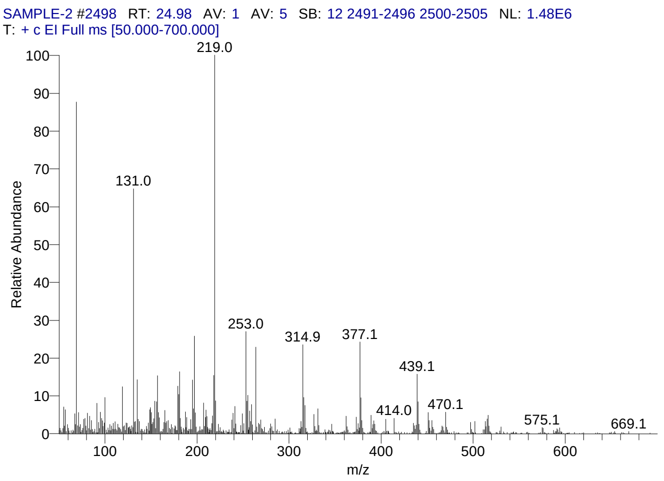


#### Library Search Results Table

| Compound Name | RT | Molecular Formula | Cas # |
|---|---|---|---|
| Cholesta-5,7,9(11)-trien-3-ol, 4,4-dimethyl-, (3á)- | 24.98 | C29H46O | 53296-72-3 |
| Cholestano[7,8-a]cyclobutane, 3-methoxy-6-oxo-2'-methylene- | 24.98 | C31H50O2 | NA |
| 3à,5à-Cyclo-ergosta-7,9(11),22t-triene-6á-ol | 24.98 | C28H42O | 118978-72-6 |

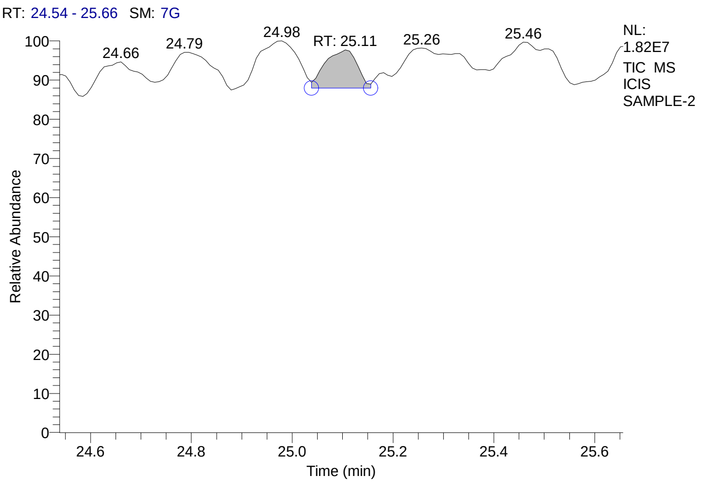


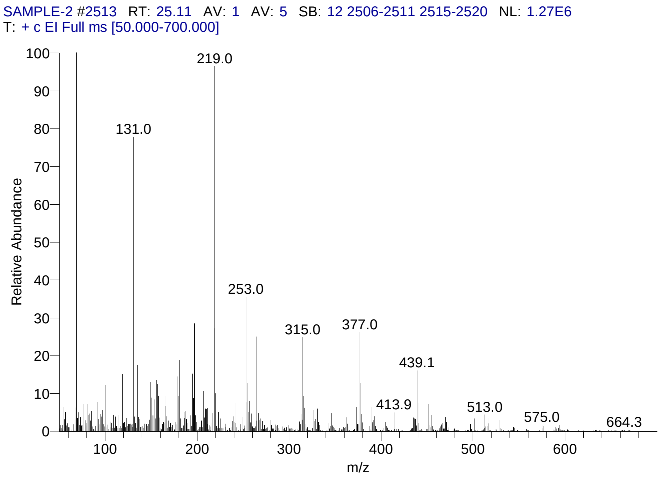


#### Library Search Results Table

| Compound Name | RT | Molecular Formula | Cas # |
|---|---|---|---|
| Cholestano[7,8-a]cyclobutane, 3-methoxy-6-oxo-2'-methylene- | 25.11 | C31H50O2 | NA |
| Cholesta-5,7,9(11)-trien-3-ol, 4,4-dimethyl-, (3á)- | 25.11 | C29H46O | 53296-72-3 |
| 3à,5à-Cyclo-ergosta-7,9(11),22t-triene-6á-ol | 25.11 | C28H42O | 118978-72-6 |

### CIL/ SAIF Panjab University Chandigarh

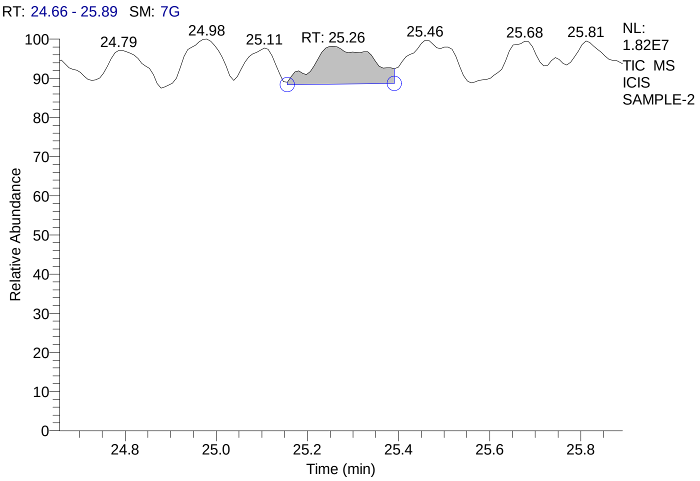


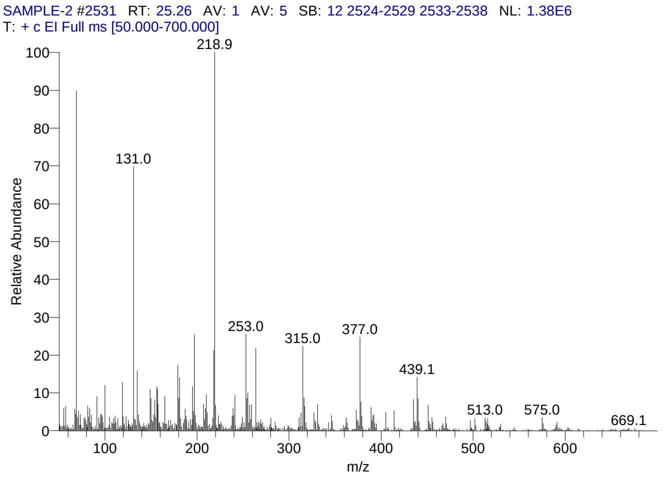


#### Library Search Results Table

| Compound Name | RT | Molecular Formula | Cas # |
|---|---|---|---|
| Cholestano[7,8-a]cyclobutane, 3-methoxy-6-oxo-2'-methylene- | 25.26 | C31H50O2 | NA |
| (14-methoxy-5,13,13-trimethyl-8-oxo-9,15-dioxapentacyclo[12.2.2.1(2,6).0(1,2).0(10,19)]nonadec-11-en-5-yl)methyl methylcarbonate | 25.26 | C24H34O7 | NA |
| Cholesta-5,7,9(11)-trien-3-ol, 4,4-dimethyl-, (3á)- | 25.26 | C29H46O | 53296-72-3 |

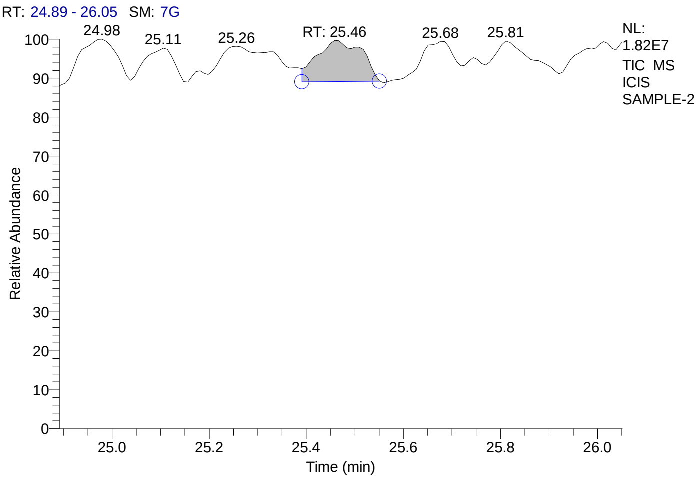


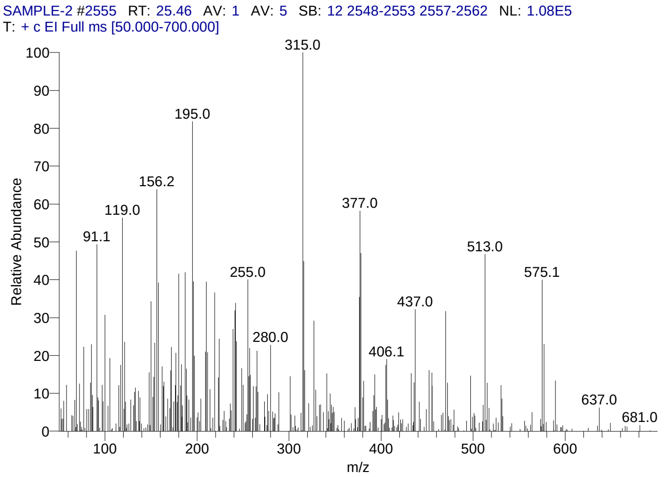


#### Library Search Results Table

| Compound Name | RT | Molecular Formula | Cas # |
|---|---|---|---|
| Urs-9(11)-en-12-one-28-oic acid, 3-acetoxy-, methyl ester (14á,20á) | 25.46 | C33H50O5 | NA |
| Cholesta-5,7,9(11)-trien-3-ol, 4,4-dimethyl-, (3á)- | 25.46 | C29H46O | 53296-72-3 |
| Cyclotrisiloxane, 2,4,6-trimethyl-2,4,6-triphenyl- | 25.46 | C21H24O3Si3 | 546-45-2 |

### CIL/ SAIF Panjab University Chandigarh

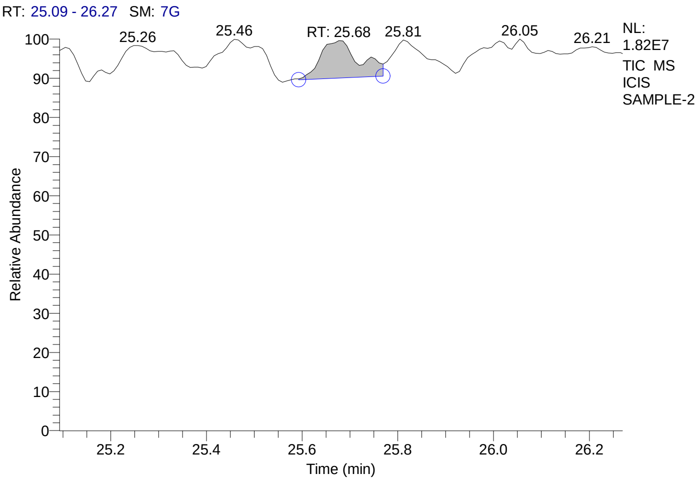


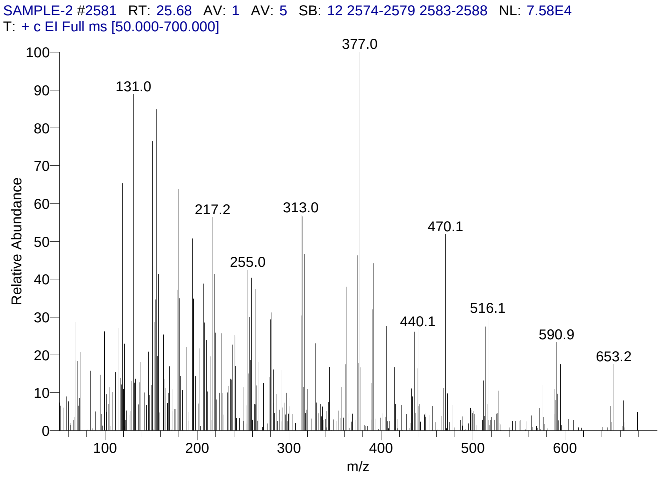


#### Library Search Results Table

| Compound Name | RT | Molecular Formula | Cas # |
|---|---|---|---|
| Milbemycin B, 5-demethoxy-5-one-6,28-anhydro-25-ethyl-4-methyl-13-chloro-oxime | 25.68 | C32H44ClNO7 | NA |
| Rhodoxanthin | 25.68 | C40H50O2 | 116-30-3 |
| Pregn-4-en-18-al, 3,20-bis(methoxyimino)-11,21-bis[(trimethylsilyl)oxy]-, O-methyloxime, (11á,17à)- | 25.68 | C30H53N3O5Si2 | 69854-81-5 |

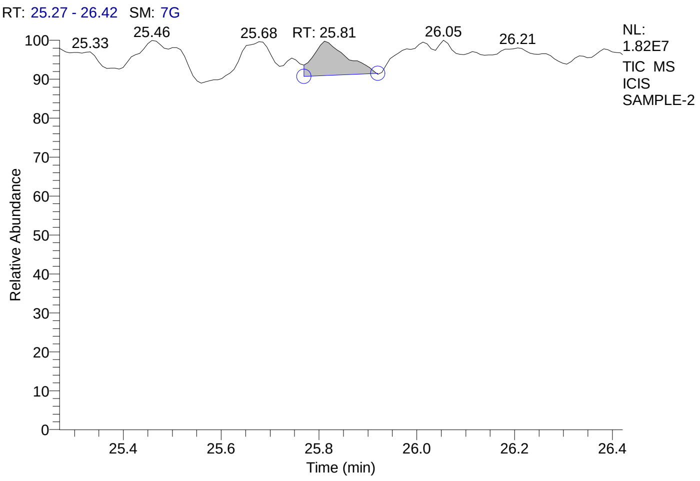


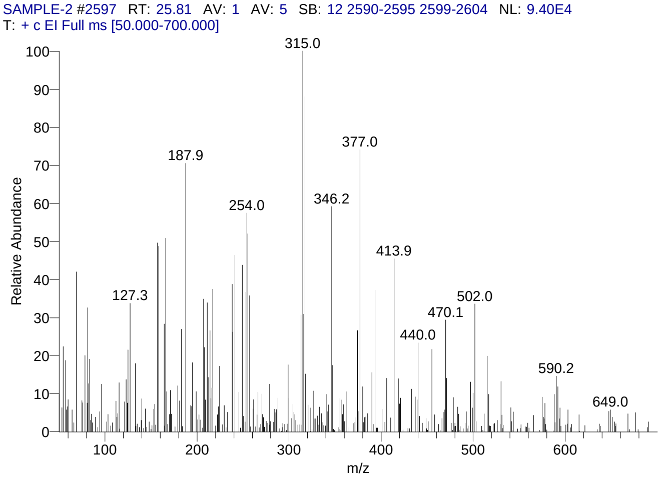


#### Library Search Results Table

| Compound Name | RT | Molecular Formula | Cas # |
|---|---|---|---|
| Rhodoxanthin | 25.81 | C40H50O2 | 116-30-3 |
| 2-(5-(5-[Cyano-(9,9-dimethyl-1,4-dioxa-7-aza-spiro[4.4]non-7-en-8-yl)-methylene]-3,3-dimethylpyrrolidin-2-ylidenemethyl)-3,3-dimethyl-ë1-pyrrolin-5-ylidenemethyl-4,4,5-trimethyl-ë1-pyrroline-5-carbonitrile] | 25.81 | C32H42N6O2 | NA |
| Dexamethasone-21-acetate | 25.81 | C24H31FO6 | 1177-87-3 |

### CIL/ SAIF Panjab University Chandigarh

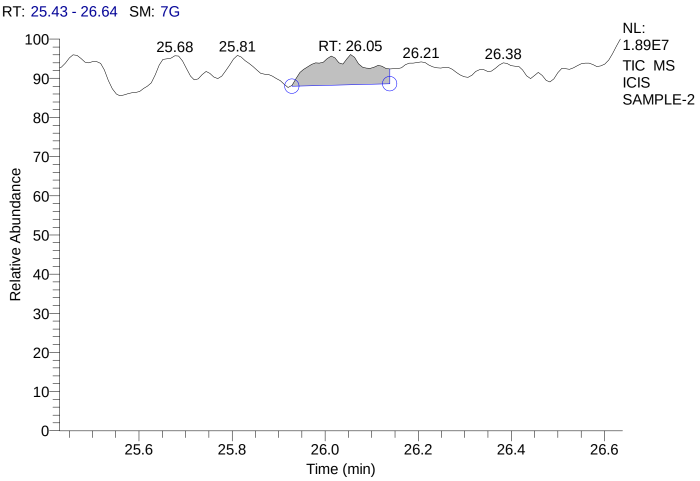


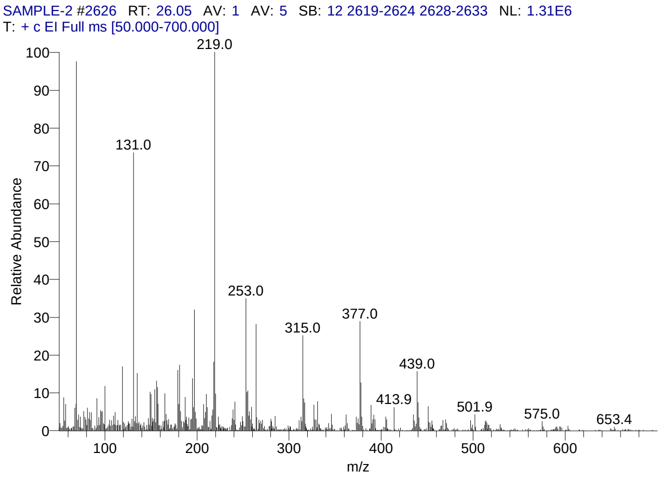


#### Library Search Results Table

| Compound Name | RT | Molecular Formula | Cas # |
|---|---|---|---|
| Cholestano[7,8-a]cyclobutane, 3-methoxy-6-oxo-2'-methylene- | 26.05 | C31H50O2 | NA |
| 3à,5à-Cyclo-ergosta-7,9(11),22t-triene-6á-ol | 26.05 | C28H42O | 118978-72-6 |
| Cholesta-5,7,9(11)-trien-3-ol, 4,4-dimethyl-, (3á)- | 26.05 | C29H46O | 53296-72-3 |

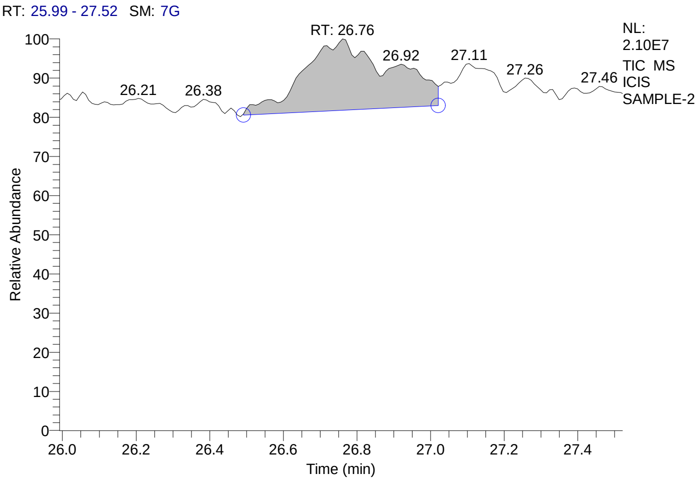


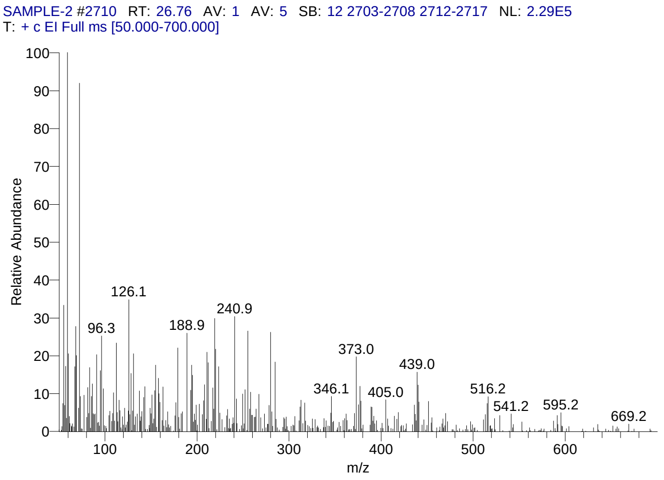


#### Library Search Results Table

| Compound Name | RT | Molecular Formula | Cas # |
|---|---|---|---|
| Cholestane, 3,5-dichloro-6-nitro-, (3á,5à,6á)- | 26.76 | C27H45Cl2NO2 | 15505-92-7 |
| 5H-Cyclopropa(3,4)benz(1,2-e)azulen-5-one, 1,1a-à,1b-á,4,4a,7a-à,7b,8,9,9a-decahydro-7b-à,9-á, 9a-à-trihydroxy-3-hydroxymethyl-1,1,6,8-à-tetramethyl-4a-methoxy-, 9,9a-didecanoate | 26.76 | C41H66O8 | 54870-24-5 |
| Hexadecanoic acid, 1a,2,5,5a,6,9,10,10a-octahydro-5a-hydroxy-4-(hydroxymethyl)-1,1,7,9-tetramethyl-6,11-dioxo-1H-2,8a-methanocyclopenta[a]cyclopropa[e]cyclodecen-5-yl ester, [1aR-(1aà,2à,5á,5aá,8aà,9à,10aà)]- | 26.76 | C36H56O6 | 77508-69-1 |

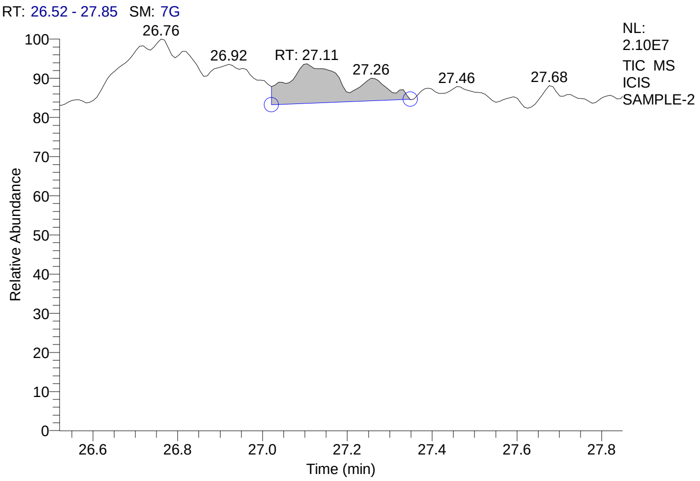


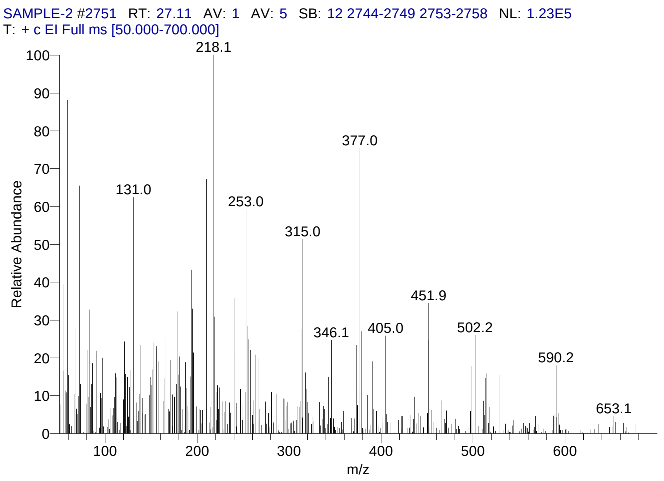


#### Library Search Results Table

| Compound Name | RT | Molecular Formula | Cas # |
|---|---|---|---|
| Milbemycin B, 5-demethoxy-5-one-6,28-anhydro-25-ethyl-4-methyl -13-chloro-oxime | 27.11 | C32H44ClNO7 | NA |
| Urs-9(11)-en-12-one-28-oic acid, 3-acetoxy-, methyl ester (14á,20á) | 27.11 | C33H50O5 | NA |
| 19-Norpregna-1,3,5,7,9-pentaen-21-al, 3,17-bis[(trimethylsilyl)oxy]-, O-methyloxime, (17à)- | 27.11 | C27H41NO3Si2 | 74299-04-0 |

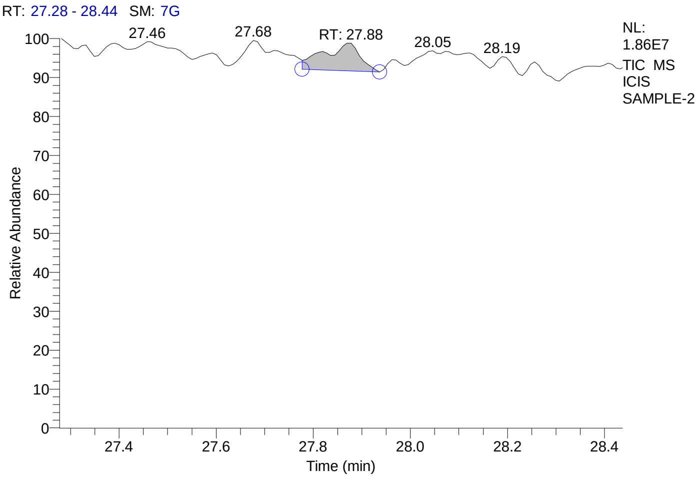


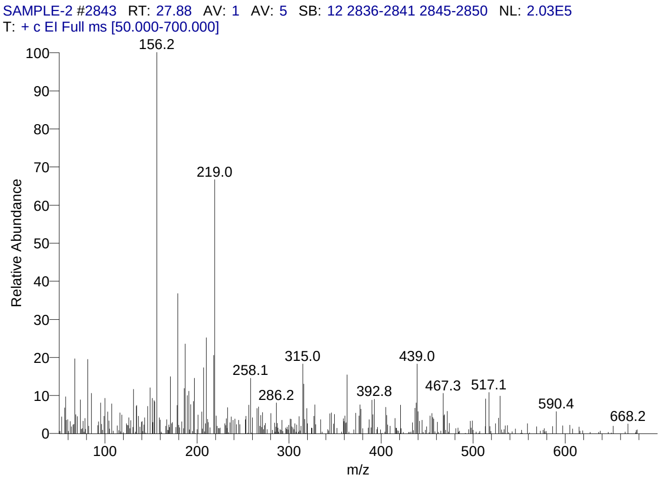


#### Library Search Results Table

| Compound Name | RT | Molecular Formula | Cas # |
|---|---|---|---|
| 3-[18-(3-Hydroxy-propyl)-3,3,7,12,17-pentamethyl-2,3,22,24-tetrahydro-porphin-2-yl]propan-1-ol | 27.88 | C31H38N4O2 | NA |
| Pregn-4-ene-3,11,20-trione, 6,17,21-tris[(trimethylsilyl)oxy]-, 3,20-bis(O-methyloxime), (6á)- | 27.88 | C32H58N2O6Si3 | 57326-06-4 |
| 18-Norcholest-17(20),24-dien-21-oic acid, 16-acetoxy-4,8,14-trimethyl-3,11-dioxo-, methyl ester | 27.88 | C32H46O6 | NA |

### CIL/ SAIF Panjab University Chandigarh

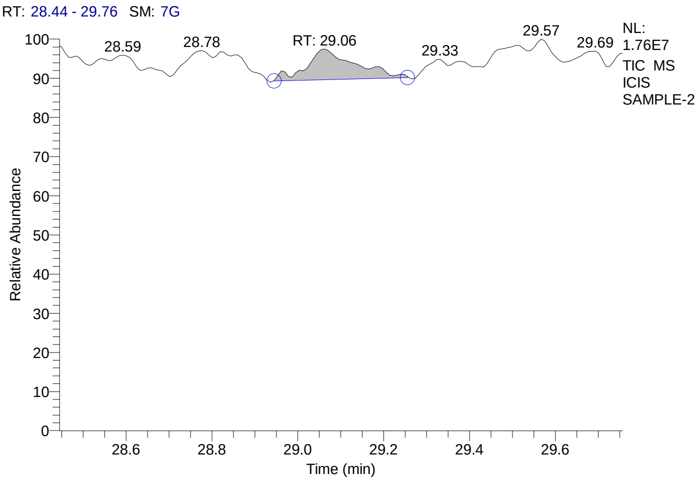


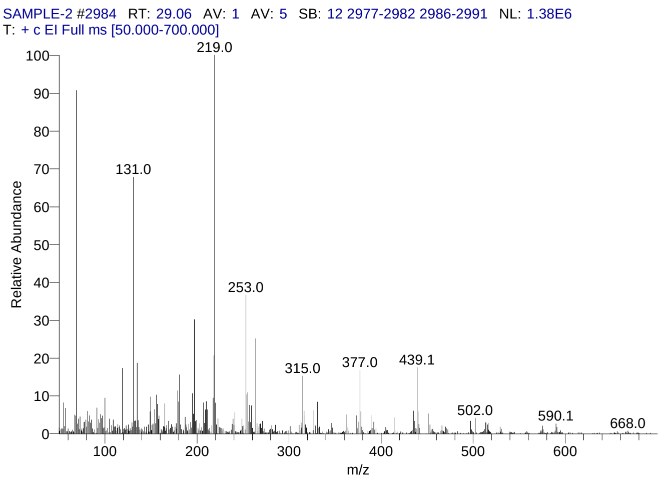


#### Library Search Results Table

| Compound Name | RT | Molecular Formula | Cas # |
|---|---|---|---|
| 3à,5à-Cyclo-ergosta-7,9(11),22t-triene-6á-ol | 29.06 | C28H42O | 118978-72-6 |
| 3-Hydroxy-1-(4-{13-[4-(3-hydroxy-3-phenylacryloyl)phenyl]tridecyl}-phenyl)-3-phenylprop-2-en-1-one | 29.06 | C43H48O4 | NA |
| 1H-Cyclopropa[3,4]benz[1,2-e]azulene-5,7b,9,9a-tetrol, 1a,1b,4,4a,5,7a,8,9-octahydro-3-(hydroxymethyl)-1,1,6,8-tetramethyl-, 9,9a-diacetate, [1aR-(1aà,1bá,4aá,5á,7aà,7bà,8à,9á,9aà)]- | 29.06 | C24H34O7 | 77508-65-7 |

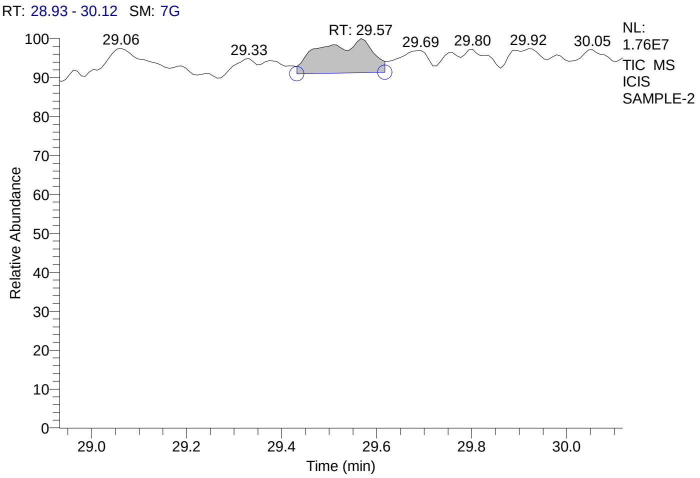


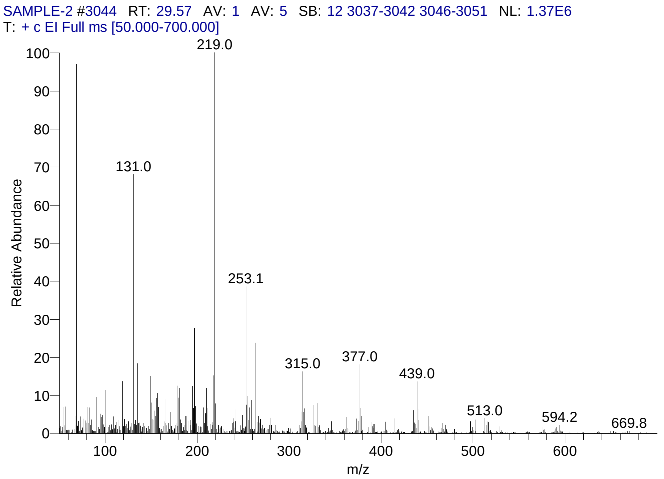


#### Library Search Results Table

| Compound Name | RT | Molecular Formula | Cas # |
|---|---|---|---|
| 3à,5à-Cyclo-ergosta-7,9(11),22t-triene-6á-ol | 29.57 | C28H42O | 118978-72-6 |
| 18-Norcholest-17(20),24-dien-21-oic acid, 16-acetoxy-4,8,14-trimethyl-3,11-dioxo-, methyl ester | 29.57 | C32H46O6 | NA |
| Cholestano[7,8-a]cyclobutane, 3-methoxy-6-oxo-2'-methylene- | 29.57 | C31H50O2 | NA |

### CIL/ SAIF Panjab University Chandigarh

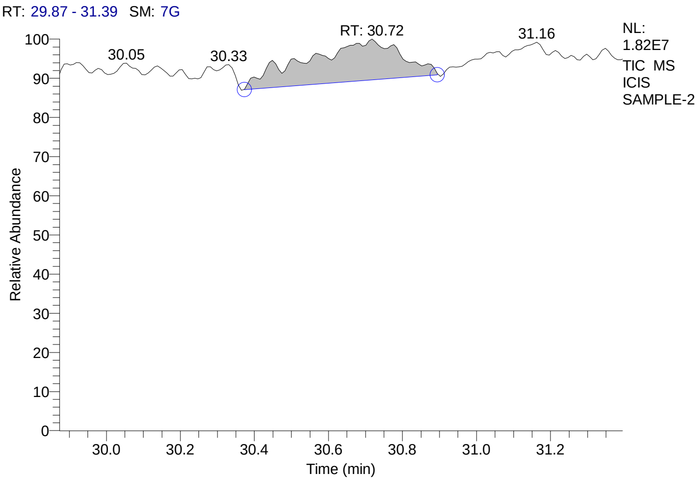


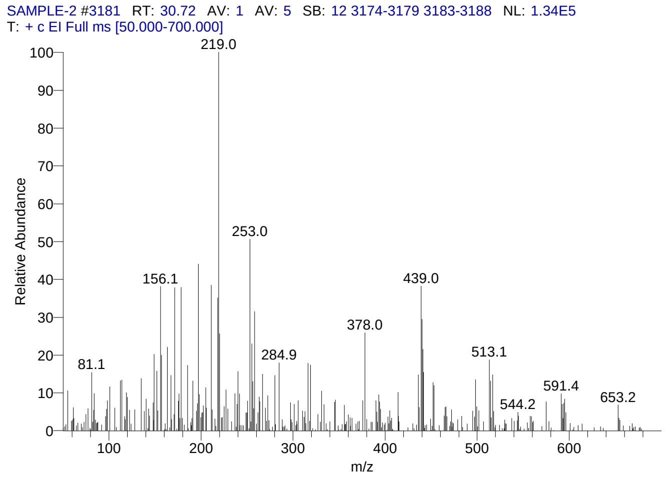


#### Library Search Results Table

| Compound Name | RT | Molecular Formula | Cas # |
|---|---|---|---|
| 3-[18-(3-Hydroxy-propyl)-3,3,7,12,17-pentamethyl-2,3,22,24-tetrahydro-porphin-2-yl]propan-1-ol | 30.72 | C31H38N4O2 | NA |
| Cyclotrisiloxane, hexaphenyl- | 30.72 | C36H30O3Si3 | 512-63-0 |
| (Vanadium, cyclopentadienyl cyclooctatetraenyl), bis | 30.72 | C26H26V2 | NA |

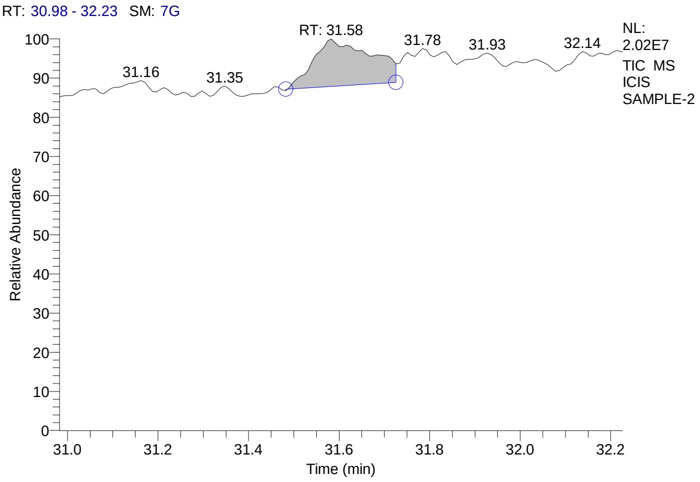


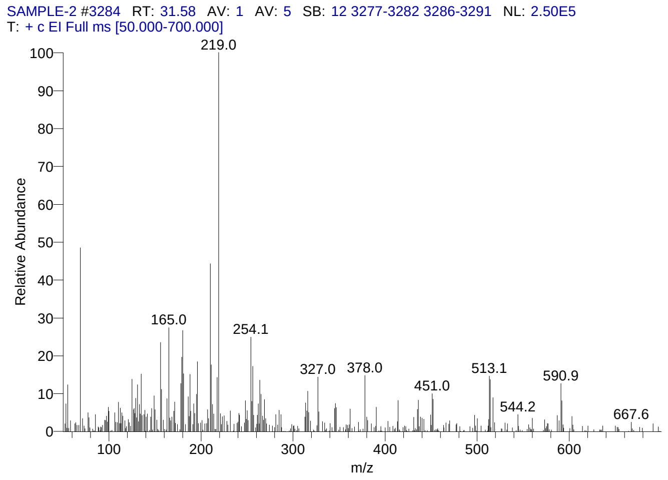


#### Library Search Results Table

| Compound Name | RT | Molecular Formula | Cas # |
|---|---|---|---|
| Milbemycin B, 5-demethoxy-5-one-6,28-anhydro-25-ethyl-4-methyl -13-chloro-oxime | 31.58 | C32H44ClNO7 | NA |
| 1'-Carboethoxy-1'-cyano-1á,2á-dihydro-17á-propion oxy-3'H-cycloprop[1,2]androsta-1,4,6-trien-3-one | 31.58 | C27H33NO5 | 75857-77-1 |
| Docosanoic acid, 1,2,3-propanetriyl ester | 31.58 | C69H134O6 | 18641-57-1 |

### CIL/ SAIF Panjab University Chandigarh

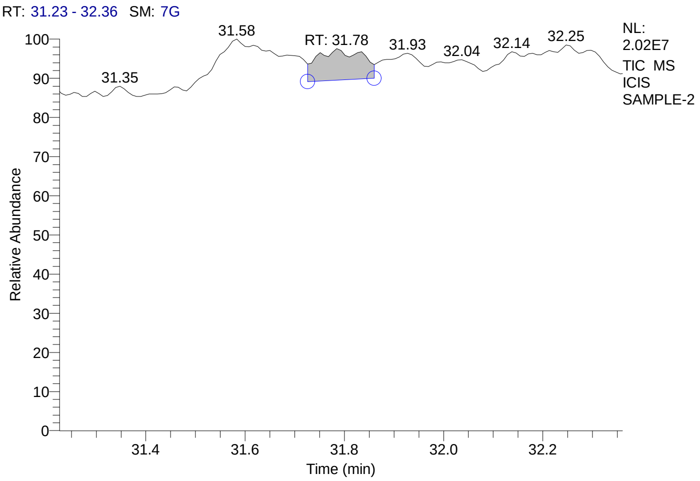


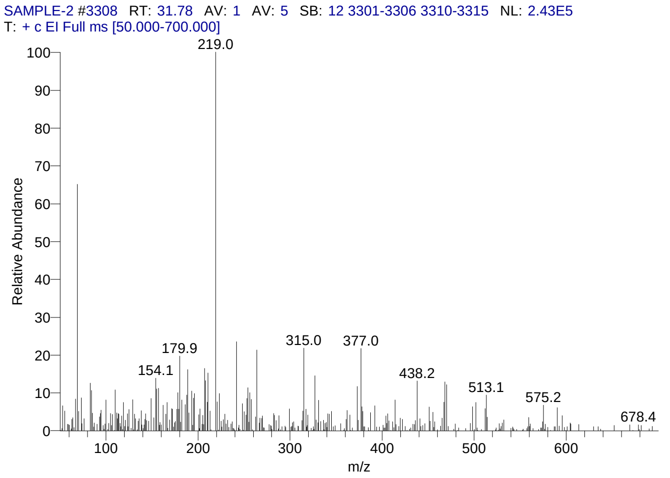


### Library Search Results Table

| Compound Name | RT | Molecular Formula | Cas # |
|---|---|---|---|
| [5-(3-Methoxymethoxy-10,13-dimethyl-2,3,4,9,10,11,12,13,14,15,16,17-dodecahydro-1H-cyclopenta[a]phenanthren-17-yl)-hex-1-ynyl]-trime | 31.78 | C30H48O2Si | NA |
| Urs-9(11)-en-12-one-28-oic acid, 3-acetoxy-, methyl ester (14á,20á) | 31.78 | C33H50O5 | NA |
| 9,19-Cyclolanostane-6,7-dione, 3-acetoxy- | 31.78 | C32H50O4 | NA |

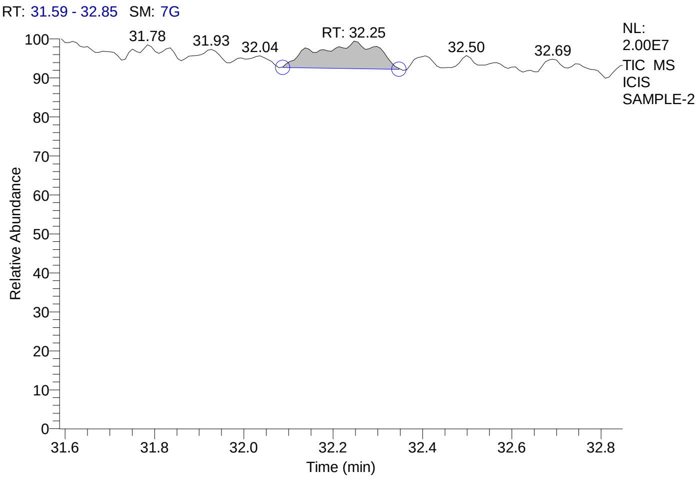


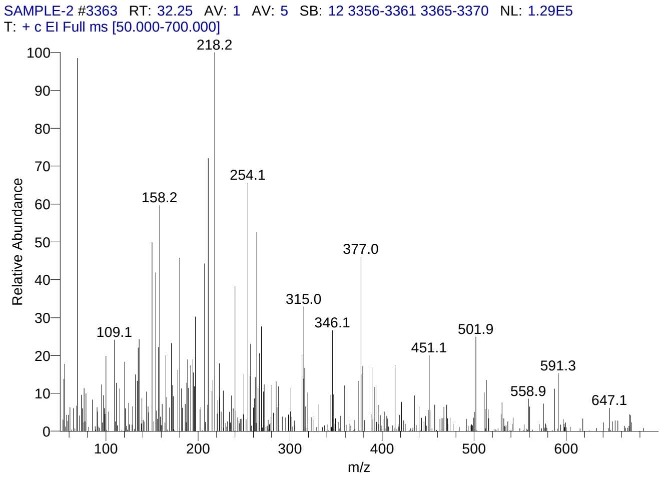


### Library Search Results Table

| Compound Name | RT | Molecular Formula | Cas # |
|---|---|---|---|
| Urs-9(11)-en-12-one-28-oic acid, 3-acetoxy-, methyl ester (14á,20á) | 32.25 | C33H50O5 | NA |
| Milbemycin B, 5-demethoxy-5-one-6,28-anhydro-25-ethyl-4-methyl-13-chloro-oxime | 32.25 | C32H44ClNO7 | NA |
| Oleic acid, eicosyl ester | 32.25 | C38H74O2 | 22393-88-0 |

### CIL/ SAIF Panjab University Chandigarh

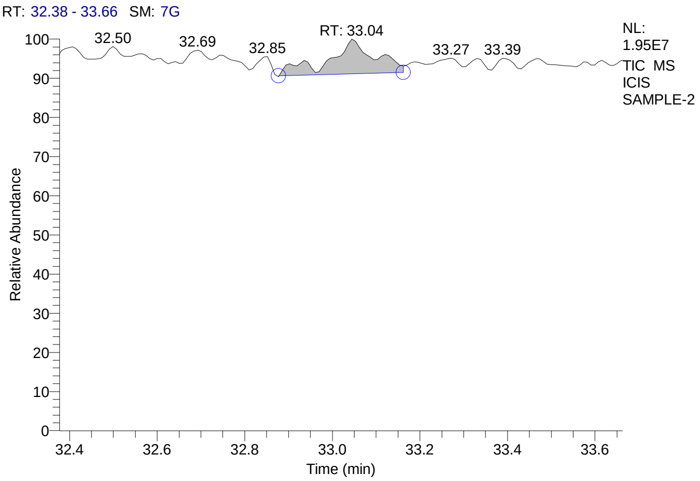


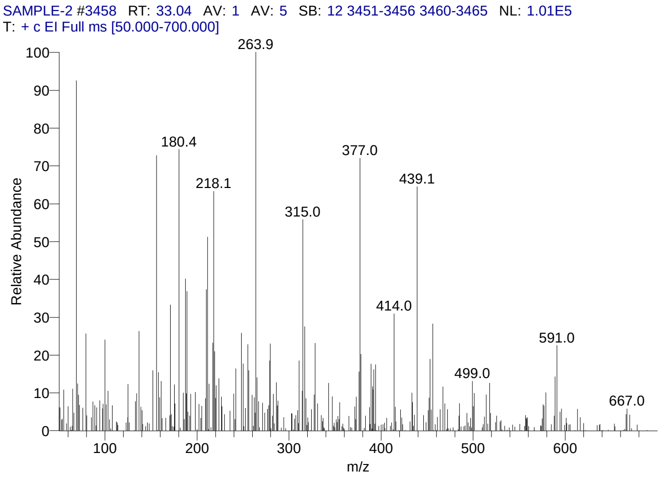


#### Library Search Results Table

| Compound Name | RT | Molecular Formula | Cas # |
|---|---|---|---|
| Cholestano[7,8-a]cyclobutane, 3-methoxy-6-oxo-2'-methylene- | 33.04 | C31H50O2 | NA |
| Rhodoxanthin | 33.04 | C40H50O2 | 116-30-3 |
| 1,4,10,13-Tetraoxa-7,16-diazacyclooctadecane, 7,16-bis(1-oxodecyl)- | 33.04 | C32H62N2O6 | 105400-04-2 |

**Supplementary Figure 7. Zone of inhibition (ZOI) of antibiotics used in anthrax control and AGE**

| Well constituents | Stock concentration (µg/ml or as indicated) | Vol. added in well (µl) | ZOI (mm) |
|---|---|---|---|
| Amoxicillin (Am) | 1000 | 20 | 28 |
| Cefixime (Ce) | 40,000 | 20 | 17 |
| **Ciprofloxacin (C)** | 100 | 20 | 25 |
| **Doxycycline (Dox)** | 20 | 20 | 31 |
| **Levofloxacin (L)** | 100 | 20 | 25 |
| **Penicillin (P)** | 800 | 2 | 40 |
| Rifampicin(R) | 90 | 20 | 12 |
| Sulfamethoxazole (S) | 160000 | 20 | - |
| Tetracycline(T) | 100 | 20 | 32 |
| Aqueous garlic extract (AGE) | 40% (w/v) | 100 | 27 |
| Water (W) | | 100 | - |

### Supplementary Table 1. Traditional use of plants as indicated in the literature

| Supplementary Table 1: Traditional use of plants used in the current study as indicated in literature | | | | |
|---|---|---|---|---|
| **S. No.** | **Name of the plant as per http://www.theplantlist.org** | **Common Name** | **Traditional use of plants as/in** | **References** |
| 1. | *Aegle marmelos* (L.) Correa | Bael, Bengal Quince | 1. Stomachic, Diarrhea, Dysentery, Stomachache<br>2. Antimicrobial and anthelmintic properties<br>3. Antidiarrheal | 1. Duke *et al.*, 2002 and references therein<br>2. Singh *et al.*, 1983; Parekh *et al.*, 2007; Venkatesan *et al.*, 2009<br>3. Mazumder *et al.*, 2006; Brijesh *et al.*, 2009 |
| 2. | *Allium cepa* L. | Onion | 1. **Anti-anthrax**<br>2. Antibacterial, cholera, dysentery, stomachache Stomachic,<br>3. Antimicrobial activity | 1. Duke *et al.*, 2008 and references therein<br>2. Duke *et al.,* 2002 and references therein<br>3. Hughes *et al.*, 1991 |
| 3. | *Allium sativum* L. | Garlic | 1. Antibacterial, cholera, dysentery, stomachache, gastroenterosis<br>2. Antimicrobial activity | 1. Duke *et al.,* 2002 and references therein<br>2. Hughes *et al.*, 1991 |
| 4. | *Azadirachta indica* A. Juss. | Neem | 1. **Anti-Anthrax:** India (Santal)<br>2. Stomachic, Diarrhea, Dysentery | 1. https://phytochem.nal.usda.gov/phytochem/ethnoPlants/show/1230<br>2. Duke *et al.,* 2002 and references therein |
| 5. | *Berberis asiatica* Roxb. ex DC. | Asian Barberry*,* Daruharidra | 1. Stomachic, Antibacterial, Dysentery, Stomachache | 1. Duke *et al.*, 2002 and references therein |
| 6. | *Coriandrum sativum* L. | Coriander; Cilantro | 1. Stomachic, Antibacterial, Diarrhea, Dysentery, stomachache | 1. Duke *et al.*, 2002 and references therein |
| 7. | *Curcuma longa* L. | Turmeric | 1. **Anti-Anthrax**: India (Santal)<br>2. Antibacterial, gastroprotective, stomachic and diarrhea<br>3. Antimicrobial | 1. https://phytochem.nal.usda.gov/phytochem/ethnoPlants/show/884<br>2. Duke *et al.*, 2002 and references therein<br>3. Eigner *et al.*, 1999 |
| 8. | *Cynodon dactylon* (L.) Pers. | Bermuda grass, Doob | 1. Diarrhea, Dysentery | 1. Chopra *et al.*, 1956; Khare, 2007 |
| 9. | *Mangifera indica* L*.* | Mango | 1. Antibacterial, Stomachic, Cholera, Dysentery, Diarrhea | 1. Duke *et al.*, 2002 and references therein |
| 10. | *Ocimum tenuiflorum* L. *(Ocimum sanctum L.)* | Holy Basil, Tulsi | 1. Antibacterial, Cholera, Dysentery, | 1. Duke *et al.*, 2002 and references therein |
| 11. | *Ocimum gratissimum* L. | Ram Tulsi | 1. Antidiarrheal<br>2. Antibacterial and antifungal | 1. Offiah *et al.*, 1999<br>2. Ndounga *et al* 1997 |
| 12. | *Morus indica L.* | Black Mulberry, Shetuta | 1. Diarrhea, Dysentery, Stomachache<br>2. Anti-inflammatory, Antipyretic activities | 1. Duke *et al.*, 2008 and references therein<br>2. Chatterjee 1983 |
| 13. | *Psidium guajava L.* | Guava | 1. Anti-bacterial, Anti-diarrheic, Dysentery, Cholera and Stomachache<br>2. Antimicrobial<br>3. Antidiarrheal | 1. Duke *et al.*, 2002 and references therein<br>2. Jaiarj *et al* 1999<br>3. Ghosh *et al.*, 1993 |
| 14. | *Zingiber officinale* Roscoe | Ginger | 1. Antibacterial, Antidiarrheal, Dysentery, Stomachache | 1. Duke *et al.*, 2002 and references therein |

**Supplementary Table 2. The aqueous extracts of different plants (40% w/v) differentially inhibited the growth of B. anthracis in agar-well diffusion assay (AWDA).**

| **Supplementary Table 2. Inhibition of *Bacillus anthracis* growth by aqueous extract of indicated plant-part** | | | | | | | |
|---|---|---|---|---|---|---|---|
| **S. No.** | **Name of the plant as per http://www.theplantlist.org (common name) - plant part used (L: Leaf; B: bulb; Fruit: F; Rhizome: R)** | **Site of collection/ Voucher No. of samples** deposited in Herbarium of Botany Department, Panjab University, Chandigarh, India **(PAN No.)** | **Zone of Inhibition (mm) after incubation for** | | | | **Volume of crude aqueous extract** |
| | | | **12 h** | **24 h** | **48 h** | **72 h** | |
| 1. | *Aegle marmelos* (L.) Correa (Bael, Bengal Quince) – L | Panjab University campus, Chandigarh (Chd.) / PAN No. 21713 | 0 | 0 | 0 | 0 | 50µl |
| 2. | *Allium cepa* L. (Onion) – B | Local market, Chd. / PAN No. 21709 | 0 | 0 | 0 | 0 | 50µl |
| 3. | *Allium sativum* L. (Garlic) – B | Local market, Chd. / PAN Nos. 21146, 21147, 21148 | 19 | 19 | 18 | 18 | 50µl |
| 4. | *Azadirachta indica* A. Juss. (Neem) – L | Panjab University campus, Chd./ PAN No. 21715 | 11 | 10 | 0 | 0 | 50µl |
| 5. | *Berberis asiatica* Roxb. ex DC. *(*Daruharidra) - L | Panjab University campus, Chd./ PAN No. 21717 | 0 | 0 | 0 | 0 | 50µl |
| 6. | *Coriandrum sativum* L. (Coriander, Cilantro) – L | Local market, Chd./ PAN No. 21710 | 0 | 0 | 0 | 0 | 50µl |
| 7. | *Curcuma longa* L. (Turmeric) – R | Local market, Chd./ PAN No. 21712 | 0 | 0 | 0 | 0 | 50µl |
| 8. | *Cynodon dactylon* (L.) Pers. (Bermuda grass, Doob) - L | Panjab University campus, Chd./ PAN No. 21711 | 0 | 0 | 0 | 0 | 50µl |
| 9. | *Mangifera indica* L.(Mango)- L | Panjab University campus, Chd./ PAN No. 21719 | 10 | 10 | 0 | 0 | 50µl |
| 10. | *Ocimum tenuiflorum* L. *(Ocimum sanctum* L.*;* Holy Basil, Tulsi) - L | Panjab University campus, Chd./ PAN No. 21720 | 0 | 0 | 0 | 0 | 50µl |
| 11. | *Ocimum gratissimum* L. (Ram Tulsi) -*L* | Panjab University campus, Chd./ PAN No. 21716 | 0 | 0 | 0 | 0 | 50µl |
| 12. | *Morus indica* L. (Black Mulberry, Shetuta) -L | Panjab University campus, Chd./ PAN No. 21714 | 0 | 0 | 0 | 0 | 50µl |
| 13. | *Psidium guajava* L. (Guvava) - L | Panjab University campus, Chd./ PAN No. 21718 | 0 | 0 | 0 | 0 | 50µl |
| 14. | *Zingiber officinale* Roscoe (Ginger) – R | Local market, Chd./ PAN No. 21721 | 0 | 0 | 0 | 0 | 50µl |
| | - **Ctrl** (Solvent control - Ultrapure Water) | | 0 | 0 | 0 | 0 | 50µl |
| | **+ Ctrl** (Rifampicin 8µg) | | 18 | 18 | 18 | 17 | - |
